# Ancient DNA reveals matrilineal organisation and recurrent unions between dominant matrilines in Iron Age Britain

**DOI:** 10.64898/2026.08.03.742615

**Authors:** Iñigo Olalde, Ian Armit, Lindsey Büster, Malcolm Lillie, Estibalitz Urkixo F. de Zuazo, Harald Ringbauer, Ali Akbari, Laura Castells Navarro, Diego Esteve-Gómez, Helen Goodchild, Derek Hamilton, Núria Puig i Riera, Almudena Sánchez-Sanz, Jo Buckberry, Chelsea Budd, Anwen Caffell, Peter Halkon, Malin Holst, Philip Jerand, Eva Panagiotakopulu, Mark Stephens, Max Stubbings, Paula Ware, Madeleine Bleasdale, Tom Booth, Kim Callan, Ella Caughran, Claire-Elise Fischer, Trudi Frost, Lora Iliev, Aisling Kearns, Michael Legge, Matthew Mah, Mackenzie K. Masters, Nihal Manjila, Mariam Nawaz, Jonas Oppenheimer, Paola Ponce, Charlotte Primeau, Marina Silva, Pontus Skoglund, Pooja Swali, Lijun Qiu, Gregory Soos, J. Noah Workman, Fatma Zalzala, Nick Patterson, Swapan Mallick, Nadin Rohland, David Reich

## Abstract

Kinship practices underpin all traditional societies, forming the basis for socially sanctioned reproductive unions, residence patterns and the inheritance of rights and property^1,2^. Although the relationship between biological relatedness and kinship is not always straightforward, ancient DNA studies are increasingly used to examine the extent to which biological relatedness underpinned social constructs of kinship in prehistoric societies^3–6^. Here, we report the analysis of genome-wide data for 534 individuals from the Arras Culture of Middle Iron Age northeast England (including 390 from Wetwang Slack, 100 from Pocklington, and 29 from Melton), finding evidence for communities with kinship systems structured along matrilineal lines. At Wetwang Slack, we reconstruct a 13-generation pedigree comprising 195 individuals structured around female-line connections: matrilineal transmissions greatly outnumbered patrilineal ones and male reproductive partners were largely absent from the cemetery, plausibly because they were buried in their own natal communities. Furthermore, the three main sites with robust sample sizes were characterised by non-overlapping dominant mitochondrial haplogroups, implying a maternal clan-based structure. Reproductive unions at Wetwang Slack suggest a recurrent alliance between two dominant maternal descent groups, with members of each group never reproducing with members of their own maternal lineage. Meanwhile, three individuals from lavishly furnished ‘chariot burials’ at Wetwang Slack were close maternal relatives belonging to a lineage with consecutive generations of close kin unions, a pattern largely absent among other individuals at the site. These results indicate highly distinctive social practices among an elite group embedded in the wider kinship network of the Arras community.

## Introduction

Understanding kinship practices is essential for reconstructing how past societies organized social relations, residence and the transmission of rights and property. Kinship cannot, however, be reduced to biological relatedness^7^ and kinship practices commonly also allow for alternative, non-biological ways of creating kin^8^. Since kinship structures within any society are social constructs that may be more or less widely adhered to, they are open to challenge and subversion from disparate groups and individuals^9^. Even where kinship structures appear to be clearly formulated, human agency, along with social and economic contingency, ensures that their application in practice is ‘messy’. Genetic data are, however, of great value for investigating the extent to which biological relatedness influences kinship practices. With sufficiently large, multi-generational data, we can examine the extent to which biological relatedness is employed within kinship practice and explore the nature of biologically based kinship structures and degrees of adherence to them over time.

Genetic data have been increasingly used to explore issues of kinship practices in prehistoric societies through the analysis of biological relatedness^10–12^. For the Neolithic period, this work revealed multi-generational pedigrees for burial populations in Britain^4^, France^5^ and Anatolia^6^. Analysis of nine individuals in Iron Age Slovenia demonstrated that burial within specific barrows could be linked to biologically-related groups^13^, while work on high-status graves in Iron Age southern Germany has identified biological relationships between high-status individuals buried up to 100 km apart^14^. Recently, analysis of 57 genomes from Durotrigian cemeteries in southern Iron Age Britain identified an extended kin group structured around maternal residence patterns, with unrelated individuals being predominantly male^3^. This, together with reduced mitochondrial diversity in British Iron Age cemeteries, pointed to a widespread practice of matrilocality^3,15^. This raised the possibility that female-centred residence practices were not isolated phenomena, but broader features of Iron Age Britain, while also highlighting the need for higher-resolution analysis in order to obtain insight into the nature of kinship and descent practices driving these patterns, as well as exploring how such systems may have varied between communities.

The Middle Iron Age Arras Culture, centred in East Yorkshire (northeast England) from the fifth to first centuries BCE, includes some of the largest cemeteries from this period (containing hundreds of burials in some cases), and is unusual for the British Iron Age in practising an archaeologically visible burial rite comprising individual inhumation under small square earthen barrows^16–18^. Arras cemeteries are also well known for lavishly furnished ‘chariot burials’, long considered to represent the elite within Arras Culture society^18^ and characterised by deposition of the deceased on a complete or dismantled two-wheeled chariot or cart. These inhumations are generally the most elaborately equipped, frequently containing weaponry and mirrors, as well as a range of other objects including decorated horse-gear and chariot-fittings.

This cultural context thus offers an unprecedented opportunity for archaeogenetic studies to contribute to studies of social relations and kinship structures within the communities that occupied Britain in the centuries prior to Roman occupation.

## Data generation

We sampled bone and teeth from 495 individuals from seven cemeteries belonging primarily to the Middle Iron Age Arras Culture of East Yorkshire and neighbouring areas. After in-solution enrichment for more than a million single-nucleotide polymorphisms (SNPs), 490 individuals yielded genome-wide data of sufficient quality for analysis. Together with 44 published individuals^19^ from the same area and cultural context, the final dataset includes 534 individuals from 10 cemeteries (Supplementary Table 1): 390 from Wetwang Slack, 100 from Pocklington, 28 from Melton 1, seven from East Coast Pipeline, three from Burton Fleming, two from Nunburnholme, and one individual each from Melton 2, Wetwang Village, Ferry Fryston and Burstwick (Fig. 1a; Extended Data Fig. 1). The mean number of SNPs recovered from a core set of 1.15 million autosomal targets is 964,521, with 526 individuals having more than 600,000 SNPs covered.

**Fig. 1.**
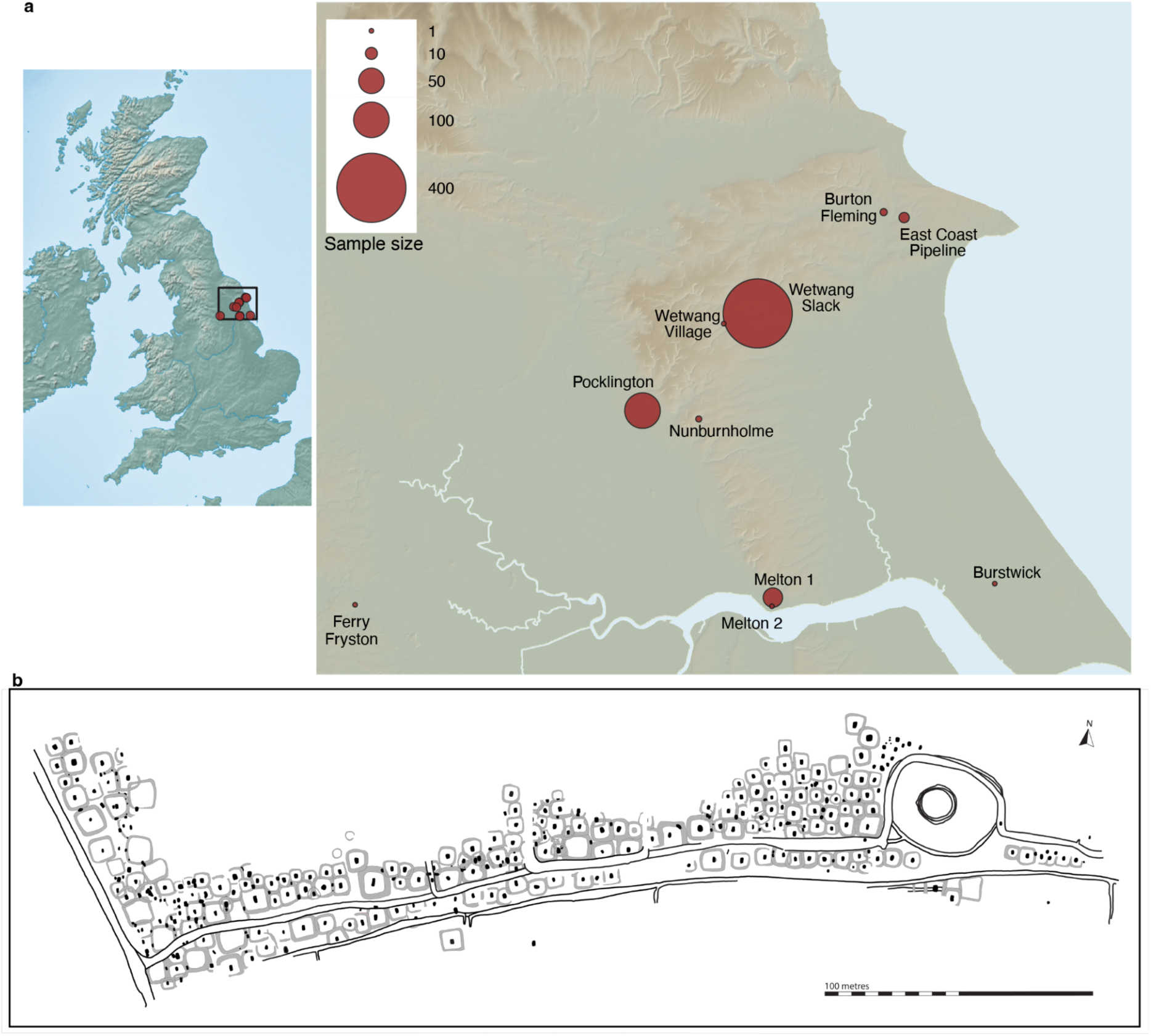
Archaeological and geographical context of the Arras Culture cemeteries. **a**, Location of the Arras Culture cemeteries analysed in this study. Circle area is proportional to the number of sequenced individuals from each site. **b,** Plan of the Wetwang Slack main cemetery area showing the distribution of burial features, including square-ditched barrows, flat graves and ditch burials. The chariot burials are located ∼200 metres to the west.

Wetwang Slack, located on the Yorkshire Wolds, is the largest excavated Iron Age inhumation cemetery in Britain and was investigated primarily during rescue work ahead of gravel quarrying in the 1970s and 1980s^20,21^. The cemetery comprised 446 burials arranged in a broadly linear fashion, following the course of a routeway through a densely settled landscape of Iron Age farms and fields^22^ (Fig. 1b), and included primary and secondary burials associated with square-ditched barrows, together with ditch burials and ‘flat’ graves. Close to the west of the main cemetery lay a separate group of five barrows, three of which each contained a single richly furnished chariot burial^21,23^. The Arras cemetery at Burnby Lane, Pocklington (hereafter referred to simply as ’Pocklington’), lies approximately 18 km southwest of Wetwang Slack (Fig. 1a) and represents the second largest excavated cemetery sample, with 100 individuals. It was separated into two groups, with a larger group to the west and a smaller group to the east, along with a number of outliers^24^. The Pocklington cemetery comprises inhumations in square and round barrows, in barrow ditches, and in flat graves between barrows^24^. The third main cemetery in our dataset, Melton 1, is located next to the Humber estuary, ∼30 km south of Wetwang Slack and Pocklington (Fig. 1a), and consists of nine Iron Age square barrows and six flat graves, alongside a number of secondary interments^25^. On the basis of modelled AMS radiocarbon dates, Wetwang Slack, Pocklington and Melton 1 were broadly contemporaneous during the fourth and second centuries cal. BCE^25–27^.

All sites were excavated in advance of either quarry works (Wetwang Slack) or housing development (Pocklington and Melton 1), and all are, to some extent, affected by plough truncation (particularly of those burials representing later insertions into upstanding barrow monuments). As such, particularly in the smaller remit of the excavations at Pocklington and Melton 1, it is likely that the full spatial extent of the cemeteries has not been explored. Nevertheless, the large number of individuals with high-quality genome-wide data, combined with the unusually well-preserved archaeological and spatial context of the excavated parts of these major cemeteries, provides a rare opportunity to examine the relationship between biological relatedness, burial location and social organisation at a population scale in Iron Age Britain.

## Lack of recent continental affinities

The cultural affiliation of the Arras Culture has long been debated because of its use of square barrows, chariot burials and La Tène-style objects, which have often invited comparison with Iron Age communities in continental Europe, particularly groups from the Paris Basin^28^. However, our genetic data do not support a model in which the individuals buried in Arras Culture cemeteries derived from a recent migration of individuals from the Continent. Genome-wide ancestry was highly homogeneous across Arras sites and burial contexts (Extended Data Fig. 3), and *qpAdm* modelling using Iron Age individuals from France as the sole ancestry source provided a poor fit (p-value < 0.001) (Supplementary Table 7). We also found no evidence of elevated IBD sharing between Arras Culture individuals and those from Iron Age France (Supplementary Table 5), and the most common Y-chromosome lineage in Arras Culture males was R1b-DF13, present in high frequencies across Bronze Age Britain (Extended Data Fig. 4)^19,29^. These results suggest that the distinctive Arras funerary tradition of East Yorkshire reflects local British communities participating in broader La Tène-connected cultural networks, rather than the establishment of a recently arrived continental population.

## Shift from patrilocality to matrilocality in Britain

We first examined whether Arras Culture cemeteries formed genetically isolated burial communities or whether they were part of a wider regional network. Identity-by-descent (IBD) network analysis revealed that genetic links were concentrated primarily within cemeteries: most individuals from the three most densely sampled sites (Wetwang Slack, Pocklington and Melton 1) formed clear site-specific clusters (Extended Data Fig. 2a). This indicates that biological relatedness was structured mainly at the cemetery level and reflected in ‘communities of the dead’. However, these clusters were not genetically isolated. We also detected multiple cross-site IBD links, particularly between Wetwang Slack, Wetwang Village, Pocklington, Burton Fleming and Nunburnholme, showing that site-specific burial communities existed within a wider network of biological connections across the Arras region (Extended Data Fig. 2b). This pattern was not mirrored by extensive sharing of mitochondrial haplotypes across Arras sites, suggesting that biological links between communities were not primarily maintained through female-mediated mobility, such as female exogamy.

Instead, the three largest Arras cemeteries were each characterised by a small number of high-frequency mitochondrial lineages, and these dominant lineages were largely non-overlapping between sites (Fig. 2a). At Wetwang Slack, two maternal haplogroups, T2e1a1b and H1ao, accounted for 51% of the cemetery population, while Pocklington and Melton 1 were each dominated by three major haplogroups accounting for 70% and 58%, respectively. At Wetwang Slack, this pattern was strongly associated with biological relatedness. Individuals with many relatives in the cemetery were predominantly associated with the two major maternal lineages, whereas individuals with few or no relatives at the site displayed substantially higher mitochondrial diversity (Fig. 2b). Thus, low mitochondrial diversity at Wetwang Slack reflects the presence of a large, biologically related burial community centred on a restricted set of maternal lineages, alongside a smaller and more diverse set of individuals with weaker genealogical ties to the rest of the cemetery population. This site-specific reduction in mitochondrial diversity contrasts with the Y-chromosome pattern: paternal lineages were more diverse, and the major Y-chromosome lineages occurred at broadly similar frequencies across the main cemeteries (Extended Data Fig. 4). The resulting asymmetry—low and site-specific mitochondrial diversity but high and regionally shared Y-chromosome diversity—is difficult to reconcile with a patrilocal model, and instead points to a practice of matrilocality and male exogamy in Arras communities.

**Fig. 2.**
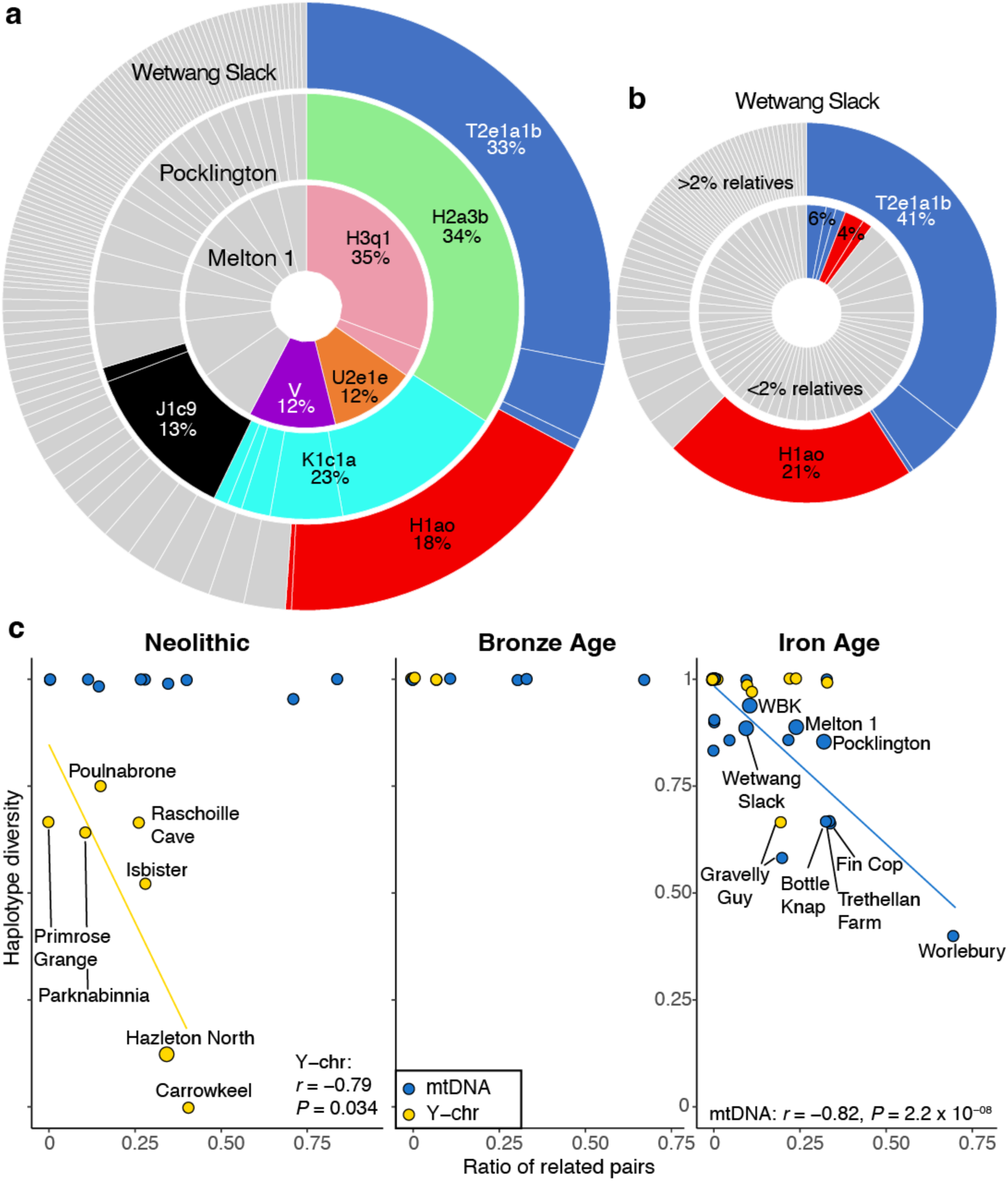
Dominant maternal lineages structure Arras Culture burial communities. **a**, Mitochondrial haplotype frequencies at Wetwang Slack, Pocklington and Melton 1. Haplotypes belonging to dominant haplogroups are coloured, while less frequent lineages are shown in grey. **b,** Mitochondrial haplotype composition at Wetwang Slack separated by within-site relatedness (individuals for whom <2% or >2% of other individuals at the site were identified as relatives). **c,** Mitochondrial and Y-chromosome within-site diversity estimates in Britain and Ireland as a function of the ratio of related pairs. Sites with <3 individuals available for diversity estimation were excluded. Sites with more than 20 individuals are displayed with larger dots. Haplotype frequencies and diversities were calculated retaining a single individual per cluster of first-degree relatives to avoid overrepresentation of large sibships.

When placed in a broader chronological framework, the Arras cemeteries form part of a wider pattern emerging for Iron Age Britain^3,15^. Developing the approach of Cassidy et al.^3^, we compared mitochondrial and Y-chromosome haplotype diversity across British and Irish prehistoric sites with sufficient genomic data. Iron Age cemeteries show a strong association between high within-site relatedness and reduced mitochondrial diversity, while Y-chromosome diversity remains comparatively high (Fig. 2c). This contrasts with the pattern observed for Neolithic sites, where mitochondrial diversity remained high and Y-chromosome diversity was reduced, especially at sites with a high number of relatives such as Hazleton North^4^. The Arras data therefore strengthen the view that Iron Age Britain saw a marked reorganisation of residence practices, with burial communities increasingly structured around matrilocality.

## Biological relatedness strongly structured cemetery space at Arras cemeteries

If we assume that cemeteries are organised according to kinship practices, then, leveraging our large sample size, we can explore the relationship between biological relatedness and kinship by testing whether biological relatedness influenced the spatial organisation of Arras cemeteries (Supplementary Table 8). At both Wetwang Slack and Pocklington, close biological relatives were buried significantly closer to one another than other pairs of individuals (p-value < 0.001) (Extended Data Fig. 5a). At Wetwang Slack, pairs related to the third degree or closer had a median burial distance of 36.9 m, compared with 127.0 m for other pairs. At Pocklington, the equivalent values were 28.3 m and 58.0 m, reflecting the overall smaller size of the excavated cemetery. This spatial pattern was not limited to close biological relatives. At Wetwang Slack, individuals sharing the same mitochondrial haplotype were also buried closer to one another than individuals with different mitochondrial haplotypes (median burial distance of 113.1 m versus 128.1 m), and this association persisted even after excluding pairs related to the third degree or closer (Extended Data Figure 5b). These differences indicate that burial location was not random with respect to biological relatedness. Rather, the people responsible for burial placement appear to have had detailed knowledge of genealogical relationships within the community. At Wetwang Slack, this was not restricted to close-family relationships, but broader maternal-line affiliations.

Burial type was also associated with relatedness. At Wetwang Slack, individuals interred as primary burials in barrows had a higher proportion of relatives within the cemetery than individuals in flat graves and ditch burials (p-value < 0.02; Figure S51 and Table S1). A similar trend was observed at Pocklington, where individuals in barrows had more within-site relatives than those in other grave types, although the difference was not statistically significant. Since primary barrow burials are present across multiple generations in the reconstructed pedigrees at Wetwang Slack (Extended Data Fig. 6c), these patterns are not the product of chronology, i.e. diminishing space for large monuments in later phases of a site attracting a more heterogeneous burial community. Rather, they suggest that primary barrows were preferentially used by particular family groups or lineages, whereas flat graves and ditch burials were used by a more heterogeneous subset of the contemporary community.

The spatial organisation of Arras cemeteries, therefore, reflects more than proximity between recently deceased family members. At Wetwang Slack especially, burial location, burial type and mitochondrial affiliation together point to a cemetery organised around long-lived, biologically related groups, with maternal lines providing a major persistent axis of kinship structure. The scale of genetic sampling at Wetwang Slack allows these aggregate patterns to be examined within a multi-generational pedigree, revealing how descent-based affiliation governed burial inclusion across generations.

## A reconstructed family of 195 members across 13 generations

Although Pocklington and Melton 1 individuals had high proportions of within-site relatives (here and throughout the manuscript, defined as such if sharing more than two IBD segments >8 cM and a total of >24 cM in IBD), with median proportions of 33% and 26%, respectively, most individuals at these sites had, at most, one first- or second-degree relative (Figure S48). As a result, the few reconstructed family pedigrees at Pocklington and Melton 1 could not be extended beyond three generations. In contrast, at Wetwang Slack, 162 individuals had multiple first- or second-degree relatives (Figure S48; Supplementary Table 4), allowing us to reconstruct, to our knowledge, the largest multi-generational pedigree yet established for a prehistoric population.

This pedigree comprises 195 individuals connected by chains of third-degree-or-closer relationships across 13 generations, representing half of all individuals with genome-wide data from Wetwang Slack (Fig. 3a). A further 93 individuals could be linked to this pedigree through fourth- or fifth-degree relationships, while the remaining 102 individuals from this site were only distantly related, or unrelated, to members of the main pedigree. Unlike individuals lying outside this pedigree (51.3% female; binomial test, p = 0.77), those included within the 195-person pedigree were biased towards females (61% female; binomial test, p = 0.0025) and were not evenly distributed through time (Extended Data Fig. 6). The first three generations contain only 14 individuals, predominantly adult females (n = 12), and all are represented by primary barrow burials, whereas the final three generations contain only ten individuals, including four infants and five primary barrow burials. By contrast, more than three quarters of the pedigree falls within generations 4–8, suggesting that these generations correspond to the floruit of burial activity in the cemetery.

**Fig. 3.**
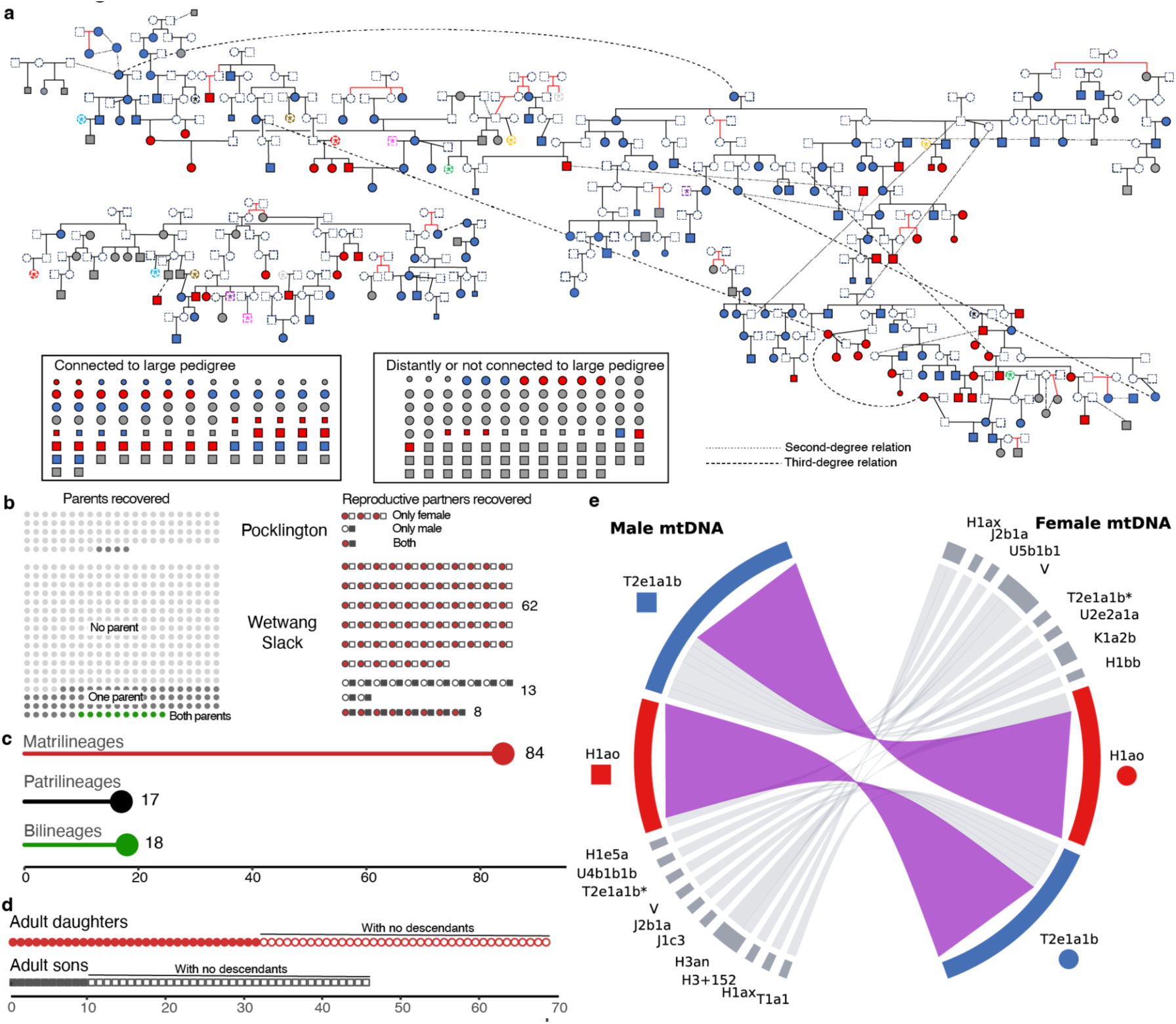
The main 13-generation pedigree reveals patterns of social organization at Wetwang Slack. **a**, Schematic representation of the largest reconstructed Wetwang Slack pedigree, comprising 195 individuals across 13 generations. Individuals are represented by symbols (squares for males and circles for females) coloured by mitochondrial lineage (red for H1ao and blue for T2e1a1b), with smaller symbols for non-adult individuals. Unsampled individuals appearing at more than one location of the tree are shown with a coloured asterisk. Second- and third-degree relations not fitted onto the tree are represented by dotted and dashed lines, respectively. Below, a group of individuals with fourth- or fifth-degree relationships with members of the pedigree is shown, together with a group with more distant or no relationships with members of the pedigree. **b,** Presence of parents and reproductive partners in the Wetwang Slack and Pocklington burial communities. Reproductive unions are classified according to whether both partners are buried at the cemetery, only the female partner is present, or only the male partner is present. **c,** Classification of reproductive unions according to the transmission of maternal and paternal lines. **d,** Retention of adult daughters and adult sons in the Wetwang Slack burial community, indicating how many of these individuals also had descendants buried at the cemetery. **e,** Mitochondrial lineages of reproductive unions at Wetwang Slack.

## Matrilineal descent and the social prominence of women

The reconstructed pedigree also reveals that genealogical continuity at Wetwang Slack was transmitted predominantly through women. We classified reproductive unions according to whether the female partner (matrilineal), the male partner (patrilineal), or both partners (bilateral) had first- or second-degree relatives in earlier generations of the cemetery. Matrilineal unions greatly outnumbered patrilineal and bilineal unions (84 versus 35; p = 8.2 × 10⁻⁶) (Fig. 3c), and this pattern was observed across the pedigree rather than being restricted to a single branch (Supplementary Table 11). This provides direct genealogical evidence that inclusion in the burial population at Wetwang Slack was determined primarily through female-line connections.

Several women appear to have acted as long-term genealogical and spatial anchors within the cemetery (Figure S57). Female I31517 (see Supplementary Table 1 for correspondence between genetic and archaeological identifiers)—the individual with the highest number of relatives (n=133) at Wetwang Slack—was the offspring of closely related parents (most likely third-degree relatives; Figure 4c and Supplementary Table 1) and was buried as a primary inhumation in a barrow in the central part of the cemetery. We identify 37 descendants of I31517 across nine generations (Extended Data Fig. 7a), many of whom were buried close to her (Extended Data Fig. 7b). These include her maternal-line great-great-great-grandson I30903, who was buried in the ditch of her own barrow, despite a time separation of likely more than 100 years. Female I36803 shows a similar, although smaller-scale, pattern: she had 11 identified descendants across seven generations, all of whom were also buried in close proximity (Extended Data Fig. 7). These cases suggest that particular women remained important reference points for burial placement long after death, implying a remarkable degree of genealogical memory within the community, together with a memory, or direct marking of, the burial location of prominent individuals.

**Fig. 4.**
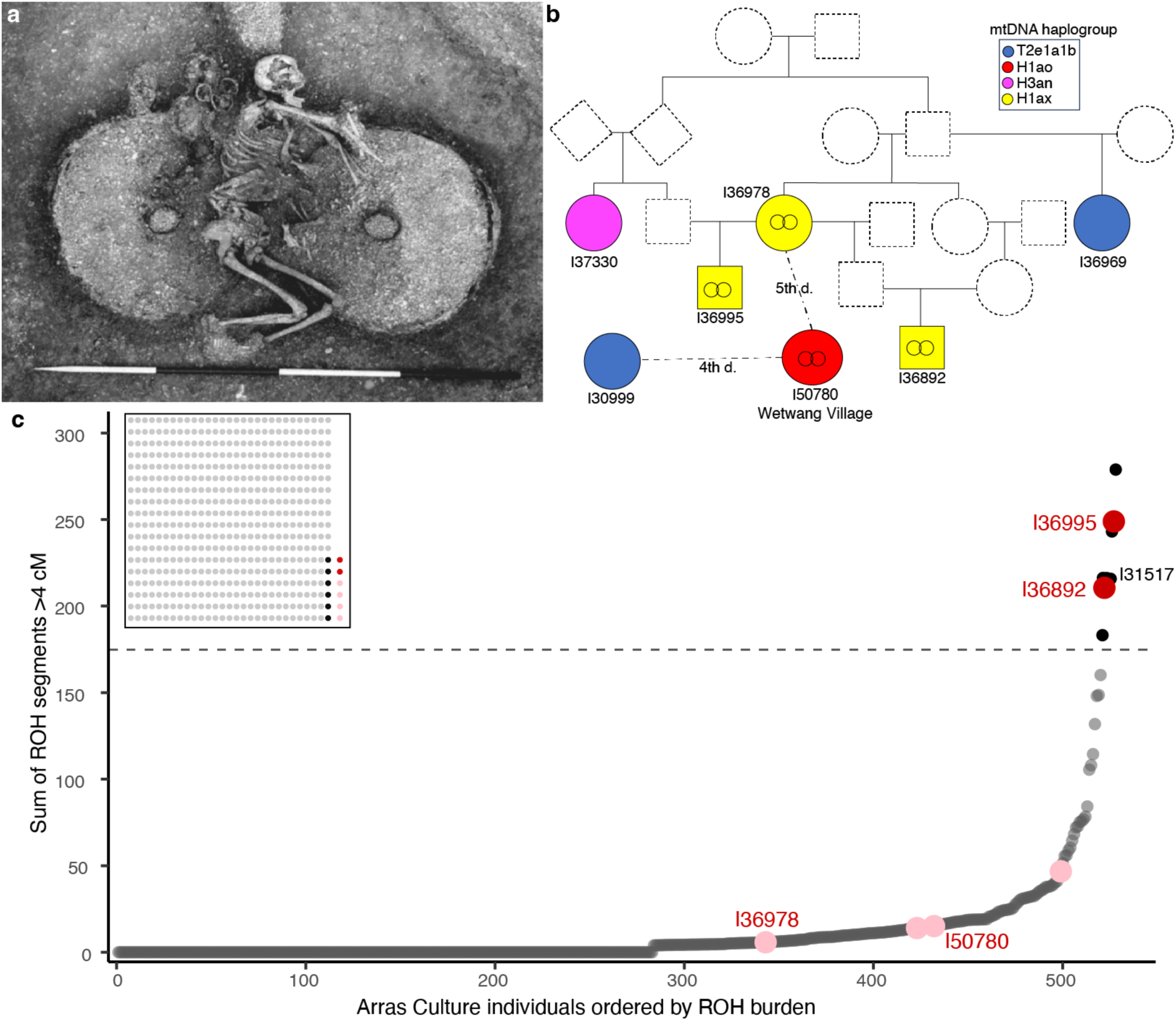
Chariot burials at Wetwang Slack belonged to an endogamous maternal lineage. **a**, Photograph of the chariot burial of female I36978 at Wetwang Slack. **b,** Pedigree showing the genealogical relationships among individuals buried with chariots at Wetwang Slack and related individuals in the wider Wetwang Slack cemetery and at Wetwang Village. Chariot burials are indicated with two circles inside the symbol. **c,** Total length of long (>4 cM) runs of homozygosity among Arras Culture individuals. Each point represents one individual, ordered by the summed length of ROH segments longer than 4 cM. Chariot burials are shown in pink/red. The dashed line marks 175 cM, used here as an approximate threshold for parental relatedness at the level of first cousins or closer. The inset shows the proportion of chariot (red) and non-chariot (black) individuals above this threshold.

## Matrilocality, male exogamy and multiple reproductive partners

Although biological relatedness clearly shaped practices of inclusion within burial communities, the Arras cemeteries did not represent burial grounds for complete residential households in which reproductive couples and their children were routinely buried together (Supplementary Table 9). While individuals below reproductive age at Wetwang Slack do not show a significant genetic sex bias (37.3% female; binomial test, p = 0.067), genetic females were clearly overrepresented among individuals above reproductive age at Wetwang Slack (59.5% female; binomial test, p = 0.00063) and Pocklington, but in this latter case without reaching statistical significance (57.6% female; binomial test, p = 0.17). At Wetwang Slack, adult female bias is particularly pronounced within the dominant T2e1a1b maternal lineage (68.0% female; exact binomial test, p = 7.0 × 10⁻^5^), while the second major lineage, H1ao, shows no evidence of significant sex bias (52.3% female; p = 0.80). At Wetwang Slack and Pocklington, 80% and 96% of individuals had neither parent identified within the cemetery, although, at Wetwang Slack, 17% of these individuals had other ancestors present. Reproductive partners were, however, rarely both present in the same burial community (Fig. 3b; Supplementary Table 10). When only one member of a reproductive pair was buried, this was more often (62 out of 75 cases at Wetwang Slack and three out of four cases at Pocklington) the female partner (Fig. 3b), suggesting that men who reproduced with women buried in the cemetery were frequently buried elsewhere, plausibly in their own natal or descent communities.

The large pedigree also provides direct evidence for matrilocality and male exogamy. Adult daughters from families already established at Wetwang Slack were more likely than adult sons to be buried in the cemetery (69 adult daughters versus 46 adult sons; p = 0.039), and this bias was strongest among individuals who themselves had descendants represented in the pedigree (32 versus 10; p = 9.4 × 10⁻⁴) (Fig. 3d; Supplementary Table 12). This pattern is consistent with a matrilocal tendency, in which daughters remained affiliated with the local burial community while sons generally reproduced elsewhere. Indeed, most of the identified adult sons, presumably having been returned to their natal community for burial, do not themselves have offspring at Wetwang Slack; these offspring may instead have been buried in their mother’s communities elsewhere. The ten adult sons who did have descendants buried at Wetwang Slack can also be accommodated within the same framework: all reproduced with women from families already represented in the cemetery, so their offspring would also be affiliated with Wetwang Slack through their maternal line. Similarly, the few sampled reproductive partners of people belonging to lineages already represented in previous generations at Wetwang Slack tended to have additional relatives (beyond their immediate descendants) buried at the cemetery (Supplementary Table 14; Extended Data Fig. 9). While most unions were likely exogamous, as suggested by the large number of missing reproductive partners, these results show that reproductive partners were sometimes embedded within pre-existing local networks.

Male exogamy is also supported by direct cross-site genealogical links among the three main sampled Arras cemeteries. The closest relationships connecting Wetwang Slack, Pocklington and Melton 1 are consistent with recent male-mediated movement between burial communities (Extended Data Fig. 8). The clearest case links adult male I30994 from the large pedigree at Wetwang Slack with adult female I21975 from the largest three-generation pedigree at Pocklington, who were themselves third-degree relatives (Extended Data Fig. 8a). Their relationship is best explained through their fathers, who were first-degree relatives. I30994’s maternal relatives, descendants and reproductive partner’s close relatives were buried at Wetwang Slack, while I21975’s close maternal relatives were buried at Pocklington. This configuration suggests that a male from Pocklington reproduced at Wetwang Slack and that his descendants were then affiliated and thus interred with the Wetwang Slack burial community. Comparable cross-site links between Wetwang Slack and Melton 1 (Extended Data Fig. 8b), and between Pocklington and Melton 1 (Extended Data Fig. 8c), also point to recent male-driven connections between burial populations at those sites.

Finally, the pedigree structure at Wetwang Slack and Pocklington provides evidence for men (n = 7) and women (n = 22) having multiple partners, with those few recovered individuals (n = 12) with more than one inferred reproductive partner all being genetic females (Supplementary Table 13). This pattern could partly reflect serial partnerships, especially in a community where adult mortality may have disrupted earlier unions. However, the absence of sampled males with multiple partners, together with the broader female-centred structure of the pedigree, raises the possibility that some multiple partnerships (i.e. exogamous male reproductive partners shared by a single female) represented polyandrous reproductive arrangements within a strongly female-line-centred social structure. At Wetwang Slack, for example, the unsampled mother of I30998 and I36874 had offspring with I37339 and with an unsampled male who was most likely a paternal second-degree relative of I37339 (Extended Data Fig. 7). This configuration is compatible with the type of arrangement described by Caesar, who claimed that among Britons, groups of men—particularly brothers, and fathers with their sons—shared wives (*De Bello Gallico*). We also identified the inverse configuration, in which an unsampled male had offspring with both I36884 and her daughter I36814 (Figure 3). These were the only two clear instances across the reconstructed pedigrees in which an individual’s multiple reproductive partners were themselves closely related, suggesting that such arrangements were not widespread within the sampled Arras burial communities.

## Recurrent unions between two matrilines

The reconstructed pedigree further suggests that reproductive unions at Wetwang Slack were structured by maternal-line affiliation. Among 45 reproductive unions for which both maternal lineages could be assigned, 25 involved one partner carrying the major T2e1a1b haplotype and the other H1ao (Fig. 3e). Thus, more than half of the reproductive unions at Wetwang Slack joined the two dominant matrilines at the site. These T2e1a1b–H1ao unions were not restricted to a single branch of the large pedigree but occurred across several families within the tree (Supplementary Table 15). The direction of the pairing was approximately balanced, with 12 unions involving an H1ao male and 13 involving a T2e1a1b male, indicating that the pattern was not driven by one maternal lineage consistently providing male or female reproductive partners. To test whether the recurrent pairing between the two dominant maternal lineages could be explained simply by their high frequencies in the cemetery, we compared the observed number of T2e1a1b–H1ao unions with a random expectation based on the frequency of these haplogroups among adult males and females at the site. The expected probability of any T2e1a1b–H1ao reproductive union is 13.4%, whereas the observed frequency is 55.6% (25/45). T2e1a1b–H1ao unions therefore occurred 4.1 times more often than expected under random pairing with respect to maternal lineage (one-sided binomial test: p = 3.2 × 10⁻^11^). The two maternal lineages do not, however, appear to have played equivalent roles within the cemetery. T2e1a1b matrilineages show a much deeper and more persistent continuity across the reconstructed pedigree, continuing for up to ten generations in one branch, nine generations in another and seven generations in a third. In contrast, H1ao matrilineages rarely persist for more than one or two consecutive generations within the cemetery. Moreover, H1ao individuals reproduced almost exclusively (89%) with T2e1a1b individuals, whereas T2e1a1b individuals had a more diverse set of partners (68% with H1ao).

A particularly illustrative example is the family of I31005, where six consecutive generations of reproductive unions involved partners from the T2e1a1b and H1ao maternal lineages (Extended Data Fig. 10a). In one fourth-generation union within this family, between I37335 and I37129, the two sons were buried closer to their mother and maternal aunt and uncle than to their father and paternal relatives (Extended Data Fig. 10b). This provides a clear spatial example of the broader genealogical pattern: reproductive unions connected the two dominant maternal lineages, but burial affiliation tended to follow the maternal line.

## An endogamous elite lineage within the chariot burials

At Wetwang Slack, a small, spatially separate group of conjoined barrows, c. 200 m west of the main cemetery area, included two male and one female chariot burials sharing mitochondrial haplogroup H1ax. Female chariot burial I36978 (Fig. 4a) was the mother of male chariot burial I36995 and most likely the paternal grandmother and maternal great-aunt of the second male chariot burial I36892, who descended from her union with a second male (Figure 4b). Both male chariot burials had parents who were closely related on the order of first cousins (Figure 4c), indicating repeated close-kin unions within this elite lineage.

The chariot burials are genetically and spatially distinct, but not totally isolated from the wider Wetwang burial community, having two detectable close relatives (Figure 4b) at the site. Female I37330 was most likely a paternal aunt of I36995 and first cousin of I36995’s mother (I36978), and was buried near them. She was interred with an elaborate iron brooch with coral inlay and copper alloy studs, suggesting a relatively wealthy status. The second close relative, female I36969, was buried in the main cemetery area, belonged to the major T2e1a1b maternal lineage and was most likely the paternal half-sister of I36978.

More distant genetic links strengthen the suggestion that those individuals afforded chariot burials represent a close-knit elite group, in that I36978 shares 7 IBD segments (and a total length of 128 cM) with female I50780 in the richly furnished chariot burial from Wetwang Village, c. 1 km to the west of Wetwang Slack. I50780 belonged to the major H1ao Wetwang Slack matrilineage and also had several relatives in the main Wetwang Slack cemetery (Figure 4b), including fourth-degree relative I30999, who was a descendant of I31517’s T2e1a1b matriline. Thus, while the chariot burials at Wetwang Slack belonged to a distinct endogamous maternal lineage, they were embedded within broader kinship networks at Wetwang Slack and its environs, and were connected to both of the major maternal lineages at the site.

The level of endogamy observed in the chariot burials at Wetwang Slack was exceptional in comparison not only with the non-chariot burials at Arras Culture cemeteries, in which only 1% (6/522) of individuals showed comparable levels of parental relatedness (Fig. 4c), but also with the three chariot burials at Wetwang Village, Ferry Fryston and Melton 1 which were not highly endogamous. Close-kin unions therefore appear to have been rare in the wider burial community, particularly with respect to matrilineal relatives. With the exception of the parents of chariot burial I36892 (Fig. 4b), we find no evidence for a reproductive union between two individuals carrying the same mitochondrial lineage. Since the two major haplogroups are present in 33% and 18% of individuals at the site, this avoidance demonstrates detailed knowledge of maternal-line affiliation across many generations and possibly hundreds of years.

## Discussion

Our genetic analyses of 534 (including 494 newly sequenced) individuals from the Arras Culture cemeteries of East Yorkshire indicate repeated and persistent patterns of female-centred social organisation in which biological relatedness played a strong role in kinship structure over multiple generations spanning several centuries, within and between communities. This is evident at the most densely sampled cemeteries of Wetwang Slack and Pocklington, where spatial organisation indicates that genealogical relationships and biological relatedness were influential in both burial placement and burial type. At Wetwang Slack, a 13-generation pedigree of 195 individuals allows us to examine the nature of these (apparently predominantly genealogically and biologically driven) kinship practices on a scale—both in terms of size and generational span—not previously possible for Iron Age Britain. Here, we observe that certain women retained social significance across multiple generations and may have functioned as enduring spatial anchors around which groups organised mortuary practices within the cemetery. This suggests that female-centred forms of social organisation characterised by the same broad signature—dominant local maternal lineages, reduced mitochondrial diversity in kin-structured cemeteries, and comparatively diverse paternal lineages—were widespread across Iron Age Britain, while also allowing for substantial regional and local variation in how such systems were organised.

The Arras burial communities, particularly Wetwang Slack and Pocklington, should not, however, be treated as a simple proxy for the full residential population. The paucity of male reproductive partners indicates that burial was primarily structured by rules of affiliation and belonging to specific maternal lineages (i.e. matriliny), not simply by residence (i.e. matrilocality) as has been demonstrated in southern England^3^. At Wetwang Slack, a lack of male reproductive partners buried within the cemetery partially explains the observed sex bias and, based on ethnographic parallels^30^, is a strong indicator of matrilineal kinship practices, where individuals are generally buried with their maternal communities. However, if this were a recurrent pattern in Arras communities, we would expect the reverse pattern where adult sons of Wetwang Slack families who reproduced at other communities were returned for burial at Wetwang Slack. Indeed, we find 46 such cases and, as would be expected under this scenario, most of these individuals themselves lack descendants buried at Wetwang Slack, apart from those who had reproduced with females who also descended from Wetwang Slack maternal lineages. However, the significant depletion of adult sons versus adult daughters further contributes to the sex bias observed within the large family pedigree at Wetwang Slack, suggesting that a substantial number of sons never returned to their maternal communities or were buried elsewhere.

Repeated reproductive unions between the two dominant maternal lineages, T2e1a1b and H1ao, are compatible with preferential alliance between maternal descent groups. The two matrilines likely fulfilled different roles, with T2e1a1b forming the principal social axis around which the Wetwang Slack burial community was structured, while H1ao individuals were repeatedly incorporated through reproductive unions. This pattern resembles exogamous matrilineal moiety systems documented ethnographically across diverse societies^1^, including the Tlingit and Haida of the Northwest Coast of North America^31^, where descent-group membership is inherited through the mother and marriage conventionally joins members of opposite moieties. Although the asymmetrical persistence and representation of the two lineages at Wetwang Slack preclude identifying a formal dual organisation, the observed pattern reflects one of its key features: recurrent reproductive alliance between complementary maternal descent groups.

The strong signal of matrilineal descent accords with the prominence of rich female burials in the Arras Culture more widely^17^. Classical textual sources record prominent female protagonists in Britain during the first century CE, notably queens Boudica of the Iceni in eastern England, and Cartimandua of the Brigantes in northern England^32^, whose regal power suggests a broader presence of matrilineal kinship structures across Iron Age Britain. Recent genetic analysis has similarly noted matrilineal kinship between elite families in southern Germany, dating to the Hallstatt D period (c.600–450 BCE)^14^, while Classical textual sources have been used to suggest female political power in a range of other European Iron Age contexts^33^. The importance of the relationship between males and their maternal uncles is a recurrent feature of matrilineal societies^1^, and the prominence of this relationship in Classical textual accounts of Iron Age Europeans^34,35^ may be a further indicator of widespread matrilineal social organisation.

The importance of matrilineal descent was again evident in the analysis of elite chariot burials at Wetwang Slack, which belonged to a distinct maternal lineage. In contrast to the wider cemetery, where reproductive unions generally avoided close relatives (and even distant relatives belonging to the same matriline), the individuals afforded chariot burials at Wetwang Slack stood apart by showing recurrent close kin unions, consistent with a reproductive strategy in which high status, identity and claims to particular forms of burial were concentrated within a restricted descent group. Palaeogenetic and documentary evidence for consanguineous unions is generally rare^36,37^ but they are a recurrent feature of high-status groups^38–40^, where reproductive alliances may be used to maintain rank, inheritance, ritual status or group boundaries^41^. The possible presence of polygamy (as opposed to serial monogamy) among the Wetwang Slack chariot burials is notable in this regard, since it has potential to complicate claims of descent and inheritance. These challenges may, however, have been less acute in a matrilineal society preferentially permitting polyandry over polygyny (i.e. females rather than males having multiple reproductive partners), in which kin affiliation proceeds only through a single female line.

The degree of variation in the application of matrilineal principles, both within and between the Arras Culture cemeteries, highlights the messiness inherent in the construction of identity and kinship across time and space. Adherence to shared conventions of behaviour and practice may frequently be overridden by human agency (including deliberate subversion), contingency, and the practical complexities of implementing social ideals consistently. The 13 generations represented at Wetwang Slack, together with the complementary results from our two other key sites of Pocklington and Melton 1, provide an unprecedented opportunity to observe this variation through time, and to examine how matrilineal descent was socially recognized and translated into funerary practice. Despite this, the scale and longevity of burial practices represented here also reveal the persistence of major underlying structuring principles: across multiple generations spanning several centuries, burial membership, spatial organisation and reproductive alliances were repeatedly shaped by maternal-line affiliation. This demonstrates that female-line descent was not an incidental feature of Arras Culture communities, but a central organising principle, as it may well have been across the communities of Iron Age Britain more broadly.

## Data availability

Genotype data for individuals included in this study can be obtained from the Harvard Dataverse repository (https://reich.hms.harvard.edu/datasets). The DNA sequences reported in this paper have been deposited in the European Nucleotide Archive under accession number PRJEB122877.

## Acknowledgements

This research received funding from the European Research Council (ERC) under the European Union’s Horizon 2020 research and innovation programme (grant agreement no. 834087; the COMMIOS Project to IA). The excavations at Pocklington, Melton 1 and Burton Fleming were undertaken by MAP Archaeological Practice Ltd with osteological analysis undertaken as an element of the post-excavation funded programme of works. The ancient DNA analysis at Harvard Medical School was funded by NIH grant HG012287; John Templeton Foundation grant 61220; a gift from Jean-Francois Clin; the Allen Discovery Center program, a Paul G. Allen Frontiers Group advised program of the Allen Family Philanthropies; and the Howard Hughes Medical Institute. The excavations at Nunburnholme were undertaken by MCL and PH with funding from the Ferens Education Trust and the Wetland Archaeological and Environmental Research Centre at the University of Hull. The excavations at Melton 2 (PH) were funded by the RAI, East Riding Archaeological Association and the University of Hull. IO was supported by the Basque Government under “Grupos Consolidados” grant no. IT1971-26 and grant RYC2019-027909-I and project PID2022-140886NA-I00 funded by MCIN/AEI/10.13039/501100011033, “ESF Investing in your future” and FEDER, UE. We thank Elizabeth Curtis and Kristin Stewardson for laboratory work. Access to the skeletal remains from Wetwang Village was by permission of The British Museum and was facilitated by museum curatorial and collections staff, with special thanks to Dr Sophia Adams and Dr Rebecca Whiting. Access to the skeletal remains from Ferry Fryston was by permission of Wakefield Museums and Castles and was facilitated by David Evans. The authors are grateful to the late Dr John Dent, excavator of the Wetwang Slack cemetery, for his help and encouragement throughout the project.

---

**Extended Data Fig. 1.**
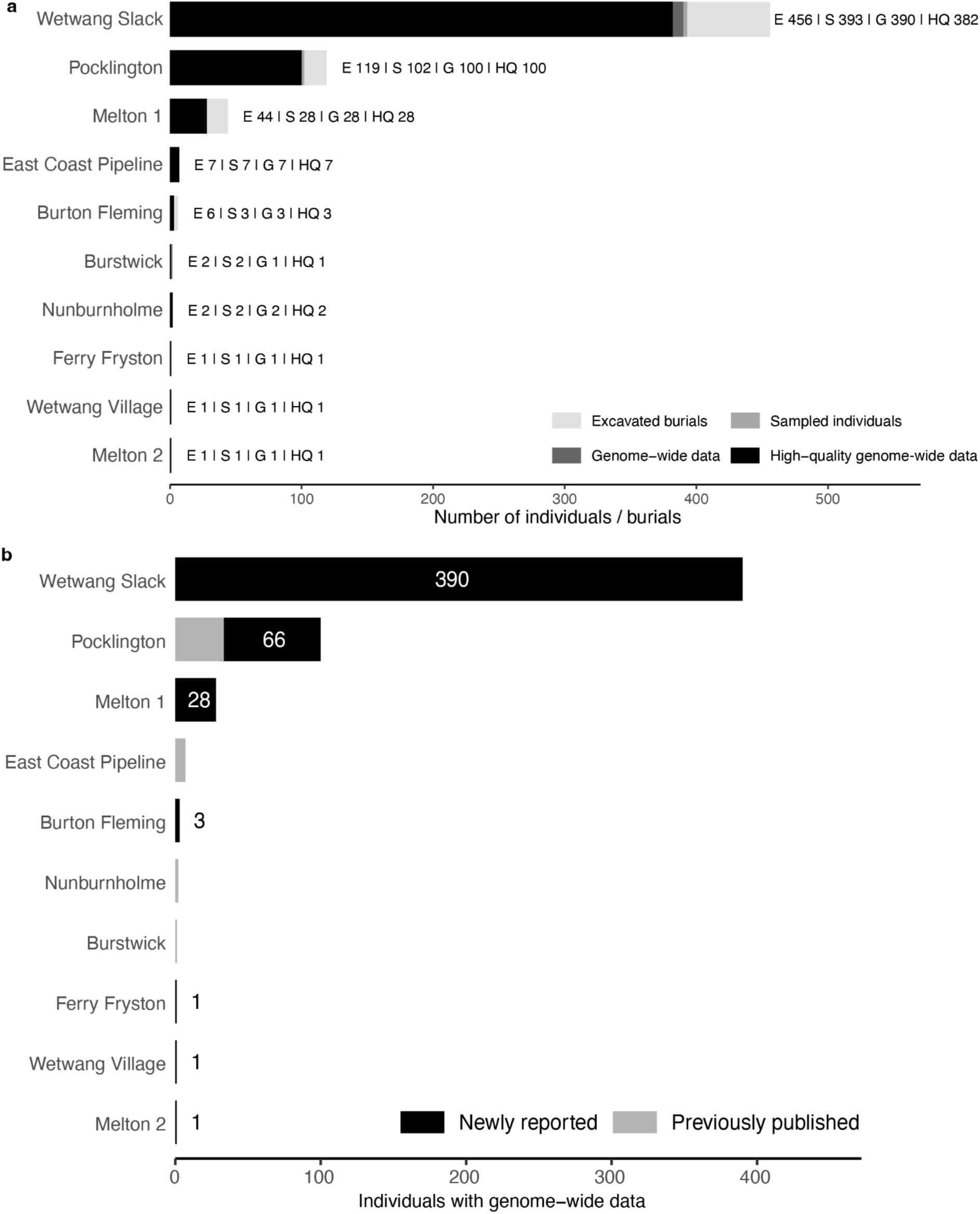
Sampling overview of Arras Culture cemeteries included in this study. **a**, Number of excavated burials, sampled individuals, individuals with genome-wide data and individuals with high-quality data amenable for IBD calling at each cemetery. **b,** Composition of the final genome-wide dataset, indicating newly reported and previously published individuals per site. Numbers inside or next to bars indicate the newly reported individuals.

**Extended Data Fig. 2.**
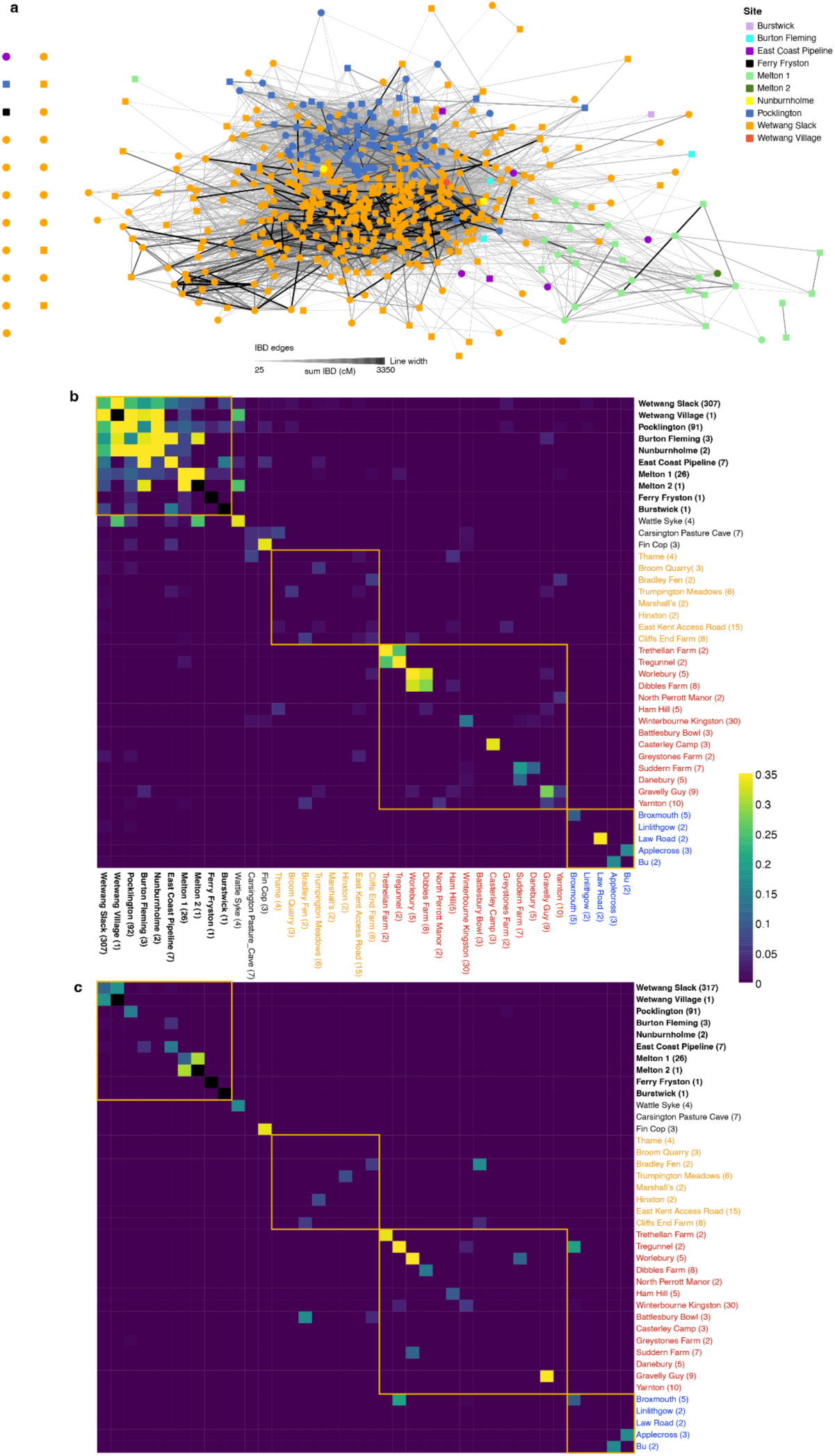
Kinship networks across Arras Culture cemeteries. **a**, IBD network of Iron Age Arras Culture sites included in the current study. Edges connect individuals sharing more than two IBD segments >8 cM and a total of >24 cM in IBD, and are weighted by the total IBD shared (sum IBD). Node colour denotes archaeological site and shape indicates genetic sex. Node positions are determined using a force-directed layout (Fruchterman–Reingold). Isolated individuals without connection meeting the threshold are shown separately. **b–c,** Pairwise mitochondrial and IBD matching across Middle and Late Iron Age sites from Britain. **b,** Ratio of pairs sharing IBD segments >12 cM and **c,** ratio of pairs sharing the same mitochondrial haplotype. Only one individual per cluster of first-degree relatives was retained. In bold, Arras cemeteries. Sites from southeast England, southwest England and Scotland are shown in orange, red and blue, respectively. The number in parentheses represents the sample size for each site.

**Extended Data Fig. 3.**
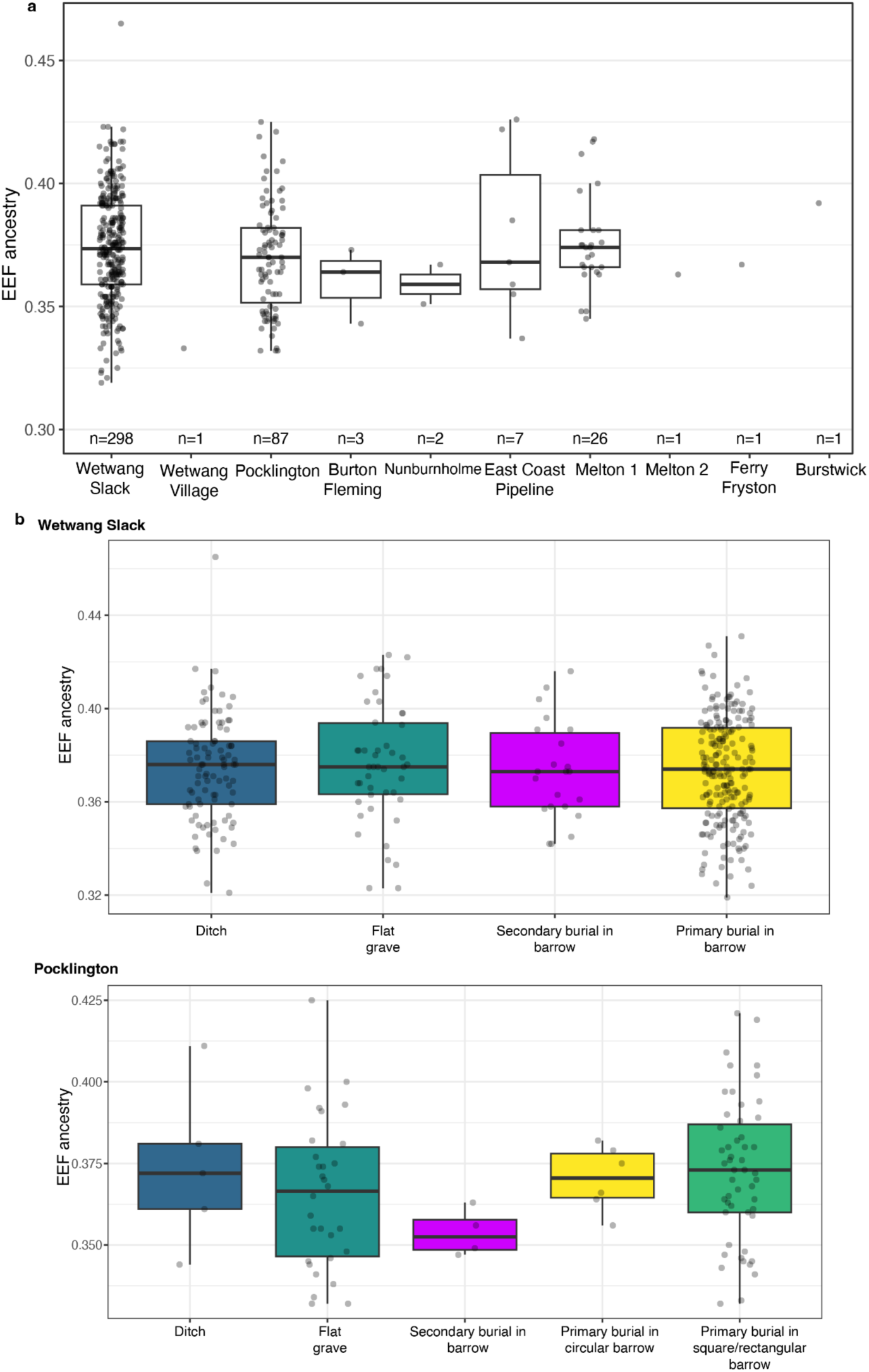
Genetic ancestry homogeneity across sites and burial contexts. Early European Farmer (EEF) ancestry proportions for Arras Culture individuals, as obtained with *qpAdm*. **a,** Across sites. **b,** Across burial types. We modelled the ancestry of each individual as a mixture of Western Hunter-Gatherer (WHG), Early European Farmers (EEF) and Steppe Early Bronze Age sources.

**Extended Data Fig. 4.**
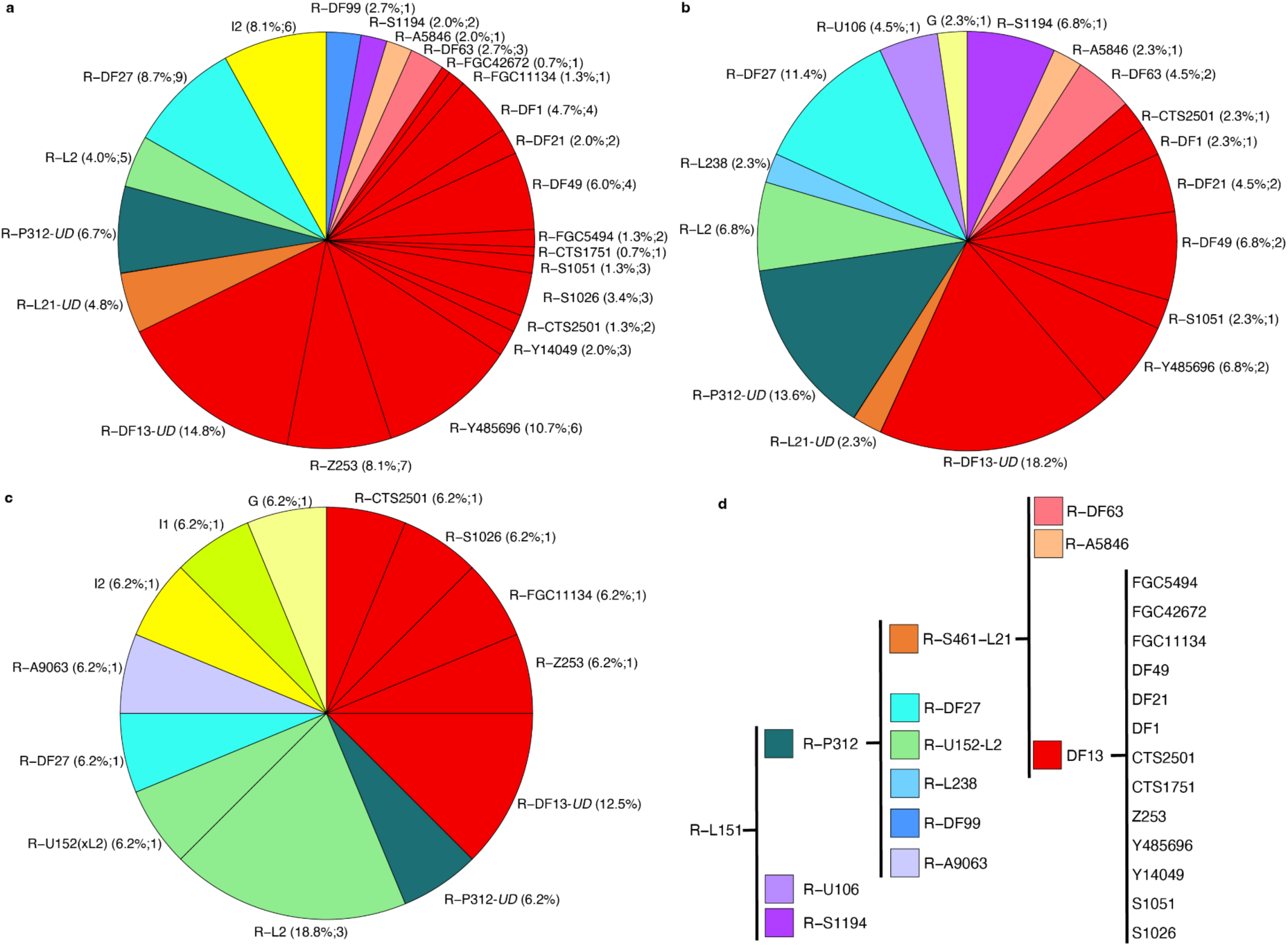
Y-chromosome patterns in Arras cemeteries. Y-haplogroup distribution at **a,** Wetwang Slack (n=149), **b,** Pocklington (n=44) and **c,** Melton 1 (n=16). For each haplogroup, we indicate in parentheses its frequency and the minimum number of distinct sublineages present in our dataset. *UD* (Undetermined) means that the sublineage cannot be identified. **d,** Schematic phylogeny of the lineages under R-L151 found in our dataset.

**Extended Data Fig. 5.**
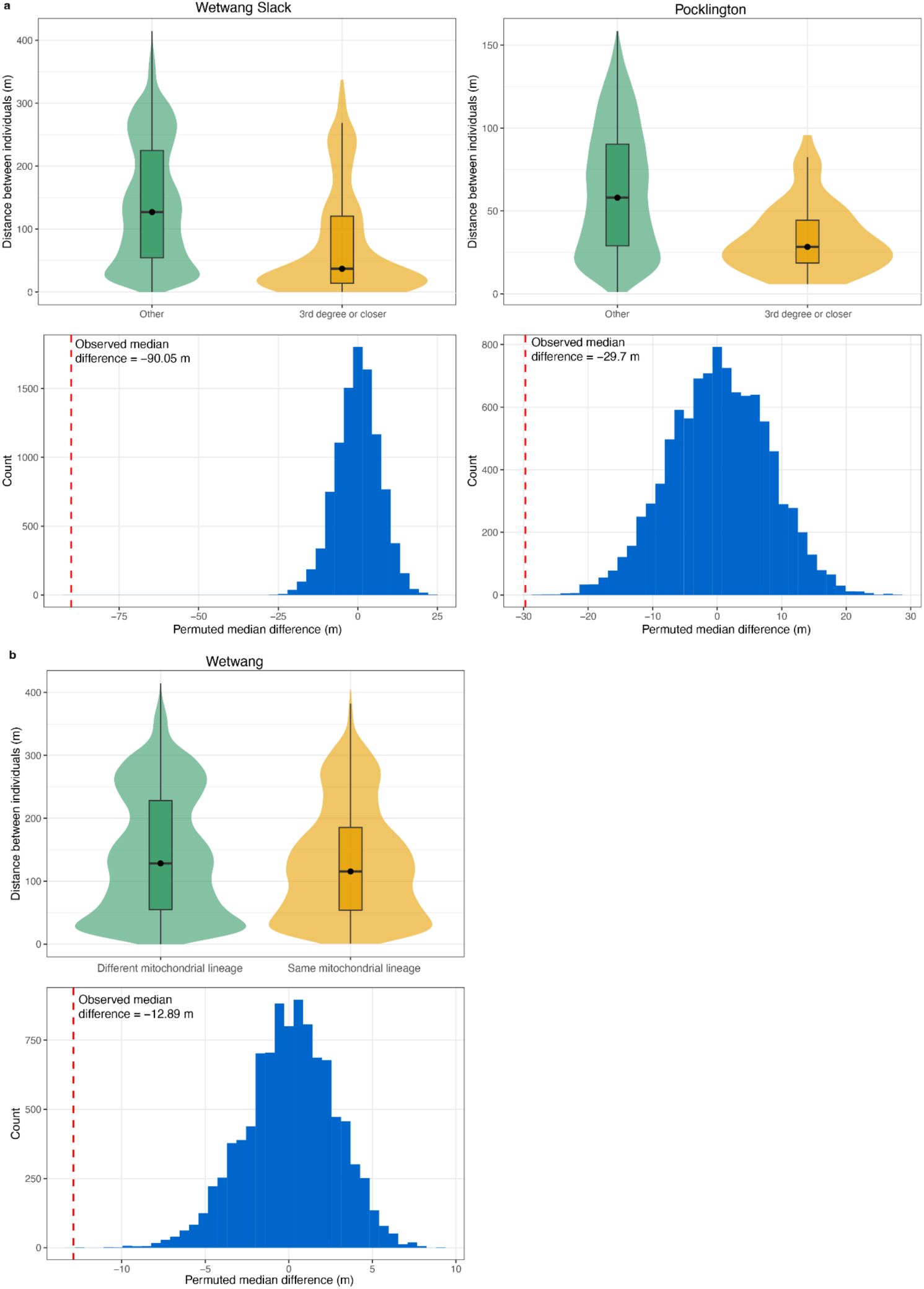
Kinship and maternal-line descent organisation of cemetery space. **a**, top: Distributions of burial distances between third-degree or closer relatives and other pairs at Wetwang Slack and Pocklington. Bottom: Histograms of the difference in median burial distance after 10,000 random permutations. The observed difference in median burial distance between third-degree or closer relatives and other pairs is shown with a red line. **b,** Distributions of burial distances between pairs sharing the same mitochondrial lineage and pairs with different mitochondrial lineages, after excluding close biological relatives up to third-degree.

**Extended Data Fig. 6.**
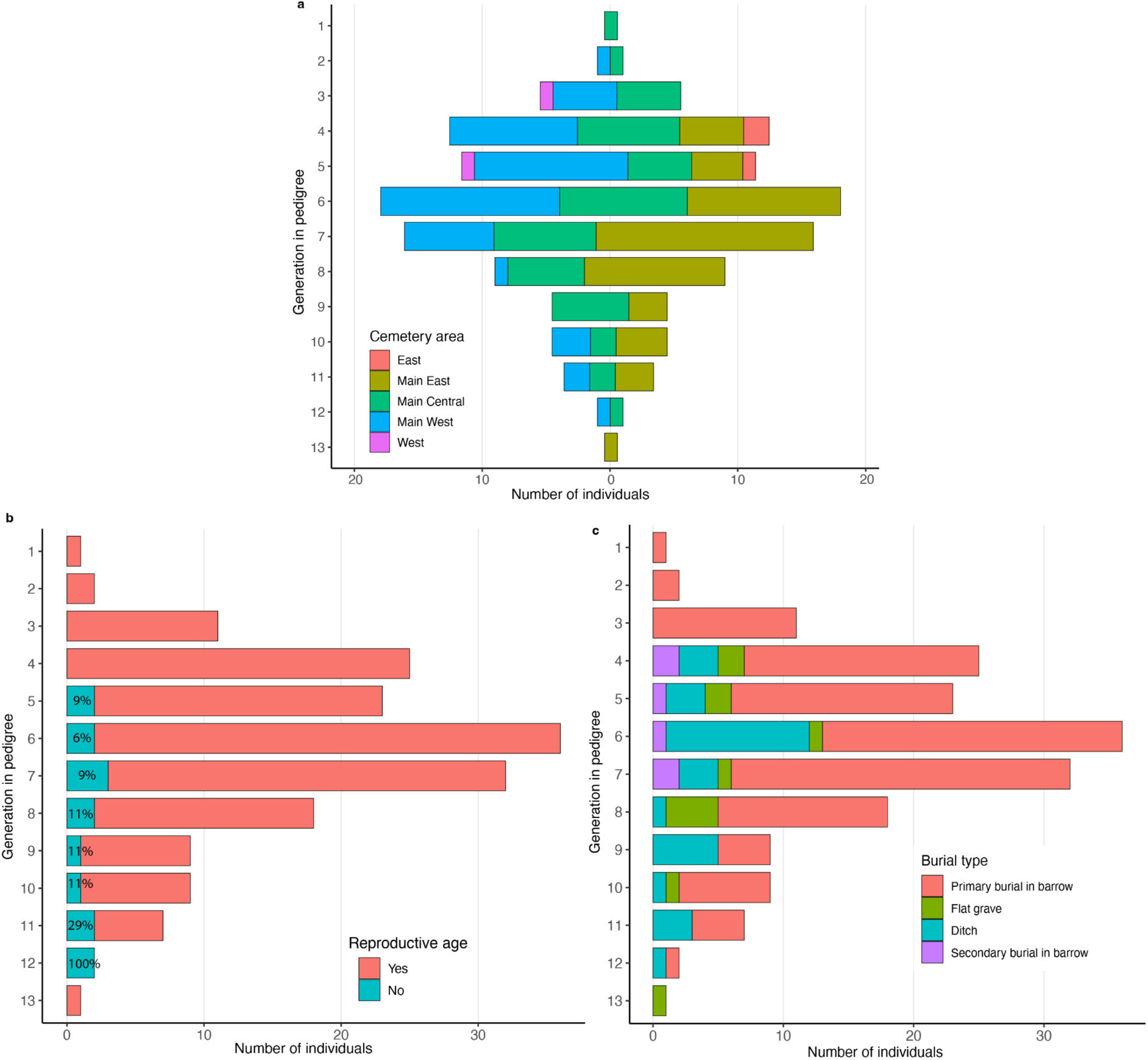
Distribution of individuals within the large family tree at Wetwang Slack. Number of individuals in each generation, coloured by **a,** location in the cemetery, **b,** reproductive age and **c**, burial type. The main burial area is divided into West, Central and East areas. The percentage of non-adult individuals is indicated in **b**.

**Extended Data Fig. 7.**
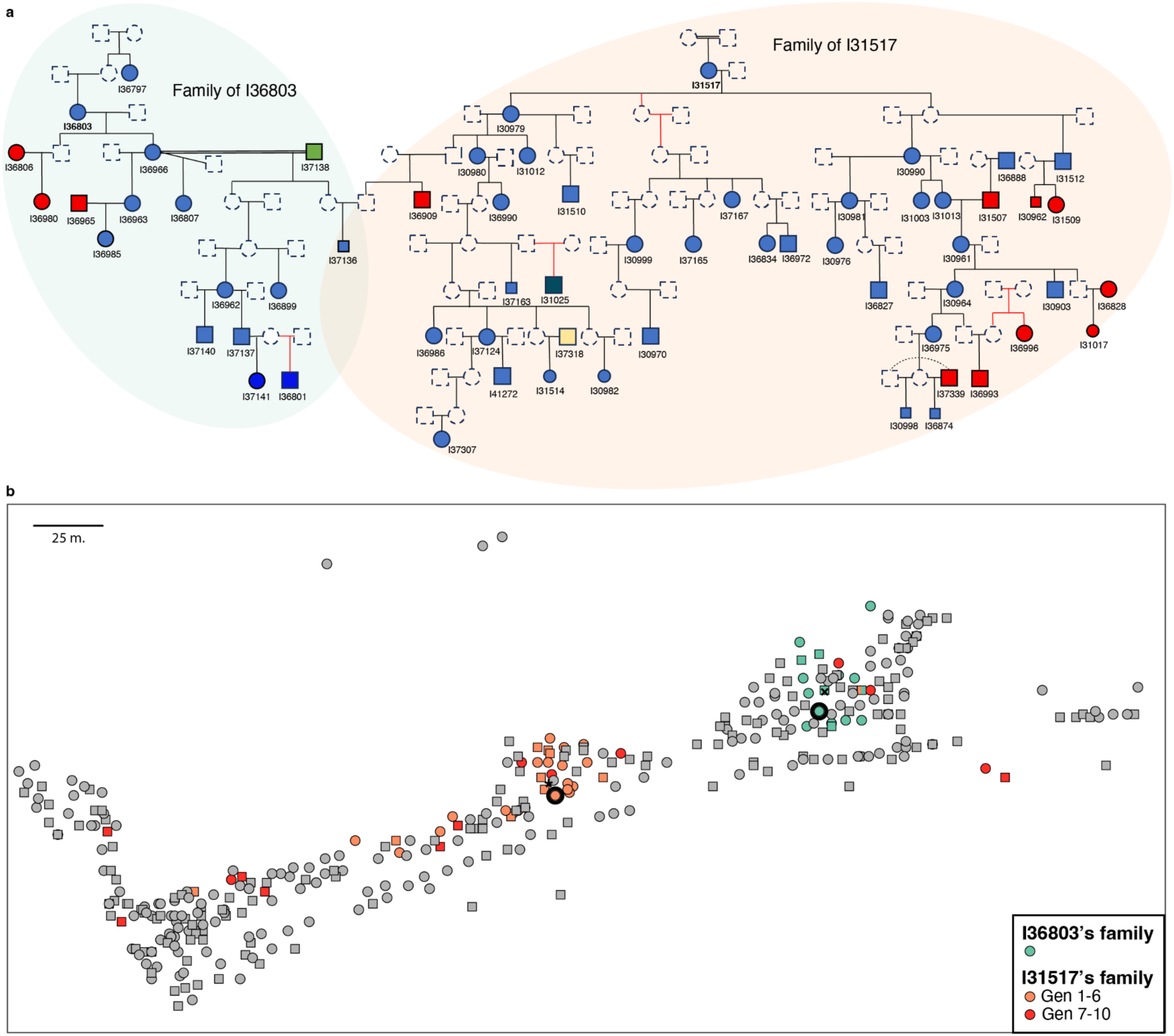
Multi-generational female-line families as spatial anchors at Wetwang Slack. **a**, Pedigrees of females I31517 and I36803. Red lines indicate uncertainty in tree topology (see Supplementary information). Double lines indicate a third-fourth-degree relationship between reproductive partners and dotted lines indicate a second-degree relationship. Colours represent different mitochondrial haplogroups. Smaller symbols represent non-adult individuals. **b,** Spatial distribution of the pedigrees of I36803 and I31517. We included the two females (highlighted with a thick black outline), their second-degree relatives, their descendants, and the reproductive partners of their descendants. Male individuals are represented by squares and female individuals by circles. Individual I37136 is a descendant of both females and is therefore shown using both colours. The median location of the relatives of I31517 and I36803 is indicated by an asterisk and a cross, respectively.

**Extended Data Fig. 8.**
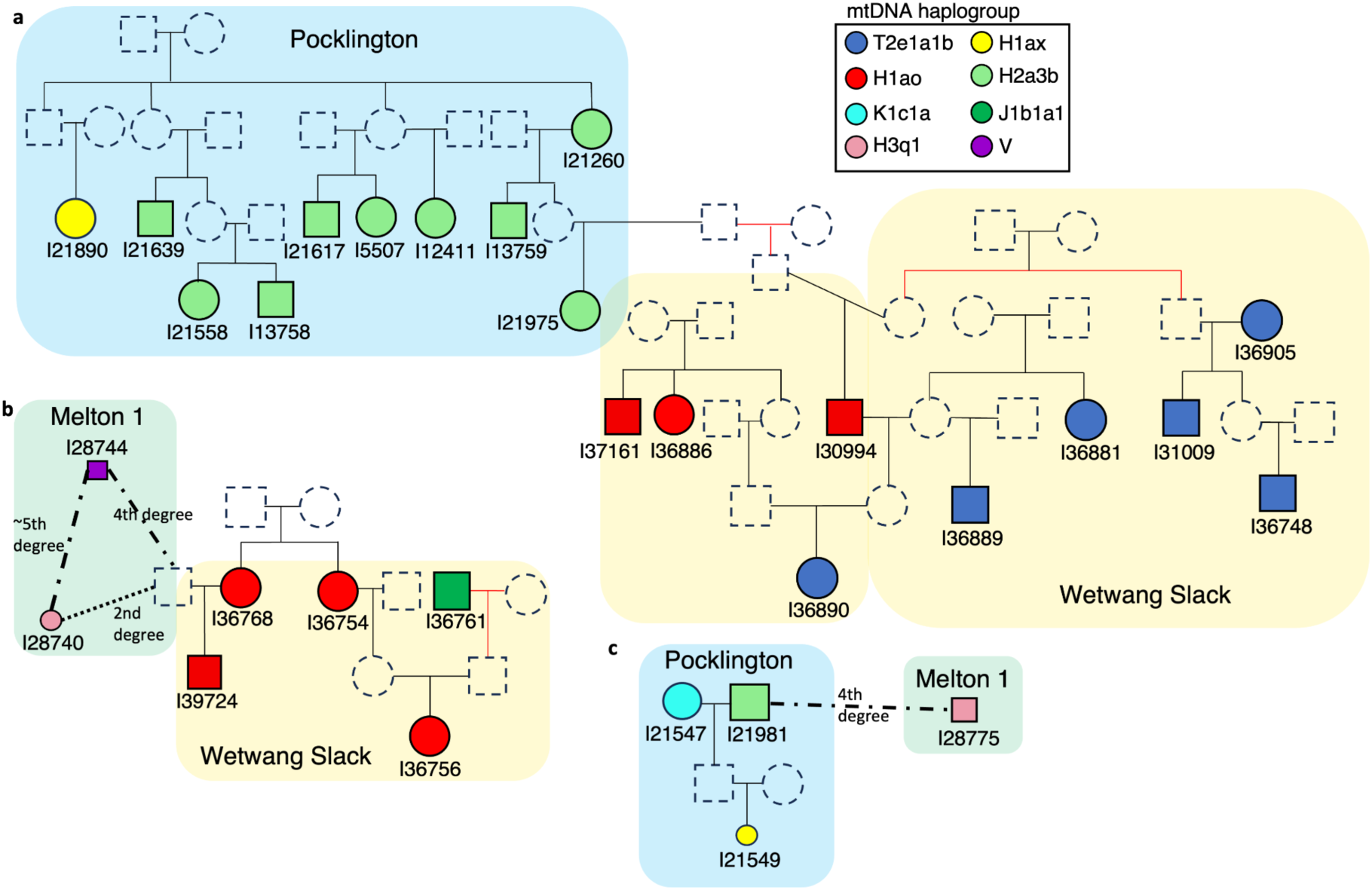
Cross-site genealogical connections among Arras Culture cemeteries. Reconstructed pedigrees linking individuals buried at different Arras Culture cemeteries, **a,** Pocklington–Wetwang Slack, **b,** Wetwang Slack–Melton 1, **c,** Pocklington–Melton 1. Red lines indicate uncertainty in tree topology (see Supplementary information). Smaller symbols represent non-adult individuals.

**Extended Data Fig. 9.**
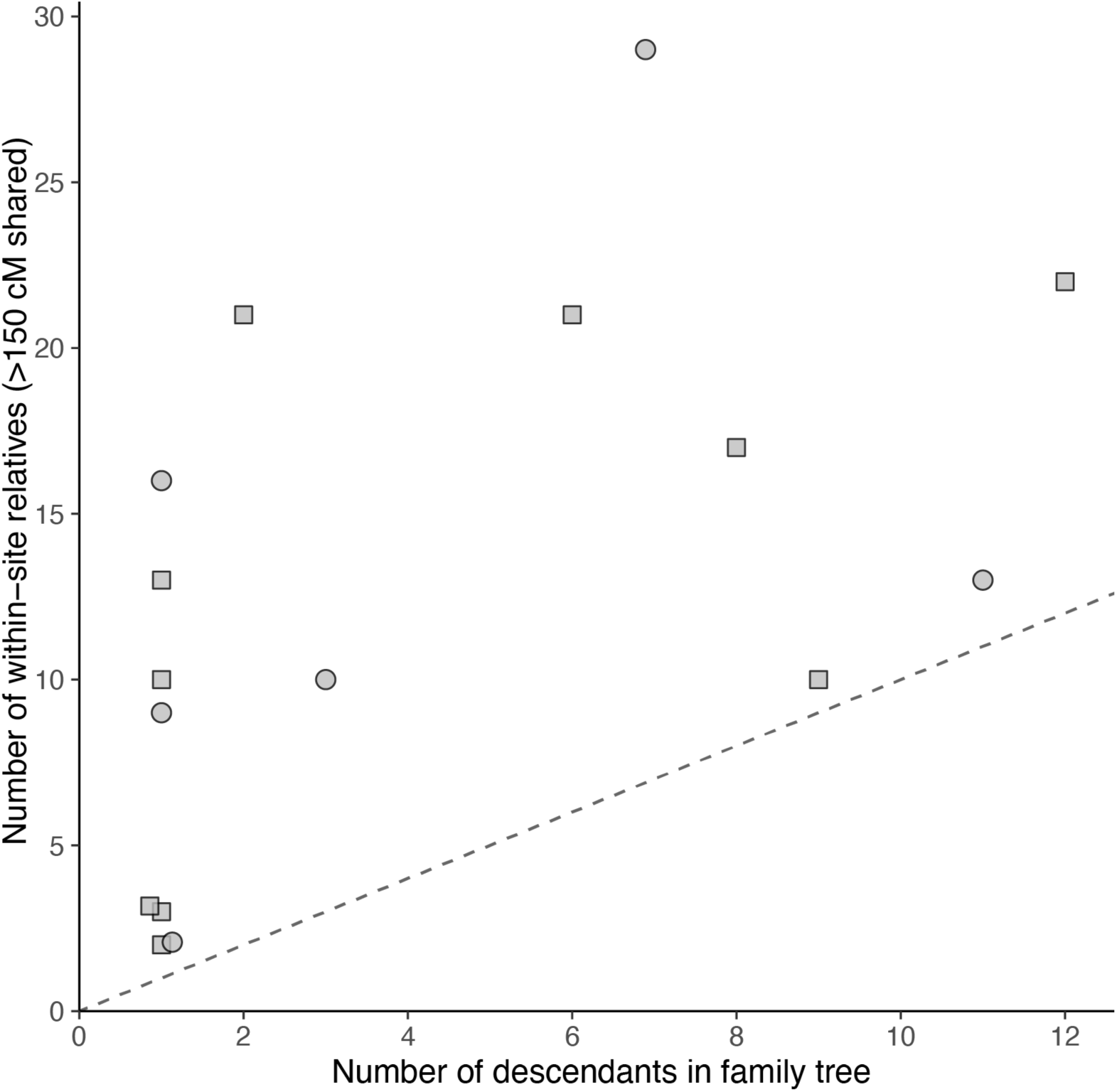
Intra-site relatives of reproductive partners at Wetwang Slack. Relationship between number of identified descendants and number of within-site relatives among reproductive partners at Wetwang Slack. Each point represents an individual reconstructed as a reproductive partner joining the Wetwang Slack pedigree. The x-axis shows the number of descendants identified in the reconstructed pedigrees, while the y-axis shows the number of within-site relatives sharing >150 cM in IBD. The diagonal line represents the expectation under which an individual’s only relatives in the cemetery are their descendants.

**Extended Data Fig. 10.**
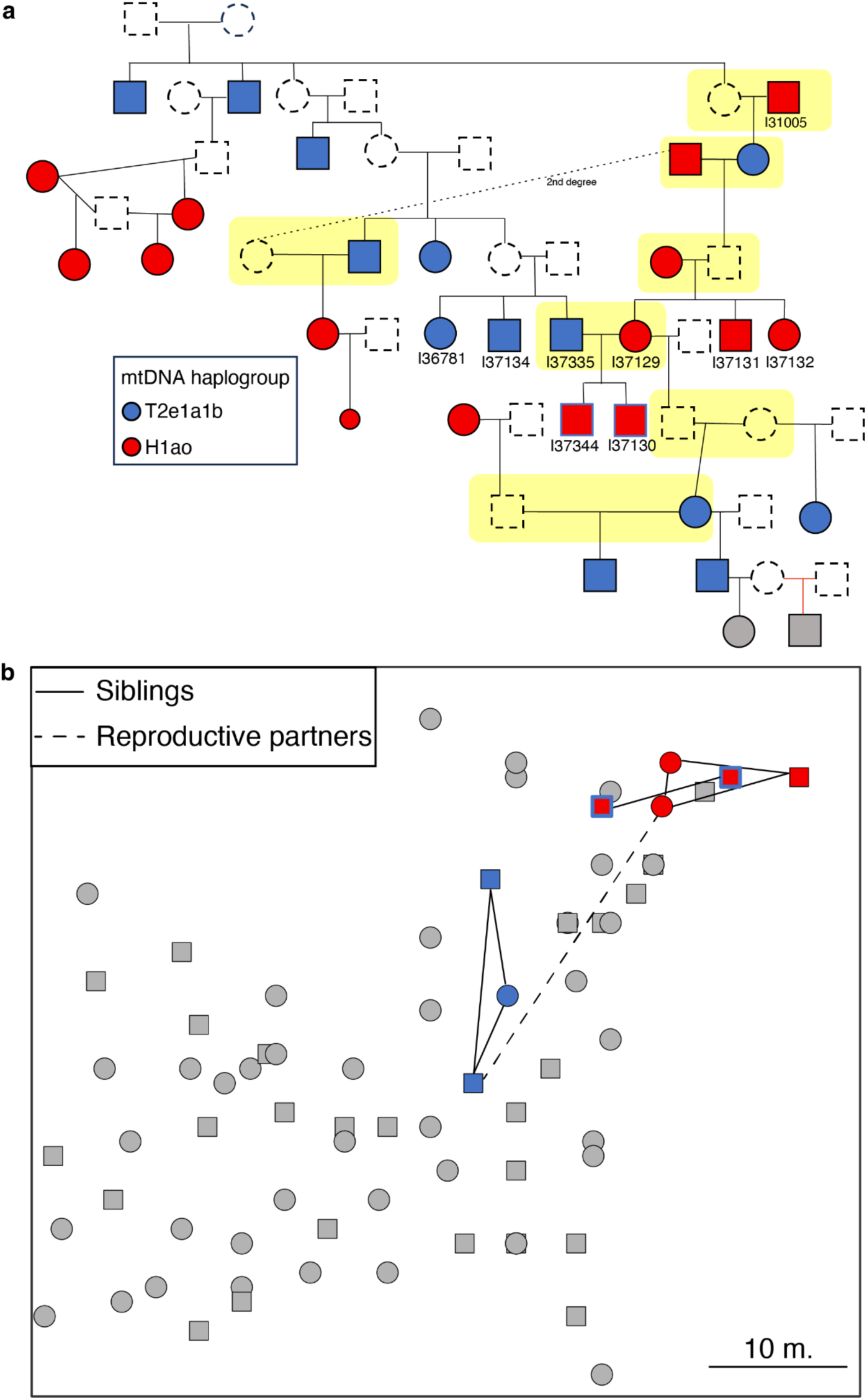
Recurrent unions between T2e1a1b and H1ao maternal lineages across consecutive generations. **a**, Detail of male I31005’s pedigree, showing repeated reproductive unions between individuals belonging to the two dominant Wetwang Slack maternal lineages, T2e1a1b (blue) and H1ao (red). Red lines indicate uncertainty in tree topology (see Supplementary information). Smaller symbols represent non-adult individuals. **b,** Detail of eastern edge of Wetwang Slack cemetery, showing how female I37129’s sons were buried closer to their mother and maternal aunt and uncle than to their father (I37335), paternal aunt and uncle. I37335 and I37129 are connected by a dashed line. I37335’s and I37129’s sons are displayed with a blue outline.

