## Supplementary Information for "Ancient DNA reveals matrilineal organisation and recurrent unions between dominant matrilines in Iron Age Britain"

#### Table of contents

SI 1. Archaeological background and context information about the sites with new genetic data

SI 2. Sampling, extraction, library preparation, capture, sequencing and bioinformatics processing

SI 3. Authenticity, sex determination and aneuploidies

SI 4. mtDNA and Y-chromosome analysis

SI 5. Kinship analysis and pedigree reconstruction

SI 6. Intra-site patterns of biological relatedness at the three main sites under study

SI 7. Inter-site patterns of biological relatedness across the Arras dataset

SI 8. Ancestry analysis

Legends of Supplementary Tables

#### S1. Archaeological background and context information about the sites with new genetic data

##### The Arras Culture

The ‘Arras Culture’, first named by Childe (1940: 16), refers to a distinctive Middle/Late Iron Age funerary tradition of individual inhumation in small square barrows, concentrated in East Yorkshire, and particularly on the Yorkshire Wolds (Stead 1991; Giles 2012; Halkon 2013). These barrows frequently cluster in extensive cemeteries containing several hundreds of burials dating broadly from the fifth to first centuries BCE (Stead 1991). While some survive as upstanding monuments, the great majority have been identified through aerial photography, where they are recognisable as small, quadrangular, ditched features, sometimes with a minority of circular or rectangular barrows. The cemeteries are set within heavily bounded landscapes of droveways, fields and enclosures, which also contain large numbers of non-nucleated roundhouse settlements (Stoertz 1997). The Arras Culture is unusual within the British Iron Age, where funerary traditions are generally highly varied and where large inhumation cemeteries are extremely rare (Harding 2016).

Arras Culture burials are generally found in cut graves on a north–south axis, in crouched or flexed positions (Stead 1991). Grave goods are found only occasionally, and are generally modest, comprising pottery vessels, brooches and other metalwork, and food offerings. A small minority of Arras Culture burials are accompanied by (usually dismantled) two-wheeled chariots, and these graves frequently contain a far richer array of diverse grave goods including weaponry and decorated metal items (e.g. Brewster 1971; Hill 2002). The occurrence of chariot burials and items decorated in La Tène style has led to long-running speculation over the degree to which the Arras Culture was related to La Tène communities in continental Europe in the second half of the first millennium BCE (Stead 1991). The burial positions (crouched rather than extended), dismantling of the chariots, and the co-occurrence of roundhouse settlement, however, are alien to contemporary continental communities and lie much more within British Iron Age traditions.

##### Wetwang Slack

Wetwang Slack, East Yorkshire, is the largest excavated Iron Age cemetery in Britain. It was excavated by archaeologist John Dent during rescue operations in advance of gravel quarrying, primarily between 1975 and 1979, with follow up investigations through to 1983 (Dent 1982, 1983, 1984, 1985; Armit 2021). The site is located in the central part of the Yorkshire Wolds, a crescentic area of rolling chalk uplands, and the nature of the underlying geology has led to excellent preservation of human remains. The excavations of the Iron Age cemetery formed part of a much longer and more extensive programme of archaeological work extending nearly 2 km along the dry valley (or ‘slack’) between the villages of Wetwang, to the west, and Garton, to the east. The eastern part of the excavations, known as Garton Slack, also contains scattered, smaller groups of Iron Age burials (including a further chariot burial), along with evidence for contemporary roundhouses and field systems; this was excavated by archaeologist Tony Brewster from 1965–1975 (Brewster 1971, 1980). The human remains from Garton Slack have not been analysed here.

The Wetwang Slack cemetery comprises approximately 240 square barrows (there are some uncertainties due to the levels of plough truncation), containing 446 burials (Dent 1984; King 2012; Armit 2021). The majority of the burials are crouched or flexed, as is normally the case in Arras Culture cemeteries. Only around 20% of the burials were accompanied by surviving grave goods (Dent 1984), and these were generally limited to dress fastenings or other small metal objects, pottery vessels and meat offerings (represented by animal bones). The majority of the barrows were tightly clustered (often conjoined and/or intercutting) and set along the course of a ditched droveway, extending for around 500 m. Aside from the primary, central burial within each barrow, numerous secondary burials had been inserted into the mounds or placed within the enclosing ditches, and apparently unmarked ‘flat’ graves were also present. Given the degree of both ancient and modern disturbance, it is likely that many more secondary burials originally existed, and even some of the central burials have been lost to the plough.

A short distance to the west of the main cemetery was a separate, smaller group of five barrows, of which three contained chariot burials: two male (Burials 453/I36995 and 455/I36892) and one female (Burial 454/I36978) (Dent 1985). Although grave goods were generally scarce in the main cemetery at Wetwang Slack, all three chariot burials were richly equipped. The female chariot burial (Burial 454/I36978), for example, was accompanied by an iron mirror and pin, two copper alloy and iron horse-bits, and the head of a broken iron and gold pin, as well as the forequarters of two pigs. Most notably, she also had a closed bronze canister, decorated with distinctive flowing, curvilinear motifs characteristic of La Tène art. This unique object has been known since its excavation as the ‘bean-tin’ (Hunter 2015: 103), and has variously been interpreted as a drum (Giles 2012: 158) or rattle (Jope 2000: 249). Both male chariot burials were interred with weaponry.

Human remains from the cemetery are held by the Biological Anthropology Research Centre (BARC) at the University of Bradford on long term loan from Hull Museums. The results of the original osteoarchaeological analysis of the assemblage, conducted by Jean Dawes in the 1980s, are available in the site archive (Armit 2021) and reassessment was more recently conducted by Sarah King (2012). Analysis of dietary isotopes on 62 human bone samples from Wetwang Slack identified a generally high terrestrial animal protein diet, with no significant variation between sexes, through time, or in relation to status as identified through grave goods (Jay 2005; Jay and Richards 2006). Strontium and oxygen isotope analysis on a small sample (three chariot burials and eight non-chariot burials) suggested that the majority of the cemetery population had spent their childhoods on the chalk landscape of the Wolds, though outliers were identified (Jay et al. 2013). Analysis of carbon and nitrogen isotopes in the bone collagen of 34 infants and children <6 years at death also demonstrated an unexpected pattern of early weaning at the site (Jay et al. 2008).

*Radiocarbon dating and chronological modelling: Wetwang Slack*

There are 22 radiocarbon results available from 20 of the burials at Wetwang Slack. These were previously presented by Jay et al. (2012) as part of a broader Iron Age East Yorkshire dataset. Here they are used on their own to develop an outline chronology of the burial activity at the site. The radiocarbon measurements were made at various times and include radiometric dates from AERE Harwell (HAR-) and accelerator mass spectrometry (AMS) radiocarbon dates from the Oxford Radiocarbon Accelerator Unit (OxA-) and the Scottish

Universities Environmental Research Centre (SUERC-). The samples dated at Harwell had the collagen extracted following the Longin (1971) method, combusted to carbon dioxide and synthesized to benzene using a method similar to that initially described by Tamers (1965) with a vanadium-based catalyst (Otlet 1977). The radiocarbon content was measured using Liquid Scintillation Counting as described by Otlet (1979). The samples submitted to Oxford were pretreated using a collagen extraction process involving acid demineralisation, gelatinization and separation by ultrafiltration (Longin 1971; Brown et al. 1988; Hedges et al. 1989; Bronk Ramsey et al. 2004). The samples were dated using Accelerator Mass Spectrometry (AMS) as described in Bronk Ramsey et al. (2004). The sample submitted to SUERC was pretreated and measured following methods described in Dunbar et al. (2016).

The chronological modelling follows a Bayesian approach (Buck et al. 1996; Hamilton and Krus 2016) and uses the simple bounded phase mode described in Hamilton and Kenney (2015). The model was produced using the computer software OxCal v4.4.4 (Bronk Ramsey 2009) and the IntCal20 calibration curve (Reimer et al. 2020). Modelled dates are presented rounded outward to the nearest 5 years. All calibrated dates have been rounded outward to the nearest 10 years.

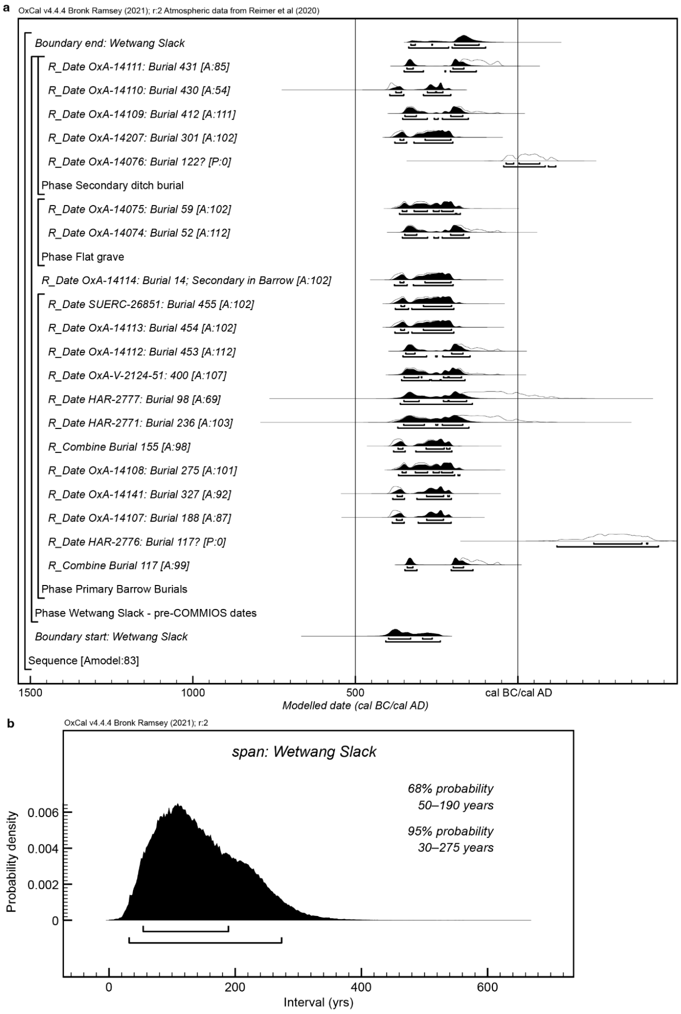


Figure S1. Visual outputs from the Bayesian chronological for Wetwang Slack on its own; a) the chronological model with the two Roman Iron Age results excluded from the overall model, such that the model is focused on the Middle Iron Age 'Arras' culture results; b) the span of Middle Iron Age burial at Wetwang Slack, as estimated from the model in S1a.

Two radiocarbon results that are too recent have been excluded from the modelling. HAR-2776 is Roman period in date, yet the more recent Oxford radiocarbon results from the same individual date to the Middle Iron Age. Furthermore, the two Oxford results are statistically consistent (T'=3.4; df=1; T'(5%)=3.8; Ward and Wilson 1978) and have been combined prior to calibration. The second Roman period date is OxA-14076, which is from Burial 122, a secondary burial in a ditch. This burial could signify later reuse of the site in the Roman period.

The model has good agreement (Amodel=83) and estimates the dated burial activity began in 410–235 cal. BCE (95% probability; Figure S1a; start: Wetwang Slack) and ended in 340–100 cal. BCE (95% probability; Figure S1a; end: Wetwang Slack). The overall period of modelled burial activity spanned 30–275 years (95% probability; Figure S1b; span: Wetwang Slack).

**Wetwang Village**

The Wetwang Village chariot burial was excavated in 2001 during excavations in advance of a housing development (Hill 2001, 2002). The site is located on high ground around 1.5 km south-east of the Wetwang Slack cemetery. The grave, which lay under a square barrow, contained the crouched remains of an adult female (I50780) with her head located, unusually, to the south, and with extensive pathological lesions. As well as the dismantled chariot, the burial was accompanied by pig remains and several objects including an iron mirror which had been deposited in a fur-lined bag closed with a string that had apparently been threaded with blue glass beads. A pig bone from the burial has been AMS-dated to 360 cal BCE–50 cal CE (95% probability; 2151±21 BP; OxA-11993) (Jay et al. 2012).

**Pocklington**

The site of Burnby Lane, Pocklington, East Yorkshire, at the foot of the Yorkshire Wolds, was excavated by MAP Archaeological Practice between 2014 and 2017 in advance of housing development (Stephens 2023). The Iron Age cemetery is located off the chalk in an area of permeable calcareous loamy soils over chalky gravel (Halkon 2023). The cemetery is of classic Arras type, comprising two principal groups (a larger western and smaller eastern group) of predominantly square barrows, with some round barrows and associated flat graves (Figure S2). A total of 82 barrows were identified during excavation, 71 of which contained surviving primary burials; a further eight secondary graves and 40 flat graves also contained burials (Stephens 2023). Many more graves are likely to have been lost through centuries of ploughing of the site, and no archaeological mitigation record exists for the earlier housing development immediately to the northwest of the cemetery area. Most burials were crouched or flexed, with heads to the north, facing east, although there were numerous exceptions. Only around 20% of the burials were accompanied by surviving grave goods, including brooches, bracelets and food offerings (ibid.: 265). Of the 119 individuals excavated, only 13 were non-adults.

Among the graves was a single chariot burial (Barrow 85; Skeleton 165: not available for aDNA sampling), containing a male individual outfitted with a sword, shield and spears. Unusually, the grave contained the articulated bones of two ponies as well as the *intact* body of the chariot.

Analysis of dietary isotopes on 54 individuals from Pocklington identified a diet based primarily on terrestrial animal protein and cereals (Hamilton et al. 2023). Analysis of sulfur isotopes from 31 individuals indicated that around a third had demonstrably moved between different geologies during their lives (ibid.).

Bayesian modelling of the radiocarbon dates from the burials, incorporating information from aDNA analysis, suggests that burial began in 470–395 cal. BCE (95% probability) and ended in 175–45 cal. BCE (95% probability) (Hamilton and Adams 2023: 207).

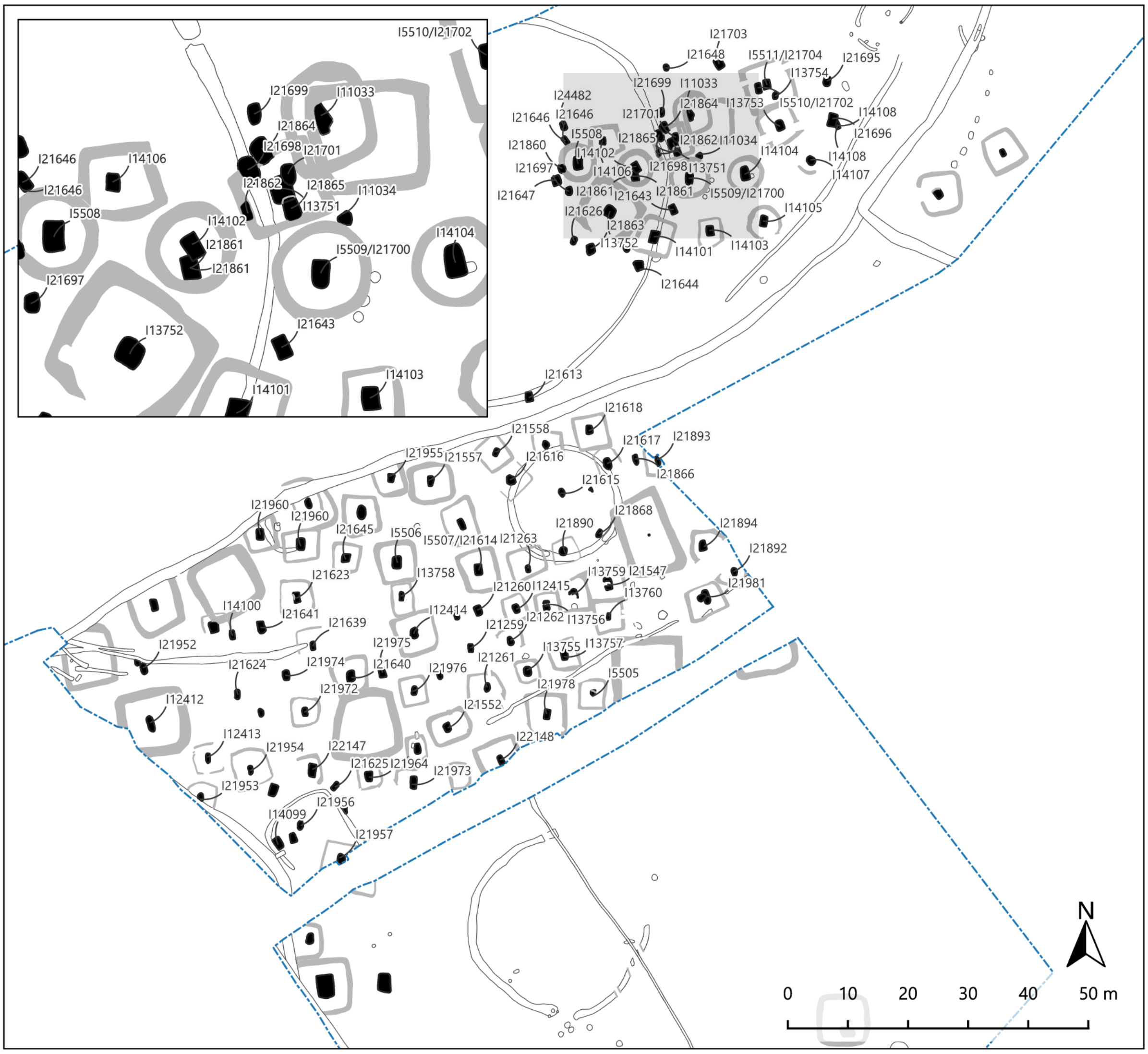


Figure S2. Plan of the Pocklington cemetery.

#### Melton 1

Excavations were carried out at this location by MAP Archaeological Practice between August 2014 and December 2016, and subsequently in February 2019, in advance of development. The site is located to the south-west of Melton, on the south side of the A63, in the East Riding of Yorkshire (Stephens 2019). It encompasses an area of 3.1 ha, and lies at an elevation of between 12 m OD to the south and 17 m OD to the north. The site is located off the chalk of the Wolds, with the bedrock geology comprising mudstone of the Ancholme Group. These are overlain by superficial deposits of sands and chalky gravels to the north and Devensian till to the south (BGS 2017).

The Iron Age cemetery identified at Melton 1 consists of nine Iron Age square barrows and six flat graves, alongside a number of secondary interments of probable Iron Age date (Stephens 2019: 28) (Figure S3a). In total, there are 44 individuals at Melton 1 that are considered to be of Iron Age date on the basis of burial architecture, mode of interment and associated grave goods. The inhumation burials included 24 adults (comprising 16 males and eight females), and twenty non-adults (comprising two adolescents, seven older juveniles, three younger juveniles, three infants, three neonates and two foetuses) (Ponce and Holst 2020: vii).

In terms of absolute chronology two individuals have been directly dated at Melton 1. Burial 27 (Sq. barrow 979) has been dated to 390–200 cal BCE (95% probability; 2229 ±29 BP; SUERC-77968), whilst Burial 7 has been dated to 380–200 cal BCE (95% probability; 2225 ±29 BP; SUERC-7796) (Stephens 2019:38).

Burial 27 (I28925) is a middle adult male who was buried with a dismantled chariot that comprised the chariot box, the axle and the pole, with associated copper alloy artefacts (terret rings, possible harness fittings or strap mounts and nave bands) and ferrous objects (bridle bit, tyres and lynch pins) (Stephens 2019: 38).

Burial 7 (I28771) is considered to be an exceptional burial deposit (Stephens 2019: 30), as three goats and three piglets were laid to the north of the inhumation. The goats overlay the piglets, and were placed on west–east alignments, with their heads to the east; two had been decapitated. The piglets lay on the reverse alignment to the goats, with their heads to the west. Additional pig and goat bones were present, suggesting that additional cuts of meat may have been present in this burial context (ibid.: 30).

A number of burials at Melton 1 were accompanied by funerary offerings, in addition to burials 27 and 7. Burial 2, for example, was a young adult female, and had an incomplete pig skull located next to her right elbow, and a copper alloy brooch in association. Burial 5 (I28769), an older middle adult female had a partial pig carcass adjacent to her knees on the east side of the grave. Three burials of probable Iron Age date are also of some interest due to the fact that they contain non-adults (secondary burials 21/I28779 and 22/I28922, and burial 33/I28741) with possible evidence for scurvy (burials 21 and 22) and/or scurvy/infection (burial 33); this is often cited as evidence for dietary stress in agricultural populations (Stephens 2019; Ponce and Holst 2020).

**Melton 2**

The site of Melton 2 is situated at Welton Common, Melton, between the southern tip of the Yorkshire Wolds and the river Humber, around 3 km to the east of Brough.
Excavation was undertaken in two short seasons in 2014 and 2015 by the University of Hull and the East Riding Archaeological Society, directed by Dr Peter Halkon and James Lyall. It lies approximately 1 km to the south of Melton 1 (see above).

In 2013, a magnetometer survey by James Lyall of Geophiz.Biz revealed a palimpsest of features including trackways, enclosures and buildings which subsequent excavation showed to date from the Iron Age to the mid–later Saxon period. These included a large square barrow around 9 m across, with a clear central anomaly. This proved to be a grave containing an adult female skeleton (I28926), only 35% complete, buried at a depth of around 1 m. With the head to the north, facing east and flexed, it was lying on its left side with its left arm straight and extended in front of the body. The right arm was bent at around forty degrees and the right hand positioned in front of the face. There was evidence of arthritis in the shoulders, wrists and knees (Loeffelmann and Holst 2016). During excavation, it was noticed that the right femur was covered with dull cut marks, running diagonally and horizontally across the entire shaft; these were similarly eroded to the rest of the cortical bone, suggesting their creation in antiquity. The skeleton was AMS dated to 540–380 cal BCE (95% probability; 2360±30 BP; Beta-437590).

**Burton Fleming**

The three individuals from Burton Fleming were recovered during a watching brief ahead of development for an agricultural building at West Hale Farm, Burton Fleming, East Yorkshire (Hunter, K. 2015). Initial topsoil removal revealed a series of pits, postholes, gullies, enclosure ditches, remains of four barrows (three with circular ditches and one with a square ditch), four inhumation burials and a pit containing disturbed human remains (ibid.: 6) (Figure S3b). The human remains consisted of four articulated individuals and two sets of disarticulated remains, with a minimum number of individuals (MNI) count for the site of six.

Three of the articulated skeletons were found in the centre of barrows; one square barrow (individual 65/I28928) and two round barrows (individuals 54/I28927 and 72/I28929). The fourth articulated skeleton was in a grave cut by the ditch of the square barrow that contained individual 65/I28928. The articulated skeletons were all crouched, and on their left sides; three were positioned along a north–south alignment, with their heads at the north end of the grave, while the fourth individual was placed with their head at the south-south-west and their feet pointing north-north-east.

The first of the two disarticulated individuals was found in a pit within a grave cut; it has been interpreted that the grave was disturbed at a later date and the remains moved into a pit in the north-east corner. Approximately 15% of the individual was recovered. The second disarticulated skeleton was found approximately 0.2 m above the grave cut into the square barrow. Very few bones were recovered from this individual, comprising predominantly one partial femur and one partial tibia.

All the individuals from Burton Fleming were excavated as part of features which have been interpreted as Iron Age, but of differing date, and all pre-date Roman features on the site, as suggested by previous phases of excavation and knowledge of the site (MAP 2016; Paris 2015).

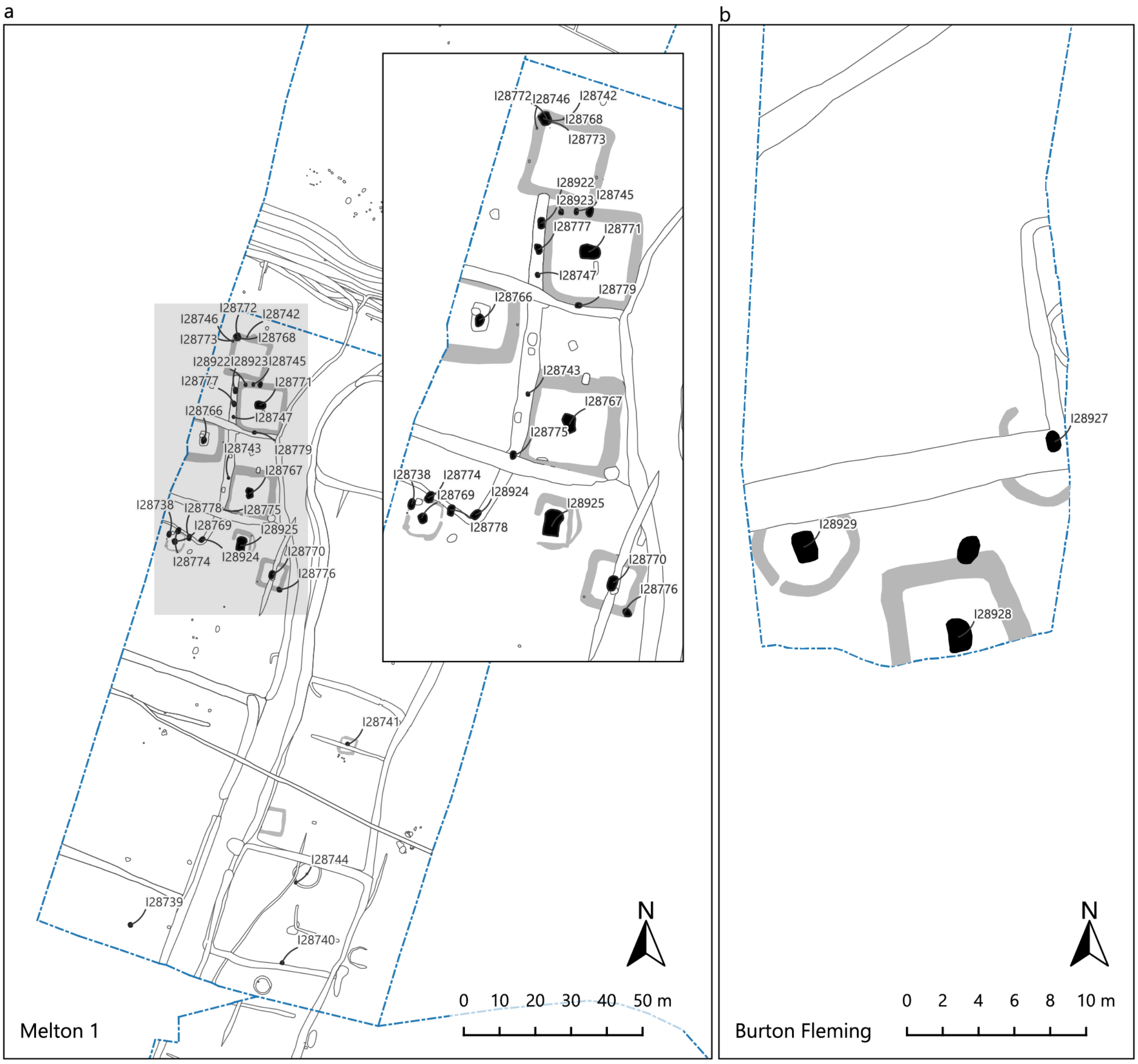


Figure S3. Plan of the a) Melton 1 and b) Burton Fleming cemeteries.

#### Ferry Fryston

An isolated square barrow containing a chariot burial was excavated in 2003 as part of a wider programme of development-led excavations along the line of the A1 (M) motorway in West Yorkshire (Brown et al. 2007). The site lay at around 20–25 m OD on ground sloping down towards the River Aire around 1 km to the north-east. The location is unusual in that it lies a considerable distance to the west of the main Arras Culture concentrations in East Yorkshire. Although only one barrow was present, other enclosures in the vicinity suggest that the site was part of a more extensive ritual and funerary complex.

The central grave comprised a large rectangular pit into which had been lowered the intact body of a chariot; this is unusual, since (unlike on the Continent) the great majority of Arras chariot burials are dismantled prior to burial. The chariot wheels were of unequal diameter, suggesting that it may have been assembled for the burial, rather than being a vehicle in use prior to the individual's death (Boyle et al. 2007: 127). Certain terret rings also appeared too weak for use, and may have been made (or co-opted) for the funeral (ibid.: 139). Laid within the chariot was the loosely flexed skeleton of an adult male (I50137) aged around 30–40 years at death. Two AMS determinations from the human remains provided a combined date of 355–110 cal BCE (95% probability; combined: 2185±35 BP, NZA-20494 and 2168±20 BP, NZA-19423-5) (Jay et al. 2012). Strontium isotopes for the individual, and for associated animal remains (see below), remain difficult to interpret (Jay and Montgomery 2020: 89) and, in this case, did not conclusively indicate an area of origin (Boyle et al. 2007: 129).

Aside from the chariot itself, with its various metal fittings, surviving grave goods were limited to an involuted brooch, made of iron, copper alloy and glass, worn at the shoulder (possibly implying that the corpse was dressed at the time of burial), and the forequarters of a pig. A collection of fragmentary metal fittings also hinted at the presence of a shield (ibid.: 145).

Remains of at least 25 cattle were recovered from the lower fill of the barrow ditch, suggesting one or more episodes of feasting, either at the time of the funeral and/or later (Boyle et al. 2007: 148–50). Remains of at least 162 cattle, dating predominantly to the Romano-British period, were recovered from higher in the ditch fills, suggesting that the monument continued to be venerated for several centuries (Orton 2007).

#### SI 2. Sampling, extraction, library preparation, capture, sequencing and bioinformatics processing

All laboratory procedures were carried out in dedicated clean-room facilities. For teeth and long bones, the external surface was removed and powder was drilled from the underlying cleaned surface, reducing the likelihood of exogenous DNA contamination. Drilling was performed at low speed to limit heat-associated damage to the DNA[1]. For petrous temporal bones, cochleae were isolated by sandblasting[2] and subsequently milled. The resulting powder was incubated in lysis buffer, after which DNA was purified and concentrated from one-fifth of the lysate. Extractions were performed either manually, using Dabney Binding Buffer[3,4], or with an automated silica magnetic bead-based protocol[5].

Ancient DNA libraries were generated using two library preparation strategies (Supplementary Table 1). Double-stranded barcoded libraries were built from the extracts using truncated adapters and were treated with partial ("half") uracil-DNA glycosylase (UDG) before blunt-end repair, substantially reducing the characteristic ancient DNA damage signal[6,7]. Single-stranded libraries were prepared using automated protocols based on Gansauge et al. and subjected to USER treatment[8].

DNA libraries were enriched for human DNA using oligonucleotide probes targeting either 1,233,013 nuclear SNPs, corresponding to the "1240k" capture set[9], or 1,352,535 nuclear SNPs using the Twist Bioscience reagent[10], together with the mitochondrial genome. Libraries enriched with the 1240k reagent underwent two rounds of capture, whereas libraries enriched with the Twist Bioscience reagent underwent a single round. Captured libraries were sequenced either on an Illumina HiSeq X10 platform using 2×101 cycles plus 2×7 cycles for dual-index reads[11], or on an Illumina NextSeq 500 platform using 2×76 cycles plus 2×7 cycles for index reads.

Sequencing reads were assigned to samples according to the indices introduced during laboratory processing, allowing up to one mismatch. Adapter sequences were trimmed and paired-end reads were merged into single sequences using a modified version of SeqPrep v.1.1, requiring a minimum overlap of 15 bp. Merging allowed either one mismatch between high-quality bases or up to three mismatches involving low-quality bases, and the highest-quality base was retained in the merged sequence at overlapping positions. Read pairs that could not be merged were excluded. Merged reads were aligned to the human reference genome hg19 and to the RSRS mitochondrial reference genome using the 'samse' command in BWA v0.7.15[12]. PCR duplicates were removed on the basis of alignment coordinates and read orientation. Finally, aligned data from multiple libraries generated from the same individual were merged into a single BAM file. The computational workflows used for these steps are available on GitHub (https://github.com/dReichLab/ADNA-Tools, https://github.com/dReichLab/adna-workflow).

#### SI 3. Authenticity, sex determination and aneuploidies

##### Authenticity analysis

We evaluated ancient DNA authenticity using multiple metrics: (i) a cytosine deamination rate at the terminal nucleotide >0.03; (ii) a ratio of Y-chromosome to combined X-chromosome + Y-chromosome 1240k targets with overlapping data <0.02 or >0.33; (iii) X-chromosome contamination estimates for sufficiently covered males <0.05 using ANGSD[13] and hapCon[14]; (iv) an upper bound of the 95% confidence interval for the mitochondrial consensus match rate >0.95 using contamMix-1.0.103[15]; and (v) an autosomal allelic mismatch rate not falling within the top 2% highest values observed in our dataset.

Applying these criteria, four individuals from Wetwang Slack (I36819, I36749, I42645, I36866) consistently showed evidence of contamination across multiple metrics (ranging between 13-22%) and were classified as "*fail*" (Supplementary Table 1). Nine additional individuals showed evidence of contamination in at least one metric and were classified as "*questionable*". In some cases—particularly when flagged solely based on autosomal mismatch rate—elevated values may reflect ancestry differences rather than substantial contamination.

Because contamination typically reduces estimated relatedness with true relatives but does not generate spurious kinship links, we retained the 13 individuals labelled as "*fail*" or "*questionable*" for kinship analyses. The extensive pedigree structure reconstructed at the site allows reliable placement of several of these individuals despite contamination (see SI 5).

Below we discuss four individuals with weak evidence of mitochondrial contamination:

-Individual I36869 belongs to haplogroup V with additional mutations 2531C and 3604A, and exhibits heteroplasmies 709A (43%) and 9629G (54%) (Supplementary Table 1). The estimated consensus match rate is [0.888, 0.932], which is driven by these heteroplasmic positions. We interpret these variants as genuine heteroplasmies rather than contamination because:

> -Haplogroup-defining positions only show 1% ancestral alleles, including 1/212 at the terminal mutation 3604A. If 709A and 9629G alleles reflected contamination from a different individual, the contaminant would need to belong to the same V+2531C+3604A haplotype present in I36869, which is highly unlikely.
>
> -A contamination level sufficient to explain ~50% discordance at these positions would be expected to severely affect other authenticity metrics (sex ratio, X-chromosome contamination estimates, damage patterns), which is not observed.

-Female I21643 belongs to haplogroup J1c9 and has a mitochondrial consensus match rate of [0.873, 0.969]. Haplogroup-defining positions (J ,J1 ,J1c ,J1c9) show only seven ancestral alleles out of 620 reads (1%). In the absence of other contamination signals, we interpret this estimate as reflecting statistical uncertainty near the upper boundary of the confidence interval rather than genuine contamination.

-Female I39729 belongs to haplogroup J1c9 with additional mutation 189G and a heteroplasmy at 16193T (45%), likely explaining the reduced consensus match rate [0.903, 0.974]. Haplogroup-defining positions (J ,J1, J1c ,J1c9, 189) show 20 ancestral alleles out of

1,322 reads (1.5%), and no other authenticity metrics indicate contamination. We therefore consider 16193T a true heteroplasmy.

-Female I50780 belongs to haplogroup H1ao and has a mitochondrial consensus match rate of [0.905, 0.945]. Haplogroup-defining positions (H, H1, H1ao) only display 10 ancestral alleles out of 622 reads (1.6%). Given the absence of additional contamination signals, we interpret this estimate as reflecting statistical variation within the confidence interval rather than significant mitochondrial contamination.

#### Sex determination

Genetic sex was inferred by assessing the relative representation of Y-chromosome sequences. Specifically, we calculated the ratio between the number of 1240k targets on the Y-chromosome with overlapping sequences relative to the total number of 1240k targets with overlapping sequences on the X- and Y-chromosomes. Individuals with a ratio greater than 0.33 were classified as genetic males, whereas those with a ratio below 0.02 were classified as genetic females (Supplementary Table 1).

#### Detection of aneuploidies

We calculated, for each autosomal chromosome, the ratio between its mean coverage and the genome-wide mean autosomal coverage, restricting the analysis to 1240k sites. The same approach was applied to the sex chromosomes, and although no individuals in the present dataset were identified as carrying sex chromosome aneuploidies, an individual with Klinefelter was identified previously at Wetwang Slack (WS 224)[16]. Using this framework, we identified one instance of Down syndrome: a newborn male (I30978; WS267) from Wetwang Slack showing 1.48× higher mean coverage on chromosome 21 relative to the autosomal average (Figure S4), consistent with trisomy 21[16].

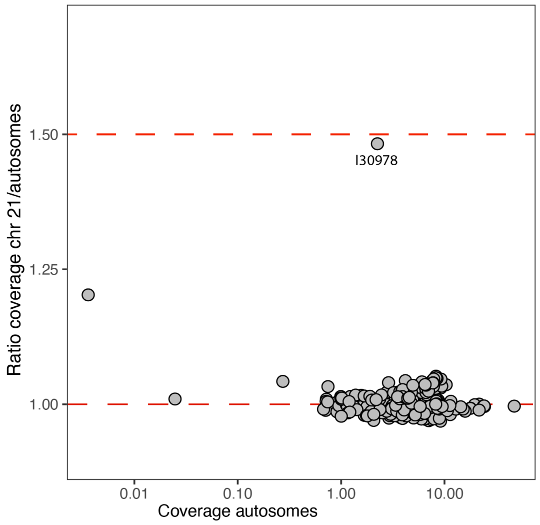


Figure S4. Ratio between the mean coverage at chromosome 21 and the mean autosomal coverage. The x-axis represents the mean autosomal coverage on a logarithmic scale.

#### SI 4. mtDNA and Y-chromosome analysis

To determine mitochondrial and Y-chromosome haplogroups, we restricted the analysis to reads with mapping quality >30 and base quality >30. Mitochondrial haplogroups were assigned using HaploGrep 3 (v3.2.1)[17], after generating a consensus sequence with bcftools[12]. Mutations at known mutational hotspots (309, 315, 16182, 16183, 16193 and 16519) were excluded for phylogenetic reconstruction. For Y-chromosome haplogroup assignment, we primarily used the YFull v.14.01 phylogeny (https://www.yfull.com/) to identify the most derived branch in each male individual, following the approach in Lazaridis *et al.*[18]. Haplogroup calls were manually curated and mapped onto the International Society of Genetic Genealogy (ISOGG) tree (version 14.76, 25 April 2019; http://www.isogg.org) to obtain the corresponding traditional 'long' haplogroup names. Because the YFull phylogeny is continuously updated, some terminal branches are not represented in the ISOGG tree. In such cases, we moved upstream in the YFull phylogeny until a corresponding ISOGG-defined branch was identified. Mitochondrial and Y-chromosome calls for each individual are included in Supplementary Table 1.

##### *Mitochondrial patterns*

We identified 144 distinct mitochondrial haplotypes among the 524 individuals from sites with newly reported data, of which only 10 were shared between sites: one between Wetwang Slack and Wetwang Village, one between Melton 1 and Melton 2, five between Wetwang Slack and Pocklington, two between Burton Fleming and Wetwang Slack, and one between Melton 1 and Wetwang Slack. Most haplotypes fall within typically European haplogroups, including H, T2, U5, K1, and J. We did not detect African-associated haplogroups such as L or U6, nor lineages commonly found in Asia, including C, D, or M.

Each of the three main sites (Wetwang Slack, Pocklington and Melton 1; reported below) we analysed was dominated by two (Wetwang Slack) or three (Pocklington and Melton 1) haplogroups, together representing more than 50% of the individuals at each site. Interestingly, none of these eight major haplogroups was shared across these three main sites, with the exception of four Wetwang Slack individuals who harboured the same J1c9 haplotype present at high frequency (12%) at Pocklington. Three of those Wetwang Slack individuals are a pair of siblings and their second-degree relative, who are among the Wetwang Slack individuals with the highest fraction of related Pocklington individuals (third, fourth and fifth position among the 382 Wetwang Slack individuals with IBD data; Supplementary Table 1). In fact, their closest Pocklington relative, sharing ~100 cM in IBD segments >12 cM with the siblings, belongs to the same J1c3 haplotype. Although the mitochondrial pools at these main sites are largely non-overlapping, this could represent a case of a female-mediated movement from Pocklington to Wetwang Slack.

###### ***Wetwang Slack***

At Wetwang Slack, we identified 107 distinct mitochondrial haplotypes in 390 individuals, of which 76 were observed only once. Two major haplogroups, T2e1a1b and H1ao, dominate the site, with frequencies of 33% and 18% respectively (Figure S5). Both lineages are rare in present-day populations and, to our knowledge, have not been reported in ancient populations elsewhere, with the exception of a single H1ao individual from Medieval Greenland[19].

We divided individuals into two groups based on intra-site relatedness: 68 individuals with less than 2% of relatives at the site (62% female) and 314 with more than 2% (55% female). Pairs were considered relatives if they shared >24 cM in IBD across more than two segments. The combined frequency of T2e1a1b and H1ao haplogroups differs markedly between these groups, reaching 10% in the low-relatedness group and 62% in the high-relatedness group (Figure 2b). Although the low-relatedness group represents only 18% of the total number of individuals, it displays 61 distinct haplotypes, compared to 62 in the much larger high-relatedness group. This indicates that the Wetwang Slack burial community comprised a large, biologically related group characterised by relatively lower mitochondrial diversity, alongside a smaller set of largely unrelated individuals exhibiting substantially higher mitochondrial diversity.

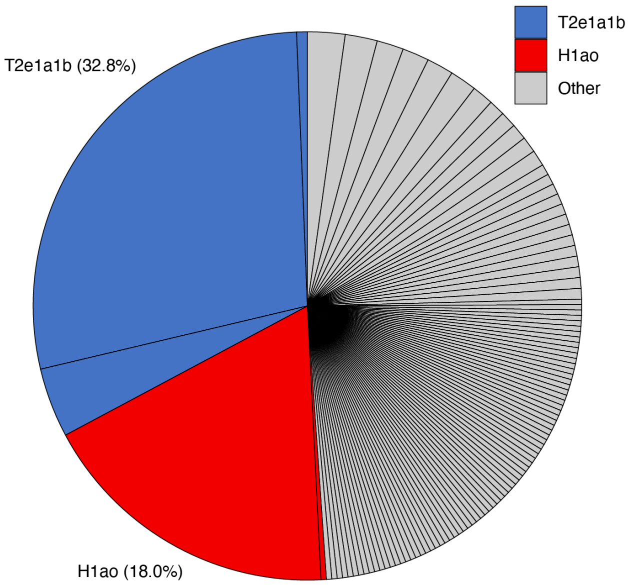


Figure S5. Mitochondrial haplotype frequencies at Wetwang Slack. Haplotypes belonging to the two major haplogroups at the site are coloured. Here, and throughout the text, haplotype frequencies were calculated retaining a single individual per cluster of first-degree relatives to avoid over-representation of large sibships. Frequencies computed without this filtering showed minimal differences.

##### *Haplogroup T2e1a1b*

A total of 142 individuals from Wetwang Slack belonged to the maternal lineage T2e1a1b. They can be assigned to three different haplotypes (Figure S6):

- T2e1a1b+14180C: Detected in two infants, I36872 and I37313. I36872's closest relative is I37313 itself (~fifth degree), while I37313's closest relatives are 30905 (fourth degree) from Wetwang Slack and I21259 from Pocklington (~fifth degree).
- T2e1a1b+14180C+4107T+9181G: Observed in 16 individuals, including I30905 and her close relatives in cluster 6 (buried in close proximity at the south-western corner of the main cemetery), as well as two brothers from cluster 4.
- T2e1a1b+14180C+4107T+2416C: Found in 124 individuals distributed across all major relative clusters, making it the most frequent haplotype at Wetwang Slack.

T2e1a1b has not been reported in other Iron Age individuals from Britain and, in fact, in any other ancient individuals so far. The presence of three closely related haplotypes at the site therefore suggests that their most recent common matrilineal ancestor—carrying the ancestral

T2e1a1b+14180C haplotype—likely lived locally prior to the formation of the documented pedigree structure. These patterns are very similar to those at the Iron Age site of Winterborne Kingston[20], southern England, where members of the dominant mitochondrial haplogroup carried five different closely related haplotypes.

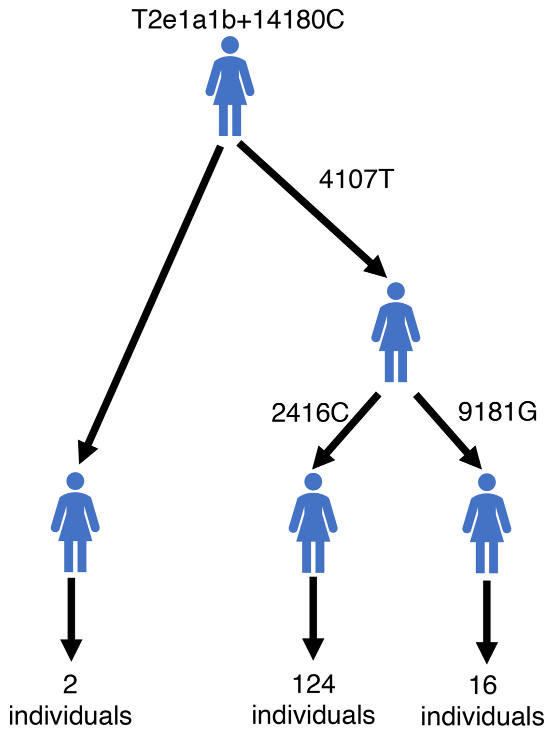


Figure S6. Phylogeny of the three T2e1a1b haplotypes at Wetwang Slack.

Individuals carrying the predominant T2e1a1b+14180C+4107T+2416C haplotype exhibit several heteroplasmies (Supplementary Table 1). Notably, positions 146 and 195 in I36803 and her descendants illustrate how heteroplasmic variants can become fixed over time and ultimately incorporated as stable mutations within a lineage (Figure S7). In I36803, two mitochondrial read populations are observed: 9% of reads carry 146C (with 195T), whereas 91% carry 195C (with 146T). Her maternal second-degree relative, I36797 (aunt or half-sister), displays the expected allelic state at both positions (146T and 195T), along with a minor fraction (~5%) of reads carrying 195C, indicating that I36803's mother likely already harboured, at the very least, the 195C heteroplasmy. I36803's maternal descendants retain both read populations, generally with a higher proportion of 195C reads. In fact, after five generations, the 195C variant becomes fixed in one of I36803's great-great-great grandsons, I37140, but not in his maternal half-brother who still displays both populations of reads. In a separate branch of I36803's matriline, the 146C variant becomes fixed in her great-granddaughter I36985. As a result of this process, I37140 and I36985—both strict maternal-line descendants of I36803 and sixth-degree relatives of each other—carry haplotypes that differ by two mutations. Because the emergence and fixation of these variants can be traced within the reconstructed pedigree, we treated these individuals as sharing the same haplotype for the purposes of mitochondrial diversity estimates.

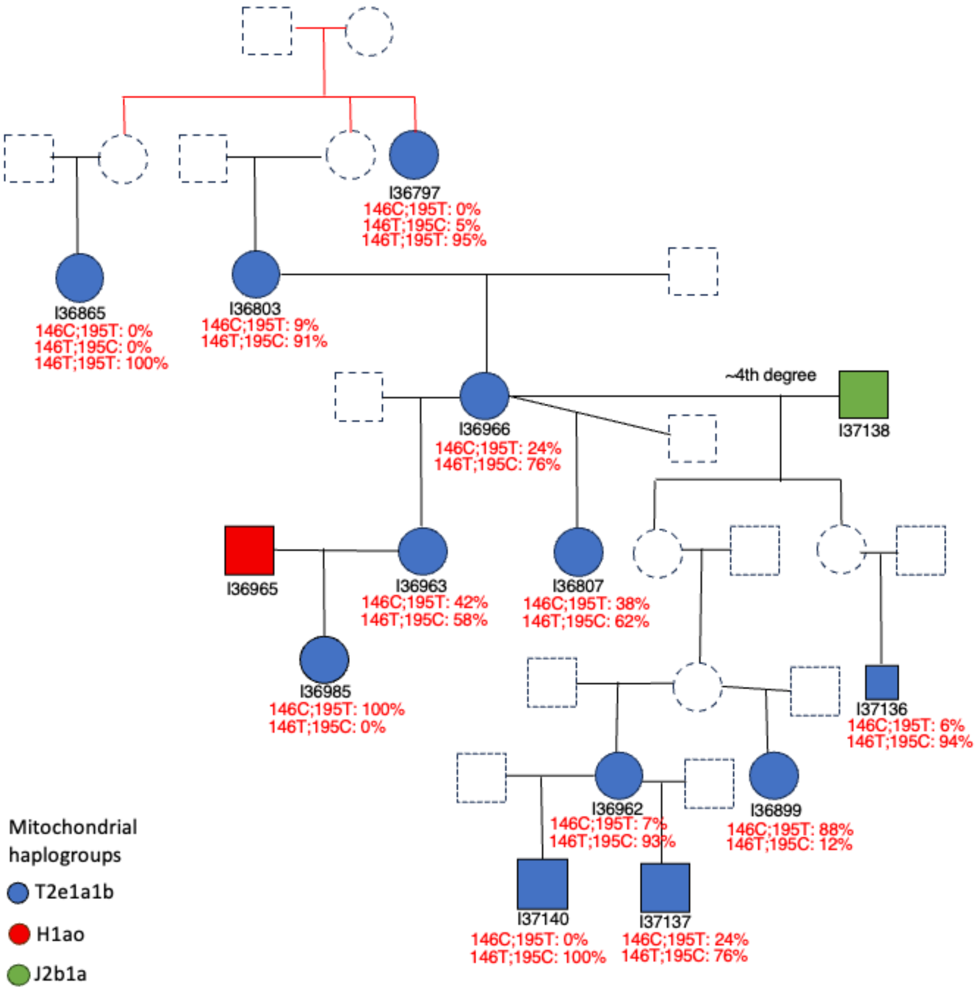


Figure S7. Female I36803's genealogical pedigree. Proportion of mitochondrial reads carrying mutations 146C and 195C are shown for each individual.

##### *Haplogroup H1ao*

A total of 80 individuals at Wetwang Slack belonged to haplogroup H1ao. All carried the same haplotype without additional mutations, with the exception of female I36839 who carried one additional mutation (14858A) and who stands out as the H1ao individual with the lowest number of relatives at the site (Supplementary Table 1). Individuals bearing the predominant H1ao haplotype appear across multiple branches of the main pedigrees and, together with T2e1a1b, this group represents a key matriline within the Wetwang Slack burial community. Consistent with its importance at nearby Wetwang Slack, the individual from the Wetwang Village chariot burial also carries this predominant H1ao haplotype.

#### ***Pocklington***

At Pocklington, we identified 28 distinct mitochondrial haplotypes among 100 individuals, 16 of which were singletons (Figure S8). Three major haplogroups—H2a3b, K1c1a, and J1c9—account for 70% of the individuals. The most frequent lineage, H2a3b, is represented by a single haplotype reaching 34% frequency and forms the backbone of the largest reconstructed pedigree at the site (Figure S43). The second most frequent haplogroup, K1c1a, comprises five different haplotypes (Figure S9). As at Wetwang Slack, the apparent absence of this lineage in previously reported ancient individuals from Britain suggests that the diversification of the K1c1a haplotypes observed here may have occurred locally. As mentioned above, the third most frequent haplogroup, J1c9, is shared with Wetwang Slack.

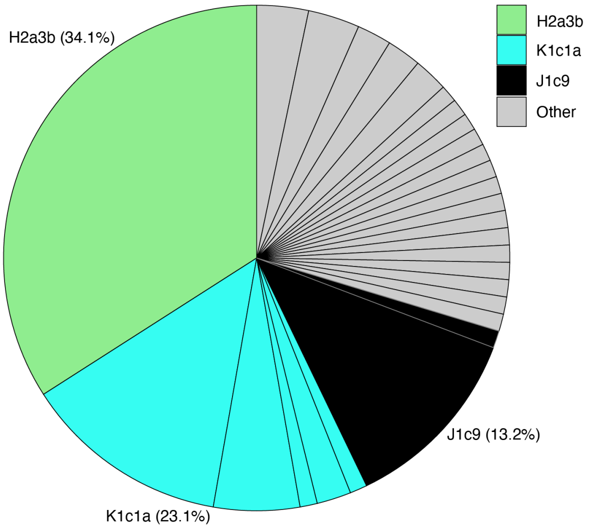


Figure S8. Mitochondrial haplotype frequencies at Pocklington. Haplotypes belonging to the three major haplogroups at the site are coloured.

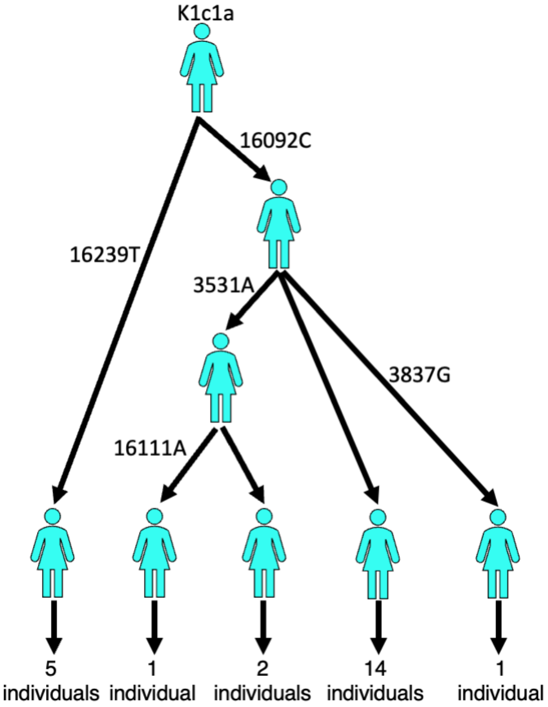


Figure S9. Phylogeny of the five K1c1a haplotypes at Pocklington.

##### ***Melton 1***

At Melton 1, we identified 13 distinct mitochondrial haplotypes in 28 individuals, of which eight were singletons (Figure S10). As at Pocklington, three haplogroups—H3q1, U2e1e and V—dominate, with a frequency of 58%. The only individual analysed from Melton 2 carried the same major H3q1 haplotype.

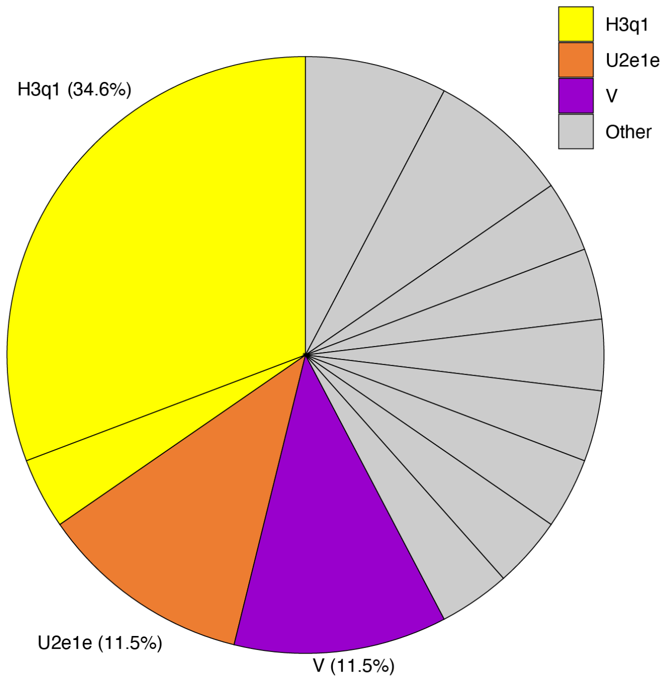


Figure S10. Mitochondrial haplotype frequencies at Melton 1. Haplotypes belonging to the three major haplogroups at the site are coloured.

##### *Y-chromosome patterns*

We identified at least 78 distinct Y-chromosome lineages among the 238 newly reported males. This represents a minimum estimate of the true number of distinct Y-chromosome haplotypes in the dataset for two main reasons. First, missing data at key diagnostic SNPs sometimes prevent precise subclade assignment. For example, although several individuals are positive for L21, incomplete coverage at downstream markers does not allow us to determine whether they belong to the same L21 sub-branches. Second, unlike the mitochondrial genome—which is typically recovered in full and allows exact haplotype determination—Y-chromosome data are limited to reads mapping to the ~82k Y-SNP targets included in the Twist capture array (or ~33k in the 1240k capture dataset), as well as nearby positions. Consequently, males assigned to the same terminal branch at our level of resolution may in fact belong to distinct sublineages within that branch.

The most common Y-chromosome haplogroup in the dataset is R-P312 (89% frequency), particularly its sublineage R-L21 (62% frequency), which rose to high frequency in Britain during the Beaker period[21]. This pattern is consistent with a substantial degree of continuity in the paternal gene pool across the Bronze and Iron Ages in Britain.

At Wetwang Slack (n=149), 58% of men belonged to at least 39 distinct sublineages within R-L21-DF13 (Extended Data Figure 4). Outside R-P312, 2% carried its sister lineage R-S1194, and 8% belonged to haplogroup I2, including six individuals assigned to I2-L1195-Y3684. Lineage I2-L1195 was common in Neolithic Britain[22] and its presence here indicates that some Neolithic-associated paternal lineages persisted, despite the major Y-chromosome turnover documented during the second half of the third millennium BC[21].

At Pocklington (n=44), the Y-chromosome composition was broadly similar to that observed at Wetwang Slack, with 43% of males carrying R-L21-DF13 lineages (Extended Data Figure 4). At Melton 1 (n=16), the proportion of R-P312-U152, which was widespread in central Europe during the Bronze and Iron Age periods[21,23,24], was higher than at the two other sites, reaching 25% and comprising four distinct sublineages (Extended Data Figure 4).

##### *Estimates of intra-site mitochondrial and Y-chromosome diversity in Britain and Ireland*

To investigate changes in mitochondrial and Y-chromosome diversity in Britain and Ireland across the Neolithic, Bronze Age and Iron Age, we compiled all previously published archaeological sites from Britain with at least two individuals with high quality data (>600k 1240k SNPs with overlapping reads) (Supplementary Table 2). Mitochondrial and Y-chromosome haplogroups were assigned using the same procedures applied to the newly reported individuals in this study.

Mitochondrial haplotypes were considered identical when individuals belonged to the same haplogroup and shared all additional mutations, excluding known mutational hotspots. For the Y-chromosome, we excluded individuals for whom missing data prevented us from determining whether they belonged to downstream branches observed in other individuals from the same site (Supplementary Tables 1 and 2). For example, at Wetwang Slack, we retained R-Z2183 individuals assigned to FGC35529, FT32305, and xFGC35529,xFGC53695, but excluded two individuals with missing data at the SNPs defining the terminal branches FGC35529 and FT32305 (Supplementary Tables 1 and 2). Following this, if more than one individual assigned only to broad terminal clades such as R-P312, R-L21, or R-DF13 remained at a given site, we retained a single representative. These haplogroups encompass numerous distinct downstream lineages in our dataset, and limited resolution at this level prevents reliable discrimination among them. Retaining multiple individuals classified only at this broad level would therefore risk artificially reducing diversity.

For each site, we calculated mitochondrial and Y-chromosome haplotype diversity ($h$) following Cassidy et al. (2025)[20], retaining only one individual per cluster of first-degree relatives. Also following Cassidy et al. (2025)[20], we assessed the extent to which burial communities were structured by biological relatedness by estimating, for each site, the proportion of genetically related pairs relative to the total number of possible pairs. Pairs were defined as related if they shared >24 cM in IBD across more than two segments.

During the Neolithic and Bronze Age, mitochondrial diversity remained close to 1 in all sites (Supplementary Table 3) (Figure 2c), including those with a high fraction of related individuals. No significant correlation was observed between mitochondrial diversity and fraction of relatives (Pearson $r$ = -0.43, $P$ = 0.21 for the Neolithic period). In contrast, during the Iron Age (comprising exclusively British sites), most sites exhibiting kinship relations (>0.04) show reduced mitochondrial diversity, with a significant correlation between the two variables (r = −0.82, P = $2.2 \times 10^{-08}$). Among sites with more than 20 individuals, Winterborne Kingston has an estimate of 0.94, while the three Arras cemeteries analysed in our study display similar values ranging from 0.86-0.89. The opposite pattern is observed for the Y-chromosome. Y-chromosome diversity is low (<0.76) in all Neolithic sites—particularly those with a high proportion of related individuals (r = −0.79, P = 0.034), such as Hazleton North and Carrowkeel—but, with one exception, exceeds 0.97 in all Bronze and Iron Age sites (Iron

Age: r = −0.28, P = 0.39) (Supplementary Table 3) (Figure 2c). Together, these results are consistent with a shift in the structuring principles of burial communities in Britain over time, from groups predominantly structured around male-line genetic relatedness during the Neolithic to groups predominantly structured around female-line genetic relatedness during the Iron Age.

As noted above, our intra-site Y-chromosome diversity estimates are conservative, since males assigned to the same branch at our level of resolution may in fact belong to distinct downstream lineages. However, this limitation is unlikely to drive the broad patterns observed. First, in Bronze and Iron Age sites, high Y-chromosome diversity is readily detected when present. Second, if the low Y-chromosome diversity were an artefact of data analysis, we would expect it to similarly affect all sites, regardless of the extent to which they are driven by biological relatedness. Instead, we observe the lowest diversities precisely in the sites with the highest proportion of relatives. Finally, at Hazleton North—the Neolithic site with the largest sample size—pedigree reconstruction shows that most males are close patrilineal relatives descending from a single founder male[22], indicating that the low Y-chromosome diversity in this case is biological rather than an artefact of limited resolution.

#### SI 5. Kinship Analysis and pedigree reconstruction

Across this and other sections, the terms 'relationship'/'relative' are being used as short-hand for 'biological/genetic relationships', with no implicit assumption of how this materialises as socially constructed kin.

We computed pairwise allelic mismatch rates[25–27] in the autosomes and X-chromosome for all pairs of individuals across Arras sites, randomly sampling one DNA sequence at each '1240k' polymorphic position (Supplementary Table 4). The mismatch rate values were converted into relatedness coefficients (*r*) following the same procedure as in Fowler et al. 2021[22]. For normalization, we used the mismatch rate value expected for two unrelated individuals from the British Iron Age population, estimated in the autosomes as the median value of the 10,892 Wetwang Slack-Melton 1 pairs with more than 100,000 overlapping SNPs, and in the X-chromosome as the median value of the 10,892 Wetwang Slack-Melton 1 pairs with more than 5,000 overlapping SNPs. This assumes a lack of relatives across the two sites—which is essentially true based on IBD results—with very few exceptions that do not affect the median mismatch rate value.

For the 515 individuals with more than 600,000 1240k SNPs with overlapping data, we called IBD segments in the autosomes and in the X-chromosome with ancIBD software v0.7 (https://ancibd.readthedocs.io/en/latest/index.html) after imputing and phasing genotypes using GLIMPSE, as described in Ringbauer et al.[28]

Relatedness coefficients and IBD statistics for each pair of individuals are included in Supplementary Table 4 and visualized in Figure S11 and Figure S12. Both analyses demonstrate the presence of several groups of closely related individuals within the three main sites: Wetwang Slack, Melton 1 and Pocklington.

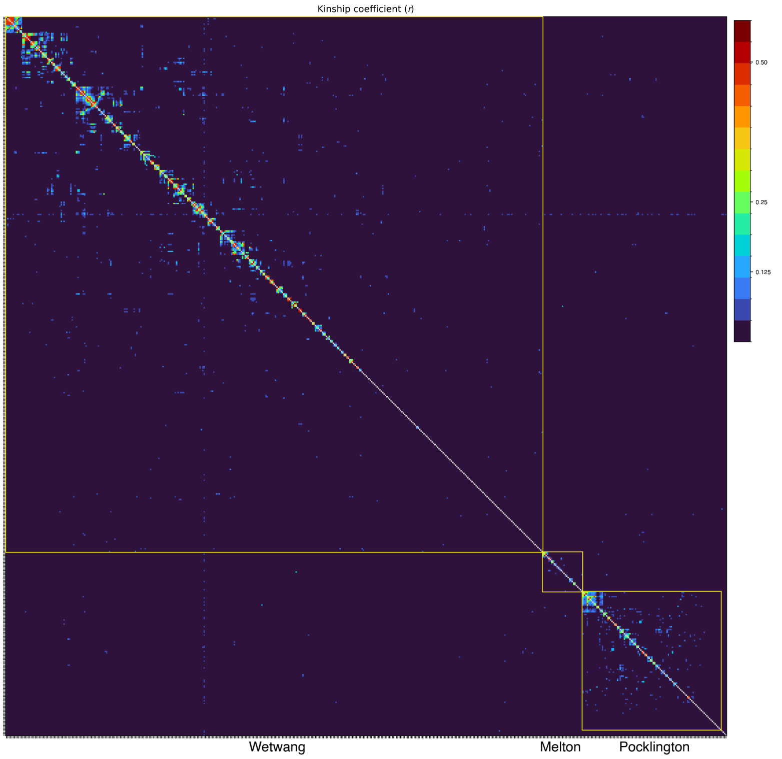


Figure S11. Heatmap of the autosomal relatedness coefficient values between pairs of individuals.

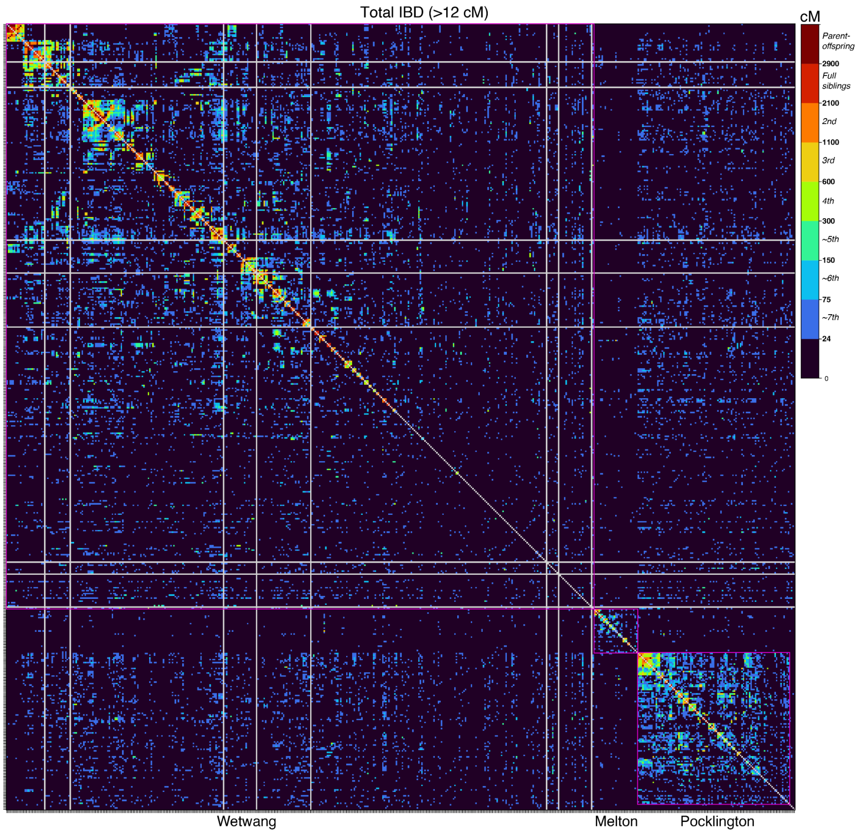


Figure S12. Heatmap of IBD sharing between pairs of individuals across the whole dataset. We plot the total length of the genome (in cM) in IBD segments >12 cM. Pairs including one individual without IBD calls are displayed in grey.

#### Pedigree reconstruction

We manually reconstructed pedigrees using the following pieces of information:

-Autosomal pairwise relatedness coefficients: inform the degree of genetic relationship up to the third–fourth degree.

-Total length of the genome in IBD segments longer than 12 cM: informs more distant degrees of genetic relationship.

-Number of IBD segments longer than 12 cM; this value is helpful for the differentiation of relationships sharing the same total amount of DNA. For instance, grandparent-grandchild relations and avuncular relationships are second-degree relationships sharing 25% of their DNA (~1600 cM), but the former tend to share fewer long IBD segments than the latter. The same goes for paternal half-siblings, who share a lower number of long segments than maternal half-siblings, even though the total amount of DNA shared is the same (25%; second-degree relatives).

-Position of IBD breakpoints: these can help differentiate genetic relationships between individuals sharing the same total amount of DNA. For instance, if a mature male is the grandfather of two young males who are themselves half-brothers, IBD breakpoints between the grandfather and one of the half-brothers should not be correlated to the IBD breakpoints between the grandfather and the other half-brother, as they are the result of two independent meiosis in the half-siblings' father[22]. If, instead of the grandfather, the mature man is their half-brother, we expect a fraction of the breakpoints (those happening in the meiosis leading to the mature man) to appear in both comparisons.

-Presence of regions of the genome where both the maternal and paternal genome share IBD: these regions are referred to as IBD2 and are characteristic of full-siblings but can also appear in other situations when two individuals are related through both their maternal and paternal sides.

-X-chromosome IBD patterns: these can inform, in some cases, on whether a specific genetic relationship is through the maternal or paternal side, and can also help clarify the exact type of relationship. For instance, a female and her paternal grandmother must share IBD along the entire X-chromosome, while this need not necessarily be the case for a woman and her paternal aunts.

-Mitochondrial and Y-chromosome haplogroups: transmitted through strictly maternal and paternal lines, respectively.

-Runs of homozygosity (ROH): indicative of parental genetic relatedness.

-Genetic sex.

-Age-at-death estimation determined by osteological analysis: in a first-degree parent-offspring relationship, for example, it would be impossible for pre-pubescent individuals to have reproduced and so they must represent offspring rather than parent.

***1. Wetwang Slack***

At Wetwang Slack, we identify a large group (195 individuals) connected by third-degree or closer genetic relationships. If we relax the threshold to include fifth-degree relatives (e.g. second cousins), the number of connected individuals increases to 288 (74% of the total number of sampled Wetwang Slack individuals). To facilitate the pedigree reconstruction, we split this large group of 195 into six smaller clusters of closely related individuals (plus a chariot burial cluster) and tackle the reconstruction of each cluster separately (Figure S13). Then, we attempted to connect these different clusters to one another.

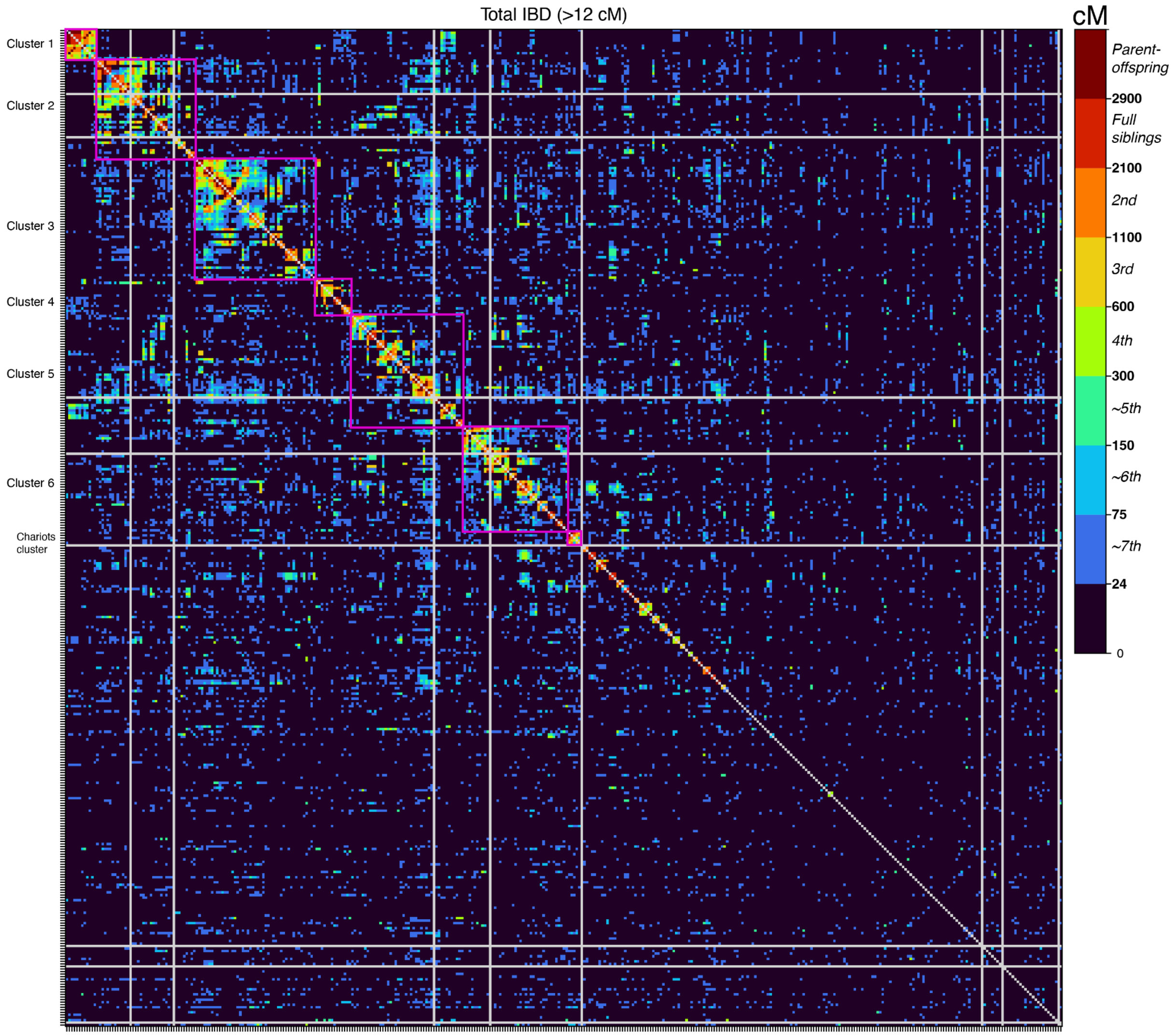


Figure S13. Heatmap of IBD sharing between pairs of individuals at Wetwang Slack and Wetwang Village. Clusters for pedigree reconstruction are indicated with magenta squares. We plot the total length of the genome (in cM) in IBD segments >12 cM.

##### 1.1. *Cluster 1*

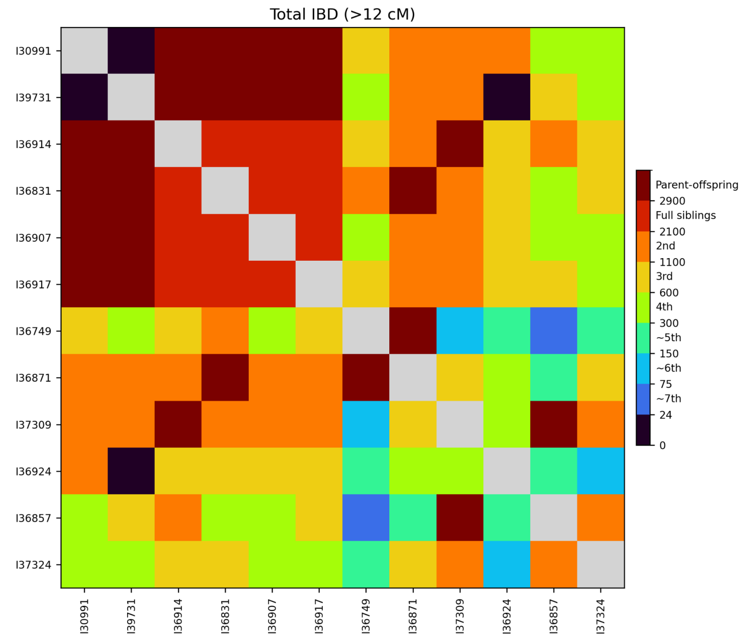


Figure S14. Heatmap of IBD sharing between pairs of individuals from cluster 1 at Wetwang Slack.

Cluster 1 comprises 12 individuals (Figure S14), detailed below.

1) Relationship between I30991 and I36924:
   They are second-degree relatives sharing 37 IBD segments. They could be maternal half-siblings, aunt-nephew or uncle-niece. We display them as aunt-nephew in Figure S15 but with red lines indicating uncertainty.

2) Relationship between I37324 and the rest of the pedigree:
   I37324 has a second-degree relationship with I37309 and I36857, and a third-degree relationship with I36914. Thus, I37324 must be I37309's grandson and I37309's nephew through I37309's unsampled son. However, several pieces of evidence suggest that I37324 is also related to the family through their mother's side:
   -I37324 has 38 cM in ROH segments >4 cM, therefore his parents must be related, most likely somewhere in the order of second-third cousins. Looking at the location of the ROHs in I37324 and the location of IBD segments between I30991-I37324 and I39731-I37324, some ROHs co-locate with I39731-I37324 IBD segments and others with I30991-I37324 IBD segments. This suggests that both I39731 and I30991 are I37324's genetic ancestors through both his paternal and maternal sides. These ROH segments also co-locate with IBD between I36914-I37324 and I37309-I37324, which means that these individuals must be part of the inbred loop.
   -I37324 shares IBD in the X-chromosome with I39731 and I30991, which would not be expected if I39731 and I30991 were I37324's great-great-grandparents through only I37324's paternal side.
   -The relatedness coefficient between I37324-I30991 is 0.12, higher than expected for fourth-degree relatives but the expected value for double fourth-degree relatives. The coefficient between I37324-I39731 is 0.08, also slightly higher than the expected value (0.0625) for simple fourth-degree relatives.
   
   Based on this evidence, we set I37324 as a descendant of I39731-I30991 through female-male-female steps. The genetic transmission cannot have occurred through three consecutive females because, if this were the case, I37324-I39731 would share

the same mtDNA lineage. Neither can the transmission have taken place through a male-female-male line of descent because, if this were the case, I37324-I39731 and I37324-I30991 would not share IBD in the X-chromosome, as they do. Under this scenario, I37324's parents would be second cousins.

I37324 is I37135's third-degree relative, and a fourth-degree relative of I37135's brother I31010 (cluster 2), which means that I37324 must be I37135's descendant. I37135 is not closely related to I37324's close paternal relatives, which means that they must be related through I37324's mother. Thus, I37135 must be I37324's maternal grandfather, through I37135's unsampled son or daughter. Given that I37135 and I37324 do not share IBD in the X-chromosome, a genetic relationship through I37135's unsampled son is more likely.

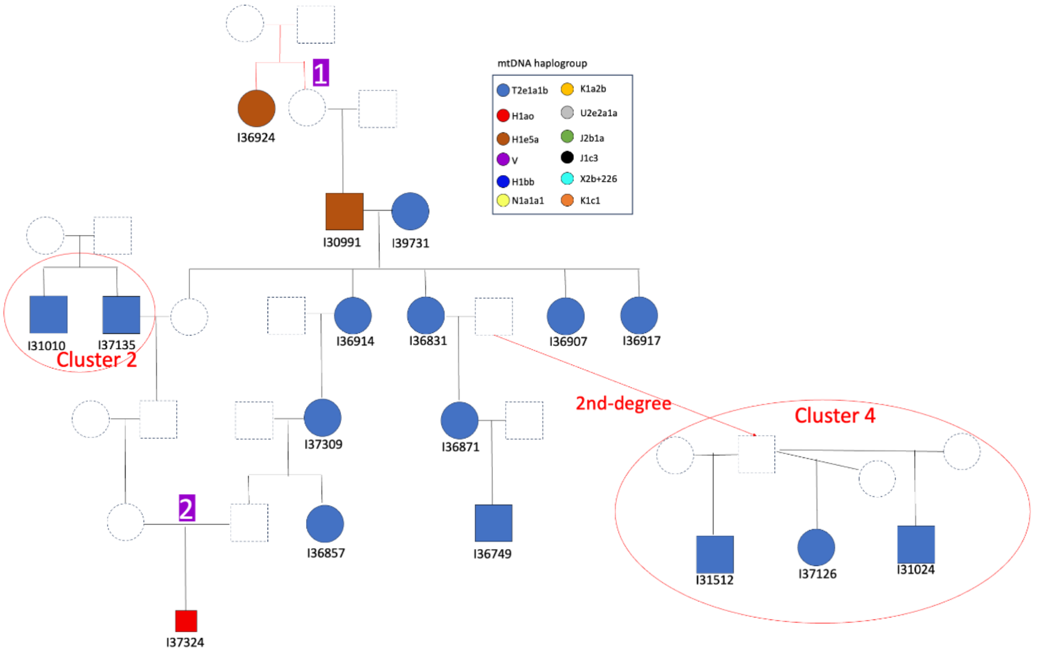


Figure S15. Reconstructed pedigree for cluster 1 at Wetwang Slack. In this and other clusters, we highlight with numbers parts of the tree that warrant some discussion. Smaller symbols represent individuals who died before reproductive age. Here, and throughout the text and figures, males and females are represented with squares and circles, respectively. Red lines indicate uncertainty in the tree topology.

##### *1.2. Cluster 2*

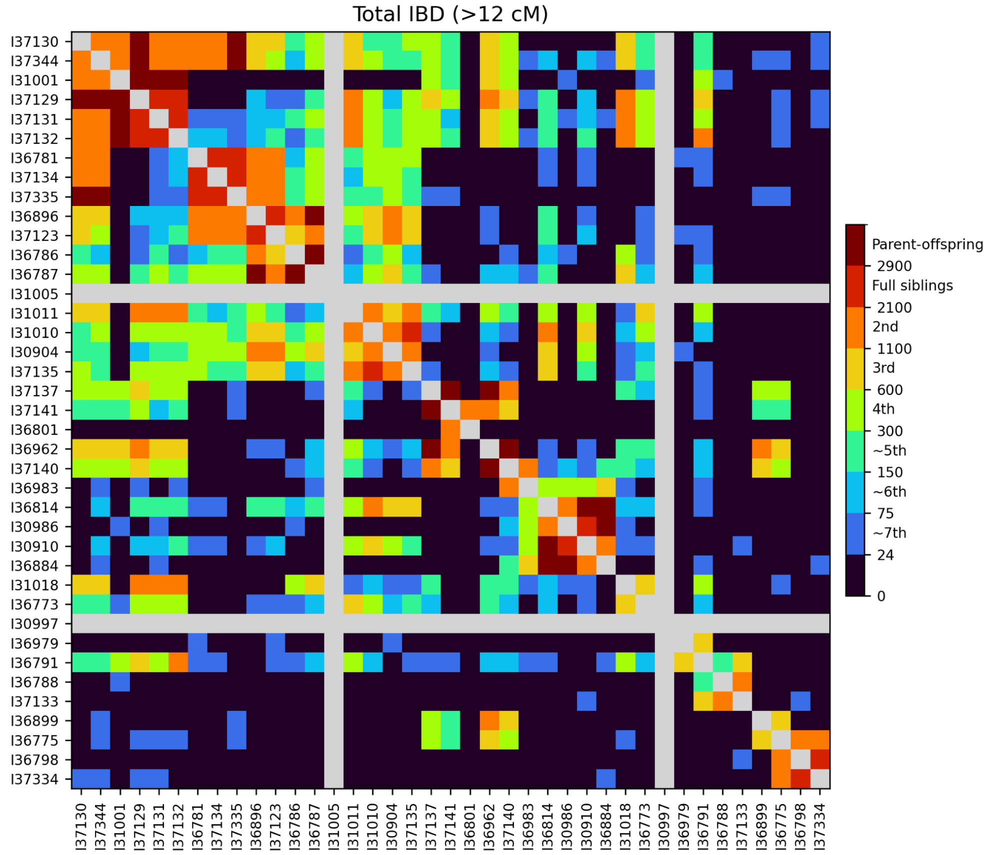


Figure S16. Heatmap of IBD sharing between pairs of individuals from cluster 2 at Wetwang Slack.

Cluster 2 comprises 39 individuals (Figure S16; Figure S17), detailed below.

1) Relationships between I31018-I31011 and I37129-I37131-I37132:
I31018 and I31011 are unrelated (except for one 60 cM IBD segment), but both are second-degree relatives of siblings I37129, I37131 and I37132 through the siblings' father; the siblings' mother (I31001) is unrelated to I31018 and I31011. Thus, I31018 and I31011 must be I37129-I37131-I37132's paternal grandparents, which is also confirmed by IBD patterns along the genome.

2) Relationship between I36962, I37137, I37140, I37141, I36801 and I36983:
I37141 is a second-degree relative of I36962 and I36801, but I36962 and I36801 are unrelated. This means that I36962 and I36801 are I37141's second-degree relatives though different sides (I36962 paternal and I36801 maternal) of the pedigree. I37137 must be I37141's father. I36801 is I37141's mother's first-degree relative, more likely her son or brother because the number of IBD segments between I36801-I37141 is high (n=34). I36962-I37140 are parent-offspring. I36983 is I37140's second-degree relative but is unrelated to I36962, so I36962 must be I37140's mother. I36983 is I37140's father's first-degree relative, more likely his mother or daughter because the number of IBD segments between I36983 and I37140 is low (n=15). I36983 cannot be I37140's granddaughter because otherwise she would be related to I36962 and she is not. As such:

-I36983 is I36884's third-degree relative.

-I36884 must be above I36983 based on the position of I36884's close relatives in the tree, specifically her daughter who is closely related to first generation I31010.

-I37140 and I36884 are ~fifth-degree relatives.

I36983 must be I37140's paternal grandmother, because if she were his paternal half-daughter, I36884 would either be equally related to them (if related through their father) or unrelated to I37140 (if related through I36983's mother).

I36962 is I37137's mother or daughter and the available evidence strongly points to I37137 being I36962's daughter and I37140's maternal half-sister because:

-I37137 and I36962 share the same mtDNA, and a father does not usually share the mtDNA with his daughter.

-In this scenario, I37137 is I37140's maternal half-sister which explains the high number of IBD segments between them (n=38). If I37137 were the father of I36962, he would be I37140's grandfather and we would expect a low number of IBD segments between them.

-I36962's relatives appear equally related to both I37137 and I37140, which suggests that they were in the same generation, rather than being separated by two generations.

3) Relationship between I37129 and I36962-I37137-I37140-I37141-I36801:
I37129 is a second-degree relative of I36962. I36962 is not, however, related to I37335, which means that she cannot be a descendant of I37129-I37335. She cannot be I37129's grandmother or aunt because she would then be a first-degree relative of I37129's mother or first-degree relative of I37129's paternal grandparents, and she is not. She cannot be I37129's paternal half-sister because then she would be a second-degree relative of I37131-I37132-I31018-I31011, and she is not. She cannot be I37129's niece because then she would be a second-degree relative of I37131-I37132, and she is only a third-degree relative of them. Thus, I36962 can only be the granddaughter of I37129 through a maternal half-brother of I37344-I37130. The X-chromosome IBD patterns and the number of IBD autosomal segments (n=18) between I36962 and I37344-I37130 support this configuration.

4) Relationship between I36899 and I36962:
I36899 is not closely related to I37129, which means that their genetic relationship is through I36962's mother, either through I36962's mother's son (sired by a different father) or I36962's mother's brother, given the high number of IBD segments shared (n=39). It is therefore likely, given their equal relationship to their cluster 5 relatives, that I36899 and I36962 are maternal half-sisters.

5) Relationship between I31018 and I30997:
I31018 and I30997 are second-degree relatives, but this cannot be through a descendant of I31018 and I31011, because I30997-I31011 are not themselves closely related.

6) Relationship between I36773 and I31018-I31011:
I36773 is third-degree relative of both I31018 and I31011, and must therefore be their descendant. This relationship cannot be through a sibling of I37129-I37131-I37132 because, if this were the case, I36773 would be their second-degree relative and she is not. Neither can the relationship be through a paternal half-sibling of I37129-I37131-I37132, because, if this were the case, she would be their third-degree (and she is

most likely a fourth-degree relative). I36773 must therefore be a descendant through I37129-I37131-I37132's paternal uncle/aunt. Given that I36773 shares IBD in the X-chromosome with I31018, the genetic relationship must have been conveyed through I37129-I37131-I37132's paternal aunt and her offspring. We display the offspring as a daughter in Figure S17 because the mtDNA haplogroup of I36773 and I31011 is the same.

7) Relationship between I36791 and I37132:
I36791 is I37132's second-degree relative and I37131-I37129's third-degree relative. Thus, he must be I37132's grandson through I37132's daughter because they share IBD in the X-chromosome and the mtDNA lineage.

8) Relationship between I36791, I36979, I37133 and I36788:
I36979 is unrelated to I37133 and I36788 (who are themselves second-degree relatives). I36791 is a third-degree relative of I36979 and I37133, and a fourth/fifth-degree relative of I36788. I36979-I37133-I36788 are unrelated to I37132 (I36791's maternal grandmother), which means they cannot be descendants of I36791. They must, therefore, be related to I36791's father and the others through I36791's maternal grandfather. I37133-I36788 share long IBD segments in the X-chromosome with I36791, which means that they must be related to I36791 via I36791's maternal grandfather. As such, I36979 must be related to I36791 via I36791's father, but not through a strict patrilineal connection, as they have different Y-chromosome lineages. Individual I37133 must, therefore, be I36791's maternal grandfather's first-degree relative, either their mother, daughter or sister. The number of shared IBD segments between I37133-I36791 (n=17) is, however, rather low for a third-degree relationship, suggesting that there is no sibling pair involved in the genetic transmission between I37133 and I36791. Furthermore, if I37133 was I36791's maternal grandfather's daughter or mother, I36791's maternal grandfather's entire X-chromosome would be present in I37133, and the X-chromosome segments not shared between I36791-I37132 would therefore be shared between I36791-I37133, as I36791 must have inherited his X-chromosome from his maternal grandparents. This is precisely what we observe, and, as such, this favours a scenario whereby I37133 is the daughter or mother of I36791's maternal grandfather. If I37133 was I36791's maternal grandfather's sister, this pattern would not necessarily hold, as I37133 and his brother could have inherited different X-chromosome segments from their common mother. We thus display I37133 as a daughter of I36791's maternal grandfather from a different reproductive union, but with red lines to indicate uncertainty.

9) Relationship between I36775, I36798, I37334,I36899 and I36962:
I36775 is I36962-I36899's third-degree relative, I37334's second-degree relative and I36798's second/third-degree relative. I36798-I37334 are unrelated to I36899-I36962, which means that I36798-I37334 must be related to I36775 through I36775's maternal line (they share the same mtDNA lineage), while I36899-I36962 are related to I36775 through I36775's paternal side. As such, I36775's father must be I36962-I36899's mother's first-degree relative, either her son (from a third reproductive union), brother or father (with I36775 being I36962-I36899's mother's paternal half-sister). The three scenarios align well with the observed X-chromosome IBD sharing between I36775 and I36962-I36899, but one piece of evidence suggests that I36775 is more likely I36962-I36899's mother's paternal half-sister. Individual I37138 from cluster 5 is a

third-degree relative of both I36962-I36899, and so I37138 has to be related through their shared mother. However, I37138 is unrelated to I36775. Thus, I36775's father cannot be I36962-I36899's mother's son or brother, because if this was the case, I36775 should be related to I37138 like I36962-I36899's unsampled mother is. If I36775 is I36962-I36899's mother's paternal half-sister, she does not need to be related to I37138, because the genetic relationship between I37138 and I36962-I36899 comes from their maternal grandmother, not shared with I36775. In this scenario, if I37138 is related to I36962-I36899 via their maternal grandmother, he must be their maternal grandmother's father because he has a different mtDNA lineage.

10) Relationship between I31010, I30986, I36884, I30910 and I36814:
I36814 is I31010's second-degree relative and I37135's third-degree relative. Given that I31010-I37135 are brothers, I36814 must be I31010's granddaughter through a son of I31010 due to the lack of X-chromosome IBD sharing between I31010 and I36814, but also because I36814's maternal side is already accounted for (see below). I36814 is a first-degree relative of both I36884 and I30910 (in both cases this is an offspring-parent relationship), who are themselves second-degree relatives. I36884 is unrelated to I31010 (I36814's second-degree relative) and must, therefore, be I36814's mother. I30910 is I31010's third-degree relative and thus can only be I36814's daughter. I30986 is I36884's mother or daughter and I36814's second-degree relative. In a regular scenario, we would expect I30986 to be I30910's third-degree relative (either I30910's great grandmother or half-aunt). However, I30986-I30910 have an r-coefficient of 0.38 (sharing 2181 cM in IBD and 34 IBD segments) and share 448 cM in IBD2, but none of the four individuals have any ROH segments. The r-coefficient of 0.38 suggests that in addition to the predicted third-degree relationship (r~0.125) between I30986 and I30910 predicted based on their relationship to I36884, they share a second-degree relationship (~0.25) through their father's line. This leaves us with two possible scenarios, depending on the relationship between I30986 and I36884:
  - -If I30986 is I36884's mother, then I30986 could be I30910's paternal aunt or paternal grandmother (or their great-grandmother through I30910's mother's side), either through I36884's unsampled brother or maternal half-brother. In these scenarios, I30910's parents would be either second-degree relatives (uncle-niece) or third-degree relatives (granduncle-grandniece or half uncle-half niece). In any of these scenarios, I30910 would display high levels of endogamy, but she does not have any ROH, which renders them impossible.
  - -If I30986 is I36884's daughter (and I36814's half-sister), then I30986 could be I30910's paternal half-sister (as well as their half-aunt through I30910's mother's side). In this scenario, none of the four individuals would be expected to show ROH and I30986-I30910 would be expected to share segments in IBD2, because they could easily inherit an identical DNA segment from their common father, and an identical segment through their mother's side. We conclude that this is, indeed, the only possible scenario: a man reproducing with his daughter's maternal half-sister. In such a case, the second-degree relation would share ~1600 cM of DNA (50% of the genome) and the third-degree relation would share ~800 cM (25% of the genome). As such, we would expect both sets of IBD to overlap in ~12.5% of the genome (i.e. ~400 cM), which falls very close to the value of IBD2 we observe (Supplementary Table 4).

So far, we have constructed the tree scaffold below individual I31005, including siblings I36781-I37134-I37335. We now need to determine the tree structure of I30904 and his close relatives (Figure S18), which connects with the existing scaffold in two places: in siblings I36781-I37134-I37335 and through I31011's mother. Based on the presence of regions (~200 cM) of the genome that display IBD0 between pairs of siblings and IBD1 between each sibling and their second-degree relative, we can determine that I36896 and I37123 must be I36781-I37134-I37335's uncle and aunt, and that I30904 must be I36896's and I37123's maternal uncle. Based on the absence of such IBD regions, I30904 and I31011 cannot be uncle/aunt of I37135-I31010, and must be either the nephew and niece of I37135 and I31010, or a maternal half-sibling (one of them) and a nephew/niece (the other one) (Figure S18). The configuration shown in Figure S17 for I30904's close relatives (with I37135's and I31010 as maternal uncles of I30904 and I31011) is the most likely, because it allows for a link with the existing scaffold without the need to shift generations, i.e. siblings I36781-I37134-I37335 occur two generations below I31011, exactly like the scaffold. Under this configuration, I37335 and I37129 are third cousins and their offspring I37344-I37130 are expected to display some ROH segments. This is, indeed, true for I37130, who presents ROH compatible with his parents being second/third cousins.

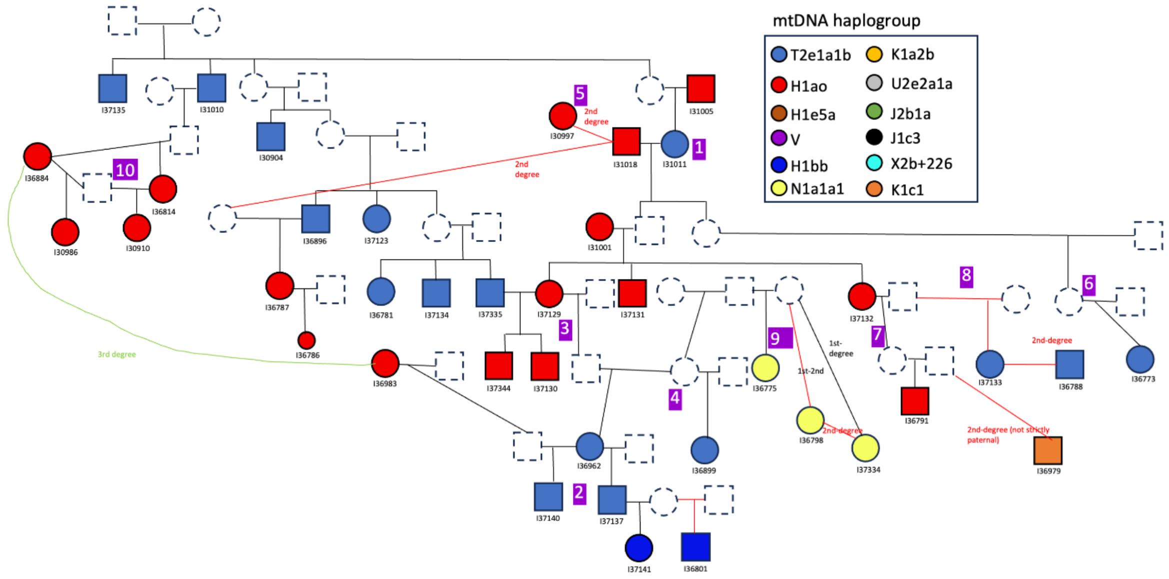


Figure S17. Reconstructed pedigree for cluster 2 at Wetwang Slack.

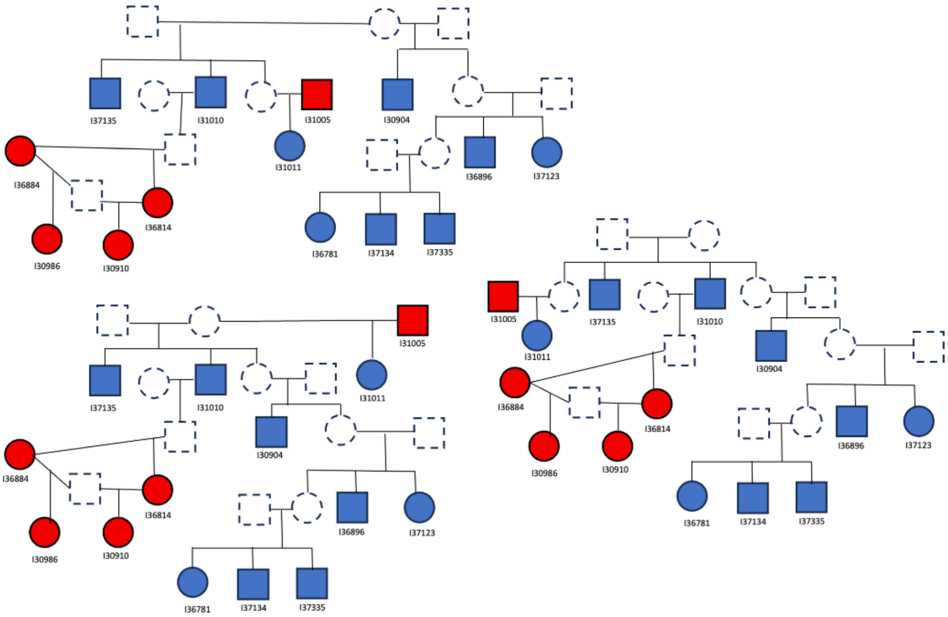


Figure S18. Possible configurations of I30904 and his closest relatives at Wetwang Slack.

##### *1.3. Cluster 3*

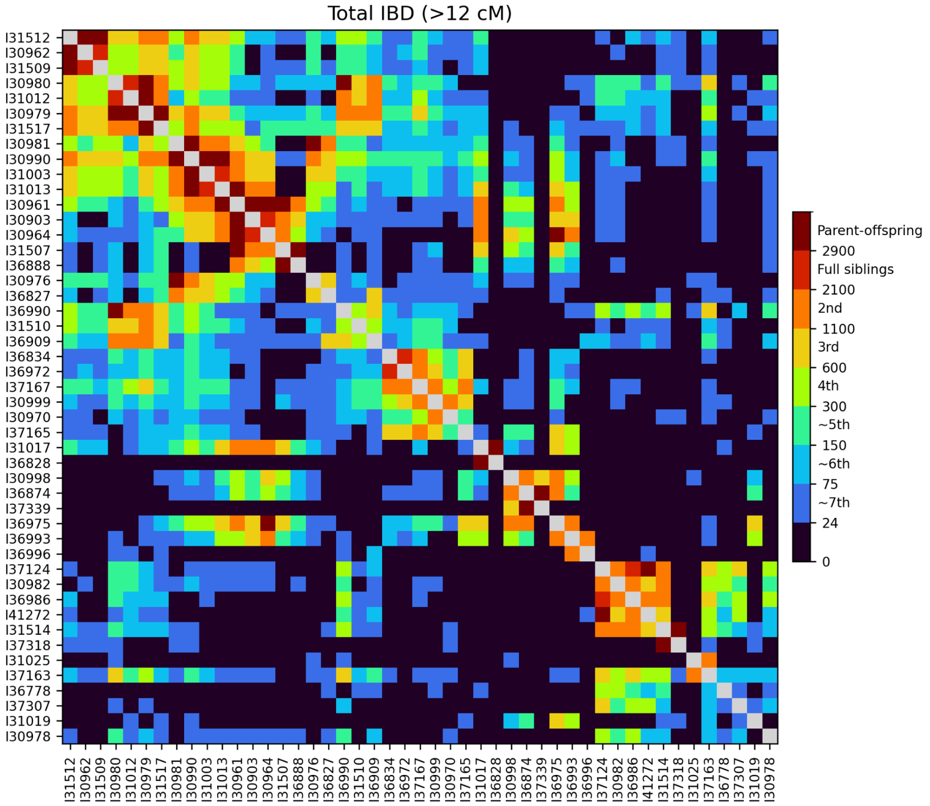


Figure S19. Heatmap of IBD sharing between pairs of individuals from cluster 3 at Wetwang Slack.

Cluster 3 comprises 47 individuals (Figure S19; Figure S20), detailed below.

1) Relationship between I31517, I30979 and I30990:

I31517 and I30979 are a mother-daughter pair (order unknown at present), and are both second-degree relatives of I30990. Thus, I30990 must be their granddaughter and niece. Given that the number of shared IBD segments between I30990-I31517 and I30990-I30979 is 26 and 40, respectively, I31517 must be I30990's maternal grandmother and I30979 must be I30990's maternal aunt.

2) Position of I37167 in the pedigree:
If I37167 is I31517's great granddaughter and I30979's grandniece, we would expect both to be equally related to I37167, but I37167-I31517 share 290 cM in IBD and I37167-I30979 share 693 cM. This suggests that I37167 is I30979's descendant, hence one degree closer to I30979 than to I31517, but in terms of distribution of IBD segments along the genome, some segments shared in I37167-I31517 are not present in I37167-I30979, which excludes the possibility that I37167 is a descendant of I30979. Furthermore, if I37167 was a descendant of I30979 (possibly her great granddaughter via a third reproductive union), I37167 would be a fourth-degree relative of I30980 and I31012, but a fifth-degree relative of I30990. This does not seem to be the case, because the three individuals share a similar amount of IBD with I37167, consistent with a fourth-degree relationship. We therefore retain the first scenario in the reconstructed tree, with I37167 as I31517's great granddaughter and I30979's grandniece, and red lines indicating uncertainty because I37167-I31517's IBD sharing appears to be too low for third-degree relatives.

3) Position of I30970 in the pedigree:
I30970 is I30999's second-degree relative (sharing the mtDNA lineage and IBD in the X-chromosome), and must be I30999's grandson or nephew through I30999's sister or daughter. He cannot be a maternal half-sibling because then I30970 and I37167 would be second-degree relatives, and they are closely related (523 cM) but definitely not second. The number of IBD segments (31) slightly favors I30999 being I30970's maternal aunt, but all the IBD segments present in I30970-I37167, I30970-I37165, I30970-I36834 or I30970-I36972 are a subset of those in I30970-I30999, which means that I30999 must be I30970's maternal grandmother, because if she were his aunt, we would expect some IBD segments in I30970 and I37167 to be absent in I30970-I30999 because they were passed through I30999's sister.

4) Position of I36888 in the pedigree:
I36888 must be I31507's son or father. We cannot identify any clear case of a sampled individual that is related to I36888 but not to I31507 or vice versa, which we would need to easily resolve these positions. I36888 and I30961 (I31507's son) share only 13 IBD segments, which suggests a grandfather-grandson relationship or a paternal half-brother relationship, but grandfather-grandson is more likely because 13 shared segments seems too low for a paternal half-brother relationship. Furthermore, there are several individuals (such as I36918 and I36925's mother) that are one degree closer in their relationship to I36888 than to I31507. If I36888 were the son of I31507, those related individuals could only be I36888's descendants. One of them is I36925's mother from cluster 4, who is I36888's 2nd-degree relative. If I36888 were the son of I31507, I36925 would be I36888's granddaughter. This is very unlikely, because in the tree connecting clusters 3, 4 and 5, I36925's mother is in the same generation as I31507. Based on these pieces of evidence, we conclude that I36888 is much more likely to be the father of I31507 than vice versa.

5) Position of I36996 in the pedigree:
I36996 is I36993's second-degree relative but is unrelated to I36993's paternal relatives. She must, therefore, be I36993's maternal aunt, half-sister or grandmother. The number of shared IBD segments between these individuals is 34, which favours a maternal aunt or half-sister relationship over a grandmother relationship. We, therefore, display I36996 as I36993's maternal aunt in the pedigree but do so with a red line to indicate uncertainty.

6) Relationship between I30998 and I36874:
The relatedness coefficient between I30998 and I36874 is 0.33, which is too high for a simple second-degree relationship but aligns well with a combined second+fourth-degree (0.25+0.0625). In such a case, the second-degree relationship would sum ~1600 cM (50% of the genome) and the fourth-degree relationship ~400 cM (12.5% of the genome), with an expected overlap in ~6.2% of the genome, i.e. ~200 cM; this is close to the amount of IBD2 sharing we observe in these individuals (Supplementary Table 4). Furthermore, comparing I30998 with I36874's father (I37339) gives a third-degree relationship, confirming that the fathers of I30998 and I36874 are second-degree relatives.

7) Relationship between I37165 and I36975:
These two individuals have a ninth-degree relationship in the pedigree through I36975's mother, but IBD sharing is compatible with a third-degree relationship. Furthermore, I36993 (I36975's nephew) and I37165 have a fourth-degree relationship. Thus, I36975's father must be I37165's second-degree relative, possibly I37165's grandson or nephew given that I37165 is one generation above I36975's father in the tree. In a scenario where I36975 is the nephew of I37165, I36975's parents would be 4th cousins.

8) Relationship between I36909 and I36827:
I36909 and I36827 are third-degree relatives, and this must be through I36827's father, since I36827's maternal relatives are more distantly related to I36909 than I36827 is to I36909. I36827's father must, therefore, be I36909's second-degree relative, possibly his nephew given that I36909 lies one generation above I36827's father in the tree.

9) Relationship between I37163 and I30980-I36990:
I37163 is a third/fourth-degree relative (sharing 600–700 cM in IBD) of both I30980 and her daughter, I36990. We detect IBD segments present in I37163-I30980 that are not present in I37163-I36990 (and vice versa) which, together with the fact that I37163 displays a similar degree of biological relatedness with I30980-I36990, indicates that I37163 must be a descendant of I36990's unsampled sibling. I37163 shares substantial IBD in the X-chromosome with both I30980 and I36990, but I37163's paternal second-degree relative is not closely related to I30980-I36990, which means that their genetic relationship must be through I37163's mother. Furthermore, X-chromosome IBD segments between I37163 and I36990 do not entirely overlap with those between I37163 and I30980, indicating that the unsampled sibling is a female; if he were a male, the shared segments between I36990 and this unsampled individual (and by extension I37163) would all derive from their mother, I30980. I37163 is a third-degree relative of both I36986 and I37124, and I36986 and I37124 are one degree

more distant to I30980-I36990 than is I37163. Thus, I37163 must be the maternal half-brother of I36986-I37124's mother.

10) Relationship between I30982, I31514, I36986 and I37124:
I30982 and I31514 are I36986-I37124's second-degree relatives. Neither of them can be their grandmother because they died as infants. I31514 cannot be a maternal aunt because her father is not related to the other individuals, and the same is also very unlikely for I30982 because we would expect ~200 cM of the genome where each of the sisters I36986-I37124 share IBD1 with I30982, and at the same time are IBD0 between the sisters, and we only detect 10 cM of such segments. I30982 and I31514 could, however, be the sister's maternal half-sisters or nieces. Since I37163 is one degree closer to I36986-I37124 than to I30982 and I31514, I30982 and I31514 are much more likely to be one generation below sisters I36986 and I37124, as their nieces.

11) Relationship between I30978 (the individual with Down syndrome) and sisters I36986-I37124:
I30978 is a fourth-degree relative of sisters I36986-I37124, and a more distant relative of the sisters' relatives, including I30980 and I36990. I30978 could be a descendant of an unsampled sibling of I36986-I37124 or a relative of I36986-I37124's mother. If a descendant of I36986-I37124's unsampled sibling, I30978 would be seventh-degree relative of I30980 and I36990 (like I37307), but their IBD sharing suggests a closer relationship. This can be accommodated by setting I30978 as a great-grandson of I36986-I37124's mother and a different man than I36986-I37124's father. In this configuration, I30978 would be I30980-I36990's sixth-degree relative, I36986-I37124's fourth-degree relative, and I37163's fifth-degree relative, which is more consistent with the pairwise IBD sharing observed. However, we display I30978's position in the tree with red lines to reflect a degree of uncertainty.

12) Relationship between I50780 (female) from the chariot burial at Wetwang Village with members of cluster 3 at Wetwang Slack.
Female I50780 belongs to mitochondrial haplogroup H1ao, the most frequent maternal lineage at Wetwang Slack after T2e1a1b. Her closest relationship is a fourth-degree relationship with an infant female (I36778) at Wetwang Slack, who is a maternal relative of I30982-I36986-I37124-I31514. I50780's second closest relationship is a fourth-degree relationship with I30999, who is part of cluster 3 and is closely linked to cluster 6 via her reproductive partner. The relationship between I50780 and I30999 is either through one of I30999's descendants (other than I30970's mother) or sibling. The genetic transmission cannot have taken place through I30999's father or mother because, while I30999's maternal relatives are also related to I50780, this relationship is more distant than I50780's relationship with I30999. Neither can it be through I30970's mother because, in this scenario, I50780 would be equally related to I30970 and I30999, and she is not. Interestingly, the three individuals from the chariot burials at Wetwang Slack (I36995, I36892, I36978) are among 10 individuals displaying strongest IBD sharing with Wetwang Village individual I50780, representing ~fifth–sixth-degree relatives.

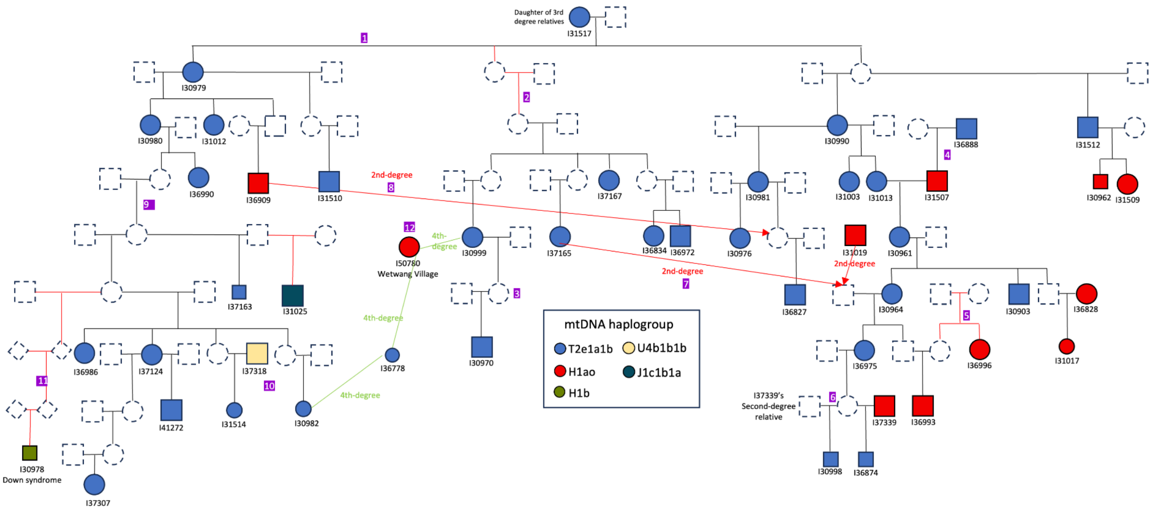


Figure S20. Reconstructed pedigree for cluster 3 at Wetwang Slack.

##### *1.4. Cluster 4*

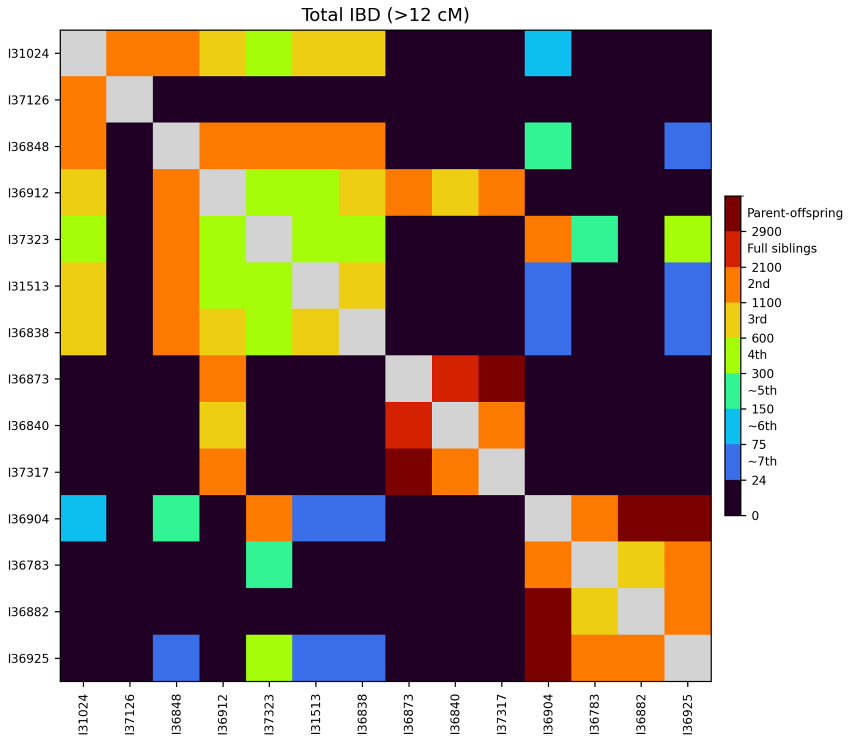


Figure S21. Heatmap of IBD sharing between pairs of individuals from cluster 4 at Wetwang Slack.

Cluster 4 comprises 14 individuals (Figure S21) (Figure S22), detailed below.

1) Relationship between I31512, I31024 and I37126:
   I31512, I31024 and I37126 are all second-degree relatives, which indicates that there must be at least one half-sibling relationship between them:
   - -One possibility is that these three individuals are paternal half-siblings from three different mothers.
   - -Another possibility is that two of them are half-siblings and the other individual is their paternal grandfather/grandmother. However, this is very unlikely because if I31512 was the grandfather of I31024 and I37126, IBD breakpoint locations in

I31512-I37126 and I31512-I31024 should not correlate, but they do. This, of course, also argues against a scenario in which one of the other two individuals (I31024 or I37126) is grandfather/grandmother to the others. Indeed, each IBD breakpoint observed in the comparisons between these three individuals appears in two of the comparisons (those involving the individual where that particular recombination event occurred) but not in the third, which is the expected pattern in the scenario with three half-siblings. For instance, recombination events in I31512's gamete appear as IBD breakpoints in I31512-I37126 and I31512-I31024 but not in I37126-I31024. This evidence suggests the presence of three half-siblings

-Another possibility is that two of the individuals are paternal half-siblings and the third is their paternal uncle/aunt. This is unlikely, however, because if that is the case, the uncle/aunt would share a high number of IBD segments with the other two, which they do not: I31512-I37126 (n=23), I31512-I31024 (n=25) and I31024-I37126 (n=30). These figures would allow for a nephew-aunt relationship between I31024-I37126, but do not support this relationship between the other two individuals.

We thus conclude that the three individuals are paternal half-siblings. The only minor misalignment in this scenario is that I31509 (I31512's daughter) and I37126 should be third-degree relatives, but they share 540 cM of IBD and have a relatedness coefficient of 0.10, both of which lie on the limit between third- and fourth-degree relatives.

2) Relationship between I31024, I31513, I36838, I36912 and I36848:
I36848 is a second-degree relative of I31024 (35 shared IBD segments), I31513 (22 shared IBD segments), I36838 (30 shared IBD segments) and I36912 (22 shared IBD segments). The other four individuals (I31024, I31513, I36838 and I36912) are all third-degree relatives of one another, except for I31513 and I36912, who share 454 cM (16 segments) of IBD, which is quite low for a third-degree relationship. However, there is no simple pedigree configuration that would satisfy these relationships if I31513 and I36912 are fourth-degree relatives. If, however, we assume that I31513 and I36912 are third-degree relatives, two simple configurations would be possible: either I36848 is a maternal aunt of I31024, I31513 and I36838 and paternal aunt of I36912, or I36848 is the paternal grandmother of I36912 and maternal grandmother of I31024, I31513 and I36838.

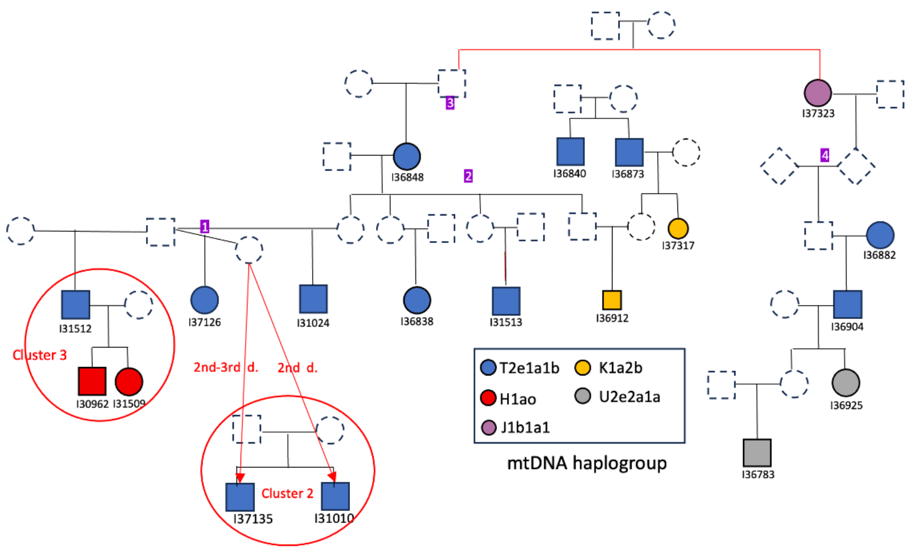


Figure S22. Reconstructed pedigree for cluster 4 at Wetwang Slack.

If I36848 is the aunt of I31024, I31513 and I36838, I37323 must be I36848‘s granddaughter, in order to satisfy I37323‘s fourth-degree relationships with I36848’s second-degree relatives. Individuals I37323 and I36848 cannot be I37323‘s paternal grandmother and granddaughter because I37323 and I36848 do not share IBD segments along the entire X-chromosome (Figure S23). Individual I36848 must, therefore, be the grandmother of I31024, I31513, I36838 and I36912.

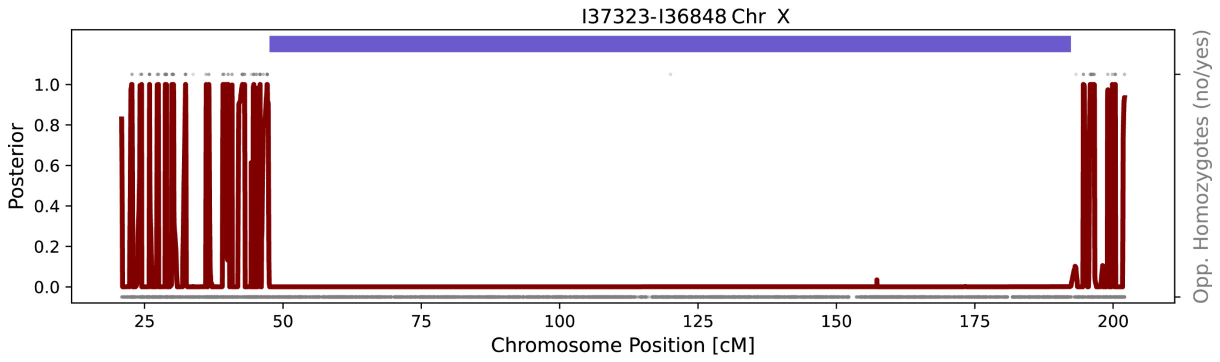


Figure S23. Posterior probability of non-IBD state on the X-chromosome for I37323 and I36848. Opposing homozygotes are shown as grey dots.

3) Relationship between I37323 and I36848:
Individual I37323 is either I36848’s paternal aunt or niece. I37323 and I36848 cannot be paternal half-sisters or granddaughter-paternal grandmother as they do not share IBD segments along the entire X-chromosome. Therefore, I37323 is displayed as the aunt, with red lines added to reflect a degree of uncertainty.

4) Relationship between I37323 and I36904:
The IBD sharing between I37323 and I36904 is halfway between what would be expected for second- and third-degree relatives, but given that I36904’s son (I36925) appears more likely to be a fourth-degree relative of I37323, we assume that I37323 and I36904 are third-degree relatives. Furthermore, we know that the genetic

relationship between I37323 and I36904 is transmitted through I36904's father, because I36904's mother is not related to them, and the relationship between I36904 and I36848's is ~fifth-degree, which means that I36904 must be I37323's direct descendant (i.e. I36904 cannot be related to I37323 via I37323's mother). It is possible that I36904 and I37323 are related through an unsampled sibling of I37323, but, if this were the case, I36904 would be a fourth-degree relative of I36848, and a fifth-degree relationship between them seems more likely. As such, we display I36904 as I37323's great grandson in the pedigree.

##### *1.5. Cluster 5*

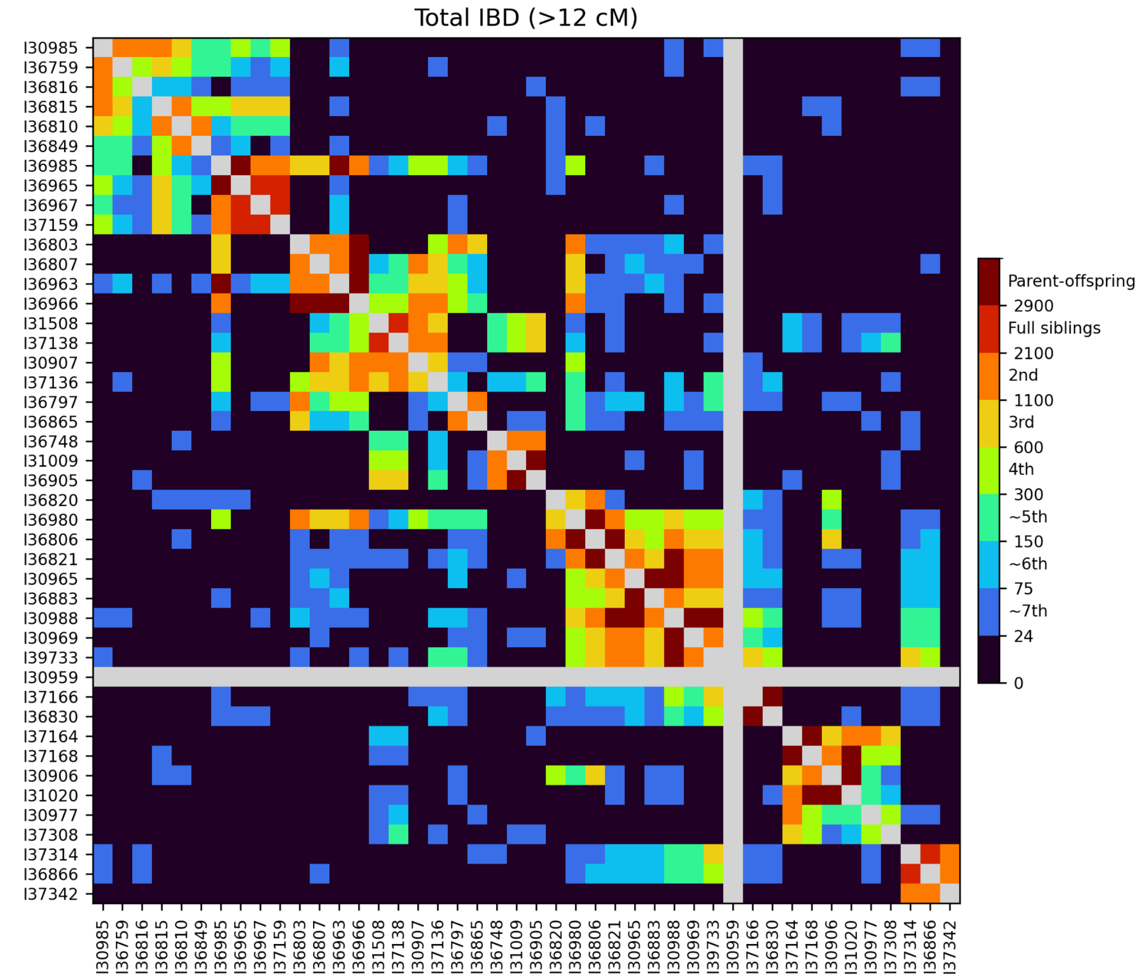


Figure S24. Heatmap of IBD sharing between pairs of individuals from cluster 5 at Wetwang Slack.

Cluster 5 comprises 44 individuals (Figure S24; Figure S28), detailed below.

1) Relationship between I37136 and I36966-I37138:
   I37136 is a second-degree relative of both I36966 and I37138. The location of shared IBD segments strongly supports the scenario whereby I36966 and I37138 (who are themselves related) are I37136's maternal grandparents.

2) Relationship between I36966, I30907, I37138 and I31508:
   Individual I30907 is a second-degree relative of I36966 and siblings I37138 and I31508. I36966 is also related to the siblings, sharing 481 cM and 577 cM in IBD with

them respectively. Siblings I37138 and I31508, themselves, display high levels of endogamy; based on the length of ROH segments, their parents must be ~third-degree relatives. The low number of IBD segments (n=20) shared between I36966 and I30907 suggests that avuncular and maternal half-sibling relationships (impossible also given the mtDNA lineages) are very unlikely. A paternal grandmother-granddaughter relationship or a paternal half-sister relationship for I36966-I30907 are plausible in principle. The evidence supports either scenario, as they share IBD segments across the whole X-chromosome. Since I36966's mother (I36803) is not related to I30907, I30907 must be the grandmother of I36966 if the paternal grandmother-granddaughter scenario is correct. Based on these two possible scenarios, together with the evidence that I36966 and I37138 had offspring together, we can exclude I37138-I31508 as I30907's maternal uncle-aunt because, if I36966 and I30907 were paternal half-sisters, I37138-I31508 would not be related to I36966, and if I36966 and I30907 were granddaughter-paternal grandmother, I37138 would need to have reproduced with a woman three generations below him in the pedigree, which is extremely unlikely. I30907 could, therefore, be the maternal grandmother, aunt or half-sister of I37138 and I31508.

Taking into account the number of shared IBD segments between I31508-I30907 and I37138-I30907, the location of shared IBD segments in I31508-I30907, I37138-I30907 and I36966-I30907, the relationship between I36966 and siblings I31508-I37138, and the X-chromosome patterns observed, the following possible scenarios are presented in Figure S25:

-Tree 1: this configuration is impossible for two reasons. All IBD segments between I31508-I36966 and I37138-I36966 are a subset of those present in I31508-I30907 and I37138-I30907, which means that genetic transmission from I36966 to siblings I37138-I31508 must take place via individual I30907, and this is not the case in this scenario. In this scenario, the predicted relationship between siblings I37138-I31508 and I36966 is at the third-degree+fifth-degree (due to I37138-I31508's parents being third-degree relatives). As such, we would expect c. ~1000 cM (800+200) of shared IBD between I37138-I31508 and I36966, which is double the amount we actually observe.

-Tree 2: this aligns well with all the evidence.

-Tree 3: this configuration is impossible because we would expect to find regions of the genome (~200 cM) in IBD0 between siblings I37138 and I31508 that also appear in IBD1 between both I30907-I37138 and I30907-I31508, but, in fact, only 4 cM was observed.

-Tree 4: this configuration is very unlikely for three reasons:

1) I37138 and I30907 share only 24 IBD segments, which seems too low for a maternal half-sibling relationship.

2) I37138 would need to have reproduced with a woman two generations below him.

3) IBD breakpoints between I37138-I30907 and I31508-I30907 are very poorly correlated. In this configuration, we would expect breakpoints derived from recombinations in the gamete leading to I30907 to be present in comparisons with both I37138-I30907 and I31508-I30907.

-Tree 5: this scenario is very unlikely because I37138 would need to have reproduced with a woman two generations above him.

-Tree 6: this configuration is impossible for the same reasons as outlined for trees 1 and 3.

-Tree 7: this scenario is impossible because it predicts no relationship between I37138-I31508 and I36966.

The only configuration which aligns with all the evidence is the one presented in tree 2 (Figure S25), in which I30907 is the paternal grandmother of I36966 and the maternal grandmother of I37138-I31508 via a different reproductive partner. Under this configuration, we can determine that I37138-I31508's father must be related to I37138-I31508's mother via her father, rather than via her mother (I30907). If the latter was the case, I30907 would be related to I37138-I31508 through a second-degree relation via the siblings' mother, and through a third-degree relation via the siblings' father. This scenario predicts ~1600+800 cM of IBD sharing between the siblings and I30907: far higher than the observed values which are compatible with a simple second-degree relationship. The siblings' (I37138-I31508) father must, therefore, be a second-degree relative of their maternal grandfather, in order to accommodate the third-degree relationship between the siblings' parents.

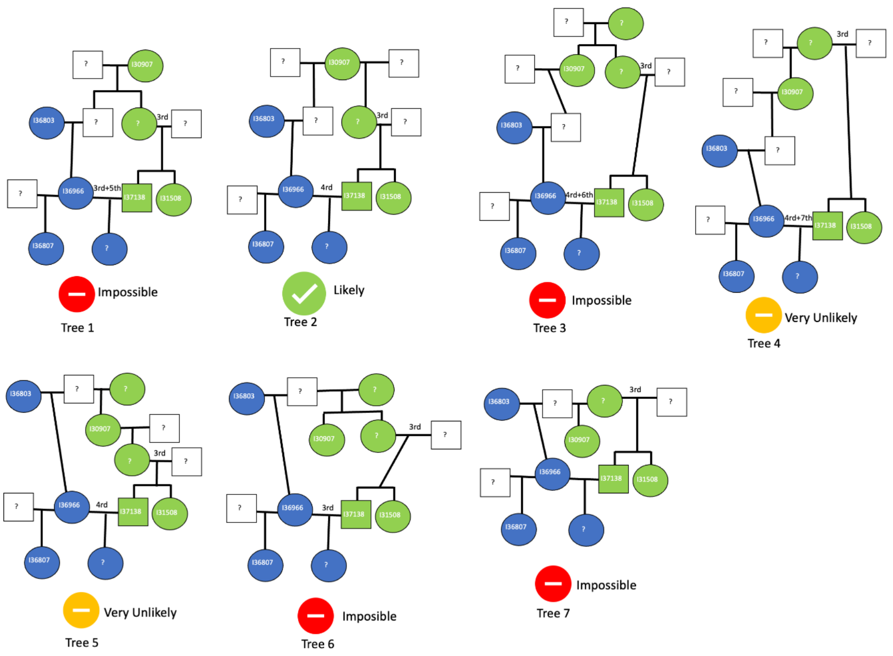


Figure S25. Different scenarios for the relationship between individuals I36966, I30907, I37138 and I31508 at Wetwang Slack.

3) Relationship between I36905 and I37138-I31508:

I31508 and I37138 are third-degree relatives of I36905 and fourth-degree relatives of I36905's son (I31009). Furthermore, I36905-I31009 have IBD sharing in the X-chromosome with I31508 but display different mtDNA lineages to I31508. As such, the genetic transmission has to be through I31508's and I37138's father. The transmission cannot be through their mother or through an unsampled sibling because, in both of these cases, I36905 would be closely related to I30907 (I31508's and I37138's maternal grandmother). For instance, in the case of a relation through an unsampled sibling of I37138-I31508, I30907 and I36905 would be fourth-degree relatives, but I30907 and I36905 are not related at all. Neither can the genetic relationship run through I37138 and I31508's maternal grandfather because, if that was the case, X-chromosome IBD segments between I31508 and I30907 (I31508's maternal grandmother) would not overlap with those between I31508 and I36905 (I31508's maternal grandfather's first-degree relative). This is because I31508's maternal X-chromosome segments would have been inherited from one of her maternal grandparents, but not from both at the same time. In fact, a large portion of the X-chromosome is shared between both I31508-I30907 and I31508-I36905 (Figure S26), which makes a transmission through I37138-I31508's maternal grandfather impossible. A relationship with I36905 through I31508-I37138's father, however, aligns with the position of siblings I30962 and I31509 in the reconstructed pedigree, who are related to I31508-I37138 via I31508-I37138's father and are also related to I36905.

I31508's and I37138's father has to be either I36905's maternal half-brother (he cannot be a paternal half-brother because, if this were the case, they would not share IBD in the X-chromosome, and they do), his maternal uncle, maternal grandfather through a third reproductive partner, his grandson through I36905's unsampled daughter (I31009's half-sister), or his nephew, through I36905's sister.

I36905 shares relatively high numbers of IBD segments with I31508 (n=23) and I37138 (n=20) for third-degree relatives, which provides additional supporting evidence that an unsampled full-sibling relative is involved in the genetic transmission. Thus, we display the scenario with I31508's and I37138's father as I36905's nephew through I36905's sister, but with the addition of red lines to indicate uncertainty.

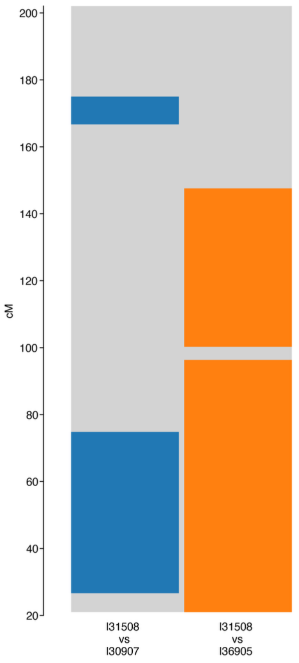


Figure S26. X-chromosome IBD segments for individuals I31508, I30907 and I36905 at Wetwang Slack.

4) Relationship between I36966 and I36807-I36963:

The correct configuration between these individuals is I36803 as the mother of I36966, with I36807-I36963 as I36966's daughters sired by two different fathers. This is because:

-The number of shared IBD segments between I36963 and I36807 is higher (n=32) than between I36963 and I36803 (n=25), I36803 and I36807 (n=24), and I36966 and I36985 (n=22), and maternal half-siblings have more IBD segments than grandparent-grandchildren relationships.

-More importantly, the IBD breakpoints between I36803-I36807 and between I36963-I36803 are closely correlated with those in I36963-I36807, but those between I36803-I36807 are not closely correlated with those between I36963-I36803. This conforms to the expected pattern if I36803 is the grandmother of the other two, who are themselves half-sisters.

5) Relationship between I30985, I36759, I36816, I36815, I36810 and I36849:

- Position of I36810, I36849 and I36815:

  -I36810 is a second-degree relative of I36815 (IBD sharing: 1575 cM, 32 segments) and I36849 (IBD sharing: 1595 cM, 31 segments).

  -I36849 is I36815's third–fourth-degree relative (IBD sharing: 594 cM, 20 segments) and I36810 is a fifth-degree relative (IBD sharing: 175–227 cM) of siblings I36965, I36967 and I37159.

  -Also constraining the model is the relationship between these individuals and I30985, who is a second-degree relative of I36815, a third-degree relative of I36810, and a fifth-degree relative of I36849 (253 cM).

Given these constraints, I36810 must lie one generation below I36815, and, therefore, be a niece of I36815 through I36815's sister. This is the only way to satisfy a third-degree relationship between I36810 and I30985, and a fifth-degree relationship between I36810 and I36965-I36967-I37159. In this scenario, for I36849 to have a fourth-degree relationship with I36815, I36849 must be I36810's grandson.

- Position of I30985, I36759 and I36816:

  -Individuals I30985, I36759 and I36816 have the same Y-chromosome lineage.

  -I30985 is a second-degree relative of I36759 (1647 cM, 29 segments), I36816 (1877 cM, 17 segments) and I36815 (1638 cM, 36 segments).

  -I30985 and I36815 must have an avuncular or maternal half-sibling relationship based on a high number of shared IBD segments (n=36), while I30985 and I36816 must have a grandparent-grandchild or paternal half-sibling relationship based on the low number of shared IBD segments (n=17).

  -I36759 is a third-degree relation of I36815 (621 cM, 16 segments) and a fourth-degree relation of I36816 (549 cM, 12 segments).

  -I36816 is a fifth–sixth-degree relative of I36815 (138 cM, 6 segments).

Given these constraints, I30985 is a paternal or maternal grandfather of I36816 (we display paternal in Figure S28 because they share the Y-chromosome haplogroup and do not share IBD in the X-chromosome) and a maternal uncle of I36815 (they share IBD in the X-chromosome and have the same mtDNA lineage). I36759 could be I30985's paternal nephew or paternal half-brother. The only minor issue is that this configuration dictates that I36816 and I36815 are fourth-degree relatives, but they share 137 cM in IBD, which is very low for a fourth-degree relation, although not impossible.

6) Relation between I36815 and siblings I36965, I36967 and I37159.

The three siblings (I36965, I36967 and I37159) are third-degree relatives of I36815. The possible scenarios are:

Hypothesis 1: I36965, I36967 and I37159 are I36815's great-grandchildren.

Hypothesis 2: I36965, I36967 and I37159 are I36815's brother's descendants

Hypothesis 3: I36965, I36967 and I37159 are I36815's half-brother's descendants.

Hypothesis 4: I36965, I36967 and I37159 are I36815's paternal cousins.

The most likely scenario is Hypothesis 1 because it is the only one that allows a fifth-degree relationship between siblings I36965, I36967 and I37159, and individual I36810 (with prior knowledge that I36810 and I36815 cannot be half-sisters), which is the observed degree of relation between the siblings and I36810.

7) Relationship between I36820, I36806 and I30906:
I30906 and I36806 share 630 cM in IBD, but I30906 is not closely related to I36806's father (I36821), and I36806 is not closely related to I30906's mother (I31020). Individual I30906 can, therefore, be neither I36806's paternal relative (because he would be even more closely related to I36821 than I36806 is to I36821), nor the descendant of I36806's sibling (because, then, he would be equally related to I36821 and I36806, and he is not), nor the descendant of I36806 (because, then, he would be one degree more distant to I36821 than I36806 is to I36821, and he is not). I30906 must, therefore, be a maternal relative of I36806. He cannot, however, be I36806's maternal ancestor or I36806's maternal ancestor's sibling, because, then, I30906's mother (I31020) would be one degree more distant to I36806 than I30906 is to I36806, or equally related to I36806, and she is not. Individual I30906 must, therefore, be the paternal half-brother of I36806's maternal grandparent. I36820 and I36806 must be maternal half-sisters because:

-I36806 cannot be I36820's grandmother or aunt because I36806's father (I36821) and I36820 are not third-degree relatives.

-I36806 cannot be I36820's granddaughter or niece because, then, I36820 would be unrelated to I30906 (in the granddaughter scenario) or I30906's third-degree relative (in the niece scenario). I36820 and I30906 share ~400 cM, compatible with a fourth-degree relationship.

8) Relationship between I30988 and her first-degree relatives:
I30988 is the first-degree relative of I36821, I30965, I30969 and I39733; in all four cases this is a parent-offspring relationship. All of the relationships between I36821, I30965, I30969 and I39733 are second degree, which leaves only two possible scenarios:
   -The four individuals are sons/daughters of I30988, each from a different father.
   -One of the four is the mother/father of I30988, and the other 3 are I30988's offspring, each from a different father.
The number of IBD segments shared between the individuals is as follows: I36821-I30965 (n=35), I36821-I30969 (n=35), I30965-I30969 (n=28), I36821-I39733 n=(28), I39733-I30969 (n=20) and I30965-I39733 (n=23). IBD sharing between the first two pairs (n=35) is too high for a grandparent-grandchild relationship, and this evidence suggests instead that I36821, I30965 and I30969 are maternal half-siblings, while I39733 could be the grandmother of the other three. Now we must consider the IBD breakpoint distribution:
   -The IBD breakpoints correlate extremely well between pairs I36821-I30969, I30965-I30969 and I30965-I36821, which is expected if the three individuals are half-siblings, because a recombination in one individual will be observed in the comparison between that individual and the other two. If, for instance, I30969 were the grandmother of I36821 and I30965, IBD breakpoints in I36821-I30969 and I30965-I30969 should not correlate.

-The IBD breakpoint locations observed in the comparison between I39733 and each of the other three individuals do not correlate, but breakpoints observed in, for example, I39733-I36821 (that emerged in I36821's gamete), are all seen in I30965-I36821 and I30969-I36821, which is the expected pattern if I39733 is their grandmother. If I39733 was a half-sister, some breakpoints observed in I39733-I36821 (those emerging in I39733), would not appear in I30965-I36821 and I30969-I36821.

The evidence, therefore, suggests that individual I39733 is the grandmother of I36821, I30965 and I30969, who are half-siblings from three different fathers and the same mother (I30988).

9) Relationship between I36797 and I36865-I36803:
I36797 shares 38 and 34 IBD segments with I36865 and I36803, respectively. This is too high for a grandparent-grandchild or a paternal half-sibling relation. I36797 could then be:
   -Maternal aunt to both I36865 and I36803, who would themselves be cousins. Although she could be a paternal aunt as well because, in that scenario, X-chromosome sharing would likewise be expected, we have placed I36797 as a maternal aunt due to the fact that they share the same mtDNA lineage.
   -Maternal (or paternal; see above) aunt to one of I36865 or I36803 and a maternal half-sister of the other.
I36865 and I36803 share 23 IBD segments, which is rather high if the pair were third-degree relatives. This suggests that a full-sibling relationship is involved in the genetic transmission, and is the reason why I36797 is shown as a maternal aunt of both I36865 and I36803, although with red lines to indicate uncertainty.

10) Relationship between I37166 and I36830:
Given that individual I36830 was 17–20 years old at the time of death and I37166 individual was 35–45 years old, I36830 is more likely to be the daughter of I37166. This is confirmed by IBD because I36860 and I36861 share four segments with I36830 but none with I37166. If I36830 were the mother, we would expect some of these segments to be passed to I37166.

11) Relationship between I37314, I36866 and I37342:
I37342 is the maternal half-brother of siblings I37314 and I36866. He cannot be their uncle, because the siblings share only 12 cM in IBD0 which at the same time are IBD1 in I37342-I37314 and I37342-I36866, and we would expect ~200 cM of those types of segments for an avuncular relationship. Neither can I37342 be the nephew of I37314 and I36866, because if this were the case, he would be closely related to I39733 (a close relative of the siblings), and he is not.

12) Relationship between I30977, I37164, I37168 and I37308:
I30977 and I37164 are second-degree relatives, but they do not share the entire X-chromosome and therefore, they cannot be paternal half-sisters or paternal grandmother-granddaughter. Individual I37164 can only be I30977's paternal aunt or niece. I37308 is I37164's third-degree relative (with X-chromosome IBD sharing) and I30977's fourth-degree relative (with no X-chromosome IBD sharing). If I37164 were I30977's niece, I37308 would not fit the pedigree because for him to be closer to I37164 than to I30977, he would need to be I37164's descendant and thus I30977's fifth-degree relative, not fourth-degree relative as he is. Individual I37164 must, therefore, be I30977's paternal aunt, and I37308 a second-degree relative of I37164's father.

I37308 must be a paternal relative of I37164 and I30977 because of X-chromosome sharing (Figure S27). I37164 and I30977 share two long segments in the X-chromosome, deriving from I37164's mother. I37164 and I37308 also share a long segment that overlaps with one of the two segments shared by I37164 and I30977, but I37308 and I30977 do not share IBD segments in this region. To accommodate this evidence, I37164 must share IBD with I30977 through her maternal X-chromosome, and with I37308 through her paternal X-chromosome; the latter is not shared with I30977, who only inherited her paternal grandmother's X-chromosome.

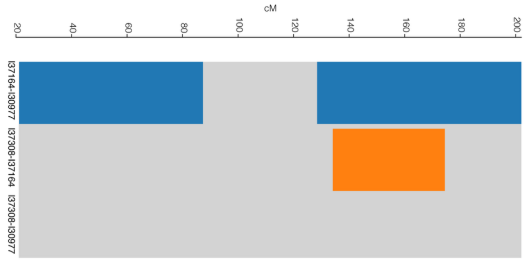


Figure S27. X-chromosome IBD segments for individuals I30977, I37164, and I37308 at Wetwang Slack.

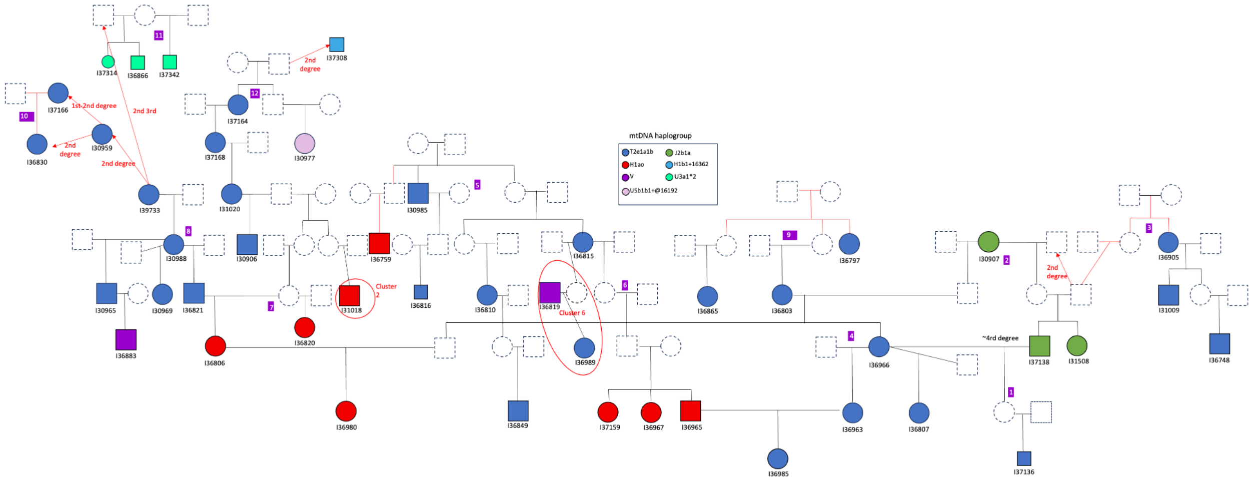

Figure S28. Reconstructed pedigree for cluster 5 at Wetwang Slack.

##### *1.6. Cluster 6*

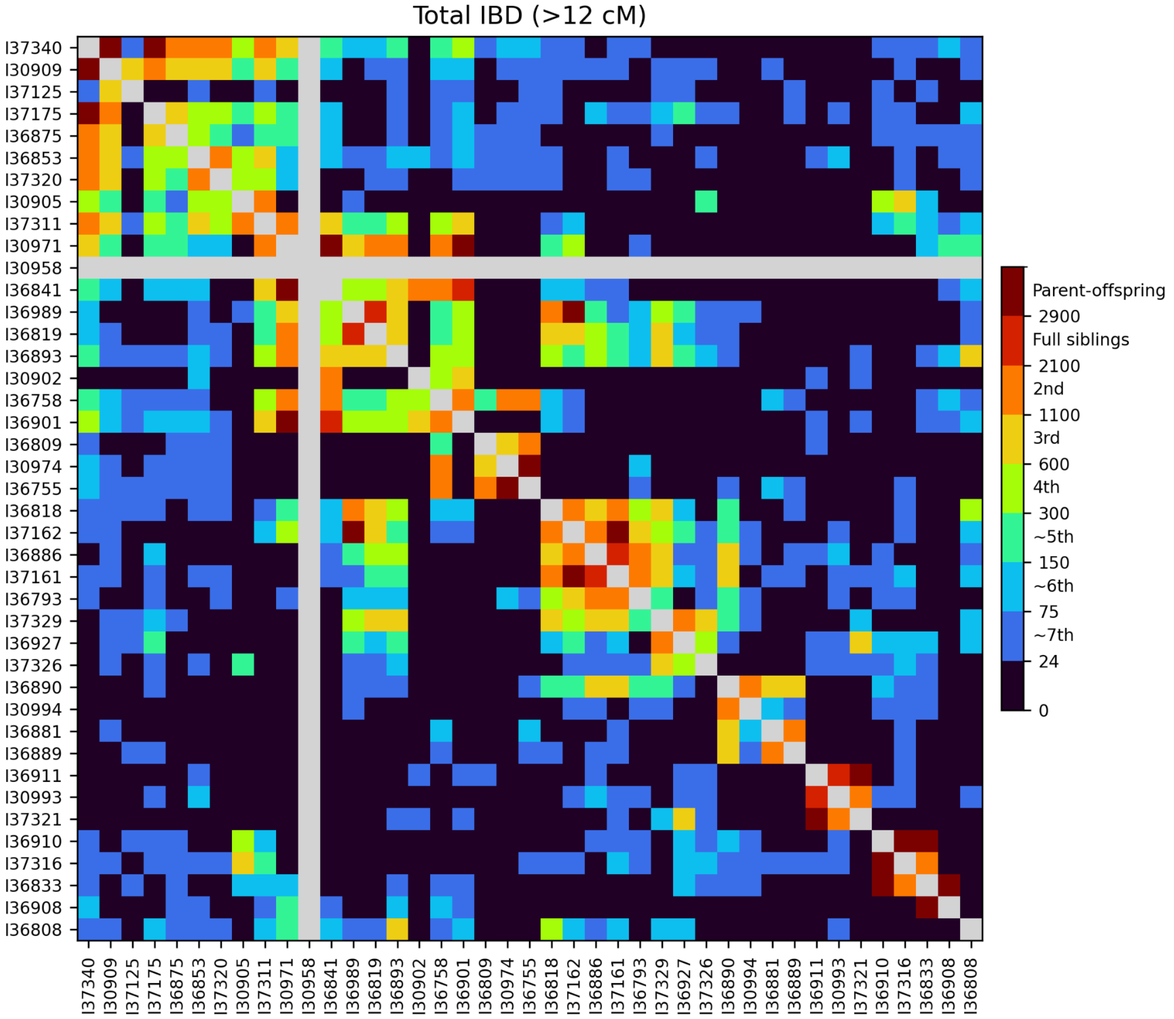


Figure S29. Heatmap of IBD sharing between pairs of individuals from cluster 6 at Wetwang Slack.

Cluster 6 comprises 41 individuals (Figure S29; Figure S36), detailed below.

1) Relationship between I30970 and I36886-I37161:

Individual I30970 is a third-degree relative of I36886 and I37161:

- The relationship cannot, however, proceed through I36886-I37161's father, because I30970 and I37161 share 49% of the X-chromosome in IBD.
- Neither can the relationship proceed only through I36886-I37161's mother, because, if this were the case, I30970 would be either:
    - I36886's and I37161's mother's half-brother: this is impossible because I36886's and I37161's mother and I30970 do not share the same mtDNA haplogroup (excluding the possibility that they are maternal half-siblings) but they do share IBD in the X-chromosome (excluding the possibility that they are paternal half-siblings).
    - I36886's and I37161's mother's nephew: this is impossible because I36886's and I37161's mother and I30970 do not share the same mtDNA haplogroup (which excludes the possibility that I30970 is their nephew via I36886's and I37161's mother's sister) and they share IBD in the X-chromosome (which excludes the possibility that I30970 is their nephew via I36886's and I37161's mother's brother).

- I36886's and I37161's mother's grandson via someone other than I36886's and I37161's father): this is impossible because I36886's and I37161's mother and I30970 do not share the same mtDNA haplogroup (excluding the possibility that I30970 is her grandson via I36886's and I37161's mother's unsampled daughter) but they do share IBD in the X-chromosome (excluding the possibility that I30970 is her grandson via I36886's and I37161's mother's son).
- I36886's and I37161's mother's grandfather: this is impossible because I36886's and I37161's mother and I30970 share IBD segments in the X-chromosome (excluding the possibility that I30970 is her paternal grandfather) and because I30999 (I30970's grandmother) is not fifth-degree relative of I36886-I37161 (25 cM in shared IBD), excluding the possibility that I30970 is I36886's and I37161's mother's grandfather, either paternal or maternal
- I36886's and I37161's mother's uncle: this is impossible because I36886's and I37161's mother and I30970 do not share the same mtDNA haplogroup (excluding the possibility that I30970 is her maternal uncle) and because I30999 is not fourth-degree relative of I36886-I37161 (25 cM in shared IBD), which excludes I30970 as I36886's and I37161's mother's paternal or maternal uncle.

The relationship must, therefore, proceed through both I36886's and I37161's mother and father, i.e. through a descendant of I36886's and I37161's parents. The only remaining possibility is that an unsampled brother of I36886 and I37161 is the maternal grandfather of I30970. The unsampled sibling cannot be the paternal grandfather/grandmother of I30970 because, in that scenario, I30970 would not share IBD in the X-chromosome with I36886-I37161. Likewise, the unsampled sibling cannot be the maternal grandmother of I30970 because, in that case, I30970 would share the mtDNA haplogroup with I36886-I37161; furthermore, this position is already occupied by I30999 who is not I36886's and I37161's sister. This scenario also aligns well with shared IBD patterns along the chromosomes, because the IBD segments in I30970-I36886 and I30970-I37161 appear exclusively in the regions of the genome that I30970 does not share with I30999 (his maternal grandmother), i.e. those regions that I30970 inherited from I30999's reproductive partner (his maternal grandfather).

I36793 (14–18 years old) is a second-degree relative of I36886-I37161 and a third-degree relative of I37162-I30970. She shares IBD in the X-chromosome with I30970 but not mtDNA haplogroup. She must, therefore, be the paternal half-sister of I30970's mother.

2) Relationship between I36818 and I37162:
I36818 is a second-degree relative of I37161, I36989 and I37162, and must, therefore, be the grandson of I37161 and I36989 through an unsampled daughter.

3) Relationship between I30974, I36755 and I36809:
I30974-I36755 are a mother-son pair (order, as yet, unknown). They are both second-degree relatives of I36758 but are not related to I36758's maternal family, which means that I36758 has to be the paternal grandson/nephew of I30974-I36755. Given that I36758-I36755 share many more IBD segments than I36758-I30974, it is likely that I30974 is I36758's grandmother and that I36755 as I36758's paternal aunt. I36809 is a second-degree relative of I36755 and a third-degree relative of I30974. As such, he must be I36755's grandson.

4) Relationship between I30902 and I30958-I36901-I36841:

I30902 is a second-degree relative of siblings I30958, I36901 and I36841 but is unrelated to the siblings' mother (I30971), which means that I30902 must be a first-degree relative of the siblings' father. She shares 28 IBD segments with I36841 and is a third-degree relative of siblings I37159, I36967 and I36965 from cluster 5. These siblings (I37159-I36967-I36965) are fifth-degree relatives of siblings I30958-I36901-I36841. Thus, I30902 could be the paternal half-sister, paternal grandmother or paternal aunt of I30958, I36901 and I36841. If I30902 was their paternal half-sister or grandmother, she would share the entire X-chromosome in IBD with sisters I30958 and I36901. If she was their paternal aunt, she would not necessarily share the entire X-chromosome with sisters I30958 and I36901, but the shared X-chromosome IBD segments between I30902-I36901 and I30902-I30958 would be the same. In fact, large portions of the X-chromosome are not shared between the sisters and I30902, but the IBD pattern in I30902-I36901 and I30902-I30958 is almost identical (the difference likely due to phasing errors) (Figure S30). Individual I30902 must, therefore, be the paternal aunt of siblings I30958, I36901 and I36841. This is further confirmed by the presence of regions of the genome in IBD1 between each of the siblings and I30902, and at the same time in IBD0 between the siblings, which would only be the case if I30902 is their aunt.

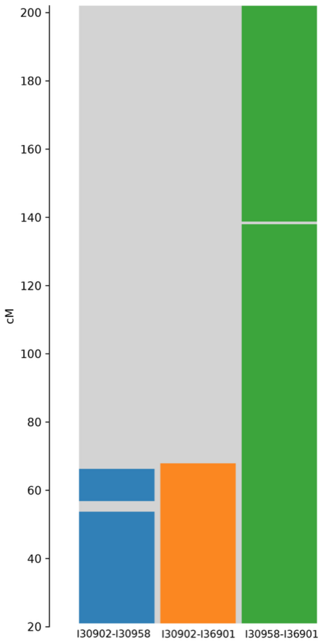


Figure S30. X-chromosome IBD patterns between I30902, I30958 and I36901 at Wetwang Slack.

5) Relationship between I30909, I37340 and I37175:

I30909-I37340 are a mother-daughter pair. Since I30909 is 17–18 years old and, more importantly, since she is a third-degree relative of I37125 but I37340 is not, I30909 must be the daughter. The available evidence suggests that I37175 is I37340's son and not her father, because:

-I30909-I37175 have 31 shared IBD segments, and are therefore more likely to be maternal half-siblings than paternal grandfather and granddaughter.

-I37175 and I37340 share the same mtDNA lineage and are therefore more likely to be son and mother.

-All four of I37340's second-degree relatives are equally related to I30909 and I37175. If I37175 were I37340's father, we would expect some of I37340's close relatives to be either more closely related to I37175 than to I37340 (if they were related to I37340 through her paternal line) or not related at all (if they were related to I37340 through her maternal line). The other possible scenario to accommodate I37175 as the father of I37340 and all I37340's second-degree relatives being equally related to I30909 and I37175, is that all of I37340's second-degree relatives are her grandchildren from different fathers (i.e. not I30909's father). This is not the case, however, because I37320, I36853 and I37340 are all second-degree relatives, which means that at least two of them must be half-siblings. Half-siblings I37320 and I36853 are not I37340's grandchildren because the IBD breakpoints between I37340-I36853 and I37340-I37320 are closely correlated, which does not align with the expected pattern for such a relationship.

6) Relationship between I30909 and I37125:
I37125 is I30909's third-degree relative, but they do not share IBD in the X-chromosome. Neither does I37125 share IBD with I37175 or I37340, and so she must be a second-degree relative of I30909's father. We can discount a scenario whereby I37125 is I30909's father's grandmother or aunt, because, if that were the case, all of I37125's relatives would also be I30909's relatives (with the addition of three degrees to each). However, although siblings I37159, I36967 and I36965 are I37125's fourth-degree relatives, they do not share a single IBD segment with I30909; as a seventh-degree relative in this scenario, we would expect at least some shared IBD segments with at least one of the siblings. We, therefore, display individual I37125 as I30909's cousin, with red lines to indicate uncertainty.

7) Relationship between I30971, I30905, I37311 and I37340:
I37311 is a second-degree relative of I30971 and I30905. While in both cases they share IBD in the X-chromosome, this relates to only a portion of the chromosome. I30971 and I30905 are unrelated. The following scenarios are possible so far: either I30971 and I30905 are I37311's maternal and paternal aunts, maternal and paternal grandmothers, maternal and paternal half-siblings, or a combination of the three possible types. I30971 cannot, however, be I37311's paternal grandmother or half-sister because they do not share IBD segments across the entire X-chromosome; I30971 must therefore be I37311's paternal aunt. We display I30905 as maternal aunt (Figure S36) but use red lines to indicate two other possible configurations (that she is a maternal grandmother or half-sister). I37340 is a second-degree relative of I37311, a third-degree relative of I30971 and a third/fourth-degree of I30905. Individual I37340 must, therefore, lie below I37311 in the pedigree, as either I37311's maternal niece or granddaughter. If I37340 were the granddaughter of I37311, she would be a fourth-degree relative of both I30971 and I30905, whereas she is, in fact, a third-degree relative of I30971. Alternatively, if I37340 were the niece of I37311, she would be a third-degree relative of both I30971 and I30905. The more likely scenario, therefore, is that I37340 is the niece of I37311, with I37340-I30905 being third-degree relatives. Placing I37340 as the niece allows us to accommodate I36853 and I37320 as I30905's fourth-degree relatives. If I37340 was I37311's granddaughter, I36853 and I37320 would be fifth-degree or more distant relatives of I30905, not fourth-degree as the evidence suggests).

8) Relationship between I37340, I37320, I36853 and I36875:

I37320, I36853 and I37340 are all second-degree relatives, which means that at least two of them must be half-siblings.

-I37340 cannot have half-siblings because, if this were the case, then they would either be second-degree relatives of I37311 (if they were maternal half-siblings) or unrelated to I37311 (if they were paternal half-siblings). Individual I37340 could, however, be grandmother or aunt to both I37320 and I36853, if they themselves were maternal half-siblings.

-The scenario in which I37340 is grandmother of I37320 and I36853 is very unlikely because, if this were the case, the IBD breakpoints between I37340 and I37320 should not correlate with those between I37340 and I36853, and they do. Furthermore, if I37340 were the grandmother, I36853 would be a fourth-degree (not third-degree) relative to I37311 and at least a fifth-degree (not fourth-degree) relative of I30905.

-I36875 must be I37340's grandson from a third reproductive partner of I37340, to accommodate the fact that he is a fourth-degree relative of I36853 and I37320.

9) Relationship between I30971 and I36893:

I30971 and I36893 are second-degree relatives, while I37311 is a second-degree relative of I30971 and a fourth-degree relative of I36893. Individual I36893 cannot, therefore, be I30971's grandfather, uncle, maternal half-brother or nephew, because, in those scenarios, I36893 would be a third-degree relative of I37311's. As such, I36893 must be I30971's grandson through an unsampled daughter.

10) Relationship between I36819, I36989, I30971, I36893 and I36883 (the last one from cluster 5):

Although sample I36819 was contaminated, the distribution of IBD segments along the genome indicates a parent-offspring relationship with I36989. As such, they must be a father-daughter pair. Since I36819 is more closely related to I30971 (at the second–third-degree), I36893 (second–third-degree) and I36883 (third-degree) than is I36989, who is their third-, third- and fourth-degree relative respectively, the genetic relationship must pass through I36819. Furthermore, shared IBD segments between I30971-I36989 and I36893-I36989 appear to be a subset of those observed in I30971-I36819 and I36893-I36819. I36883 is I36819's third-degree relative, I36989's fourth-degree relative, I30971's and I36893's third-degree relative. These relationships are, in all cases, observed in the X-chromosome, and must therefore have progressed through I36883's mother (I36883's father is unrelated). I36883 appears equally related (at the third degree) to I30971 and I36893. I36989 is likewise a third-degree relative of I30971 and I36893, while I36819 is a second–third-degree relative of I30971 and I36893. If I30971-I36893 are indeed grandmother-grandson, then the only scenario that accommodates the relationship between I30971-I36893 and the other three (I36883, I36989 and I36819) is if it passes through I36893's half-siblings, who would be equally related to their half-brother (I36893) and to their grandmother (I30971). As such, I36893 is the maternal half-brother of I36819 and the maternal half-brother of I36883's mother.

11) Relationship between I30994, I36890, I36881, I36889, I37161 and I36886:

I30994 and I36890 are second-degree relatives sharing the same X-chromosome but not the same mitochondrial haplogroup. As such, I30994 could be either I36890's maternal grandfather or their paternal uncle. Two pairs of close relatives, siblings I37161 and I36886 and second-degree relatives I36881 and I36889, are related to I36890 via third-degree relationships, but they are not closely related to the other pair or to I30994. If I30994 were the paternal uncle, in order to accommodate the lack of relationships between I30994, I36881-I36889 and I37161-I36886, siblings I37161-I36886 would have to be related to I36890 through I36890's maternal grandfather, and to I36881-I36889 through I36890's maternal grandmother (they share mtDNA with I36890). The X-chromosome patterns indicate, however, that such a configuration is impossible because, in this scenario, I36890 would have inherited her maternal X-chromosome segment either from her maternal grandfather or from her maternal grandmother, but not both simultaneously. As such, X-chromosome regions where I36890 shares IBD segments with I37161-I36886, cannot also be shared between I36890 and I36881-I36889. We observe such regions (Figure S31), which allows us to discard this configuration. To accommodate the observed patterns, I30994 must be I36890's maternal grandfather. As such, I36890 must be related to I36881 and I36889 through her maternal grandmother and to I37161-I36886 through her father. As expected in such a scenario, X-chromosome regions shared between I36890-I30994 and between I36890-I36881 are mutually exclusive.

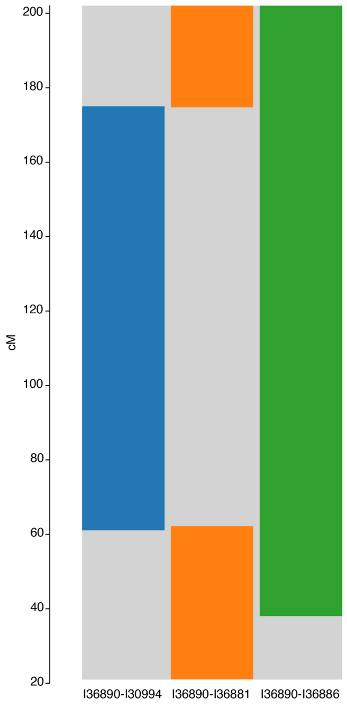


Figure S31. X-chromosome IBD segments for I36890 and individuals I36881, I30994 and I36886 at Wetwang Slack.

12) Relationship between I36890, I36889 and I36881:

We have established that I36889 and I36881 are third-degree relatives of I36890 via I36890's maternal grandmother, who must, in turn, be the mother or sister of I36889 and I36881. Since I36889 and I36881 are themselves second-degree relatives,

I36890's maternal grandmother could be either I36889 and I36881's mother (who reproduced with two different men, themselves distinct from I30994), or the mother of one of them and sister of the other. Again, X-chromosome IBD patterns allow us to determine the correct configuration. The scenario whereby I36881, I36889 and I36890's mother are maternal half-sisters from three different fathers is impossible because, if they were maternal half-siblings, I36881, I36889 and I36890's mother must have inherited one of the two X-chromosomes of their common mother. If all three did indeed inherit the same maternal chromosomal segment from their common mother, if one individual shares IBD in that specific region with the other two, the other pairwise comparison must also show IBD in that region. However, in the X-chromosome regions not shared between I36890 and her maternal grandfather (I30994) (i.e. those that I36890 inherited from her maternal grandmother), although I36881 shares a large IBD segment with both I36889 and I36890, I36889 and I36890 do not share IBD in this specific region (Figure S32). Similarly, if I36889 were I36890's mother's maternal uncle and I36881 were I36890's mother's maternal half-sister, I36890, I36881 and I36889 would only inherit one of the three X-chromosomes present in I36889's parents in any given region (and, in the case of I36889, only one of two present in his mother). In the region of the X-chromosome that I36890 has inherited from her maternal grandmother, I36881 shares one IBD segment with both I36889 and I36890. This means that the three individuals must have inherited the same chromosome, and we would therefore expect I36889 and I36890 to share IBD in this region, but they do not (Figure S32). We, therefore, conclude that I36889 is the maternal half-sister of I36890's mother and I36881 is I36890's mother's maternal aunt. Under this configuration, I36881 and I36889's mother can inherit the same maternal and paternal chromosomes at any given region. If I36889's mother then passed one of her chromosomes to I36889 and the other to I36890's mother, who would in turn pass it to I36890, I36881 would share IBD with both I36889 and I36890, but I36890 and I36889 would not share IBD in that region, which is exactly what we observe.

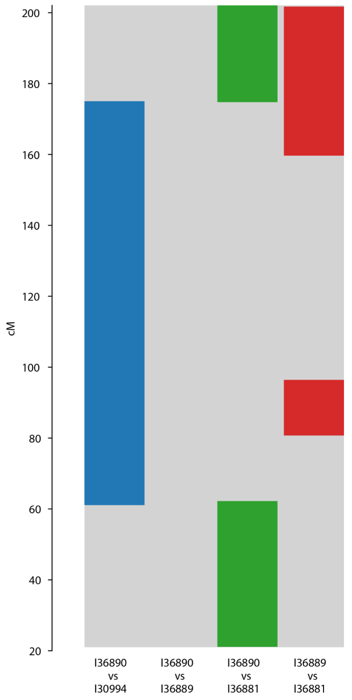

Figure S32. Shared X-chromosome IBD segments for I36890 and individuals I36881, I36889 and I30994 at Wetwang Slack.

13) Relationship between I36890, and siblings I37161 and I36886:
Siblings I37161 and I36886 are related to I36890 via I36890's father, who must be their second-degree relative. I36890's father could be their maternal half-brother (not their paternal half-brother because we detect X-chromosome IBD sharing between the siblings and I36890), their maternal nephew, their maternal uncle or their maternal grandfather. The only possible configuration is to set I36890's father as the siblings' nephew through their unsampled sister. The other three configurations are impossible due to the patterns of X-chromosome sharing (Figure S33): I36886 shares a large IBD segment with both I37161 and I36890, which derives from the siblings' mother, but I37161 and I36890 do not share IBD in this region. If I36890's father was the siblings' maternal half-brother, the shared IBD segment between I36886 and I36890 would mean that they must have inherited the same chromosome from I36890's paternal grandmother in that region. In this scenario, if I37161 shared IBD segments with I36886, then he must also share IBD with I36890 (having derived his X-chromosome only from his mother, I36890's paternal grandmother). However, I37161 and I36890 do not share IBD. If I36890's father was the siblings' nephew through their unsampled sister, then I36886 would share IBD segments with her unsampled sister (who in turn passed them to her granddaughter (I36890) via the X-chromosome coming from I36890's father). I36886 could, however, also share IBD segments with her brother via the X-chromosome coming from her mother and so, in this scenario, I36890 and I37161 need not share IBD in that region.

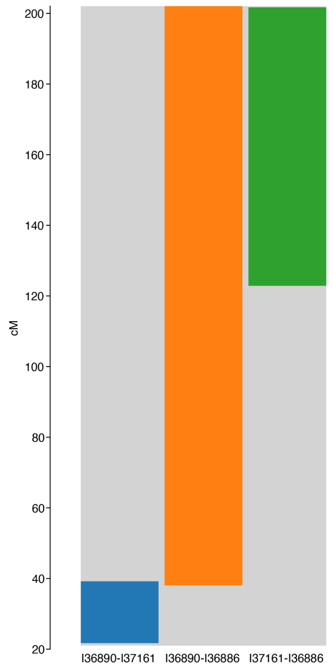


Figure S33. Shared X-chromosome IBD segments between I36890, and individuals I37161 and I36886 at Wetwang Slack.

14) Relationship between I36927, I37329, I37326, I37321, I36911 and I30993:

I36911 is the mother of I37321 and the sister of I30993. I36927 is I37321's third-degree relative but she is not closely related to I36911, which means that she is a second-degree of I37321's father, sharing IBD in the X-chromosome. I36927 is I37329's second-degree relative and I37326's fourth-degree relative (in both cases sharing IBD in the X-chromosome). I37329 is I37326's third-degree relative, with no X-chromosome sharing. There are many different possible configurations for the relationship of I36927 and I37321, and for the relationship between I36927, I37329 and I37326. However, only three scenarios accommodate the degrees of relationship between all the individuals: that I36927 is either I37329's maternal half-sister, maternal aunt or maternal niece. A scenario whereby I36927 is I37329's maternal grandmother is unlikely due to the high number of shared autosomal IBD segments. In all three plausible scenarios, I36927 is the maternal grandmother of I37321's father (she cannot be his paternal grandmother as they share IBD in the X-chromosome). I36927 cannot, however, be I37329's maternal niece, since this is not supported by patterns of X-chromosome sharing (Figure S34). In this configuration, I37326 must be a second-degree relative of I37329's father. Since I37326 and I36927 share one X-chromosome IBD segment, of the three X-chromosomes present in her maternal grandparents, I36927 must have inherited that region from her maternal grandfather (who is I37326's close relative). In this scenario, I37329 would also share IBD in that region with I36927 and I37326, having inherited that same chromosome from her father (I36927's maternal grandfather), but this is not the case. Either of the other two configurations, with I36927 as I37329's maternal aunt or maternal half-sister, are possible. We show one of them in Figure S36 but use red lines to reflect this degree of uncertainty. In both cases, I37329 is I37326's great-aunt and their genetic connection must go through both a male and a female relative (to account for the different mitochondrial haplogroups and X-chromosome sharing between I37326 and I36927).

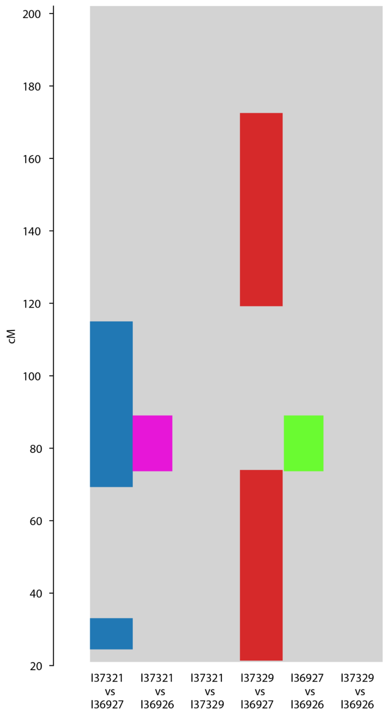


Figure S34. Shared X-chromosome IBD segments for individuals I36927, I37329, I37326 and I37321 at Wetwang Slack.

15) Relationship between I37329, and siblings I37161 and I36886:

Siblings I37161 and I36886 are third-degree relatives of I37329, and more distantly related to I37329's closest relatives (I36927 and I37326). Autosomal IBD segments shared between the siblings and I36927 are a subset of those shared between the siblings and I37329 (and the same is true for I37326) (Figure S35), which means that the relationship between the siblings and I36927-I37326 must be via I37329. As such, the siblings must be I37329's descendants and, more specifically, I37329's great grandchildren through the siblings' father and I37329's daughter. We can discount the scenario whereby the genetic transmission takes place via two male relatives or two female relatives because I36886 and I37329 have different mitochondrial haplogroups and because patterns of X-chromosome sharing do not support this configuration. Neither can the genetic transmission take place through the siblings' mother because of a lack of X-chromosome sharing between I37161 and I37329. Nor can the transmission progress through I37329's parents or siblings, because, in this scenario, the siblings would share IBD segments with I36927 and I37326 that are not present between I37329 and the siblings, and no such segments are detected. The configuration that places siblings I37161 and I36886 as the great grandchildren of I37329 through I37329's daughter aligns well the third-degree relationships observed between I37329 and I36819-I36893, who are approximately fourth-degree relatives of one of I37161-I36886's parents based on the sixth-degree relationship between I37161 and his reproductive partner I36989 (I36819's daughter).

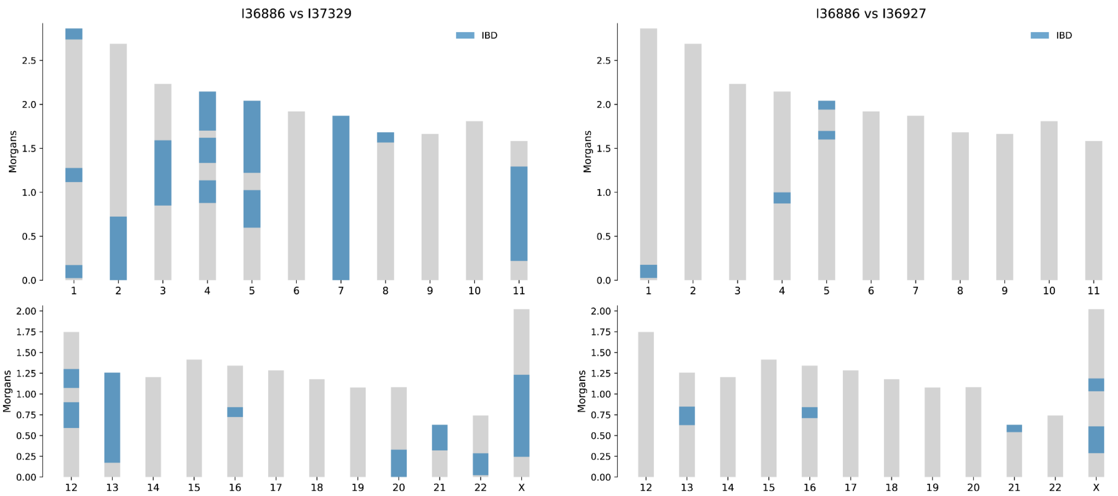


Figure S35. IBD segments between pairs of individuals I36886-I37329 and I36886-I36927 at Wetwang Slack.

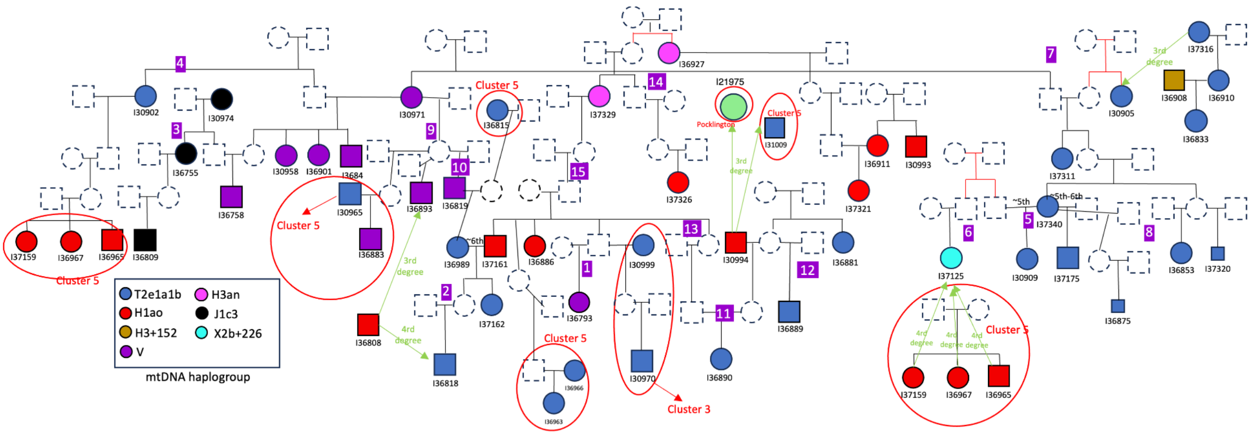


Figure S36. Reconstructed pedigree for cluster 6 at Wetwang Slack.

##### *1.7. Chariots cluster*

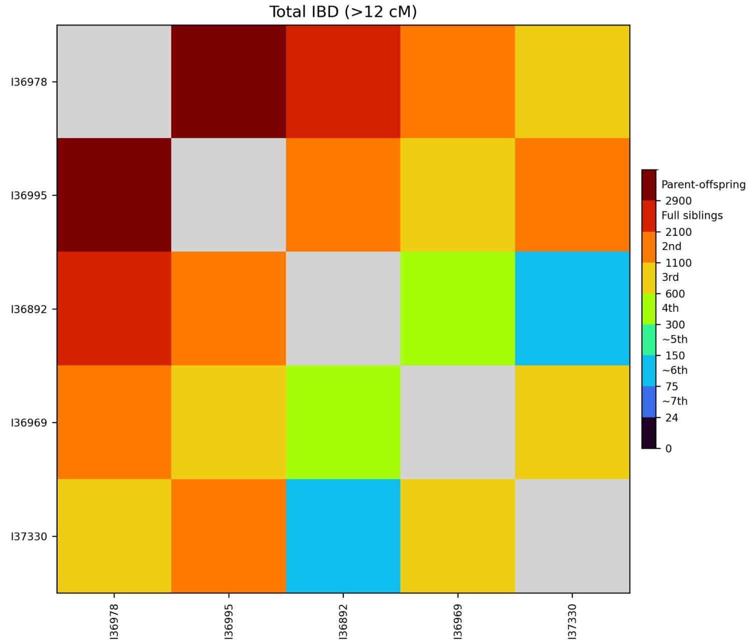


Figure S37. Heatmap of IBD sharing between pairs of individuals from the chariots cluster at Wetwang Slack.

The chariots cluster consists of five individuals (Figure S37): three chariot burials (I36995, I36892 and I36978) and two individuals from non-chariot burials. Of the non-chariot burials, I37330 was buried 25 m east of the chariot burials, while the other (I36969) was buried in the main cemetery, 480 m east of the chariot burials. The relationships between these individuals are reconstructured as follows:

I36978 is the mother of I36995, since they share the same mitochondrial haplogroup and IBD segments across the entire autosomal genome. They also share 290 cM in IBD2, which indicates that I36995's father was closely related to I36978. This is confirmed by ROH segments in I36995 (Supplementary Table 1), which strongly suggest a third-degree relationship between his parents. Female I37330 is a second-degree relative of I36995 and a third-degree relative of I36978, with a different mitochondrial haplogroup from both I36995 and I36978. I37330 cannot be a descendant of I36978 through I36995's unsampled full-sibling because, if this were the case, she would have the same degree of relationship to I36995 as to I36978. Neither can I37330 be I36995's descendant because, in this scenario, IBD segments shared between I36978 and I37330 would be a subset of those shared between I36995 and I37330, and this is not the case because some IBD segments (including in the X-chromosome) shared between I36978 and I37330 are not present in the I36995-I37330 comparison. I37330 could be both I36995's paternal aunt and I36978's first cousin. This resolves the endogamic loop observed in I36995 because, in this scenario, I36995's father would also be I36978's first cousin, supporting the observed shared IBD2 segments and the results of endogamy analysis. Furthermore, it accommodates both the predicted second+fourth-degree relationship and the observed shared IBD2 segments between I36995 and I37330. Female I36969 is a second-degree relative of I36978 and a third-degree relative of I37330. I36978 and I36969 have different mitochondrial haplogroups but share the entire X-chromosome in IBD, which strongly suggests a paternal grandmother or paternal half-sister relationship. If, however, I36969 were I36978's granddaughter sired by an individual other than I36995's father, I37330 and I36969 would be fifth-degree relatives, rendering this configuration extremely unlikely. Similarly, if I36969 were I36978's grandmother, I36969 would need to be I37330's paternal grandmother in order to accommodate the third-degree relationship between I36978 and I37330 (and I36995's father). However, I37330 and I36969 are more likely third-degree (not second-degree) relatives. Even if we accept a second-degree relation, they cannot be paternal grandmother-granddaughter because they do not share IBD segments across the entire X-chromosome. Thus, the only possible configuration is that I36969 is I36978's paternal half-sister and, like I36978, a first cousin of I37330 (Figure 4b). This relationship must pass through the father of I36978 and I36969, and either through the mother or father of I37330. This configuration necessitates that IBD segments shared in the X-chromosome between I37330 are I36978 are identical to those shared between I37330 and I36969, and that those shared between I37330 and I36995 identical or a represent a subset of those shared between I37330 and I36978, and this is precisely what is observed.

The only remaining individual to be placed in the tree is male I36892, who shares the same mitochondrial haplogroup as I36978 but does not share IBD in the X-chromosome with them. I36978 and I36892 have IBD values at the boundary between full-siblings and second-degree relatives. However, they share 124 cM in IBD2, far less than the expected value for a sibling relationship (~800 cM). This suggests, therefore, that they are second-degree relatives, but with the possibility of additional relationships deriving from I36892's high levels of inbreeding (his parents are approximately third-degree relatives based on the length of ROH segments. Since I36892 is one degree closer in relationship to I36978 than to I36995, I36892 is very unlikely to be a descendant of I36978 through I36995's full-sibling because, if this were the case, I36892 would have the same degree of relationship with I36995 as with I36978. Likewise, I36892 is very unlikely to be a descendant of I36978 through I36995 because, in this scenario, I36892 would be more closely related to I36995 than they are to I36978. Individual I36892 could be:

- The uncle or grandfather of I36978, but this is impossible because I36892 would then be a second-degree relative of I37330 and I36969 (if related through I36978's father), or not related to I37330 and I36969 (if related through I36978's mother), rather than the fourth-fifth-degree relationship that we observe.
- The maternal half-brother of I36978, but this is impossible because, in this scenario, I36892 would not be related to I37330 and I36969, which he is.
- The nephew of I36978. This is very unlikely because, in this scenario, I36892 would be I36969's third-degree relative, and IBD sharing (433 cM) between the two individuals suggests that a fourth-degree relationship is much more likely.
- I36978's grandson and half-nephew of I36995 (Figure 4b): in order to resolve the endogamic loop observed in I36892, in this scenario, I36978 is also I36892's great-aunt, accommodating I36892's parents as first cousins. Given that I36978 and I36892 share the same mitochondrial haplogroup but do not share IBD in the X-chromosome, we can place I36978 as I36892's paternal grandmother, since, in this configuration, no X-chromosome would be transmitted, and as his I36892's maternal grandmother's sister, hence sharing the same mitochondrial haplogroup. Under this configuration, I36892 would be both a second+third-degree relative of I36978 (which aligns well with a relatedness coefficient of 0.39, 2200 cM in shared IBD and 124 cM in shared IBD2), and a third+fourth+sixth-degree relative of I36995 (which aligns well with a 0.21 relatedness coefficient and 1135 cM in shared IBD). The only minor inconsistencies in this scenario are: the relationship between I37330 and I36892, who would be double fifth-degree relatives (but who share 144 cM in

IBD), and the relationship between I36969 and I36892, who should be double fourth-degree relatives (but who share 434 cM in IBD).

Notably, individuals from the chariot burials in this cluster are fifth–sixth-degree relatives of I50780, the female from the chariot burial at Wetwang Village. As such, and like I50780 (Figure S20), they are also related to members of cluster 3 in the main cemetery at Wetwang Slack, especially I36969 who was buried in close proximity to most members of cluster 3, and whose closest (fifth–sixth-degree) relative outside of the chariots cluster is I37167 from cluster 3, with whom she shares the same mtDNA haplogroup (T2e1a1b).

##### *1.8. Connections between clusters 1, 2, 3, 4, 5 and 6*

Clusters 1, 2, 3, 4, 5 and 6 are all connected to one another through third-degree or closer relationships. Thus, Figure S39 displays clusters 3, 4 and 5 in a single tree because they connect in relatively simple ways that allow for easy visualization. Clusters 1, 2 and 6, however, connect with clusters 3, 4 and 5 in more complex configurations that makes it impossible to visualize them together in a unified two-dimensional tree and, as such, we indicate where clusters 1,2 and 6 connect with clusters 3, 4 and 5 in Figure S39. In Figure 3a we display clusters 1-6 in a unified image.

###### *Connections between clusters 3 and 4*

Clusters 3 and 4 connect via individual I31512, who is I30990's (cluster 3) maternal half-brother and I37126-I31024's (cluster 4) paternal half-brother.
They also connect via I36925's (cluster 4) unsampled father, who is I36888's (cluster 3) second-degree relative.

###### *Connections between clusters 3 and 5*

Clusters 3 and 5 connect via male infant I37136, who is the maternal grandson of I36966 and I37138 (cluster 5) and the paternal nephew of I36909 (cluster 3). I36909 cannot be I37136's paternal grandfather because I30980 and I37136 would be fourth-degree (rather than third-degree) relatives, and neither can they be paternal half-brothers because I30980 and I37136 would, in that case, be second-degree (not third-degree) relatives.
These two clusters also connect via:

-Female I39733 (cluster 5) and I31517 (cluster 3) who are third-degree relatives.

-Siblings I30962 and I31509 (cluster 3) and siblings I37138 and I31508 (cluster 5), who are third-degree relatives. Since I31512 (I30962 and I31509's father) is not related to siblings I37138 and I31508, the relationship has to proceed through I30962/I31509's mother. Furthermore, since I30907 (I37138 and I31508's maternal grandmother) is not related to I30962/I31509, the relationship must pass through I37138/I31508's father, who could be the maternal half-brother, maternal uncle or maternal grandfather of I30962 and I31509. A scenario in which I37138/I31508's father is the maternal half-brother of I30962 and I31509 is highly unlikely because siblings I30962 and I31509 would lie one generation above siblings I37138 and I31508. Based on the connection between clusters 3 and 5 via infant I37136, I30962 and I31509 lie, in fact, one generation below siblings I37138 and I31508, because I30962/I31509's father (I31512) has a maternal first cousin whose son (I36909's brother) reproduces with one of I37138's daughters. This configuration connecting both pairs of siblings is unambiguous and so, if I37138/I31508's father was the maternal half-brother of I30962 and I31509, I30962/I31509's mother would need to have reproduced with a man

(I31512) two generations below her. Similarly, in a scenario in which in I37138/I31508's father is the maternal uncle of I30962 and I31509, I30962-I31509's mother would need to have reproduced with a male one generation below her. As such, placing I37138/I31508's father as the maternal grandfather of I30962 and I31509 is far more plausible, for two reasons: first, in this scenario, I30962-I31509's mother can reproduce with a male from her own generation; and, more importantly, this configuration accommodates the observed X-chromosome sharing patterns between these individuals. Under this scenario, I31508 inherits one of her X-chromosomes directly from her father (I30962-I31509's maternal grandfather), which allows us to determine whether I30962 and I31509 inherited their maternal X-chromosomes from their maternal grandfather or grandmother. I30962 and I31508 share a short IBD segment at the beginning of the X-chromosome that I30962 must have inherited from his maternal grandfather, indicating that the rest of his X-chromosome came from his maternal grandmother. I31509 and I31508 share a large IBD segment (not overlapping with I30962-I31508's segment) that I31509 must have inherited from her maternal grandfather. Under this scenario, the regions of the X-chromosome outside those two segments must be shared in IBD between I30962 and I31509 because both siblings inherited them from their maternal grandmother. This is exactly what we observe (Figure S38). This scenario also aligns well with the observation that X-chromosome IBD breakpoints between I30962 and I31508 are different to those between I31509 and I31508 (because they derive from two distinct meiosis in I30962/I31509's mother), and that the IBD breakpoints observed between I30962 and I31509 will only be those in the I30962-I31508 and I31509-I31508 comparisons, which is exactly what we observe.

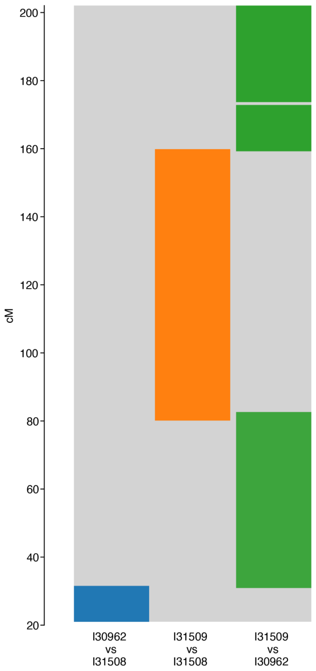


Figure S38. X-chromosome IBD segments shared between siblings I31509-I30962 and I31508 at Wetwang Slack.

##### ***Connections between cluster 1 and other clusters***

Cluster 1's connections with other clusters are described as follows:

-I37324 is a third-degree relative of I37135 (cluster 2) and a fourth-degree relative of I31010 (I37135's brother). I37324 must, therefore, be I37135's great-grandson.

-I36871's father is a second-degree relative of I31512's, I37126's and I31024's (cluster 4) father.

##### *Connections between cluster 2 and other clusters*

Cluster 2's connections with other clusters are described as follows:

- I37135 and I31010 are third-degree relatives of I37126 (cluster 4).
- I30986's and I30910's father is a third-degree relative of I36810 (cluster 5).
- I36962's and I36899's maternal grandmother's parents are I37138 and I36966 (cluster 5). Since I36962 and I36899 are both third-degree relatives of I36966 and I37138, they must be I36966's and I37138's descendants through I37136's maternal aunt. Indeed, 146C and 195C heteroplasmies observed in individuals I36962 and I36899 are also present in I36803's descendants (including I36966), support the suggestion that they are descended, through a strict maternal line, from I36803.
- I36773 is a third-degree relative of I31510 (cluster 3).

##### *Connections between cluster 6 and other clusters*

Cluster 6's connections with other clusters are described as follows:

- I36893, I36819 and I30971 are second-degree relatives of I36883's mother (cluster 5).
- I30902 is a third-degree relative and I37125 is a fourth-degree relative of siblings I37159, I36967 and I36965 (cluster 5). I30902 must be the siblings' great grandmother because all of the other possibilities (i.e. relationships through one of I30902's parents, I30902's sibling or I30902's half-sibling) result in fourth-degree relationships between the siblings and I30902's nieces and nephew (I30958, I36901 and I36841), not fifth-degree as observed. This is confirmed by the observed shared IBD segments between each of the siblings I37159-I36967-I36965 and each of the siblings I30958-I36901-I36841, in that they are, in all cases, a subset of those between I30902 and I37159-I36967-I36965; this is expected if I37159-I36967-I36965 are I30902's descendants. Furthermore, the connection between I30902 and I37159, I36967 and I36965 must be through their mother and through I30902's son, because this is the only configuration which allows I36965 and I30902 to share IBD in the X-chromosome but not share a mitochondrial haplogroup.
- I36989's mother is a first-degree relative of I36815 (cluster 5). Given that I36989 and I30985 are fourth-degree relatives, I36815 must be I36989's maternal grandmother because the other possibilities (such as I36815 as I36989's maternal aunt or maternal half-sister) would result in I30985 and I36989 having third-degree or second-degree (not fourth-degree) relationships.
- Siblings I37161 and I36886 are second-degree relatives of I36963's father (cluster 5).
- Siblings I37161 and I36886 are the great uncle and aunt (respectively) of I30970 (cluster 3)
- I30994 is a third-degree relative of I31009 (cluster 5).

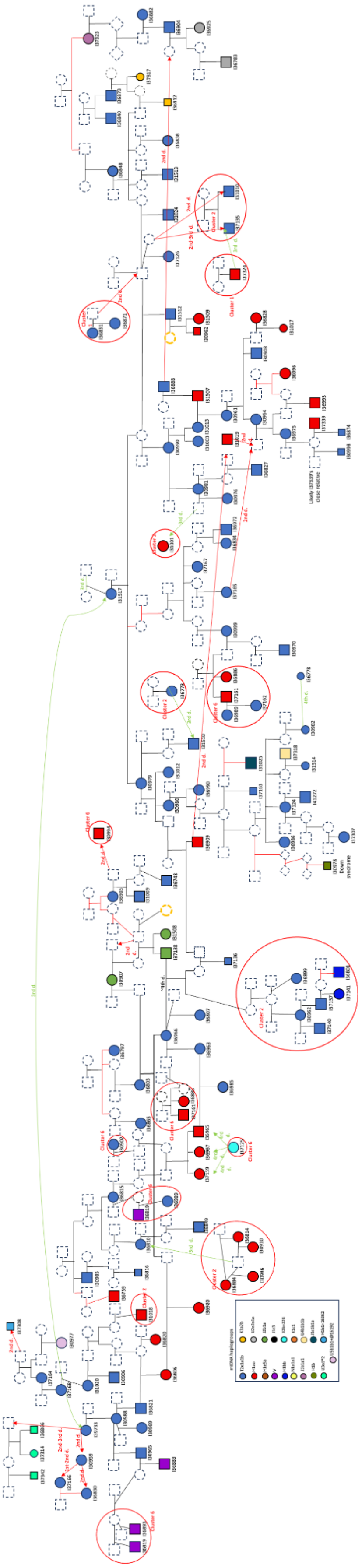

Figure S39. Combined pedigree for cluster 3, 4 and 5 at Wetwang Slack.

#### ***2. Pocklington and Melton 1***

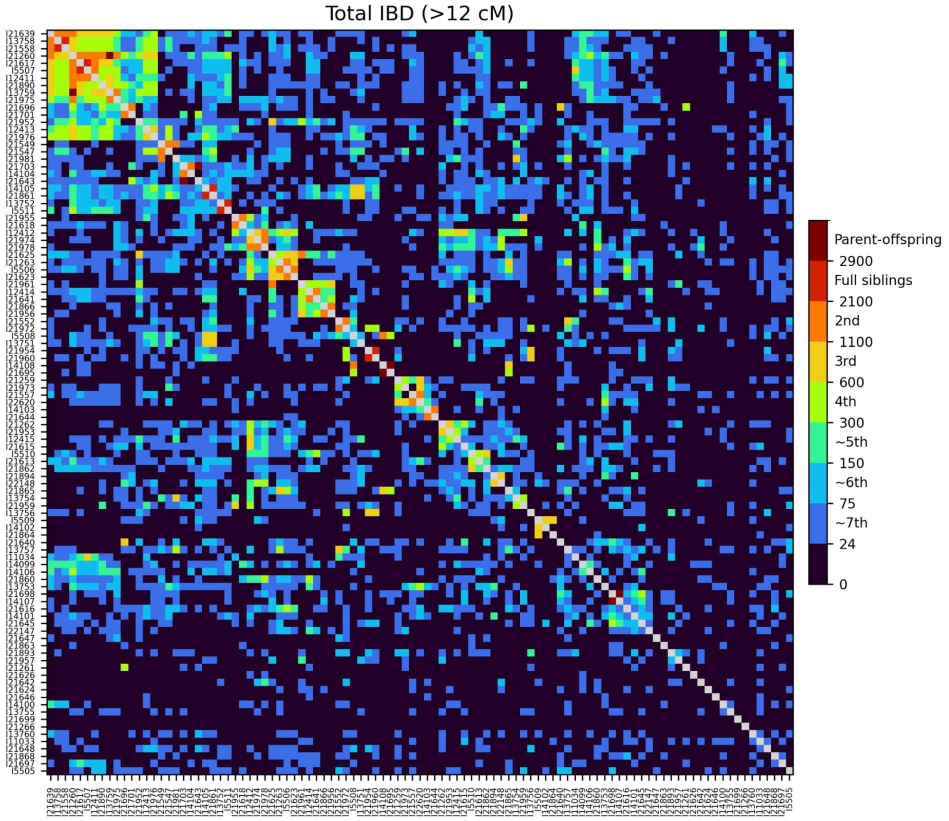


Figure S40. Heatmap of IBD sharing between pairs of individuals at Pocklington.

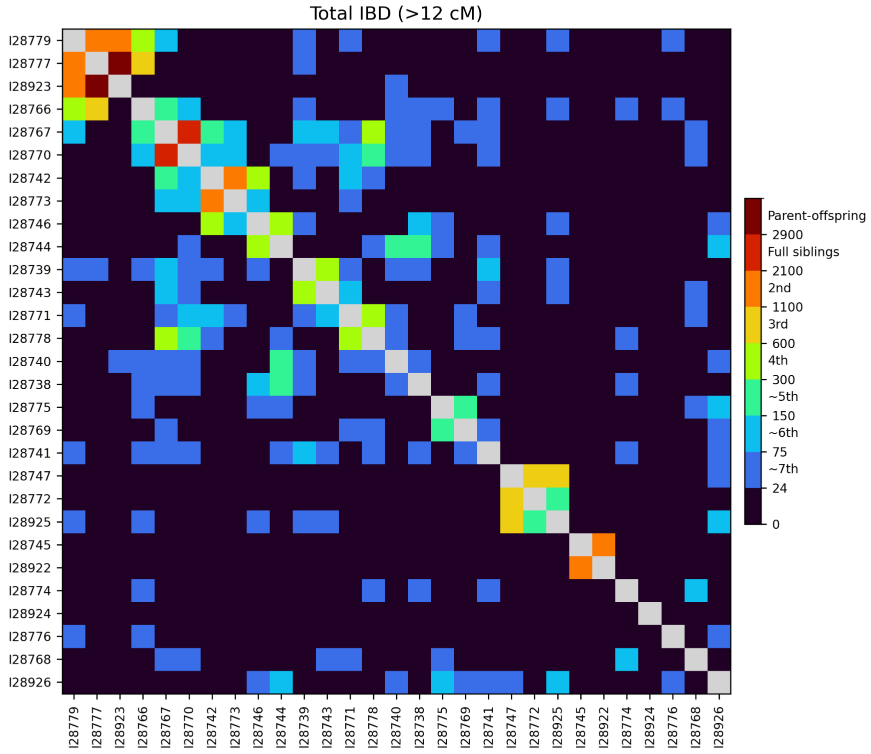


Figure S41. Heatmap of IBD sharing between pairs of individuals at Melton 1.

At both Pocklington and Melton 1, we detect many intra-site pairs of close relatives (Figure S40 and Figure S41), but the relative lack of individuals with several first- and second-degree relationships prevents us from reconstructing large family pedigrees like those at Wetwang Slack, with one exception: individual I21260's relatives at Pocklington (Figure S43), as detailed below.

Adult female I21260 is a second-degree relative of I21975, I21890, I21639, I21617, I5507 and I12411, and a third-degree relative of I21558 and I13758. I13759 is I21260's son, and both are second-degree relatives of I21975; they must therefore be I21975's maternal grandmother and uncle (respectively). I21558 and I13758 are siblings and second-degree relatives of I21639. Individual I21639 must be their maternal uncle, because this scenario allows for the presence of ~400 cM of shared segments in IBD0 between the siblings and at the same time in IBD1 between each sibling and I21639. Such regions are only expected for a pair of siblings and their uncle/aunt. For the same reason, I21260 is the maternal aunt of siblings I21617 and I5507. I12411 is a second-degree relative of I21260, I21617 and I5507. As such, she must be I21617 and I5507's maternal half-sister. Uncertainties remain, however, regarding the position of I21890 and I21639 with respect to I21260. Individual I21890 could be I21260's niece through her unsampled brother, or I21260's paternal aunt. I21890 and I21260 cannot be paternal half-sisters or paternal grandmother-granddaughter because they do not share IBD segments across the entire X-chromosome. Furthermore, X-chromosome patterns allow us to discard one of the other suggested possibilities for the position of I21890 in the tree. If I21890 were I21260's paternal aunt, then all IBD segments shared between I21890 and I21617, I5507 or I12411 must also be shared also between I21890 and I21260. This is because these segments would derive from I21890's brother and I21617-I5507-I12411's maternal grandfather, who would have passed his entire X-chromosome to I21260. However, if I21260 were I21890's paternal aunt, segments shared between I21890 and I21617, I5507 or I12411 need not be shared between I21890 and I21260, because they would be derived from I21260's mother, who may well have passed one of her X-chromosomes to I21890's father and to I21617-I5507-I12411's mother, and the other one to I21260. Since we observe several of this type of segments (Figure S42), shared between I21890 and either I21617, I5507 or I12411, but not shared between I21890 and I21260, I21890 cannot be I21260's paternal aunt and must be I21260's niece through I21260's unsampled brother.

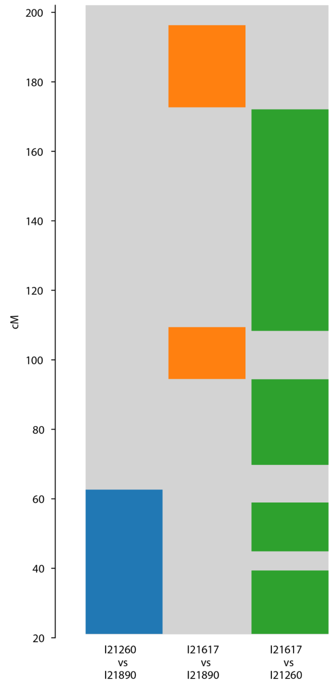


Figure S42. X-chromosome IBD segments for I21260's, I21890, and I21617 at Pocklington.

The only remaining ambiguity in the pedigree is the position of I21639, who could be I21260's maternal uncle, nephew or half-brother. The number of shared segments allows us to strongly disfavour the scenario with I21639 as I21260's half-brother, due to the following reasoning: all unambiguous third-degree relationships in the tree (I13759-I21617, I13759-I5507, I12411-I13759, I21890-I12411, I21890-I5507, I21890-I21617 and I21890-I13759), progress through a pair of full-siblings, sharing 22, 19, 28, 20, 19, 20 and 19 IBD segments longer than 12 cM, respectively. If I21639 were I21260's half-brother, we would expect I21639 to share fewer IBD segments with his five third-degree relatives than those reported above for the unambiguous third-degree relationships, since the relationship would not progress through a pair of full-siblings. In fact, these individuals share 22, 27, 30, 23 and 28 IBD segments longer than 12 cM, similar to (and, in some cases, even higher than) the unambiguous third-degree relationships progressing through a pair of siblings. This suggests that the relationship between I21639 and I21260 must also progress through a pair of full-siblings, in a scenario whereby I21639 is either I21260's maternal uncle or nephew.

The relationship between I21639 and other individuals strongly argues against the scenario in which I21639 is I21260's uncle. Individual I14100 shares nine IBD segments longer than 12 cM, and a total of 230 cM, with I21639. I14100 must be either I21639's relative through I21639's unsampled sibling or a relative via I21639's parents, but not through a I21639's descendant because firstly, I14100 shares approximately 100 cM in IBD with I21639's nephew (I13758) and niece (I21558), which suggests a sixth-degree relationship, i.e. only one-degree lower than the relationship between I21639 and I14100, not the expected two-degrees if I14100 were a descendant of I21639. Secondly, and more importantly, I14100 shares IBD segments with I13758 and I21558 that are not shared between I21639 and I14100, and this is not possible if I14100 is I21639's descendant. Given that I14100 is related to I21639 via I21639's parents or siblings, if I21639 is the maternal uncle of I21260, then I14100 should also be related to I21260, to the order of at least one degree more distant than I14100 is related to I21639 (i.e. sixth degree), or potentially more closely if the connection is through I21260's mother. The absence of shared IBD segments between I14100 and I21260 makes this scenario unlikely, as sixth-degree relatives are generally expected to share some detectable IBD.

In summary, the remaining likely configuration for I21639, is that he is I21260's nephew through an unsampled sister of I21260 (Figure S43), which accommodates the fact that I21639 has paternal relatives not shared with his maternal aunt (I21260).

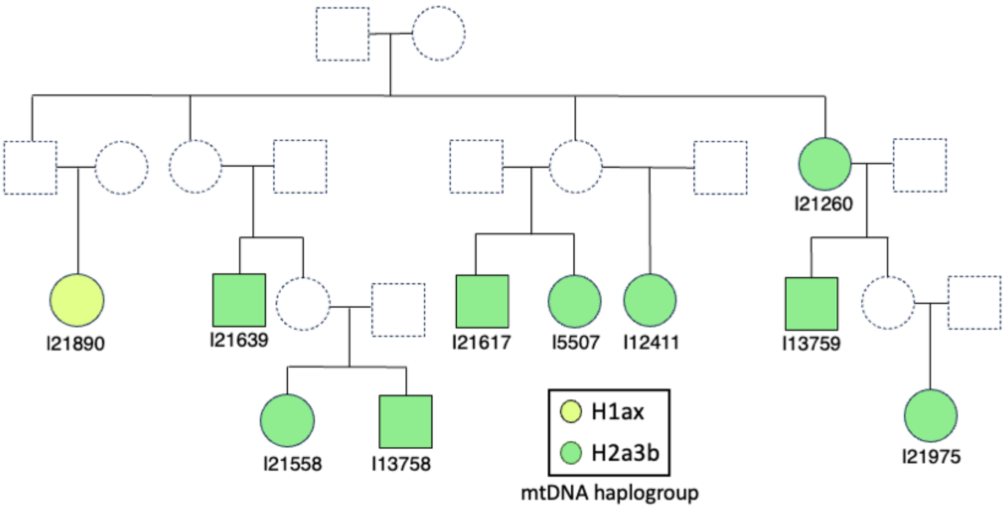


Figure S43. Reconstructed pedigree for I21260's relatives at Pocklington.

#### *3. Inter-site pedigrees*

There are widespread inter-site genetic links between the three main cemeteries (Wetwang Slack, Pocklington and Melton 1; Figure S12), but most are too distant to allow for reconstruction of the pedigrees connecting these individuals. The closest links between each pair of cemeteries are two sets of third-degree relatives and one fourth-degree relationship, the configurations of which are explored below.

##### 3.1. Connection between I39724 (Wetwang Slack) and I28740 (Melton 1)

I39724 is a 35–45 year-old male from Wetwang Slack who belongs to the second most frequent mitochondrial lineage (H1ao) at the site. He shares 1061 cM in IBD with I28740, a 9–11 year-old girl buried at Melton 1 who belongs to the dominant mitochondrial lineage (H3q1) there. They are most likely third-degree relatives, although a second-degree relationship cannot be completely ruled out.

I39724's closest relatives at Wetwang Slack can be confidently placed within a pedigree (Extended Data Figure 8b). I36768 and I36754 are I39724's mother and maternal aunt, respectively, while I36756 is I36754's granddaughter via an unsampled daughter. I36761 is a second-degree relative of I36756 which must progress through the paternal line, as he is not related to I36756's maternal relatives. I36761 could be I36756's paternal grandfather, half-brother or uncle, although the first two scenarios are more likely based on the low number of shared IBD segments (n=19) between them.

I39724's relatives do not appear to be related to the large group (n=195) of sampled Wetwang Slack individuals connected by third-degree or closer relationships (clusters 1–6). They are,

however, related to this group via a fourth–fifth-degree relationship between I36756 and I37329 from cluster 6 at Wetwang Slack.
I39724's maternal family is not related to I28740, which means that I28740 must be a second-degree relative of I39724's father, or a first-degree relative of I39724's father if I39724 and I28740 are second-degree relatives. Assuming that I28740 and I39724's father are second-degree relatives, and since I28740 is a (probably pre-pubescent) non-adult, I39724's father could be: I28740's grandfather (through a reproductive partner other than I39724's mother), I28740's uncle, I28740's half-brother or I28740's nephew. I28740's closest (approximately fifth-degree) relative at Melton 1 is infant I28744, who is also related to I39724's father via an approximately fourth-degree relative.

##### 3.2. Connection between I30994 (Wetwang Slack) and I21975 (Pocklington)

Adult male I30994 from Wetwang Slack (cluster 6) is a third-degree relative of adult female I21975 from Pocklington, who is, herself, part of the reconstructed pedigree at Pocklington (Figure S43). I30994 from Wetwang Slack is also a third-degree relative of female I31009 from Wetwang Slack (cluster 5), which helps in the reconstruction of I30994's relationship with I21975.
Three pieces of evidence allow us to narrow down the number of possible tree configurations:
- I30994 is a third-degree relative of both I21975 from Pocklington and I31009 from Wetwang Slack, but I21975 and I31009 do not share any IBD segments, indicating that I21975 and I31009 must be related to I30994 through I30994's parents. In all the other scenarios, for instance, in which one of I21975 and I31009 is related to I30994 through I30994's father or mother and the other is related to I30994 through one of I30994's descendants (or through a descendant of I30994's unsampled sibling), I21975 and I31009 would be sixth-degree relatives or closer, and we would expect detectable IBD sharing, which we do not identify.
-I30994 is not related to I36905 (I31009's mother) but I30994 is related to I36748 (I31009's nephew through a sister), indicating that the relationship between I30994 and I31009 progressed through I31009's father, who is I30994's second-degree relative.
- I21975's close maternal relatives (I13759 and I21260) are not related to I30994, which means that the relationship between I30994 and I21975 is transmitted through I21975's father, who is I30994's second-degree relative.
Based on this evidence so far, the fathers of I21975 and I31009 are both second-degree relatives of I30994: one each through the maternal and paternal lines. The fathers of I21975 and I31009 could, therefore, be half-brothers, grandfathers or uncles of I30994. The number of IBD segments (n=15) shared between I30994 and I21975 is, however, rather low for third-degree relatives, which favours the half-brother and grandfather scenarios over those that place I21975 and I31009 as uncles, because the genetic transmission need not pass through a pair of full-siblings. I30994 shares neither a mitochondrial haplogroup nor any X-chromosome IBD segment with I21975 or I31009, precluding the assignment of these individuals as maternal and/or paternal relatives. If I21975's father were I30994's maternal half-brother or grandfather, only two or one meioses respectively would separate the X-chromosomes of I21975 and I30994. This would likely result in shared X-chromosome IBD segments between I21975 and I30994. Since this X-chromosome IBD sharing is absent, it is more likely that I21975's father is a paternal relative and I31009's father is a maternal relative of I30994. In this scenario, three, two or four meioses would separate the X-chromosomes of I36748 and I30994 (depending on whether I31009's father is I30994's half-brother, grandfather or uncle). The more plausible scenario is one in which I31009's father is the uncle

of I30994, as it maximizes the probability that I36748 and I30994 do not share X-chromosomal IBD. Extended Data Figure 8a displays the most likely tree connecting these individuals, with red lines indicating more ambiguous configurations.

##### 3.3. Connection between I28775 (Melton 1) and I21981 (Pocklington)

The closest genetic link between Pocklington and Melton 1 is a fourth-degree relationship between adult male I21981 (Pocklington) and I28775, a 10–12 year-old boy from Melton 1 (Extended Data Figure 8c). Both carry Y-chromosome haplogroup G-CTS4803, a rare lineage present only in these two individuals within the entire Arras dataset. This strongly suggests a recent male-mediated genetic connection between the two sites. I21981 also had an infant granddaughter (I21549) in common with I21547, who was herself also buried at Pocklington. I21549 shares mutually exclusive IBD segments with I21547 and I21981, indicating that I21547 and I21981 are her grandparents from the same (paternal) side.

**Inter-site connections**

Beyond the close inter-site relationships that could be fitted into pedigrees, we wanted to identify individuals who, as well as having many intra-site genetic connections, stood out as having more between-site genetic connections to the other main sites (Wetwang Slack, Pocklington and Melton 1). For each individual, we calculated: (i) the mean amount of IBD shared with individuals from the same and other sites (sum_IBD>8 + sum_IBD2), and (ii) the proportion of relatives at each site. Relatives were defined as individuals sharing more than two IBD segments of >8 cM and a total of >24 cM in IBD in total. Mean IBD is strongly influenced by close genetic ties involving long shared segments. Meanwhile, the proportion of relatives is less sensitive to single close relationships but does not capture the intensity of relatedness.

<u>*Wetwang Slack-Pocklington*</u>

The most striking pattern is the generally high genetic connectivity (i.e. beyond those individuals who can be fitted into a pedigree) between Wetwang Slack and Pocklington. Individuals from both sites show overlapping distributions in the proportion of relatives at Wetwang Slack (Figure S44), with nearly all Pocklington individuals and the vast majority of Wetwang Slack individuals falling between 0–21% of Wetwang Slack relatives. This indicates that most Pocklington individuals have multiple (mostly distant) relatives at Wetwang Slack. Moreover, the proportion of relatives at Wetwang Slack is positively correlated with the proportion of relatives at Pocklington, both among Wetwang Slack individuals (Spearman's $\rho = 0.622$, $P = 6.27 \times 10^{-42}$) and Pocklington individuals ($\rho = 0.7$, $P = 2.49 \times 10^{-16}$). In other words, an individual who has lots of relatives at Wetwang Slack, is also likely to have lots of relatives at Pocklington.

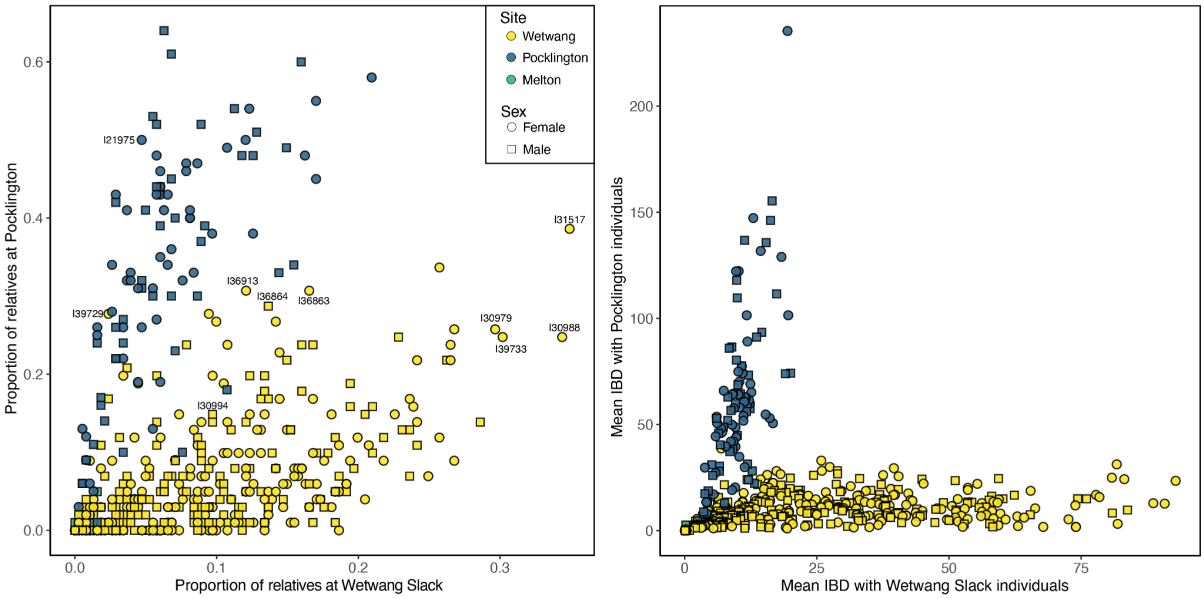


Figure S44. Comparison of individual-level relatedness to Wetwang Slack and Pocklington burial communities. **a)** For each individual, the proportion of relatives at Wetwang Slack is plotted against the proportion of relatives at Pocklington. The proportion represents the number of related individuals divided by the total number of individuals sampled at that site. Relatives are defined as pairs sharing >2 IBD segments of >8 cM and >24 cM total IBD. **b)** For each individual, the mean IBD sharing (sum_IBD>8 + sum_IBD2) with Wetwang Slack is plotted against the mean IBD with Pocklington individuals.

Wetwang Slack individuals with the highest intra-site proportion of relatives also tend to have the highest proportion of relatives at Pocklington (Figure S44). Female I31517, along with her third-degree relative I39733 and their respective daughters (I30979 and I30988), show particularly high proportion of relatives at Pocklington (Figure S44). Notably, I31517 sits at the top of cluster 3 (Figure S20), which spans nine generations, and is the Wetwang Slack individual with the largest number of descendants buried at the site (n=37), while her third-degree relative I39733 sits at the top of a pedigree spanning five generations (Figure S39). This likely reflects their position in the earliest generations of the Wetwang Slack pedigree. If most Wetwang Slack–Pocklington connections pre-date the main use of both cemeteries, individuals in the early generations of the Wetwang Slack cemetery would be expected to show stronger links to Pocklington. Importantly, the mean IBD sharing of these females is substantially higher within Wetwang Slack (~80 cM) than with Pocklington (~30 cM) (Figure S44), indicating that their closest ties remain within their site of burial.

Wetwang Slack individuals with a very low proportion of intra-site relatives also show a low proportion of Pocklington relatives. Among the 68 Wetwang Slack individuals with <2% intra-site relatives, only three exceed 5% of relatives at Pocklington. This pattern suggests the coexistence of two broadly connected communities at Wetwang Slack and Pocklington alongside a subset of Wetwang Slack individuals with limited connections to both sites.

Several Wetwang Slack individuals display average intra-site proportion of relatives but display elevated connections with Pocklington:
-Individuals I36863, I36913, I36864, and I39729 rank (at third, fourth, fifth and sixth, respectively) among the Wetwang Slack individuals with the highest proportion of Pocklington relatives. I39729 is also the Wetwang Slack individual with the highest mean IBD value shared with the community at Pocklington (Figure S44). All belong to haplogroup J1c9, the third most

frequent haplogroup at Pocklington (13%). This evidence is consistent with recent female-mediated mobility from Pocklington to Wetwang Slack, particularly I39729, whose 12 closest relatives were buried at Pocklington, including an approximately fifth-degree relative sharing 63% of the X chromosome in IBD.

-The closest Wetwang Slack–Pocklington link is a third-degree relationship between male I30994 (Wetwang Slack) and female I21975 (Pocklington). Their fathers are first-degree relatives, most likely father and son (Extended Data Figure 8a). I30994's close maternal relatives, his descendants and his reproductive partner's close relatives are all buried at Wetwang Slack, whereas I21975's close maternal relatives are buried at Pocklington. The proportion of relatives of each individual at both sites (Figure S44) can inform us about the possible direction of movement. If I30994's father was born among the Pocklington community and moved to Wetwang Slack, we would expect I30994 to stand out among Wetwang Slack individuals in terms of his proportion of Pocklington relatives, since his paternal family would come from Pocklington. On the other hand, we would expect I21975 to fall within the average range in terms of her proportion of relatives at Wetwang Slack, since only I30994 and his descendants would be her relatives (not those higher up the pedigree). Among the Pocklington individuals, I21975 lies at the 41st percentile for the proportion of Wetwang Slack relatives, whereas among Wetwang Slack individuals, I30994 lies at the 88th percentile for the proportion of Pocklington relatives. Despite substantial background relatedness between sites, this asymmetry suggests recent male movement from Pocklington to Wetwang Slack.

_Wetwang Slack-Melton 1_

The closest connection between these sites is a third-degree relationship between an adult male (I39724; Wetwang Slack) and a 9–11 year-old female (I28740; Melton 1). I39724's maternal relatives are buried at Wetwang Slack. I39724's father could be related to I28740 as her grandfather (through union with a different reproductive partner to I39724's mother), or he could be her uncle, half-brother, or nephew (Extended Data Figure 8b). I39724 is the Wetwang Slack individual with the highest proportion of Melton 1 relatives, whereas I28740 lies within the 64th percentile of the proportion of Wetwang Slack relatives within the Melton 1 burial community (Figure S45). This asymmetry again suggests male-mediated movement, potentially involving I39724's father relocating from Melton 1 to Wetwang Slack.

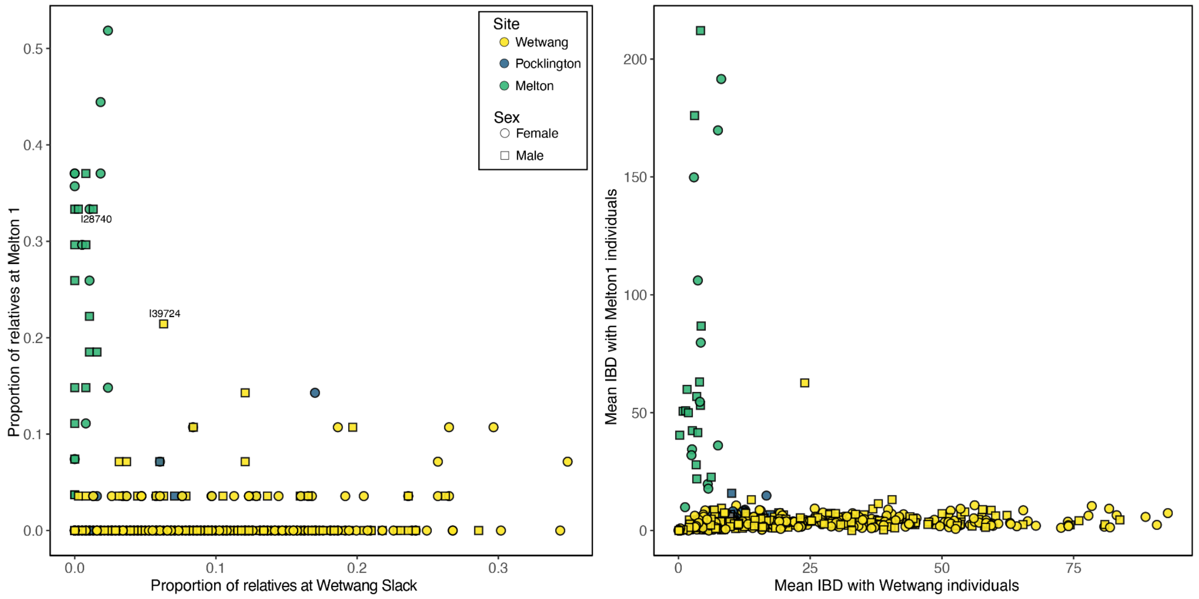

Figure S45. Comparison of individual-level relatedness to Melton 1 and Wetwang Slack. **a)** For each individual, the proportion of relatives at Melton 1 is plotted against the proportion of relatives at Wetwang Slack. The proportion represents the number of related individuals divided by the total number of individuals sampled at that site. Relatives are defined as pairs sharing >2 IBD segments of >8 cM and >24 cM total IBD. **b)** For each individual, mean IBD sharing (sum_IBD>8 + sum_IBD2) with Wetwang Slack is plotted against mean IBD sharing with Melton 1 individuals.

#### *Pocklington-Melton 1*

I28775, a 10–12-year-old boy from Melton 1—who has a fourth-degree relationship with I21981 from Pocklington (Extended Data Figure 8b)—has the highest proportion of Pocklington relatives among all individuals from his burial community (Figure S46).

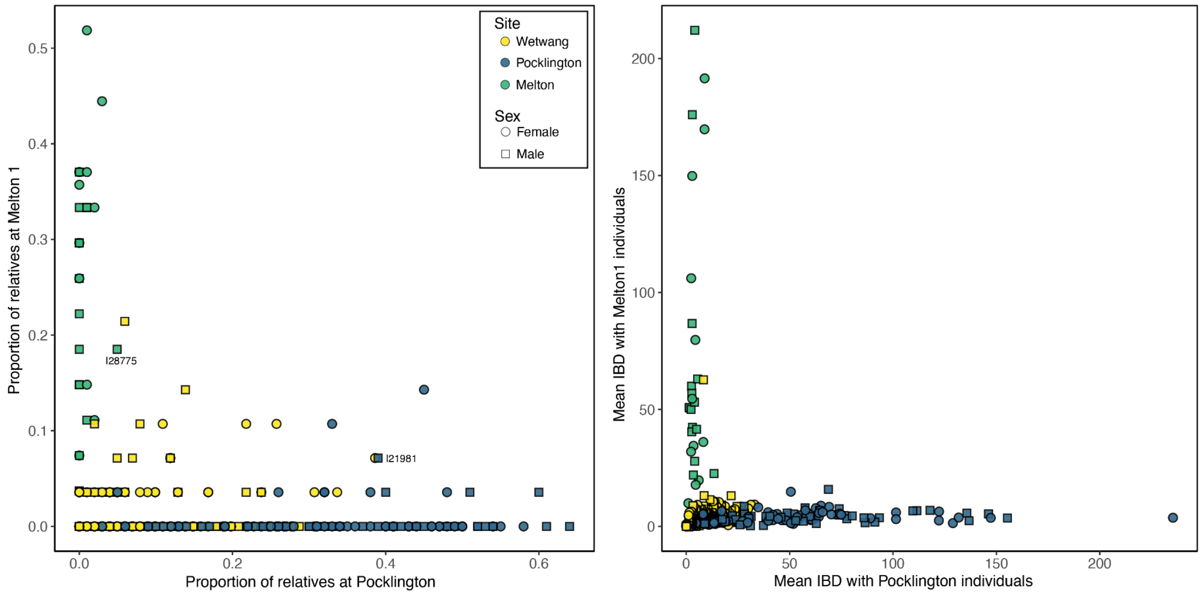


Figure S46. Comparison of individual-level relatedness to Melton 1 and Pocklington. **a)** For each individual, the proportion of relatives at Melton 1 is plotted against the proportion of relatives at Pocklington. The proportion represents the number of related individuals divided by the total number of individuals sampled at that site. Relatives are defined as pairs sharing >2 IBD segments of >8 cM and >24 cM total IBD. **b)** For each individual, the mean IBD sharing (sum_IBD>8 + sum_IBD2) with Melton 1 is plotted against the mean IBD with Pocklington individuals.

#### SI 6. Intra-site patterns of biological relatedness at the three main sites under study

In this section, we explore the correlation of biological relatedness with other traits at the three main sites (Wetwang Slack, Pocklington and Melton 1).

For each individual, we again calculated the proportion of relatives within the same site. Two individuals were considered related if they shared more than two IBD segments of >8 cM and a total of >24 cM in IBD. Pairs were classified as first- or second-degree relatives if their relatedness coefficient exceeded 0.20.
The median proportion of intra-site relatives was 0.26 at Melton 1 and 0.33 at Pocklington (Figure S47). These values indicate that individuals buried at Melton 1 and Pocklington were typically embedded within extended kin groups, with a substantial fraction of biological relatives interred at the same site. In contrast, Wetwang Slack shows a lower overall median proportion of relatives (0.08); which is examined in more granular detail below.

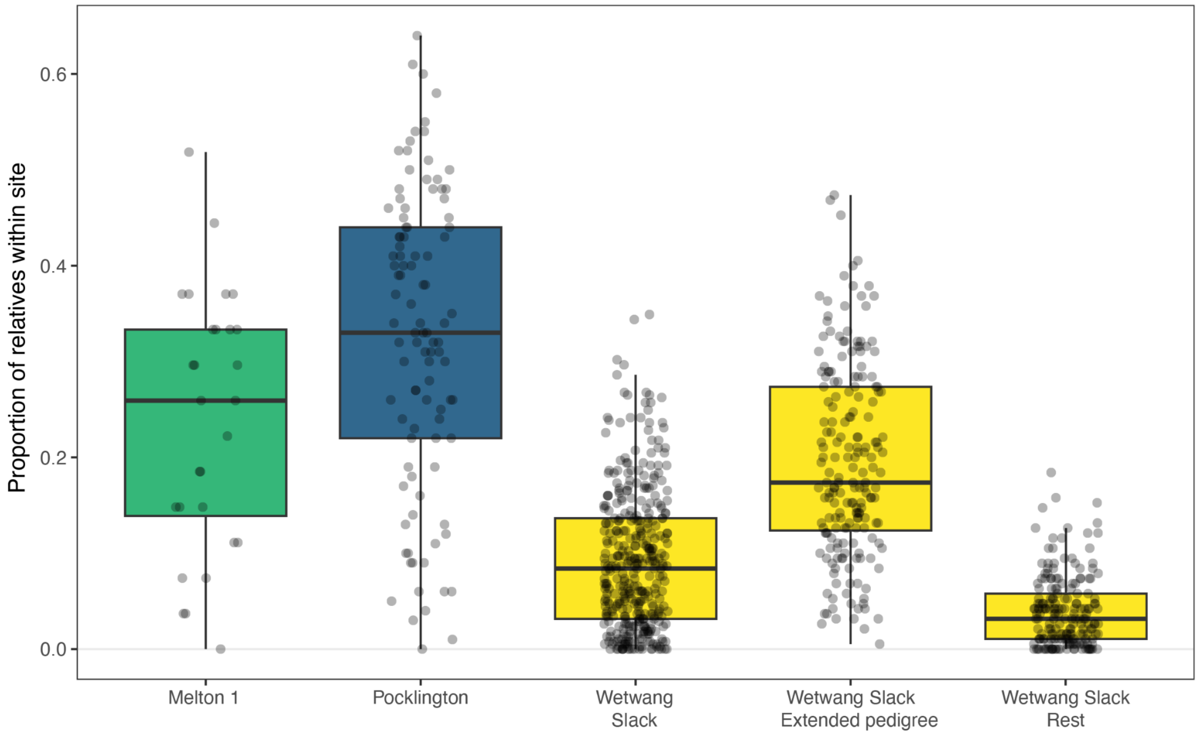


Figure S47. Proportion of intra-site relatives per individual at Melton 1, Pocklington and Wetwang Slack. Wetwang Slack is shown both overall and subdivided into individuals who belong to the large reconstructed pedigree (n=195) and those who do not. Relatives are defined as pairs sharing >2 IBD segments longer than 8 cM and >24 cM in total IBD. The proportion of intra-site relatives is calculated by dividing the number of related individuals by the total number of individuals sampled at the same site.

Pedigree reconstruction at Wetwang Slack revealed a large family comprising 195 individuals connected through third-degree or closer relationships. We therefore divided individuals at this site according to whether or not they belonged to this extended pedigree and recalculated relatedness proportions within each group. The large family displays a pattern comparable to that observed at Pocklington and Melton 1, with a median proportion of relatives within-group of 0.17 (Figure S47). By contrast, individuals outside this extended pedigree show very low proportions of within-group relatives (median = 0.03).

As outlined in SI4, despite the high proportion of related individuals at Melton 1 and Pocklington, we were unable to reconstruct multi-generational pedigrees comparable to those at Wetwang Slack. This difference reflects the distribution of close genetic links. At Wetwang Slack, many individuals have more than two first- or second-degree relatives (Figure S48), providing the connections needed to reconstruct extended pedigrees spanning several generations at this site. By contrast, at Melton 1 and Pocklington, most individuals have at most one first- or second-degree relative, limiting pedigree reconstruction despite the overall high prevalence of biological relatedness.

Although plausibly a real archaeological difference between the sites, we must also bear in mind that this contrast could also be strongly influenced by sample size. When we randomly downsample Wetwang Slack (n=390) to match the sample size at Pocklington (n=99), we recover a mean of 23±6 first- or second-degree relative pairs (versus 333 pairs of first- or second-degree relatives across the full dataset), close to the 37 observed at Pocklington. Thus, if Wetwang Slack had been only partially excavated, or if only a quarter of individuals had been sampled, we would not have been able to reconstruct the large multi-generational pedigrees available in the full dataset.

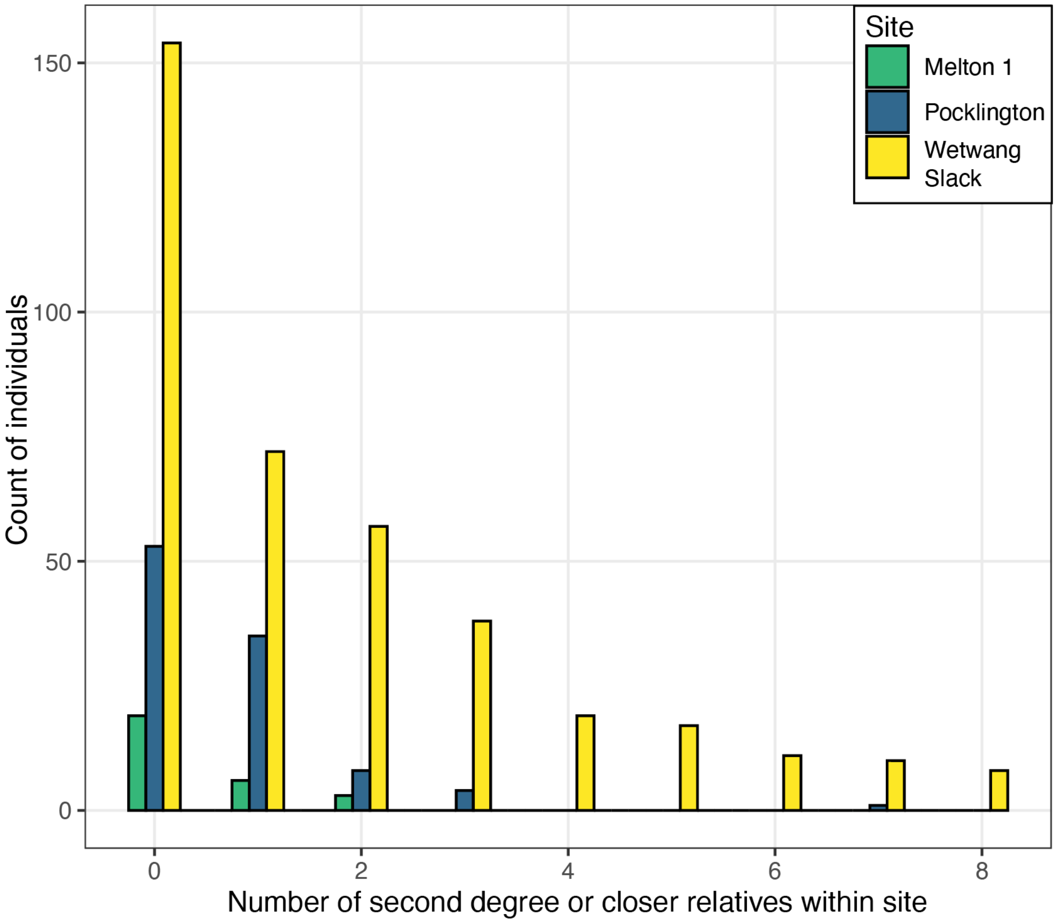


Figure S48. Histogram of the number of second-degree or closer relatives (i.e. those with a relatedness coefficient >0.20) per individual at Melton 1, Pocklington and Wetwang Slack.

*Sex-specific relatedness patterns at Wetwang Slack and Pocklington*

To assess sex-specific patterns in intra-site biological relatedness among individuals in whom social practices such as exogamy, residence and reproductive affiliation would have had time to be expressed, we calculated, for each adult individual, the mean amount of IBD shared with adult males, adult females and all adults from the same site. IBD sharing between individuals, was calculated using the sum of the genomic length in IBD segments of >8 cM (sum_IBD>8) and the total length of IBD2 segments (Supplementary Table 4), which accounts for the fact that two individuals can share IBD on both homologous chromosomes in IBD2 regions. To assess whether the observed differences could arise by chance, we performed a permutation test (n=100,000) in which sex labels were randomly reassigned among adults while preserving the number of individuals assigned to each sex-based category. For each permutation, per-individual mean IBD values were recalculated using the permuted labels, distributions of intra-site relatedness among adult females and distributions of intra-site relatedness among adult males were reconstructed, and the difference between medians of both distributions was recomputed. The same procedure was, then, applied for the comparison between adult males with all adults and adult females with all adults. This procedure generates a null distribution (Figure S50) under the hypothesis that sex is unrelated to patterns of IBD sharing, and allows for the computation of empirical two-sided p-values. Given that the structure of pairwise IBD sharing at the site is preserved, the resulting null distribution also inherently reflects differences in male vs female group sizes, sex-based heterogeneity in individual relatedness, and the underlying network structure of IBD sharing.

At Wetwang Slack, the median IBD shared between adult females and other adult females was 28.9 cM, whereas the corresponding value for adult males was 15.8 cM (Figure S49). The observed difference in median IBD sharing was rarely reproduced under permutation (P-value = 0.036) (Figure S50), indicating that adult females were more closely related to one another than adult males were to each other. The difference in IBD sharing between adult females and all adults (median = 29.13) versus adult males with all adults (median = 18.72) was also statistically significant (P-value = 0.041), indicating that females tend to share more IBD across the entire adult burial population than do males. These patterns persisted, even after excluding first-degree relationships (P-values 0.020-0.029), indicating that the observed sex-based differences are not driven solely by close kin relationships but reflect broader patterns of relatedness within the community.

At Pocklington, adult females shared more IBD with other adult females (median = 48.9 cM) than adult males did with other adult males (median = 42.1 cM) (Figure S49), but this difference was not statistically significant (permutation P = 0.58) (Figure S50). At Melton 1, the small sample size precluded meaningful statistical testing.

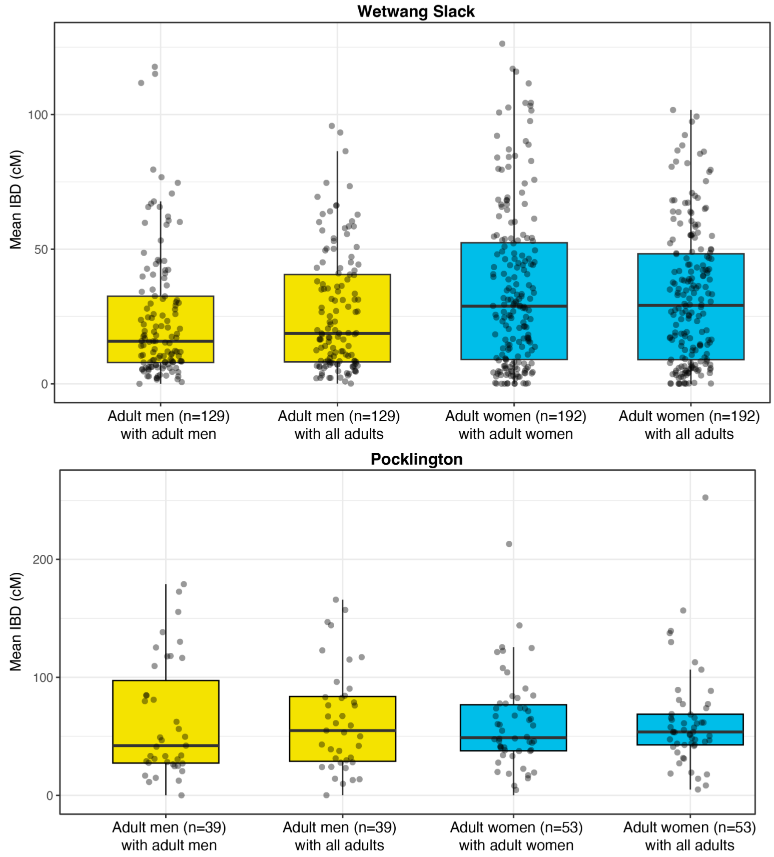


Figure S49. Distribution of per-individual intra-site mean IBD sharing at Wetwang Slack and Pocklington. For each adult individual, the mean genomic length shared in IBD (sum_IBD>8 cM + sum_IBD2) with adult individuals of the same sex, of the opposite sex and of both sexes was computed.

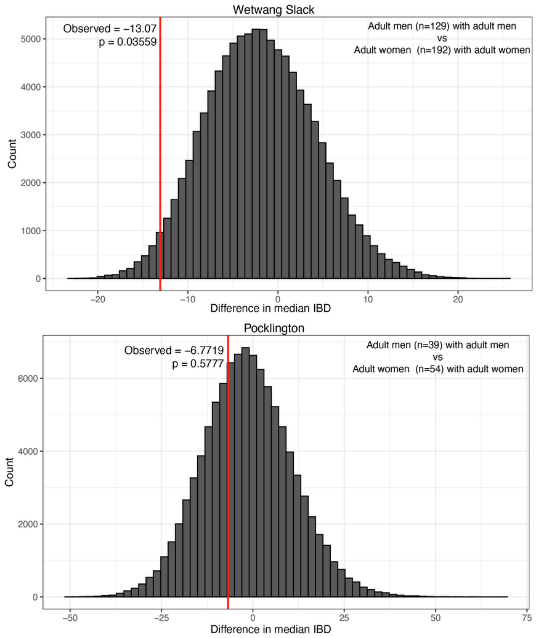


Figure S50. Permutation-based null distribution of the difference in median of same-sex IBD sharing between adult males and adult females at Wetwang Slack. Sex labels were randomly reassigned among adults (n=100,000 permutations) while keeping pairwise IBD values fixed. The vertical line indicates the observed difference. The empirical two-sided P-value corresponds to the proportion of permuted differences which matched or exceeded the observed value.

*Relatedness by burial type*

To investigate whether patterns of genetic relatedness vary according to burial type within each site, we compared the proportion of intra-site relatives across the different burial types—as assigned in the original excavation reports—represented (Figure S51). At Wetwang Slack, the median value of the proportion of intra-site relatives was 0.1 for primary burials in barrows, compared to values ranging between 0.04 and 0.06 for secondary burials on barrow platforms, flat graves, and ditch burials. After correcting for multiple testing, the differences in median values were found to be statistically significant for comparisons between individuals afforded primary burials in barrows with individuals buried in both barrow ditches and flat graves (Table S1). Although the comparison between primary burials (n=209) and secondary burials in barrows (n=23) showed a difference in median values (0.042) comparable to that observed (0.045) between primary burials in barrows and flat graves (n=47), the difference was not statistically significant, likely due to the relatively small sample size of secondary burials in barrows.

At Pocklington (unlike Wetwang Slack) barrows take both square/rectangular and circular forms. Individuals buried in square or rectangular barrows displayed the highest median proportion of intra-site relatives (0.38) (Figure S51), while flat graves showed the lowest value (0.26). Meanwhile, the difference between median proportions of relatives for individuals interred as primary burials in barrows (n=60; circular, square and rectangular; 0.36) and other burial types (n=41; flat grave and secondary barrow burials; 0.32) was not statistically significant (Table S1). This analysis indicates that, at Wetwang Slack and Pocklington, individuals interred as primary burials in barrows had substantially more intra-site relatives than individuals in other burial types, but only at Wetwang Slack was this difference statistically significant, possibly due to lower sample sizes at Pocklington.

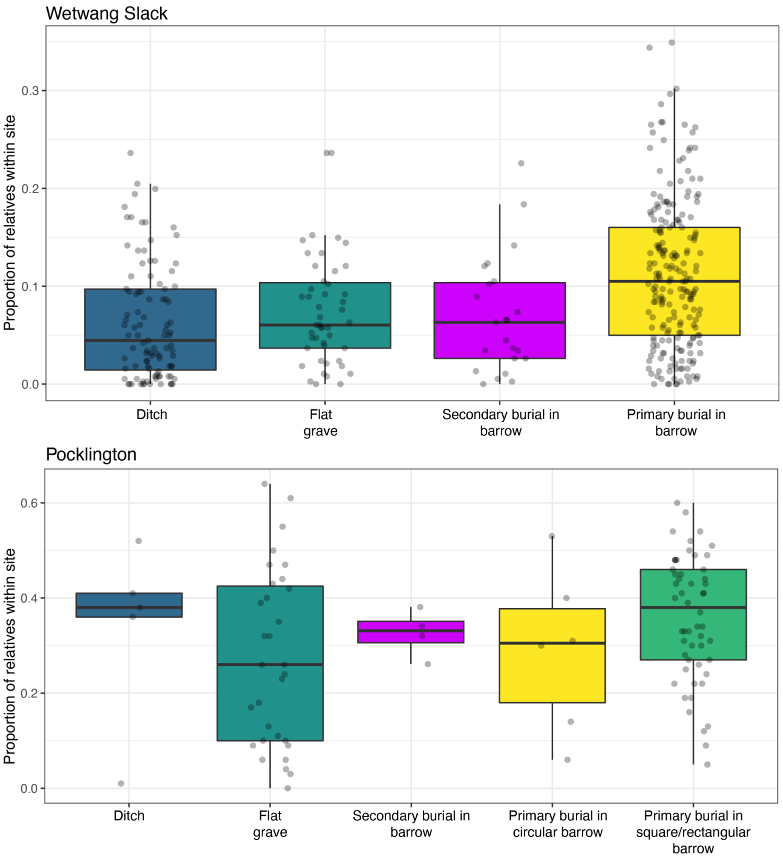

Figure S51. Proportion of intra-site relatives per individual at Wetwang Slack and Pocklington, grouped by burial type.

Table S1. Comparing the proportion of intra-site relatives between different burial types at Wetwang Slack and Pocklington.

| Site | Burial type 1 | Burial type 2 | n1 | n2 | median burial type 1 | median burial type 2 | Observed difference | P-value of the permutation test (100,000 permutations) | P-value adjusted using the Benjamini–Hochberg (BH) procedure |
|---|---|---|---|---|---|---|---|---|---|
| **Wetwang Slack** | Ditch | Barrow (primary burial) | 99 | 209 | 0.045 | 0.105 | -0.060 | 0.00001 | 0.00006 |
| **Wetwang Slack** | Flat grave | Barrow (primary burial) | 47 | 209 | 0.060 | 0.105 | -0.045 | 0.0050 | 0.0151 |
| **Wetwang Slack** | Barrow (primary burial) | Barrow (secondary burial) | 209 | 23 | 0.105 | 0.063 | 0.042 | 0.0512 | 0.1025 |
| **Wetwang Slack** | Ditch | Flat grave | 99 | 47 | 0.045 | 0.060 | -0.016 | 0.4215 | 0.5562 |
| **Wetwang Slack** | Ditch | Barrow (secondary burial) | 99 | 23 | 0.045 | 0.063 | -0.018 | 0.4635 | 0.5562 |
| **Wetwang Slack** | Flat grave | Barrow (secondary burial) | 47 | 23 | 0.060 | 0.063 | -0.003 | 0.9616 | 0.9616 |
| **Pocklington** | Square/Rectangular barrow (primary burial) | Flat grave | 54 | 31 | 0.38 | 0.26 | 0.12 | 0.0110 | 0.1214 |
| **Pocklington** | Flat grave | Ditch | 31 | 5 | 0.26 | 0.38 | -0.12 | 0.2454 | 0.8411 |
| **Pocklington** | Square/Rectangular barrow (primary burial) | Circular barrow (primary burial) | 54 | 6 | 0.38 | 0.305 | 0.075 | 0.4011 | 0.8411 |
| **Pocklington** | Ditch | Circular barrow (primary burial) | 5 | 6 | 0.38 | 0.305 | 0.075 | 0.5539 | 0.8411 |
| **Pocklington** | Square/Rectangular barrow (primary burial) | Barrow (secondary burial) | 54 | 4 | 0.38 | 0.33 | 0.05 | 0.5919 | 0.8411 |
| **Pocklington** | Flat grave | Barrow (secondary burial) | 31 | 4 | 0.26 | 0.33 | -0.07 | 0.6179 | 0.8411 |
| **Pocklington** | Flat grave | Circular barrow (primary burial) | 31 | 6 | 0.26 | 0.305 | -0.045 | 0.6462 | 0.8411 |

While the above analysis allows us to examine the general degree of intra-site connectedness of individuals from each burial type, it is not informative about the strength of relatedness between different burial types. Thus, for each individual at both Wetwang Slack and Pocklington, we calculated the mean amount of IBD shared with all individuals belonging to each burial type within the same site. For each burial type, we obtained distributions of per-individual mean IBD sharing between individuals interred in burial type *A* and individuals interred in burial type *B*, including the within-type case (*A*→*A*) (Figure S53 and Figure S54). Differences between distributions were quantified using the difference in median IBD shared and permutation tests (n=100,000) based on random reassignment of burial-type labels at the individual level (respecting the original sample size of each represented burial type) were used to assess whether the observed differences in IBD sharing could arise by chance (see Figure S52 for an example). We tested the following types of comparisons:

- comparisons between within-type and between-type relatedness for each burial type (e.g. A→A vs A→B; A→A vs A→C; A→A vs A→D).
- comparisons between within-type distributions across different burial types (e.g. A→A vs B→B).

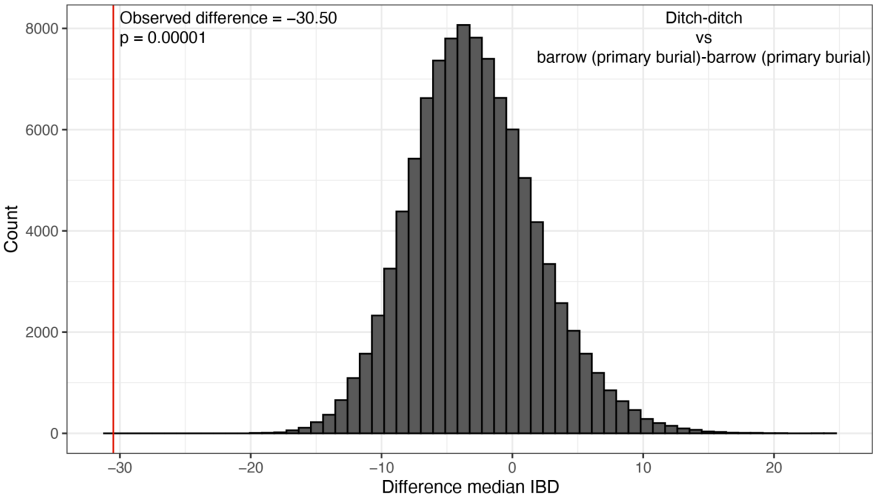


Figure S52. Permutation-based null distribution of the difference in median IBD sharing between ditch burials with other ditch burials and primary burials in barrows with other primary burials in barrows at Wetwang Slack. Burial type labels were randomly reassigned while keeping pairwise IBD values fixed. The vertical line indicates the observed difference. The empirical two-sided P-value corresponds to the proportion of permuted differences which matched or exceeded the observed value.

At Wetwang Slack, individuals from primary burials in barrows shared significantly more IBD with others from primary burials in barrows (median = 40 cM) than with individuals from other burial types (median = 13–18 cM) (Table S2). This pattern was not observed for the other burial categories (ditch graves, flat graves and secondary burials in barrows), where individuals shared similar median IBD values in both within-type and between-type burials (Table S2). Furthermore, IBD sharing between individuals in primary burials in barrows was significantly higher than in the other three within-type comparisons (i.e. ditch-ditch, flat grave-flat grave and secondary burials in barrows-secondary burials in barrows) (Table S2).

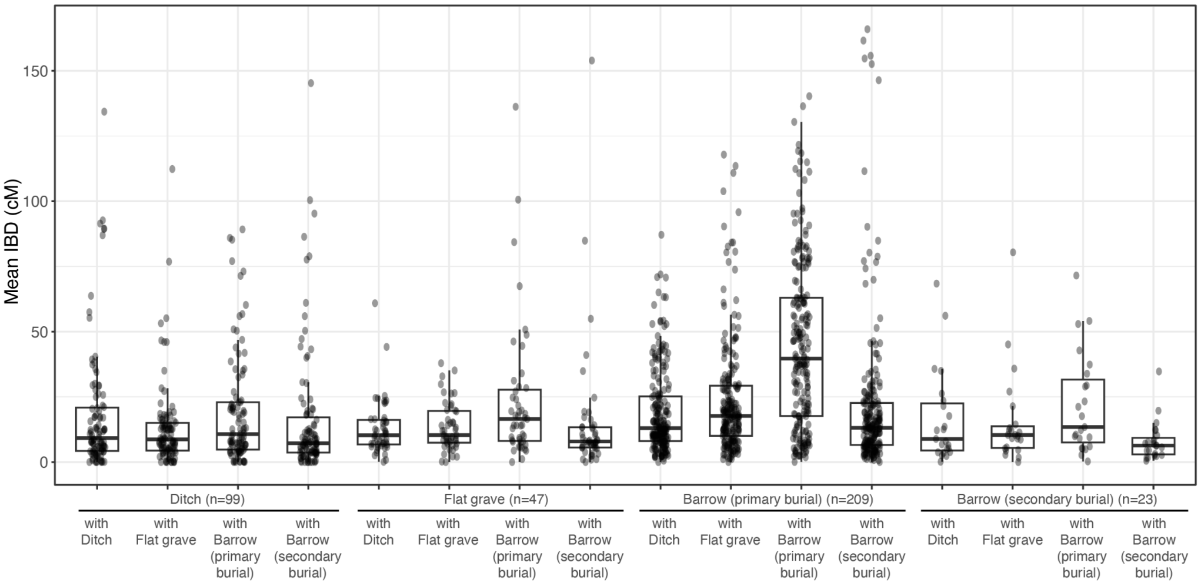


Figure S53. Per-individual mean IBD (cM) values at Wetwang Slack, grouped by burial type.

Due to smaller sample sizes, in order to further explore the pattern of IBD sharing between individuals in different burial types at Pocklington, we grouped together primary burials in square barrows with primary burials in rectangular barrows (n=54), and also grouped together individuals buried in a ditch with secondary burials in barrows (n=9). Individuals in circular barrows (n=6) share significantly more IBD with individuals from circular barrows (median = 161 cM) than with individuals from other burial types (median = 39–57 cM), including square/rectangular barrows (Table S3). This pattern suggests that circular and square/rectangular barrows may have held different social or funerary significance within the community. A similar pattern was observed for individuals buried in square/rectangular barrows (Table S3), who shared higher IBD with others from the same burial type (median = 72 cM) than with individuals from other types (median = 33–44 cM). Individuals in the remaining burial categories (flat graves and secondary burials in barrows), did not show significantly higher IBD sharing in within-type versus between-type burials (Table S3). When comparing within-type IBD values across burial types, both circular (median = 160.91 cM), and square/rectangular barrows (median = 71.52 cM) exhibited higher levels of relatedness than flat graves (median = 31.08 cM) and ditch/secondary burials in barrows (median = 45.54 cM).

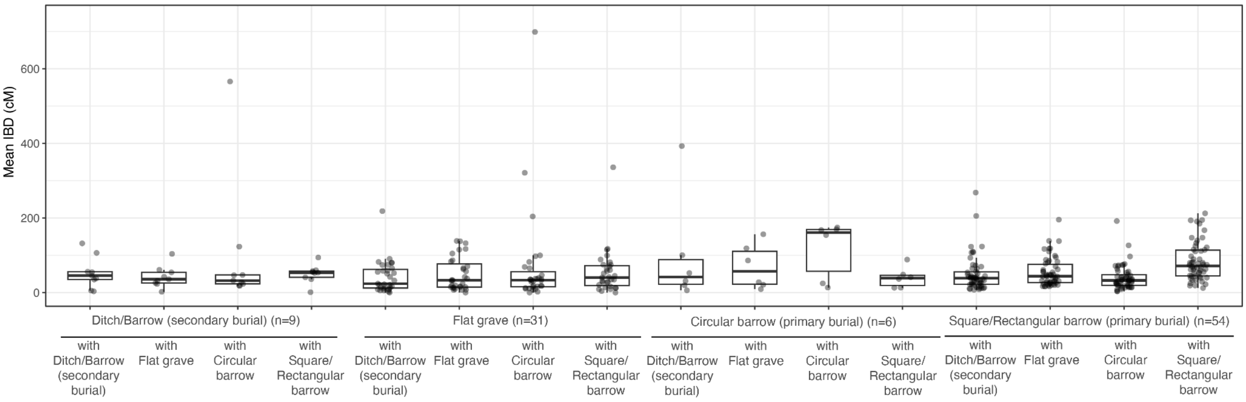


Figure S54. Per-individual mean IBD (cM) values at Pocklington, grouped by burial type.

Taken together, these patterns suggest that burial practices with regards to monument type at both Wetwang Slack and Pocklington were at least partly structured by biological relatedness. The concentration of close genetic relatives within primary barrows at Wetwang Slack, and within circular and square/rectangular barrows at Pocklington, is consistent with the idea that these funerary contexts were preferentially used by particular family groups or lineages. In contrast, the more homogeneous patterns of IBD sharing observed in flat graves, ditch burials, and secondary interments indicate that these burial types likely incorporated a more genetically heterogeneous subset of the population. Since primary barrow burials are present across multiple generations in the reconstructed pedigrees at Wetwang Slack (Extended Data Fig. 6c), these patterns are not the product of chronology, i.e. diminishing space for large monuments in later phases of a site attracting a more heterogeneous burial community, and may, therefore, reflect differences in funerary selection, with certain burial contexts representing more restricted or lineage-associated practices, while others functioned as more inclusive or community-level burial spaces.

Table S2. Comparing the distributions of per-individual mean IBD values across different burial types at Wetwang Slack.

| Group 1 | Group 2 | n1 | n2 | median group 1 | median group 2 | Observed difference | P-value permutation test | P-value adjusted using the Benjamini–Hochberg (BH) procedure |
|---|---|---|---|---|---|---|---|---|
| Ditch with Ditch | Ditch with Flat grave | 99 | 99 | 9.18 | 8.70 | 0.47 | 0.92 | 0.98 |
| Ditch with Ditch | Ditch with Barrow (primary burial) | 99 | 99 | 9.18 | 10.70 | -1.52 | 0.79 | 0.97 |
| Ditch with Ditch | Ditch with Barrow (secondary burial) | 99 | 99 | 9.18 | 7.25 | 1.93 | 0.87 | 0.97 |
| Flat grave with Flat grave | Flat grave with Ditch | 47 | 47 | 10.36 | 10.28 | 0.08 | 0.99 | 0.99 |
| Flat grave with Flat grave | Flat grave with Barrow (primary burial) | 47 | 47 | 10.36 | 16.52 | -6.16 | 0.50 | 0.97 |
| Flat grave with Flat grave | Flat grave with Barrow (secondary burial) | 47 | 47 | 10.36 | 7.90 | 2.45 | 0.55 | 0.97 |
| Barrow (primary burial) with Barrow (primary burial) | Barrow (primary burial) with Ditch | 209 | 209 | 39.67 | 13.00 | 26.67 | 0.000010 | 0.000045 |
| Barrow (primary burial) with Barrow (primary burial) | Barrow (primary burial) with Flat grave | 209 | 209 | 39.67 | 17.69 | 21.98 | 0.000010 | 0.000045 |
| Barrow (primary burial) with Barrow (primary burial) | Barrow (primary burial) with Barrow (secondary burial) | 209 | 209 | 39.67 | 13.12 | 26.55 | 0.000010 | 0.000045 |
| Barrow (secondary burial) with Barrow (secondary burial) | Barrow (secondary burial) with Ditch | 23 | 23 | 6.35 | 8.87 | -2.53 | 0.78 | 0.97 |
| Barrow (secondary burial) with Barrow (secondary burial) | Barrow (secondary burial) with Flat grave | 23 | 23 | 6.35 | 10.39 | -4.04 | 0.42 | 0.97 |
| Barrow (secondary burial) with Barrow (secondary burial) | Barrow (secondary burial) with Barrow (primary burial) | 23 | 23 | 6.35 | 13.43 | -7.08 | 0.59 | 0.97 |
| Ditch with Ditch | Barrow (primary burial) with Barrow (primary burial) | 99 | 209 | 9.18 | 39.67 | -30.50 | 0.000010 | 0.000045 |
| Flat grave with Flat grave | Barrow (primary burial) with Barrow (primary burial) | 47 | 209 | 10.36 | 39.67 | -29.32 | 0.000130 | 0.000468 |
| Barrow (primary burial) with Barrow (primary burial) | Barrow (secondary burial) with Barrow (secondary burial) | 209 | 23 | 39.67 | 6.35 | 33.33 | 0.000790 | 0.002370 |

| | | | | | | | | |
|---|---|---|---|---|---|---|---|---|
| Ditch with Ditch | Barrow (secondary burial) with Barrow (secondary burial) | 99 | 23 | 9.18 | 6.35 | 2.83 | 0.78 | 0.97 |
| Ditch with Ditch | Flat grave with Flat grave | 99 | 47 | 9.18 | 10.36 | -1.18 | 0.86 | 0.97 |
| Flat grave with Flat grave | Barrow (secondary burial) with Barrow (secondary burial) | 47 | 23 | 10.36 | 6.35 | 4.01 | 0.57 | 0.97 |

Table S3. Comparing the distributions of per-individual mean IBD values across different burial types at Pocklington.

| **Comparison 1** | **Comparison 2** | **n1** | **n2** | **median 1** | **median 2** | **Observed difference** | **P-value of the permutation test (100,000 permutations)** | **P-value adjusted using the Benjamini–Hochberg (BH) procedure** |
|---|---|---|---|---|---|---|---|---|
| Ditch/Barrow (secondary burial) with Ditch/Barrow (secondary burial) | Ditch/Barrow (secondary burial) with Square/Rectangular barrow (primary) | 9 | 9 | 45.54 | 53.27 | -7.73 | 0.78 | 0.82 |
| Ditch/Barrow (secondary burial) with Ditch/Barrow (secondary burial) | Ditch/Barrow (secondary burial) with Circular barrow (primary) | 9 | 9 | 45.54 | 32.08 | 13.46 | 0.45 | 0.779 |
| Ditch/Barrow (secondary burial) with Ditch/Barrow (secondary burial) | Ditch/Barrow (secondary burial) with Flat grave | 9 | 9 | 45.54 | 35.90 | 9.64 | 0.66 | 0.921 |
| Flat grave with Flat grave | Flat grave with Ditch/Barrow (secondary burial) | 31 | 31 | 33.08 | 23.83 | 9.25 | 0.57 | 0.625 |
| Flat grave with Flat grave | Flat grave with Square/Rectangular barrow (primary) | 31 | 31 | 33.08 | 40.22 | -7.14 | 0.564 | 0.696 |
| Flat grave with Flat grave | Flat grave with Circular barrow (primary) | 31 | 31 | 33.08 | 33.17 | -0.09 | 0.997 | 0.997 |
| Circular barrow (primary) with Ditch/Barrow (secondary burial) | Circular barrow (primary) with Circular barrow (primary) | 6 | 6 | 41.73 | 160.91 | -119.18 | 0.018 | 0.039 |
| Circular barrow (primary) with Flat grave | Circular barrow (primary) with Circular barrow (primary) | 6 | 6 | 56.93 | 160.91 | -103.98 | 0.019 | 0.039 |
| Circular barrow (primary) with Square/Rectangular barrow (primary) | Circular barrow (primary) with Circular barrow (primary) | 6 | 6 | 38.87 | 160.91 | -122.04 | 0.013 | 0.039 |
| Square/Rectangular barrow (primary) with Ditch/Barrow (secondary burial) | Square/Rectangular barrow (primary) with Square/Rectangular barrow (primary) | 54 | 54 | 38.72 | 71.52 | -32.80 | 0.027 | 0.039 |
| Square/Rectangular barrow (primary) with Flat grave | Square/Rectangular barrow (primary) | 54 | 54 | 43.90 | 71.52 | -27.62 | 0.004 | 0.039 |

| | | | | | | | | |
|---|---|---|---|---|---|---|---|---|
| | with Square/Rectangular barrow (primary) | | | | | | | |
| Square/Rectangular barrow (primary) with Circular barrow (primary) | Square/Rectangular barrow (primary) with Square/Rectangular barrow (primary) | 54 | 54 | 32.69 | 71.52 | -38.82 | 0.016 | 0.039 |
| Ditch/Barrow (secondary burial) with Ditch/Barrow (secondary burial) | Circular barrow (primary) with Circular barrow (primary) | 9 | 6 | 45.54 | 160.91 | -115.36 | 0.034 | 0.049 |
| Flat grave with Flat grave | Circular barrow (primary) with Circular barrow (primary) | 31 | 6 | 33.08 | 160.91 | -127.83 | 0.017 | 0.039 |
| Circular barrow (primary) with Circular barrow (primary) | Square/Rectangular barrow (primary) with Square/Rectangular barrow (primary) | 6 | 54 | 160.91 | 71.52 | 89.39 | 0.029 | 0.051 |
| Ditch/Barrow (secondary burial) with Ditch/Barrow (secondary burial) | Square/Rectangular barrow (primary) with Square/Rectangular barrow (primary) | 9 | 54 | 45.54 | 71.52 | -25.97 | 0.34 | 0.557 |
| Flat grave with Flat grave | Square/Rectangular barrow (primary) with Square/Rectangular barrow (primary) | 31 | 54 | 33.08 | 71.52 | -38.43 | 0.007 | 0.039 |
| Ditch/Barrow (secondary burial) with Ditch/Barrow (secondary burial) | Flat grave with Flat grave | 9 | 31 | 45.54 | 33.08 | 12.46 | 0.64 | 0.921 |
| Barrow with Barrow | Barrow with Other | 60 | 60 | 66.35 | 44.64 | 21.71 | 0.009 | 0.039 |
| Barrow with Barrow | Other with Other | 60 | 40 | 66.35 | 37.26 | 29.10 | 0.017 | 0.039 |
| Other with Barrow | Other with Other | 40 | 40 | 48.25 | 37.25 | 10.99 | 0.285 | 0.387 |

Finally, we explored whether burial type differed between the two dominant maternal lineages at Wetwang Slack, and between individuals in the 195-individual pedigree and outside the pedigree. We found no significant difference in burial-type between individuals belonging to T2e1a1b and H1ao ($\chi^2$ test; df = 3, p = 0.477). Although T2e1a1b individuals (belonging to the most dominant of the two lineages) were slightly more frequently interred as primary barrow burials (68% in T2e1a1b versus 59% in H1ao) and H1ao individuals were slightly more frequently interred as ditch burials (18% in T2e1a1b versus 25% in H1ao), these differences do not provide statistical evidence for differential funerary treatment between the two maternal lineages and, as noted previously, can neither be explained as a product of chronology. Restricting the comparison to the predominant T2e1a1b haplotype (with mutation 2416C; Figure S6) increased the descriptive difference between the two maternal lineages, with T2e1a1b individuals more often buried as primary barrow burials than H1ao individuals (73.4% versus 58.8%), and H1ao individuals more often represented in ditch burials (25.0% versus 12.9%). However, differences in the overall burial-type distribution between the two lineages were not statistically significant ($\chi^2$ test; df = 3, p = 0.114). Conversely, burial-type distributions differed markedly between individuals inside and outside the 195-individual pedigree ($\chi^2$ test; df = 3, $p = 1.08 \times 10^{-8}$). Pedigree individuals were substantially enriched in primary barrows: 71% were buried in primary barrows, compared to 39% of non-pedigree individuals. Conversely, non-pedigree individuals were more commonly found in ditch burials (36%), compared with 17% pedigree individuals. Flat graves were also less frequent among pedigree individuals (8%) than among non-pedigree individuals (16%), while secondary barrow burials showed a weaker difference (4.1% versus 8.7%). This pattern suggests that primary barrows formed a more lineage-restricted component of the cemetery, while ditch and flat-grave contexts included a higher proportion of individuals outside the main reconstructed pedigree.

#### Correlation between genetic relatedness and burial distance

We assessed the spatial structure of biological relatedness within Wetwang Slack and Pocklington by testing the association between pairwise genetic sharing and geographic distance between individuals. Individuals I36995, I37331, I36978, I36892, I37330, I36927, I37329, I37326, I37328, I36823 and I36926 were excluded from the analysis, as they were lay outside the main burial area at Wetwang Slack. Pairwise distances were calculated from latitude and longitude coordinates and genetic relatedness was measured by total length of the genome in shared IBD (sum_IBD>8 cM + sum_IBD2). We performed Mantel tests to assess the correlation between the geographic distance matrix and the matrix of pairwise IBD values, while accounting for the non-independence of pairwise comparisons (Figure S55). At Wetwang Slack, a weak but significant negative association ($\rho = -0.044$, Mantel $p = 1.0 \times 10^{-4}$) between genetic relatedness and burial distance was detected (i.e. as biological relatedness increased, distance between burials decreased). At Pocklington, a similar weak (though not statistically significant) association was detected ($\rho = -0.034$, Mantel p = 0.13).

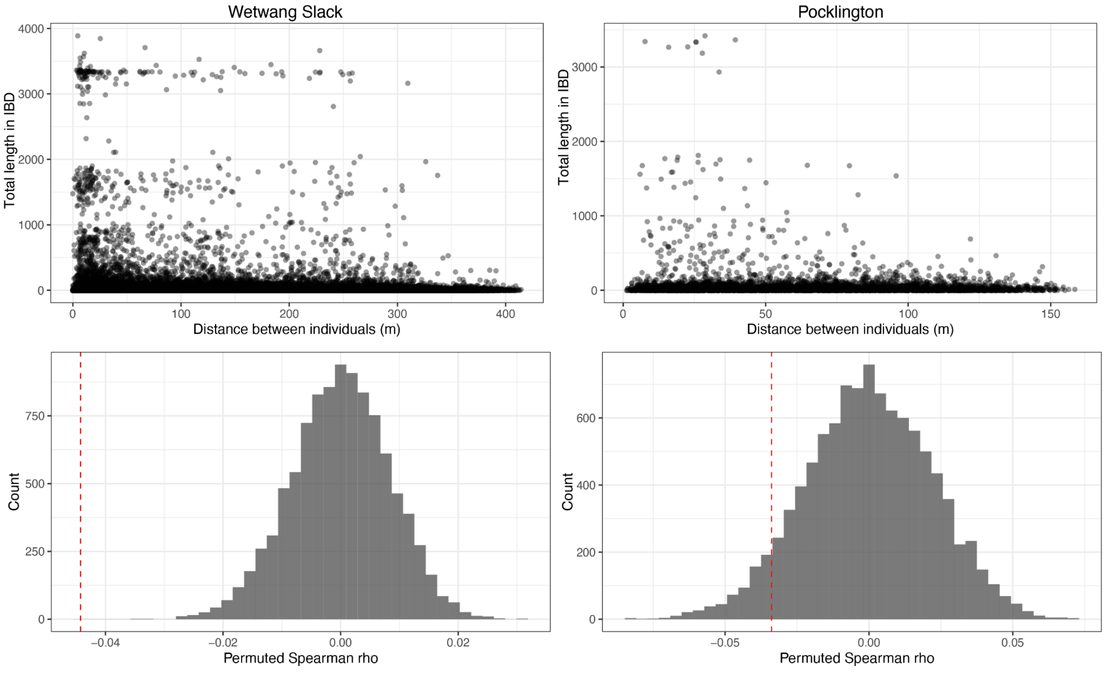


Figure S55. Relationship between pairwise IBD sharing and spatial distance at Wetwang Slack and Pocklington. Scatterplots show total IBD sharing as a function of inter-individual distance, and histograms show null distributions of Spearman's ρ under 10,000 permutations. Red dashed lines indicate the observed correlation.

We can look at this in a different way by grouping all intra-site pairs of individuals into 5 m distance bins and calculating, for each bin, the number of first-, second-, and third-degree relatives. Across both sites, close biological relationships peak within the first few distance bins and decline rapidly with increasing distance (Figure S56). At Wetwang Slack, this signal is especially strong, reflecting both the larger sample size and the extensive internal pedigree structure at the site. This indicates that close biological relatives were more often buried in spatial proximity, although closely related individuals at Wetwang Slack can also be identified up to 300 m apart.

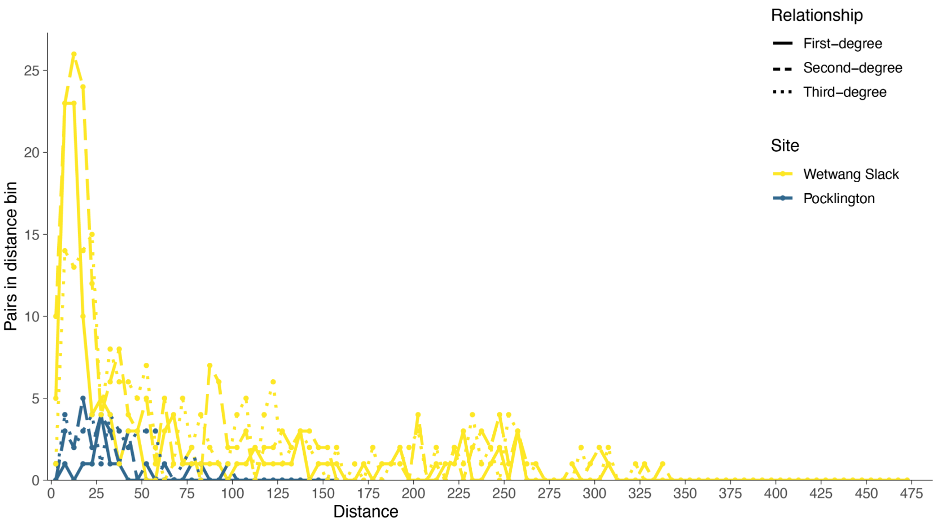


Figure S56. Distribution of close biological relationships by burial distance. Within each site, all pairs of individuals were grouped into 5 m distance bins according to the distance between their burials. Lines show the absolute number of pairs classified as first-degree, second-degree, or third-degree relatives.

To explore the spatial organisation of the cemeteries in more detail, we next tested whether close relative pairs were buried significantly closer to one another in the cemetery compared to other pairs of individuals. Statistical significance was assessed using a permutation test. For each site, the geographic distance matrix was kept fixed, while the relationship-classification matrix was permuted by jointly permuting rows and columns, thereby preserving the internal structure of the pairwise relationship matrix while breaking its correspondence with spatial location. For each permutation, the group labels were recalculated and the difference in median distance between the two groups was recomputed. One-sided p-values were calculated as the proportion of permutations in which the permuted median difference was equal to or smaller than the observed median difference.

The analysis demonstrated that close biological relatives were consistently buried closer to one another than other pairs of individuals, at both Wetwang Slack and Pocklington (Supplementary Table 8). At Wetwang Slack, first-degree relatives had a median burial distance of 21.2 m, compared with 126.5 m for other pairs; this corresponds to a median difference of −105.3 m ($p = 1.0 \times 10^{-4}$). This pattern is replicated for third-degree or closer relatives (Extended Data Figure 5a), with a median distance of 36.9 m versus 127.0 m for other pairs ($p = 1.0 \times 10^{-4}$). At Pocklington, closely related pairs of individuals were also buried significantly closer to one another than other pairs of individuals, with third-degree or closer relatives having a median burial distance of 28.3 m, compared with 58.0 m for other pairs ($p = 1.0 \times 10^{-4}$). Thus, both sites show a clear spatial clustering based on biological relatedness.

Pairs of individuals sharing the same mtDNA haplotype were also buried significantly closer to one another than non-matching pairs, but the strength and robustness of this signal differed between sites (Supplementary Table 8). At Wetwang Slack, mtDNA-matching pairs had a median burial distance of 113.1 m, compared with 128.1 m for non-matching pairs ($p = 1.0 \times 10^{-4}$). This pattern persisted, even after excluding third-degree or closer relatives ($p = 1.0 \times 10^{-4}$) (Extended Data Figure 5b). This indicates that the spatial clustering of mtDNA-sharing individuals at Wetwang Slack is not driven solely by close relatives. At Pocklington, the mtDNA signal was weaker, with mtDNA-matching pairs having a median distance of 51.7 m compared with 58.3 m for non-matching pairs ($p = 0.032$). However, after excluding third-degree or closer relatives, this difference reduced and was no longer statistically significant (Supplementary Table 8): mtDNA-matching pairs had a median burial distance of 55.6 m compared with 58.4 m for non-matching pairs ($p = 0.194$). This suggests that the apparent relationship between mtDNA haplotype and spatial structure at Pocklington is largely explained by close genetic relatives rather than broader matrilineal clustering (although, as noted previously, small sample size might also be a factor).

Finally, we tested whether the proximity between father–offspring pairs was significantly different than mother–offspring pairs (Supplementary Table 8). At Wetwang Slack, 87 first-degree parental pairs could be classified, comprising 63 mother–offspring and 24 father–offspring pairs. Mother–offspring pairs had a lower median burial distance than father–offspring pairs, at 19.5 m versus 29.2 m, respectively. However, this difference was not statistically significant under the permutation test ($p = 0.386$). At Pocklington, this comparison could not be formally tested because only four mother–offspring pairs and no father-offspring pairs were available. Overall, this analysis does not provide evidence for a significant difference in burial proximity between mother–offspring and father–offspring pairs, although statistical power is limited by the small number of classifiable father–offspring relationships.

*Distance between reproductive partners*

At Wetwang Slack, in only eight cases were both reproductive partners buried in the cemetery (Figure S57). The mean distance between them was 23 m, which is statistically significantly closer (p-value $< 0.001$) than that expected for eight random pairs of individuals in the cemetery (Figure S58). We can, therefore, conclude that, at Wetwang Slack, although it was rare to bury both reproductive partners in the same cemetery, in cases where this happened, they were buried close to one another.

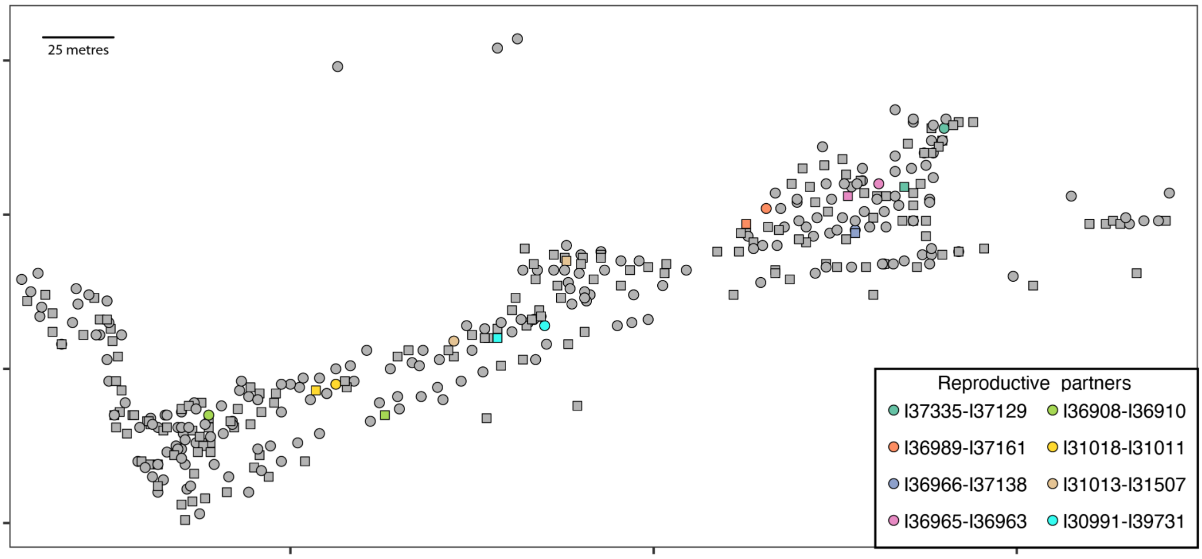


Figure S57. Spatial distribution of the eight inferred reproductive pairs at Wetwang Slack in which both partners were interred in the cemetery. Grey symbols show other sampled individuals, while coloured symbols indicate members of each reproductive pair. Male individuals are represented by squares and female individuals by circles.

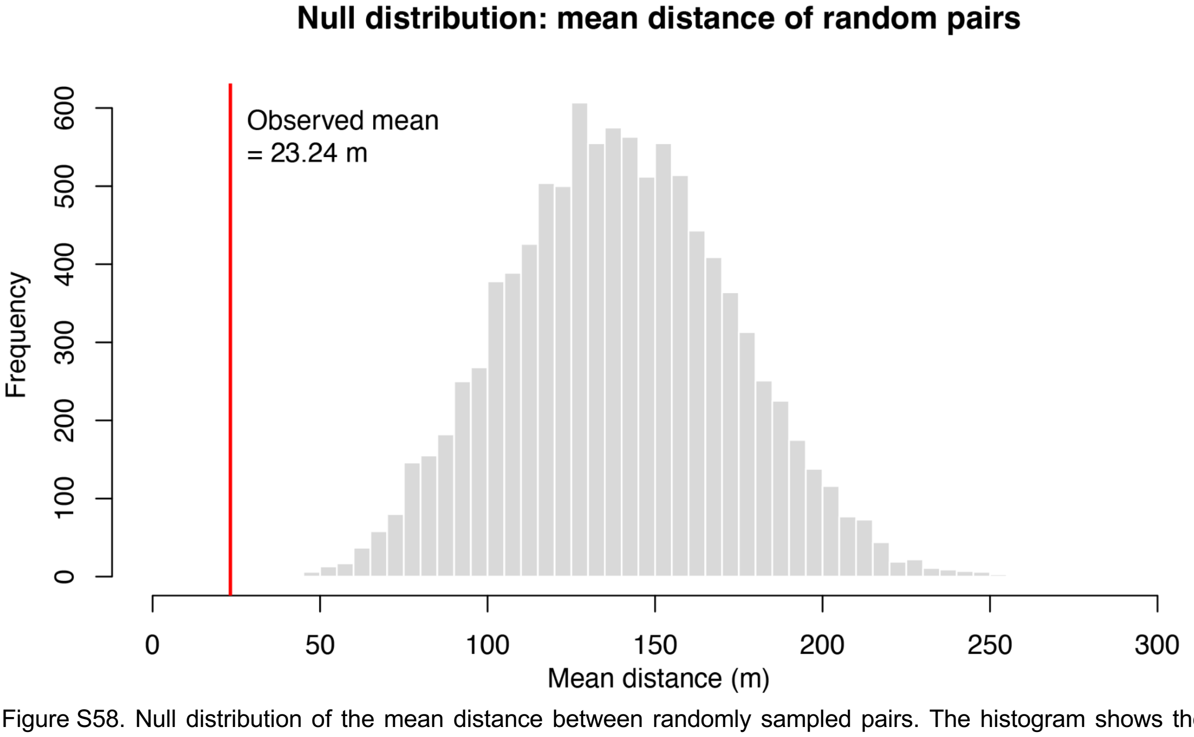


Figure S58. Null distribution of the mean distance between randomly sampled pairs. The histogram shows the distribution of mean pairwise distances obtained by randomly sampling, in each permutation (n=10000), the same number of pairs as the observed reproductive pairs. The red vertical line marks the observed mean burial distance between reproductive partners.

*Cemetery location of biological relatives of females I31517 and I36803*

In this section we focus on two women who sit at the top of the large pedigree and who have a large number of descendants across many generations. Female I31517 is the individual from Wetwang Slack with the highest number of intra-site relatives. She was the offspring of closely related parents (approximately third-degree) (Supplementary Table 1), and the founder of a family with 37 descendants buried at Wetwang Slack over 10 generations. Female I36803 has 11 descendants buried at Wetwang Slack over 7 generations, and shares one descendant with I31517 (Extended Data Figure 7a).

Both females tend to have close relatives, and descendants and their reproductive partners, buried in close proximity to them within the cemetery (Extended Data Figure 7b). In the case of I31517, this pattern is particularly evident during the earlier generations, whereas descendants from generation seven onwards appear substantially more dispersed across the cemetery.

To further investigate whether the burial locations of these females acted as focal points for their close relatives, we computed, for each individual at Wetwang Slack, median burial distance to members of the I31517 and I36803 lineages. Among the 390 sampled individuals at the site, I31517 has the third lowest median distance to members of her lineage (20.2 m), while I36803 has the eighth lowest median distance to members of her lineage (13.6 m) (Figure S59). These patterns suggest that the burial locations of certain females retained genealogical significance across multiple generations and may have functioned as enduring spatial anchors around which the mortuary practices of close biological relatives within the cemetery were organised.

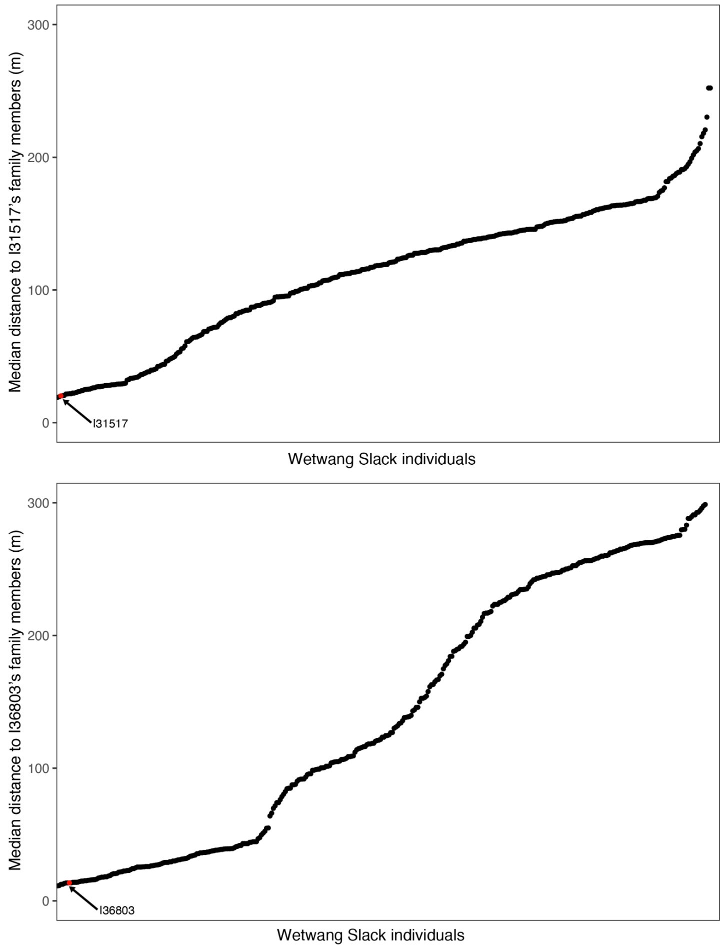


Figure S59. Median distance between Wetwang Slack individuals and members of the I31517 (top) and I36803 (bottom) pedigrees. Individuals on the x-axis are ordered according to their median burial distance to members of I31517's lineage (top) and I36803's lineage (bottom). Red dots indicate the two focal women, I31517 and I36803.

*Cemetery location by generation*

To explore how the reconstructed pedigree relates to spatial organisation within the cemetery, we counted the number of individuals assigned to each of the 13 generations in the pedigree and grouped them by burial area within the cemetery (Figure S60; Supplementary Table 1). We excluded individuals whose position in the pedigree was speculative and who could conceivably be assigned to more than three possible generations. The distribution shows that the pedigree is not evenly represented across the cemetery through time. The earliest generations are represented by very few individuals and are concentrated in the central and western parts of the cemetery (Extended Data Figure 6a). From generation four onwards, the number of individuals increases markedly, reaching the highest values between generations six and eight. These middle generations are distributed across several cemetery areas, but are dominated by individuals from the main central and eastern areas. In later generations, the total number of individuals declines, with most individuals located in the main central and eastern areas.

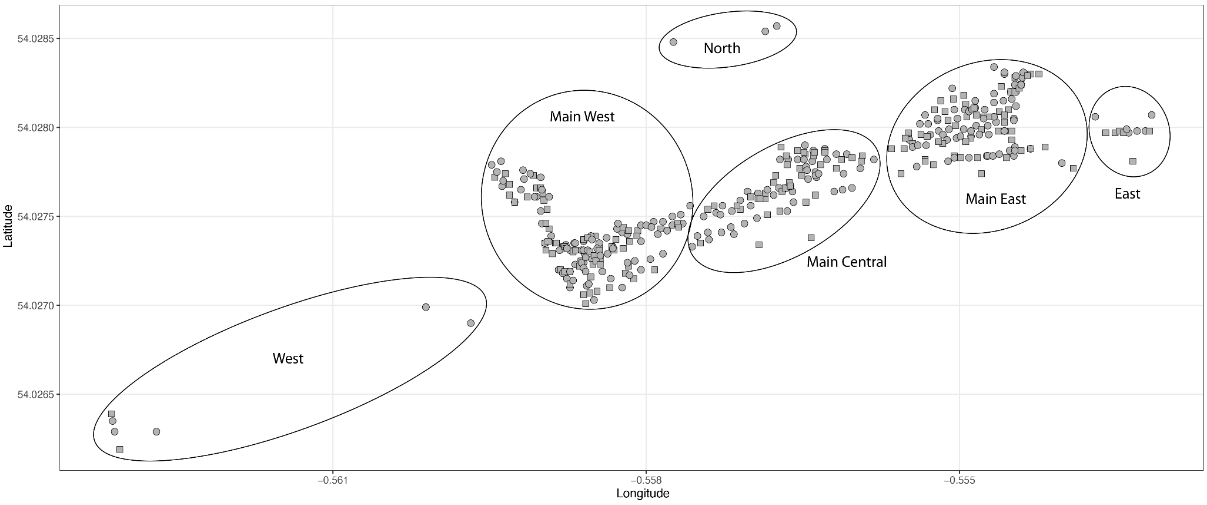


Figure S60. The Wetwang Slack cemetery is divided into six areas.

#### Patterns of social organisation

*Sex bias*

We tested for sex bias within each of the three main cemeteries (Supplementary Table 9). Wetwang Slack showed a significant excess of females, with 219 (56.2%) females and 171 males (exact binomial test, $p = 0.017$). If we focus on adult individuals, the number of females was also significantly higher than males (59.5% females; $p = 0.00063$) Among non-adults, males were more frequent than females (37 males versus 22 (37.3%)) females), but this was not statistically significant ($p = 0.067$). In contrast, no significant deviation from a balanced sex ratio was observed at Pocklington (54 females versus 46 males; $p = 0.484$) or Melton 1 (11 females versus 17 males; $p = 0.345$). Within Wetwang Slack, the female bias was concentrated among individuals belonging to the large pedigree, with 119 (61.0%) females and 76 males ($p = 0.0025$), whereas individuals outside this pedigree showed an approximately balanced sex ratio (100 (51.3%) females versus 95 males; $p = 0.775$). It is also particularly pronounced within the dominant T2e1a1b maternal lineage, which includes 92 (64.8%) females and 50 males ($p = 5.3 \times 10^{-4}$). In contrast, the second major lineage, H1ao, shows no evidence of sex bias, with 38 females (47.5%) and 42 males ($p = 0.738$). These results indicate that the female bias at Wetwang Slack is primarily associated with the large group of individuals connected by close biological relatedness, particularly within the dominant T2e1a1b matriline.

*Presence of parents and reproductive partners*

In this section we assess the extent to which the burial population at Wetwang Slack represents a complete residential community. If the cemetery largely reflected a living community, we would expect many individuals to have both biological parents also buried at the site (i.e. many mother-offspring and father-offspring pairs). Instead, most individuals (80%) have no parents represented in the cemetery; 68 individuals (17%) have one parent and only 10 (3%) have both parents (Supplementary Table 10). This pattern is unlikely to be explained solely by unsampled individuals, given that we have recovered data from 390 out of 446 excavated burials. At Pocklington we observe a similar pattern, with 96% of individuals having no parent buried in the cemetery, and only 4% with one parent.

To explore these patterns in more detail, we then turned the focus from site-wide analysis of parent-offspring pairs to reproductive partners as observed in the reconstructed pedigrees. At Wetwang Slack, reproductive partners were very rarely both buried in the cemetery. Among the reproductive unions which could be genetically reconstructed, in only eight cases were both interred in the cemetery. In contrast, 13 unions had only the male partner buried, whereas 62 had only the female partner buried. At Pocklington, in only one case were both partners of a reproductive union buried at the cemetery, three had only the female buried and one had only the male buried.

This strong asymmetry suggests that the cemeteries at Wetwang Slack and Pocklington are not a straightforward reflection of co-resident nuclear families. Instead, many reproductive unions involved one partner who was absent from the cemetery, and this absent partner was most often male. In other words, females who reproduced within the Wetwang Slack pedigree were much more likely than their male reproductive partners to be buried at Wetwang Slack. This pattern is consistent with the cemetery being organized around local female lines, with many male reproductive partners being buried elsewhere, perhaps in their own natal communities. The Wetwang Slack burial community should not, therefore, be considered as a simple proxy for the full residential population. Rather, it appears to represent a socially selected burial community based on particular genealogical lines.

*Number of matrilineages, patrilineages and bilineages*

To assess whether reproductive unions at Wetwang Slack were preferentially structured through maternal or paternal lines, we classified each union according to the presence of close biological relatives of the various reproductive partners within the cemetery. A union was classified as matrilineal if the mother had first- or second-degree relatives in earlier generations, and the father did not. Conversely, patrilineal descent was ascribed when the father had first- or second-degree relatives in earlier generations, but the mother did not. Finally, bilateral descent was ascribed when both reproductive partners had first- or second-degree relatives in earlier generations buried at the site.

We identified 84 matrilineages, 17 patrilineages and 18 bilineages in the Wetwang Slack cemetery (Supplementary Table 11). Thus, matrilineages accounted for 71.2% of detectable reproductive unions. The distribution differed strongly from equal representation of the three categories ($\chi^2$ goodness-of-fit test: $\chi^2 = 74.3$, df = 2, $p = 7.2 \times 10^{-17}$), and matrilineal unions were significantly more frequently present than those reflecting patrilineal and bilineal unions combined (exact binomial test: 84/119, $p = 8.2 \times 10^{-6}$).

These results indicate strong asymmetry in the transmission of local genealogical connections among the burial population at Wetwang Slack, with reproductive unions much more often linked to pre-existing maternal than paternal relatives. Importantly, this pattern is not restricted to a single family, but is observed in all clusters within the main pedigree of 195 individuals (Supplementary Table 11). At Pocklington, matrilineages (n=3) were also more common than patrilineages (n=1), but the lack of extended pedigrees prevented us from studying this pattern more formally.

*Exogamy*

Direct evidence for male exogamy derives from the closest biological relationships across sites. The closest genetic link between Wetwang Slack and Pocklington (Extended Data Figure 8a), between Wetwang Slack and Melton 1 (Extended Data Figure 8b), and between Pocklington and Melton 1 (Extended Data Figure 8c), all involve connections through male individuals.

To assess whether we could identify indirect evidence of exogamy in the three largest burial populations (at Wetwang Slack, Pocklington and Melton 1), we examined the presence of any sex-bias among adult individuals within the reconstructed pedigrees who had first- or second-degree relatives buried in earlier generations of the same cemetery. We interpret the presence of such relatives in earlier generations as evidence that an individual was descended from a locally established genealogy. At Wetwang Slack, adult daughters (n=69) were more frequently represented than adult sons (n=46) with first- or second-degree relatives in earlier generations of the cemetery (Supplementary Table 12). This represents a significant excess of females among adult individuals with demonstrable local genealogical continuity (60%; $p = 0.039$). The sex bias becomes stronger when considering only individuals who themselves had descendants represented in the reconstructed pedigrees (Supplementary Table 12). At Wetwang Slack, 32 adult females with first- or second-degree relatives in earlier generations had identified descendants, compared with only 10 adult males (76.2% female; $p = 9.4 \times 10^{-4}$). This pattern is not restricted to a single branch of the pedigree, but is observed across most of the major clusters. The signal is weaker at the other sites. At Pocklington, females with close relatives in earlier generations were more frequent than males, but the numbers are small (nine females and five males), and the difference is not statistically significant ($p = 0.424$). Meanwhile, at Melton 1, only one adult male met this criterion and no meaningful test could be performed.

The under-representation of adult sons from established lineages at Wetwang Slack is consistent with male exogamy. However, the presence of some adult sons at Wetwang Slack indicates that burial location need not necessarily correspond directly to residence during life. One explanation is that many sons from Wetwang Slack lineages joined other communities during life, and some were returned or selected for burial in the cemetery of their natal family.

Of the 46 adult sons buried at Wetwang Slack, 36 (78%) had no descendants buried at the cemetery, compared with 54% of equivalent adult daughters. In a matrilineal descent system, the offspring of exogamous males would be expected to be buried elsewhere, most likely in the cemetery associated with their reproductive partner's maternal lineage, and this was possibly the case for the offspring of those 36 adult sons with no descendants at Wetwang Slack. The remaining 10 adult sons who did have descendants buried at Wetwang Slack all reproduced with females who also had close relatives buried in the cemetery. These cases suggest that adult sons from Wetwang Slack lineages were more likely to have descendants buried at the cemetery when their reproductive partner also belonged to a locally established genealogy. In such cases, the cemetery of the male's natal lineage and that of his reproductive partner's lineage overlapped, potentially explaining why these males, unlike most adult sons from Wetwang Slack families, were buried in the same cemetery as their descendants.

Below we summarize the 10 adult males who have both close biological relatives in earlier generations and descendants buried at Wetwang Slack:

-I36896 (cluster 2) belonged to a lineage established at Wetwang Slack two generations before him (Figure S17). He had a daughter (I36787) with an unsampled woman who also had at least two close relatives at Wetwang Slack: male I31018 (a second-degree relative) and female I30997 (a ~third-degree relative).

-I37335 (cluster 2) was I36896's nephew through an unsampled sister, and had two siblings. I37335 had two sons (I37344, I37130) with I37129 who, like I37335, also had two siblings and belonged to a lineage established at Wetwang Slack three generations before her (Figure S17). The two reproductive partners were themselves third cousins. I37335, I37129, their respective siblings and their sons were all buried at the eastern edge of the cemetery. However, their sons were buried closer to their mother I37129 (5 m away) and her siblings than they were to their father (I37335), who was buried closer to his own siblings (Extended Data Figure 10b). This is a good example of a male from an established maternal lineage at Wetwang Slack who was buried in the same cemetery as his sons because his reproductive partner (their mother) was also a descendant of the Wetwang Slack pedigree.

-I37137 (cluster 2) belonged to a lineage that can be traced back six generations through both his maternal grandfather and maternal grandmother (Figure S17 and Figure S28). He had one daughter (I37141) who was buried close to him. I37141's mother was not sampled, but one of I37141's mother's first-degree relatives (male I36801) was also buried close to I37141, indicating that I37141's maternal family was also present in the cemetery.

-I31512 (cluster 3) is the only male descendant of I31517 whose offspring (I30962 and I31509) were buried at Wetwang Slack. He reproduced with an unsampled female with several close paternal relatives buried in the cemetery, including her half-siblings I37138 and I31508.

-I31507 had his father (I36888) (cluster 3) and two of his father's close relatives (I36925 and I36783) (cluster 4) buried in the cemetery. He also had other more distant relatives at Wetwang Slack, including his fourth/fifth-degree relative I31011 (cluster 2). I31507 had a daughter (I30961) with I31517's great-granddaughter. Unlike his father, who was buried at the western edge of the cemetery, I31507 was buried close to his daughter and to I31517.

-I36904 (cluster 4) was buried at Wetwang Slack, as was his mother and close paternal relatives. He reproduced with an unsampled female who was I36888's second-degree relative.

-I36819 (cluster 6) had several maternal relatives buried at the cemetery, including his grandmother and half-brother. He had one daughter (I36989) with an unsampled woman whose mother (I36815) and other maternal relatives were also buried at Wetwang Slack.

-I37138 (cluster 5) had several close relatives buried at the cemetery, including his sister and maternal grandmother. He reproduced with I36966, a daughter of I36803 who also had several close relatives in the cemetery. I37138 was buried in the ditch of I36966's barrow, close to his descendants and away from his sister and grandmother.

-I30965 (cluster 5) was the son of female I30988 (Figure S28), the Wetwang Slack individual with the second highest proportion of intra-site relatives. He had a son (I36883) with an unsampled female from a family with many members buried in the cemetery, including the unsampled female's grandmother (I30971). I30965 was buried in the central part of the cemetery, close to his mother and half-siblings, but far from his son and his son's maternal relatives, who were buried in the western part of the cemetery.

-I36821 was also I30988's son and I30965's half-brother (Figure S28). Like I30965, he reproduced with an unsampled female from a family present at Wetwang Slack with members such as I31018 or I30906. I36821 was buried in the central area of the cemetery, very close to his mother and maternal half-siblings.

*Multiple reproductive partners*

We identified individuals in the reconstructed pedigrees who had offspring with more than one reproductive partner. We use the term "multiple reproductive partners" in a strictly genetic sense, referring to individuals inferred to have had offspring with more than one partner represented in the pedigree. This does not necessarily imply socially recognized (or socially sanctioned) polygyny, polyandry, simultaneous unions, or co-residence.

At Wetwang Slack, we identified 27 individuals with multiple reproductive partners (Supplementary Table 13). Of these, 20 were females and seven were males. Among the females, 12 of these individuals were physically present/sampled, while eight were inferred from the pedigree structure. In contrast, none of the seven males with multiple female reproductive partners was recovered in the cemetery; all of them were inferred from genetic analysis of their offspring. Most individuals with multiple reproductive partners had offspring with two partners, although a smaller number had offspring with three partners. Among females with multiple male partners, 16 had two reproductive partners and four had three. Among inferred males with multiple female partners, six had two reproductive partners and one had three.

A striking feature of these unions is that multiple reproductive partners of the same individual were almost never jointly represented in the cemetery. In other words, although several individuals had offspring with more than one partner, we do not observe cases in which two or more of those partners were themselves buried at Wetwang Slack. The only exception is I36814 and I36884 (cluster 2) (Figure S17), who were mother and daughter who both had offspring with the same male, who was not recovered at the site. Besides this case, we found another case of a person reproducing with two partners who were themselves related: the unsampled mother of I30998 and I36874 (cluster 3) (Figure S20) who reproduced with I37339 and with an unsampled male who was most likely I37339's second degree relative.

This pattern reinforces the broader observation that reproductive partners were often absent from the cemetery, especially males. The absence of any recovered male with multiple reproductive partners suggests that males involved in these repeated reproductive links were not selected for burial within the excavated cemetery. Conversely, the recovery of several females with multiple reproductive partners is consistent with the central role of female lineages within the Wetwang Slack pedigree.

Similar cases were much rarer to detect outside Wetwang Slack, due to the lack of extended reconstructed pedigrees (Supplementary Table 13). At Pocklington, we identified two unsampled females with two reproductive partners each, whereas no individuals with multiple reproductive partners were identified at Melton 1.

*Individuals joining Wetwang Slack families as reproductive partners*

To assess whether reproductive partners entering the reconstructed pedigrees derived from outside communities or instead belonged to local Wetwang Slack lineages, we compared the number of their identified descendants in the pedigree with the total number of their moderately close intra-site relatives. We focused on sampled individuals identified as reproductive partners of people belonging to lineages already represented in previous generations of the cemetery. If both members of a reproductive partnership had relatives in previous generations, we chose one of them for this analysis. For each of these individuals, we counted the number of relatives at Wetwang Slack sharing >150 cM in IBD (corresponding to approximately fifth-degree relatives or closer), and compared this value with the number of descendants identified in the reconstructed pedigrees.

If these reproductive partners had originated from outside Wetwang Slack and had no pre-existing biological ties to the burial community, we would expect most of their intra-site relatives to be their own descendants. In that scenario, individuals should fall close to the 1:1 expectation between number of descendants and number of intra-site relatives. Instead, all analysed reproductive partners have more intra-site relatives than identified descendants (Extended Data Figure 9). Several individuals show a particularly large excess of relatives compared with descendants (Supplementary Table 14), including: I37129, who has seven identified descendants but 29 intra-site (fifth-degree or closer) relatives; I37161, with two descendants but 21 relatives; and I36806, with one descendant but 16 relatives. Although some unidentified descendants not incorporated into the reconstructed pedigrees may contribute to intra-site relative counts, they are unlikely to explain the large, systematic excess of relatives over identified descendants. This pattern suggests that these individuals were not generally unrelated outsiders. Rather, many appear to have belonged to broader kin groups already represented in the cemetery at Wetwang Slack. For instance, female I36803 has 11 descendants, all buried close to her in the cemetery (Extended Data Figure 7b). Her unsampled son had a daughter (I36980) with I36806, a female belonging to the H1ao mtDNA haplogroup whose maternal and paternal relatives are well-represented at Wetwang Slack. I36806 and her daughter were buried very close to I36803 at the eastern part of the cemetery, unlike any of I36806's close maternal and paternal relatives.

In other cases, such as male I37318 who reproduced with one of I31517's descendants, none of his relatives (besides his daughter) could be placed within the family tree, but he shares 300 cM in IBD with male I36835 and 172 cM with female I37171, themselves third-degree relatives and both from Wetwang Slack, which indicates that I37318 had other relatives at the site and was not a complete outsider.

We also have additional evidence of reproductive partners (in this case unsampled) joining reconstructed pedigrees at Wetwang Slack who had other relatives at Wetwang Slack. For example, the father of female I30976 (cluster 3; I31517's great-great-granddaughter) has not been identified among the sampled individuals (Extended Data Figure 7a), but we know he was a third-degree relative of female I31001 from cluster 2 (Figure S17), who reproduced with a male in the third generation of a T2e1a1b matriline.

This pattern indicates that reproductive partnership at Wetwang Slack was sometimes embedded within pre-existing local kinship networks rather than involving individuals from other communities whose only biological connection to the cemetery population was through their descendants. However, these observations do not contradict the fact that most unions were likely exogamous because 1) most reproductive partners have not been recovered from the cemetery population at Wetwang Slack and so their genetic ties to the community cannot be studied, and 2) those few reproductive partners that are present at Wetwang Slack, and therefore available for analysis in this section (Extended Data Figure 9; Supplementary Table 14), are precisely those individuals who we would expect to have close pre-existing biological links with the Wetwang Slack community, because it is this fact that would have granted them eligibility for inclusion in the cemetery.

*Recurrent unions between the two dominant matrilines at Wetwang Slack*

To investigate whether reproductive unions at Wetwang Slack were structured by maternal-line affiliation, we examined all reproductive unions for which mitochondrial haplogroups of both reproductive partners could be determined. In a few cases, both partners were present among the sampled individuals, but in others, one or both mitochondrial haplogroups were inferred from their position within the reconstructed pedigree. We focused on the two dominant maternal lineages (T2e1a1b and H1ao) at Wetwang Slack, which together account for a large proportion (62%) of the biologically related individuals at the site (Figure 2b).

Among 45 reproductive unions for which both maternal lineages could be assigned, 25 involved one partner carrying T2e1a1b and the other H1ao, whereas 20 involved other combinations of maternal lineages (Supplementary Table 15) (Figure 3e); no union in which bother partners carried the same dominant (i.e. T2e1a1b or H1ao) maternal lineage were detected. Thus, more than half of the reproductive unions for which both maternal lineages could be assigned joined the two dominant matrilines at the site. These T2e1a1b–H1ao unions were not restricted to a single branch of the large pedigree, but occurred across several of the main clusters (cluster 1=1; cluster 2=7; cluster 3=8; cluster 5=5; cluster 6=4). The haplogroups of these pairings were approximately balanced between the sexes, with 12 out of 25 unions involving H1ao males and 13 involving T2e1a1b males. This indicates that the pattern is not simply driven by one maternal lineage consistently providing male or female reproductive partners.

To test whether the recurrent reproductive unions between the two dominant maternal lineages could be explained simply by their high frequencies in the cemetery, we compared the observed number of T2e1a1b–H1ao unions with a random expectation based on the frequency of these haplogroups among adult males and females at the site. Among adult individuals at Wetwang Slack, 35 of 134 males carried the main T2e1a1b haplotype and 31 carried H1ao, while 76 of 197 females carried T2e1a1b and 34 carried H1ao. Under random unions with respect to maternal lineage, the expected probability of a union between a T2e1a1b male and a H1ao female is therefore 4.5%. Among the 45 reproductive unions for which the mitochondrial haplogroups of both partners could be assigned, 13 (rather than the two expected under random pairing) involved a T2e1a1b male and an H1ao female: corresponding to 28.9% of informative unions. This is a 6.4-fold higher-than-expected ratio (one-sided binomial test: $p = 5.9 \times 10^{-8}$). We then tested the reciprocal combination. The expected probability of a union between an H1ao male and a T2e1a1b female is 8.9%. In the observed data, 12 (rather than the four expected under random pairing) of 45 informative reproductive unions involved an H1ao male and a T2e1a1b female, corresponding to 26.7%. This represents a 3-fold enrichment over expectation, (one-sided binomial test: $p = 0.0012$). Combining both types of pairing, the expected probability of any T2e1a1b–H1ao reproductive union is 13.4%, whereas the observed frequency is 55.6% (25/45). Thus, T2e1a1b–H1ao unions occurred 4.1 times more often than expected under random pairing based on adult haplogroup frequencies at the site (one-sided binomial test: $p = 3.2 \times 10^{-11}$).

These results show that the recurrent pairing between T2e1a1b and H1ao cannot be explained solely by the high frequencies of these two maternal lineages at Wetwang Slack. Instead, reproductive unions between the two dominant matrilines were strongly overrepresented, consistent with a socially structured pattern of recurrent alliance between maternal descent groups.

A particularly illustrative example is the family of I31005 in cluster 2 (Extended Data Figure 10a), where six consecutive generations of reproductive unions involved partners from the T2e1a1b and H1ao maternal lineages. Such repeated pairings across multiple generations are difficult to explain as random unions, especially given the reconstructed depth of the pedigree and the strong spatial and genealogical structure of the cemetery. Instead, they suggest that members of this community maintained detailed knowledge of maternal-line affiliation across many generations.

This recurrent association between the two dominant matrilines is compatible with a social structure in which the community was organized around two major maternal descent groups. Under such a model, repeated reproductive unions between T2e1a1b and H1ao individuals would reflect socially preferred or permitted pairings between complementary maternal groups.

Despite this recurrent pairing, the two maternal lineages do not appear to have played equivalent roles within the cemetery. T2e1a1b matrilines show a much deeper and more persistent presence at Wetwang Slack, continuing for up to 10 generations in one branch, nine generations in another and seven generations in another. In contrast, H1ao matrilines rarely persist for more than one or two consecutive generations within the cemetery. This asymmetry suggests that T2e1a1b formed the principal maternal core of the Wetwang Slack burial community, while H1ao and other lineages were repeatedly incorporated through reproductive unions but were less consistently maintained across generations.

Together, these results suggest that reproductive choices at Wetwang Slack were structured by maternal-line affiliation. The repeated pairing of T2e1a1b and H1ao across multiple branches of the pedigree points to an exceptional degree of genealogical memory. At the same time, the long-term persistence of T2e1a1b indicates that the cemetery was not simply organized around two equivalent maternal lineages, but around a dominant local matriline that repeatedly formed reproductive links with other major maternal lineages.

#### SI 7. Inter-site patterns of biological relatedness across the Arras dataset

##### IBD network analysis

*IBD network construction*

We constructed undirected networks of genetic relatedness based on pairwise sharing of IBD segments among individuals from the ten Arras Culture sites represented in the dataset (n=522), excluding those with evidence of contamination and those with fewer than 600,000 SNPs. Individuals were represented as nodes, and edges (connections) were defined between pairs sharing at least three IBD segments longer than 8 cM and a total of >24 cM in IBD. Edge weights correspond to the total IBD shared between individuals, providing a proxy for the degree of relatedness.

The Arras Culture network of all individuals (Extended Data Figure 2) comprises 22,028 edges and shows clear clustering by site, particularly for the three largest sites: Wetwang Slack, Pocklington and Melton 1. Pocklington and a large group of individuals from Wetwang Slack show particularly high connectivity, whereas individuals from Melton 1 appear more isolated within the network. Notably, however, a subset of Wetwang Slack individuals is positioned at the periphery of the network, with relatively few connections to other individuals.

To facilitate focusing on only the closest genealogical connections, we generated a restricted network of edges exceeding 150 cM in total shared IBD (Figure S61), thereby emphasising recent kinship ties of approximately fifth-degree or closer. In this refined network, site-specific clustering becomes more pronounced. Individuals from Melton 1 retain only three edges connecting them to other sites, while Pocklington and Wetwang Slack maintain 47 edges between them, indicating substantial mobility between these two communities within approximately three to five generations prior to  creation of the cemeteries. To explore sex- and age-specific patterns, we additionally constructed subset networks including only adult males or only adult females (Figure S62). The adult male and adult female networks are broadly similar, but with closer connections (edges with darker colour, higher IBD shared) in the female network.

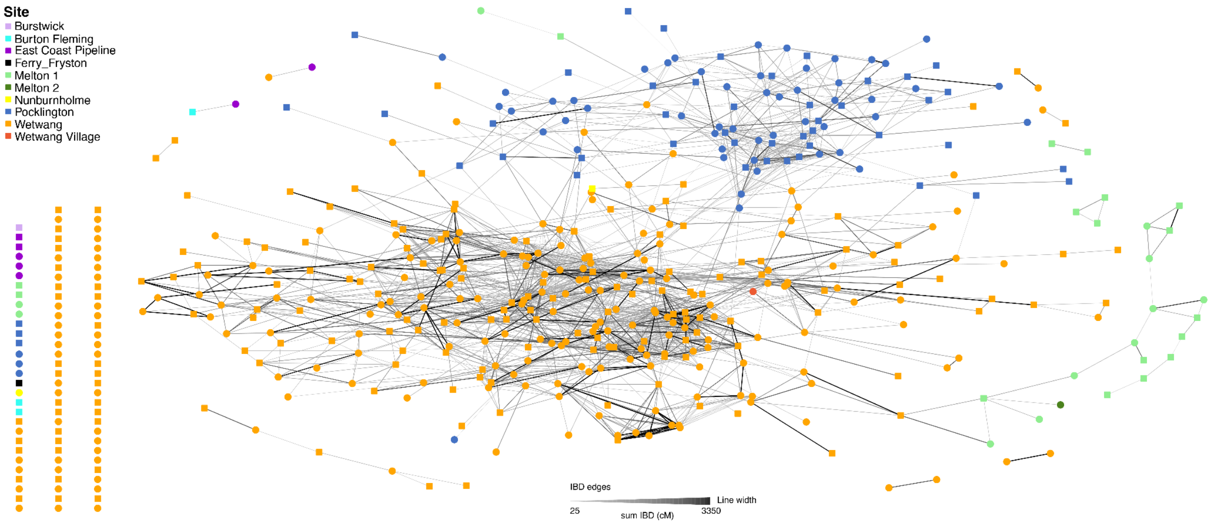


Figure S61. IBD network of the Arras Culture sites represented in the dataset, featuring approximately fifth-degree or closer biological relatedness. Edges connect individuals sharing more than two IBD segments of >8 cM and a total of >150 cM in IBD (approximately fifth-degree or closer relationships), and are weighted by the total IBD shared (sum_IBD). Node colour denotes archaeological site and shape indicates genetic sex: squares indicate males and circles indicate females. Node positions are determined using a force-directed layout (Fruchterman–Reingold). Isolated individuals without connections meeting the threshold are shown separately (left).

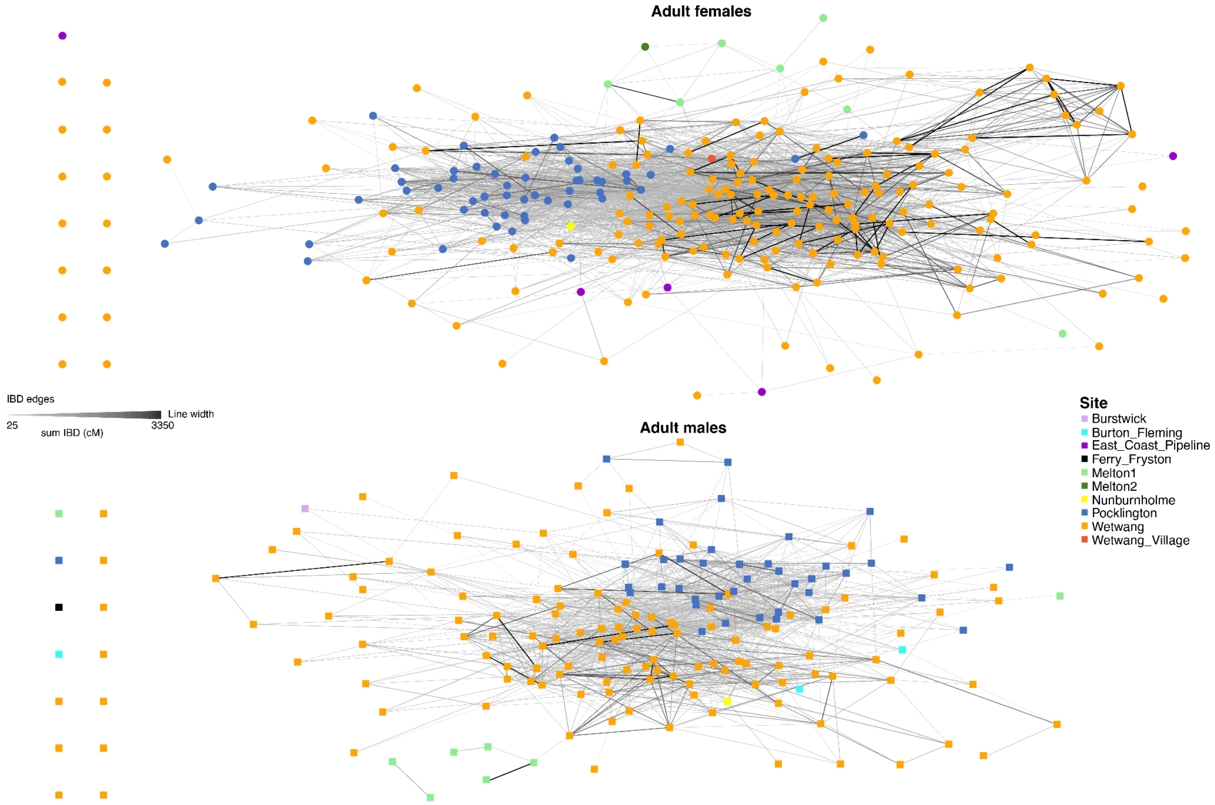


Figure S62. IBD network of Arras Culture sites represented in the dataset for adult females (top) and adult males (bottom) only.

*Network metrics*

For each individual, we computed two node-level measures of connectivity. Degree centrality ($k$) was defined as the number of IBD connections per node, and strength ($w$) as the sum of edge weights (total IBD shared) across all connections. For analyses restricted to adult individuals, degree and strength were recalculated considering only connections between adults, excluding adult–non-adult links; this avoided potential biases due to incomplete life (i.e. reproductive) histories for pre-pubescent individuals.

*Sex-specific patterns of connectivity and relatedness*

To compare connectivity patterns between sexes, we analysed the empirical survival distributions of degree and strength, defined as $P(X>x)$, where X corresponds to $k$ or $w$. Survival functions were computed separately for male and female individuals, as well as for the combined population. Differences between male and female distributions were quantified using the two-sample Kolmogorov–Smirnov (KS) D statistic, which measures the maximum deviation between cumulative distributions. Statistical significance was assessed using a permutation framework: sex labels were randomly reassigned among individuals while preserving the network structure, and the KS statistic was recalculated for each permutation (10,000 replicates). Empirical p-values were obtained as the proportion of permutations yielding KS values equal to or greater than the observed statistic.

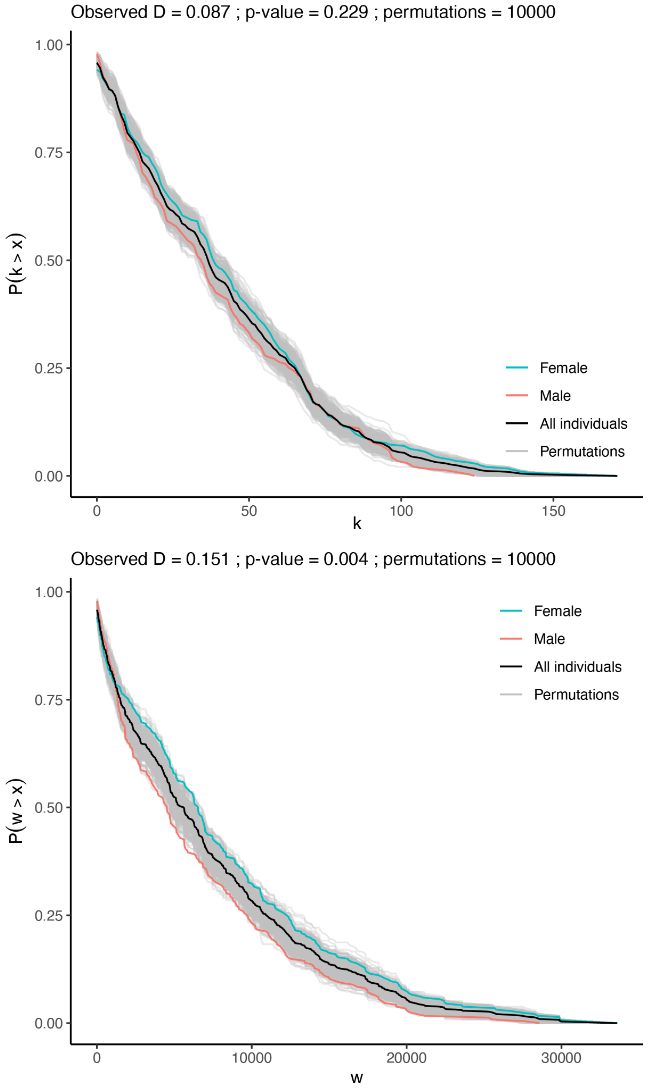


Figure S63. Sex-specific differences in network connectivity and relatedness in the entire IBD network for the Arras Culture sites represented in the dataset. The survival distributions of node degree (*k*: number of connections per individual) and strength (*w*: sum of total IBD shared in all edges per individual) are shown for females (blue), males (red), and all individuals (black). Grey curves represent the null expectation under random assignment of sex labels. The Kolmogorov–Smirnov statistic (D) and empirical p-value assess whether the observed differences between sexes exceed those expected under random assignment. The threshold for defining edges is more than two IBD segments of >8 cM and a total of >24 cM in shared IBD.

In the full Arras Culture network, the survival distributions of strength (*w*) differ significantly between males (n=235) and females (n=287) (D=0.15; P-value =0.0045), with females showing consistently higher cumulative IBD sharing (Figure S63). In contrast, the distributions of degree (*k*) do not differ significantly between sexes (Table S4). These results indicate that, although females tend to accumulate greater total genetic relatedness within the network (mean *w*: females=8.14 cM versus males=6.31 cM), the number of connections per individual is broadly comparable between sexes. This pattern is replicated when restricting the analysis to the adult-only network (n=442) and to Wetwang Slack alone (n=378) (Table S4). In both cases, strength remains significantly higher in females than in males, whereas degree does not differ significantly between sexes. This consistency suggests that sex-biased patterns of relatedness are present within the site with the higher sample size (i.e. Wetwang Slack) and persist at the scale of the full Arras Culture network.

Table S4. Comparing connectivity patterns between sexes in different subsets of the Arras Culture network

| | **Arras** | | **Wetwang Slack** | | **Wetwang Slack Clusters 3-5** | |
|---|---|---|---|---|---|---|
| | **n IBD>=3; sum IBD>24** | | **n IBD>=3; sum IBD>24** | | **n IBD>=3; sum IBD>150** | |
| | **All network** | **Adults only** | **All network** | **Adults only** | **All network** | **Adults only** |
| **n** | 523 | 442 | 378 | 321 | 89 | 75 |
| **n males** | 235 | 182 | 164 | 129 | 35 | 26 |
| **n females** | 287 | 260 | 214 | 192 | 54 | 49 |
| **Males mean k** | 40 | 37 | 33 | 30 | 9 | 7 |
| **Males median k** | 35 | 32 | 29 | 26 | 9 | 7 |
| **Males mean w** | 6311 | 6170 | 6639 | 6352 | 7526 | 6772 |
| **Males median w** | 4525 | 3884 | 4550 | 3825 | 7493 | 6825 |
| **Females mean k** | 44 | 40 | 36 | 32 | 13 | 11 |
| **Females median k** | 39 | 36 | 34 | 30 | 11 | 9 |
| **Females mean w** | 8136 | 7664 | 8641 | 8200 | 11369 | 10747 |
| **Females median w** | 6535 | 6001 | 7091 | 6855 | 10225 | 8817 |
| **Kolmogorov Smirnov D k** | 0.087 | 0.056 | 0.067 | 0.059 | 0.184 | 0.272 |
| **P-value k** | 0.229 | 0.812 | 0.699 | 0.883 | 0.296 | 0.078 |
| **Kolmogorov Smirnov D w** | 0.151 | 0.143 | 0.168 | 0.157 | 0.325 | 0.307 |
| **P-value w** | 0.004 | 0.020 | 0.009 | 0.038 | 0.015 | 0.062 |

The absence of significant differences in degree centrality across sexes (Table S4) can be explained within the matrilineally focused cemetery organisation observed. Although several families, most clearly at Wetwang Slack, exhibit clear matrilineal organisation—where successive generations are structured around female lineages (e.g. clusters 3 and 5; Figures S20 and S28)—adult males within these clusters often represent the biological sons of the lineage (e.g. I31512, I36972, I36909) and therefore possess a high number of connections. Additionally, some adult males who enter these groups through marriage or mobility (e.g. I31507, I37138) also have relatives buried within the cemetery beyond their immediate descendants, further increasing their connectivity. When analyses are restricted to clusters 3 and 5 (n=89), which exhibit strong matrilineal organisation, females display higher degree values (mean=13) than males (mean=9), consistent with their central role in structuring the kinship networks in these families. However, these differences do not reach statistical significance in the KS framework (Table S4), likely due to the presence of highly connected males within these clusters.

#### IBD sharing across Iron Age Britain

To explore genealogical connections across Iron Age Britain, we quantified IBD sharing between Middle and Late Iron Age sites with high-quality data (Supplementary Table 5). For each pair of sites (including intra-site comparisons), we computed the fraction of individual pairs sharing IBD relative to the total number of comparable pairs (Extended Data Figure 2b). As we are interested in connections across Britain, which are likely more distant than those within the Arras Culture sites under study, we applied a more permissive threshold than in previous analyses: pairs were classified as sharing IBD if they exhibited at least one segment of >12 cM. To avoid over-representation of close kin, we retained only one individual per cluster of first-degree relatives. In parallel, we assessed strictly matrilineal connections by computing the fraction of individual pairs sharing the same mitochondrial haplotype relative to the total number of comparable pairs (Extended Data Figure 2c).

Arras Culture sites displayed a high level of IBD sharing, particularly Wetwang Slack, Wetwang Village, Pocklington, Burton Fleming and Nunburnholme, where the fraction of pairs sharing IBD remained above 0.16 in all pairwise comparisons (Extended Data Figure 2b). This pattern indicates that these cemeteries were not used by isolated communities but instead formed part of a tightly interconnected regional network, within which individuals maintained genealogical ties across multiple sites. The strength and consistency of these connections suggest sustained interaction over several generations, rather than sporadic movement. This high degree of connection is not, however, mirrored in the mtDNA. In this analysis, the fraction of pairs sharing mtDNA haplotypes never exceeds 0.005 (Extended Data Figure 2c), with the only exception of the comparison between Burton Fleming and East Coast Pipeline (at 0.048) (and Melton 1-Melton 2 and Wetwang Slack-Wetwang Village, but in these pairs this reflects their close spatial proximity ~1 km). In contrast, intra-site mtDNA sharing remains high, exceeding 0.11 in all Arras Culture sites with more than five individuals.

Together, these patterns indicate that most genealogical connections across Arras Culture sites are not mediated through strict maternal lines. While this suggests that maternal links are not the primary driver of inter-site connectivity, they appear to play an important role within sites. For example, at Wetwang Slack, the fraction of pairs sharing IBD (0.24) is nearly identical to that observed between Wetwang Slack and Pocklington (0.25), highlighting the strong connectivity between these sites, as well as the presence of individuals at Wetwang Slack with few or no detectable links to either site. In contrast, the fraction of mtDNA matches between Wetwang Slack and Pocklington is extremely low (0.002), reinforcing the limited contribution of maternal connections to the intense IBD sharing. This supports the hypothesis that many of the connections were forged through a practice of male exogamy, although individual cases—such as female I39729 from Wetwang Slack who most likely represents a recent arrival from Pocklington—may reflect recent female movement between sites.

Arras Culture sites display a very low degree of IBD sharing with other Middle and Late Iron Age sites in Britain, with the exception of the (relatively) nearby site of Wattle Syke in West Yorkshire, which displays elevated IBD sharing with Melton 2 and Wetwang Village.

#### SI 8. Ancestry analysis

We used *qpAdm* to study the ancestry make-up of individuals buried in the Arras Culture sites represented in the dataset. Following Patterson *et al.* 2021[23], we modelled the ancestry of each individual as a mixture of Western Hunter-Gatherer (WHG), Early European Farmer (EEF) and Steppe Early Bronze Age sources. In the outgroup set we included four populations: ancient individuals from Cameroon (Old Africa), Afanasievo individuals from Russia (Russia_Afanasievo), Neolithic individuals from Anatolia (Anatolia Neolithic), and Hunter-Gatherer individuals from the Iron Gates region in south-eastern Europe (IronGates_HG). In source and outgroup populations, only individuals with shotgun or Twist capture data were included (Supplementary Table 6). Arras Culture individuals labelled as FAIL or QUESTIONABLE were, however, excluded. The model provided a good fit for the vast majority of individuals, with only 14 out of 521 individuals with p-values lower than 0.01 (Supplementary Table 7). One individual from Wetwang Slack (I36776) was a clear outlier with higher (47%) EEF ancestry and lacked relatives both at Wetwang Slack and other Arras Culture sites.

We compared the distribution of EEF ancestry proportions across Arras Culture sites, keeping only individuals with p-values>0.01 and, to avoid over-representation of close relatives, keeping only one representative per cluster of first-degree relatives (Extended Data Figure 3a). All sites with more than five individuals displayed mean values of 37–38%, with no significant differences between them (Welch's *t*-test, two-tailed; $p > 0.05$). This analysis demonstrates that the ancestry of individuals buried at Arras cemeteries was highly homogeneous.

Within the context of Middle–Late Iron Age Britain, EEF ancestry displayed a significant negative correlation with latitude (Figure S64). As such, Arras Culture cemeteries had intermediate EEF values, higher than Scottish sites but lower than sites in southern England, likely due to the stronger continental connectivity of southern England sites[20].

Based on material culture similarities with La Tène cemeteries in France,— notably the square-ditched barrows and the chariot burials—with La Tène cemeteries in France, the East Yorkshire Arras Culture has been hypothesised to have a recent origin on the Continent[29]. Ancestry analysis does not support a recent origin of Yorkshire Arras Culture individuals on the Continent. Individuals in the Iron Age France group (24 individuals; Supplementary Table 2) not only display higher (45%) EEF ancestry, but a very poor fitting model (P-value <0.001) was produced when Iron Age France was used as the only source of ancestry for Yorkshire Arras Culture individuals (Supplementary Table 7). Neither did we find any indication of increased IBD sharing (Supplementary Table 5) when computing the fraction of individual pairs sharing IBD between Iron Age France and Arras Culture cemeteries.

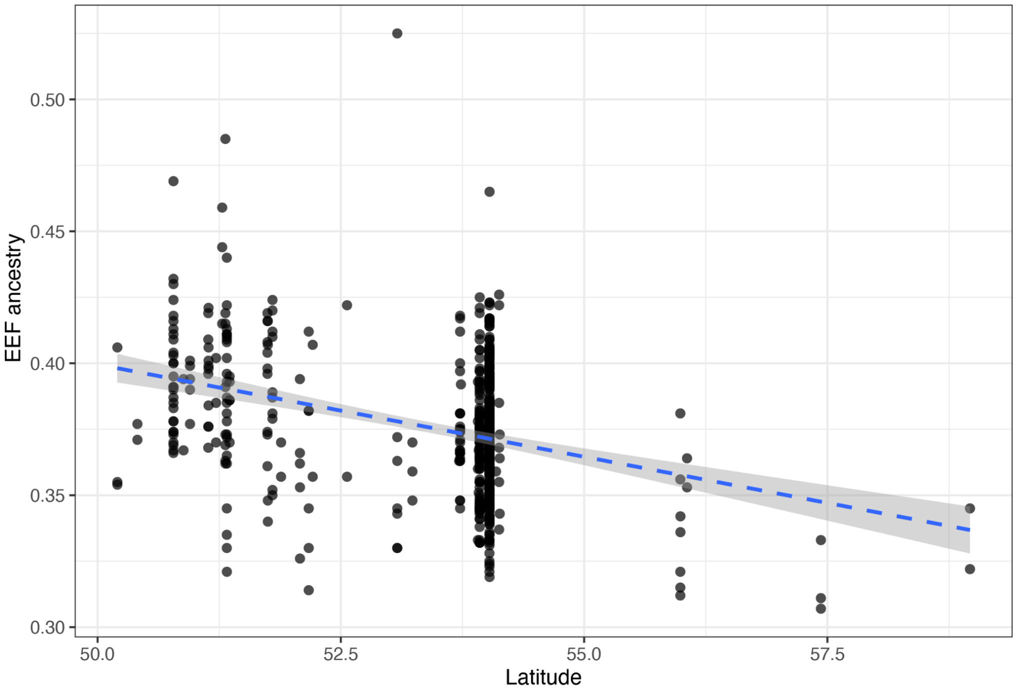


Figure S64. Relationship between EEF ancestry proportion and latitude in Middle–Late Iron Age individuals from Britain. Each point represents one individual, with ancestry proportions estimated using *qpAdm*. The dashed line shows a linear regression fit with 95% confidence interval (shaded area), displayed for visualisation purposes. Statistical association was assessed using Spearman's rank correlation ($\rho = -0.218$, *p-value* = $8.75 \times 10^{-8}$; n=591), indicating a weak but statistically significant decrease in EEF ancestry with increasing latitude.

##### *EEF ancestry proportions across burial types*

At Pocklington and Wetwang Slack, we tested whether individuals interred in different burial types displayed differences in ancestry proportions (Extended Data Figure 3b). We found no statistically significant differences.

### Legends of Supplementary Tables

**Supplementary Table 1**. Ancient individuals from Arras Culture cemeteries (both published and unpublished) included in this study.

**Supplementary Table 2**. Previously published ancient individuals from other cultural contexts used in this study.

**Supplementary Table 3**. Mitochondrial and Y-chromosome haplotype diversities for archaeological Neolithic, Bronze Age and Iron Age sites in Britain.

**Supplementary Table 4**. Pairwise kinship statistics for Arras Culture sites.

**Supplementary Table 5**. IBD sharing between Middle and Late Iron Age sites in Britain and and French Iron Age sites in aggregate. Fraction of individual pairs sharing IBD relative to the total number of comparable pairs. Pairs were classified as sharing IBD if they exhibited at least one segment of >12 cM.

**Supplementary Table 6**. Individuals included in source and outgroup populations for *qpAdm* analysis.

**Supplementary Table 7**. *qpAdm* results for individuals from Arras Culture cemeteries.

**Supplementary Table 8**. Tests for the difference in grave distances between relatives and other individuals.

**Supplementary Table 9**. Sex bias analysis for different groupings.

**Supplementary Table 10**. Counts of individuals with no parent, one parent and two parents in the same cemetery, and count of reproductive partners with both members, only male and only female present in the same cemetery.

**Supplementary Table 11**. Number of matrilineages, patrilineages and bilineages for the three main cemeteries, Wetwang Slack, Pocklington and Melton 1.

**Supplementary Table 12**. At the three main cemeteries: Wetwang Slack, Pocklington and Melton 1, number of adult males and females with first- or second-degree relatives in earlier generations.

**Supplementary Table 13**. Number of males and females with multiple reproductive partners at the three main cemeteries, Wetwang Slack, Pocklington and Melton 1.

**Supplementary Table 14**. For reproductive partners entering the reconstructed pedigrees at Wetwang Slack, count of identified descendants in the pedigree and total number of moderately close intra-site relatives.

**Supplementary Table 15**. Mitochondrial haplogroups of reproductive partners at Wetwang Slack.
